# Anomalous phylogenetic behavior of ribosomal proteins in metagenome assembled genomes

**DOI:** 10.1101/731091

**Authors:** Sriram G. Garg, Nils Kapust, Weili Lin, Fernando D. K. Tria, Shijulal Nelson-Sathi, Sven B. Gould, Lu Fan, Ruixin Zhu, Chuanlun Zhang, William F. Martin

## Abstract

Metagenomic studies have claimed the existence of novel lineages with unprecedented properties never before observed in prokaryotes. Such lineages include Asgard archaea^1–3^, which are purported to represent archaea with eukaryotic cell complexity, and the Candidate Phyla Radiation (CPR), a novel domain level taxon erected solely on the basis of metagenomic data^4^. However, it has escaped the attention of most biologists that these metagenomic sequences are not assembled into genomes by sequence overlap, as for cultured archaea and bacteria. Instead, short contigs are sorted into computer files by a process called binning in which they receive taxonomic assignment on the basis of sequence properties like GC content, dinucleotide frequencies, and stoichiometric co-occurrence across samples. Consequently, they are not genome sequences as we know them, reflecting the gene content of real organisms. Rather they are metagenome assembled genomes (MAGs). Debates that Asgard data are contaminated with individual eukaryotic sequences^5–7^ are overshadowed by the more pressing issue that no evidence exists to indicate that any sequences in binned Asgard MAGs actually stem from the same chromosome, as opposed to simply stemming from the same environment. Here we show that Asgard and CPR MAGs fail spectacularly to meet the most basic phylogenetic criterion^8^ fulfilled by genome sequences of all cultured prokaryotes investigated to date: the ribosomal proteins of Asgard and CPR MAGs do not share common evolutionary histories. Their phylogenetic behavior is anomalous to a degree never observed with genomes of real organisms. CPR and Asgard MAGs are binning artefacts, assembled from environments where up to 90% of the DNA is from dead cells^9–12^. Asgard and CPR MAGs are unnatural constructs, genome-like patchworks of genes that have been stitched together into computer files by binning.

---

The sequencing of environmental DNA (metagenomics) has become an essential tool of modern science because it allows microbiologists to uncover the existence of genes and species in environments such as marine sediment or the deep biosphere from which representatives cannot readily be cultured^13,14^. Initially an endeavor involving rRNA sequences^15^, metagenomic investigations have led to the binning-assembly and deposition in databases of MAGs, sequences of similar GC content and stoichiometry across samples. Because rRNA has limited phylogenetic resolution, concatenated sequences of ribosomal proteins (r-proteins) and other universally distributed proteins are commonly used for phylogeny. This practice is well established with over 20 years of tradition, whereby the validity of using concatenated r-proteins for phylogeny lies in the reproducible crosschecking result that individual r-proteins from the same sequenced genome give the same or very similar trees^16–21^. Based in such precedence, it became common practice to use concatenated r-proteins from sequenced genomes for microbial phylogeny without first crosschecking whether the individual proteins gave similar trees. That no-crosscheck practice has however been blindly extended to MAGs, which is unfounded because there is no independent evidence from overlapping genome assembly procedures that the 20-30 r-proteins used for phylogeny in a given MAG are encoded in one and the same genome. They could easily stem from DNA fragments of 20-30 different genomes that simply occur in the same environment. This problem is exacerbated by the circumstance that up to 90% of sequenceable DNA in some environments such as marine sediment is not packaged in cells but is extracellular DNA (eDNA) from dead cells or biofilms^9–12^, whereby the proportion of eDNA varies across environments and changes over geological time^13–14^.

Given the unprecedented nature of claims regarding complex archaea in Asgard^5,6^ and the existence of the CPR, methods to validate phylogenies based on universal and r-proteins from MAGs are needed. We turned to a fundamental principle known from the earliest days of phylogenetic testing: proteins with a shared evolutionary history should generate the same or similar trees^8^. For genomic DNA from cultivated prokaryotic organisms, hereafter referred to as organismal DNA (orgDNA) it is known that different r-proteins produce the same tree or very similar trees^16–21^. Yet even for r-proteins that share a common evolutionary history, their trees will differ to some extent owing to practical and theoretical limitations of phylogenetic inference^16–23^. The extent of natural variation across r-protein trees for orgDNA can however be determined empirically by simply comparing the r-protein trees for a given genome set. Using that natural distribution as a reference, we can then ask whether MAGs fulfill the same criterion, that is, do trees for r-proteins from MAGs resemble each other to the same or to a lesser degree than trees based on orgDNA, and if they differ, is the difference significant?

We assembled alignments of proteins from orgDNA and from MAGs (see Methods) that are universal or nearly so among archaeal genomes. That is the procedure underpinning the taxon assignment in the phylogenetic trees that purport to describe the relationship of MAGs from Asgard samples to orgDNA^1,7^. We first generated alignments for 39 different proteins that are present in all members of a large and diverse sample of archaeal orgDNA and archaeal MAGs. For orgDNA, we generated phylogenetic trees for each alignment individually and asked how similar the trees are to one another using the simple but robust Robison-Foulds pairwise distance measure ^24^ (Fig. 1a). We asked the same question for the same proteins using archaeal MAGs. The result (Fig. 1b) shows that in phylogenetic terms, archaeal MAGs behave in a fundamentally different manner from orgDNA in two respects. First, orgDNA trees are much more similar to one another than MAG trees are. Second, the 23 ribosomal proteins (including secY, which is ribosomal for co-translational insertion) and 16 other proteins universally distributed within the genome sample show no difference in their ability to recover approximately the same tree for orgDNA but the same is not true for archaeal MAGs (light shading in Fig. 1b, see scale bar). This distinctly bimodal distribution of phylogenetic behavior for the 39 Asgard MAG proteins is not observed for orgDNA.

**Fig. 1.**
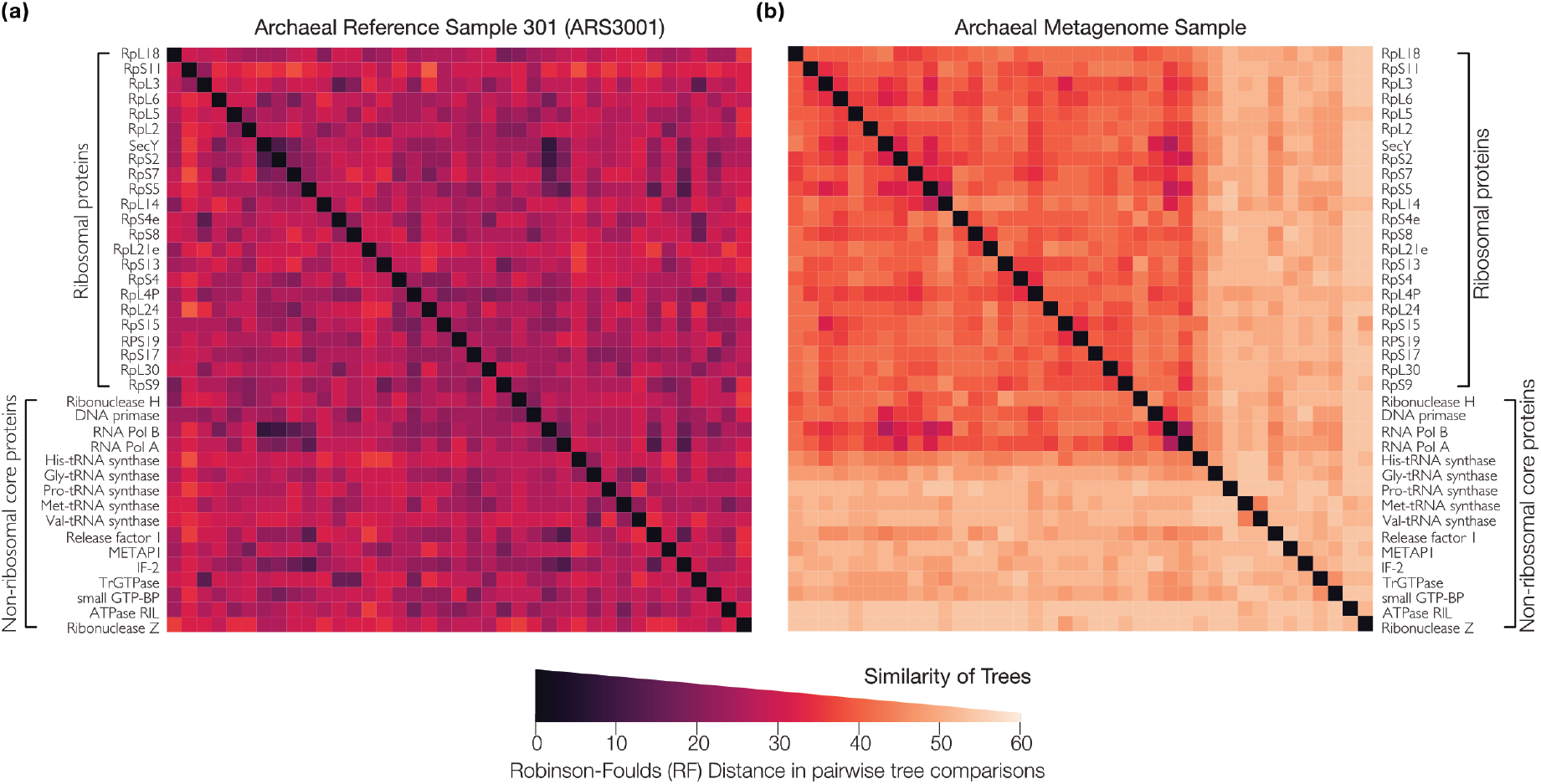
Pairwise Robinson-Foulds distance between trees for universal archaeal proteins. Pairwise distance between phylogenetic trees of 39 universal proteins were calculated using the Robinson-Foulds metric and plotted in **(a)** for a sample of 30 archaeal genomes from RefSeq and **(b)** for a sample of 30 archaeal MAGs. The differences among trees for these archaeal proteins in RefSeq reflects the natural variation for sequenced genomes from cultured archaea. The archaeal MAGs, while having a lower degree of congruence between the trees overall, cluster into two major discernable groups with one composed largely of ribosomal proteins. It is evident that r-protein trees in archaeal MAGs are more similar to each other than trees for non-ribosomal proteins. The scale bar applies to both panels.

To quantify these differences, we plotted the distribution of pairwise similarity across 39 trees for ten different samples of 30 genomes each from RefSeq archaea (Supplementary Fig. S1) and 30 archaeal MAGs. The distribution for archaeal MAGs, which include Lokiarchaeum and Asgard archaea, is distinctly bimodal using four different tree comparison methods (Fig. 2a). Note that the distributions for trees from archaeal MAGs are always shifted towards higher pairwise distances between trees. We plotted the curves for the 23 ribosomal proteins against the 16 other universal proteins (including RNaseH, DNA primase, polA and polB) (Fig. 2b, Fig. 2c). The phylogenetic behavior of archaeal MAGs is obviously and significantly different from archaeal orgDNA. The distribution of tree dissimilarity scores across r-proteins from MAGs is significantly larger than the corresponding values for orgDNA, with p-values ranging from 10^−26^ to 10^−139^ (two-tailed Kolmogorov-Smirnov test) depending on the tree comparison metric (Fig. 2b) (see Methods and Supplementary Table S3b). The probability that the distribution of tree dissimilarities across non-ribosomal proteins from MAGs is drawn from the same distribution as the corresponding value for orgDNA ranges from 10^−48^ to 10^−66^, depending on the tree comparison metric (see Methods and Supplementary Table S1c).

**Fig. 2.**
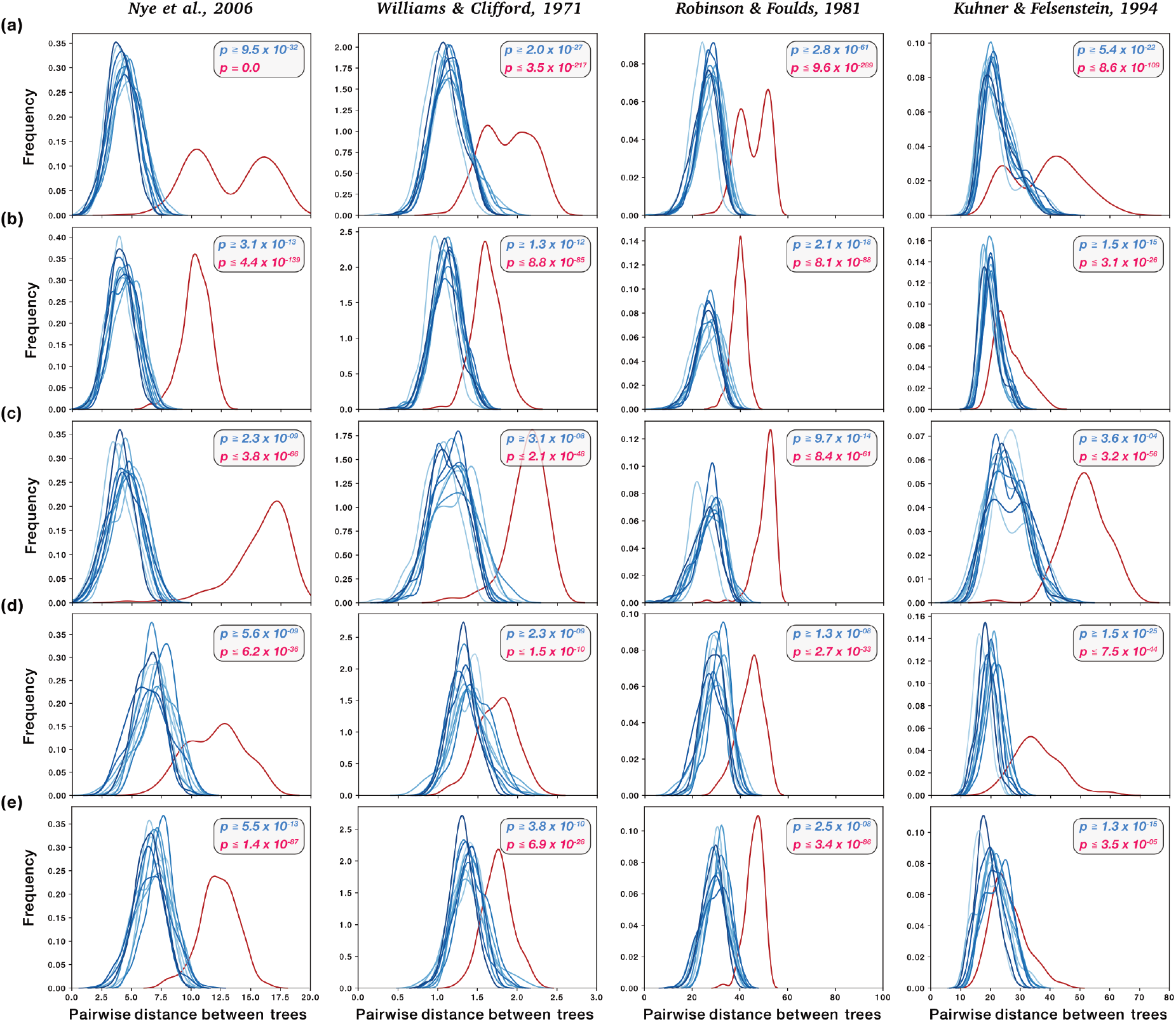
Distribution of pairwise tree-distances between RefSeq and metagenomes: The pairwise comparisons of tree distances computed using four different metrics (see Methods) are shown. In each case, a matched set of proteins present in MAGs and 10 random samples from RefSeq are taken to plot comparable distributions. The MAG sample is always shown in red. **(a)** Trees for 39 universal proteins from 30 archaeal MAGs are compared with trees for the 39 homologues from 10 samples of 30 archaeal RefSeq genomes **(b)** Trees for 23 ribosomal proteins from 30 archaeal MAGs are compared with those from 10 samples of 30 archaeal RefSeq genomes. Note that the mean topological distance between trees is higher for the MAGs when compared with RefSeq genomes. **(c)** Trees for 16 non-ribosomal proteins from 30 archaeal MAGs are compared with those from 10 samples of 30 archaeal RefSeq genomes. **(d)** Trees for 16 ribosomal proteins from 30 candidate phyla radiation CPR MAGs from ref. 4 are compared with those from 10 samples of 30 bacterial RefSeq genomes. **(e)** Trees for 20 ribosomal proteins from non-CPR 30 bacterial MAGs are compared with those from 10 samples of 30 RefSeq genomes. In all panels, blue curves represent the 10 independent reference samples while the red curve represents the MAGs. The largest FDR corrected p-value (two-tailed Kolmogorow-Smirnow test) from comparisons between the MAG sample and each RefSeq sample is indicated in red while the smallest p-value is indicated in blue (see Methods).

This effect is not due to greater phylogenetic depth or greater sequence divergence of ribosomal proteins in archaeal MAGs relative to orgDNA, as shown by the distributions of pairwise uncorrected p-distances for Asgard MAGs and orgDNA for each protein in the sample (Supplementary Fig. S4, S5). Many phylogenetic analyses involving MAG data employ site-filtering procedures to remove sites from the sequence alignment^1,2,17,25,26^. To check whether site filtering affects the phylogenetic anomaly of Asgard MAGs, we trimmed alignments (see Methods) as in earlier analyses^1,2^ and recalculated the trees and comparisons. As shown in Extended data 1, site filtering does not improve MAG phylogenetic behaviour. The Asgard MAGs have a severe, systematic and previously undetected phylogenetic anomaly.

Is the MAG phylogenetic anomaly archaeal specific? For bacteria, we investigated whether MAG bins from the Candidate Phyla Radiation (CPR) group showed the same effect. The probability that the distribution of tree similarities across 16 ribosomal proteins from CPR MAGs is drawn from the same distribution as the corresponding value for bacterial orgDNA ranges from 10^−10^ to 10^−44^, depending on the tree comparison metric (Fig. 2d). For bacteria there are more metagenomes available than for archaea. We obtained metagenomic assemblies for bacteria from RefSeq that do not belong to the CPR and applied the same test. Similar to archaeal MAGs and CPR MAGs, for bacterial MAGs that do not stem from the CPR, the distribution of tree similarities across r-proteins from MAGs is different from bacterial orgDNA for the four tree comparison metrics when compared to the reference (Fig. 2e).

To see if all MAGs reveal the same binning problem, we repeated the tree comparison metrics on 10 additional samples each of non-asgard archaeal MAGs, CPR MAGs and non-CPR bacterial MAGs (Supplementary Fig. S7). We find that archaeal MAG r-protein samples from the Tara ocean genome project generated by binning methods are not fundamentally anomalous in their phylogenetic behavior, although trees for non-r-proteins from the Tara data are significantly different (Supplementary Fig. S37). The data behind the two most spectacular phylogenetic claims reported so far from metagenomics — the Asgard and CPR MAGs — behave in an unnatural manner (Fig. 1, Fig. 2, Extended data 1, Supplementary Fig. S7).

In the worst case, the phylogenies generated by MAGs could share no better than random similarity, reflecting trees of proteins encoded by DNA of similar sequence properties but of unlinked phylogenetic histories. To visualize the effect of randomization we employed a standard network method, Neighbor-Net (NNet)^27^. The NNet for r-proteins sampling crenarchaeal and euryarchaeal orgDNA is extremely tree-like, with 16 strong, low conflict splits indicated in the figure (Fig. 3a), reflecting highly congruent phylogenetic signals across r-proteins from diverse cultivated archaea (Fig. 1a, Fig. 2b). An NNet for Asgard archaeal MAGs is shown in Fig. 3b, revealing one split in the data, that separating Thorachaea MAGs from the rest. The phylogenetic congruence across r-proteins in the remaining 21 archaeal MAGs is not different from random, as shown by randomizing the source of sequences in individual ribosomal protein alignments, used for concatenation and NNet plots (Extended Data 2). NNets for bacterial orgDNA from RefSeq comapred to CPR MAGs reveal the same result (Extended Data 3).

**Fig. 3.**
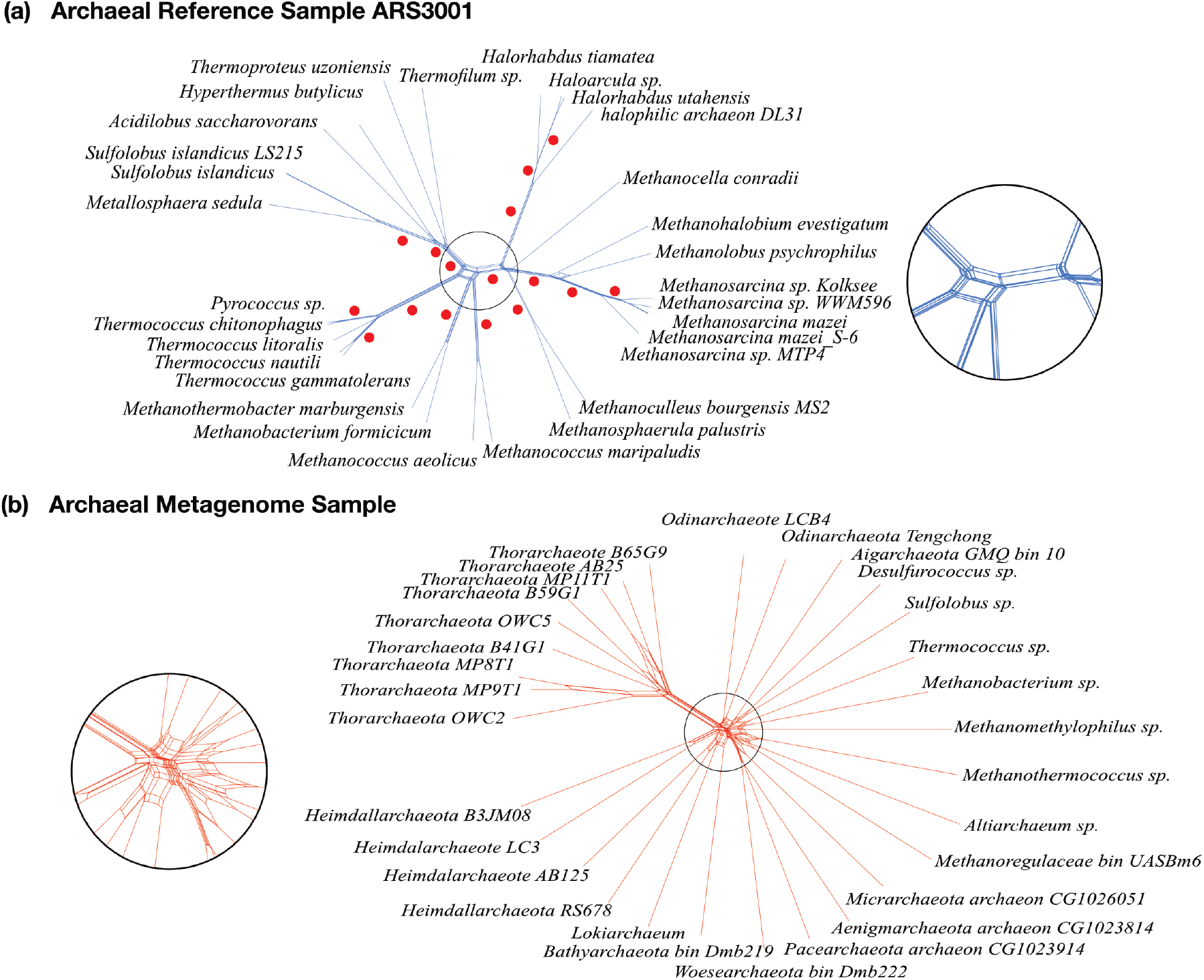
Neighbor-Nets reconstructed from concatenated alignments of 23 ribosomal proteins for archaeal reference samples and archaeal MAGs. **(a)** The Neighbor-Net of a concatenated alignment of 23 ribosomal proteins in the archaeal reference sample ARS3001 shows very little conflict throughout, resulting in a tree-like network with 16 well supported splits (indicated with red dots). **(b)** A Neighbor net drawn from a concatenated alignment of the same 23 ribosomal proteins from Asgard archaeal MAGs results in a network with a star-like structure. The insets magnify the central area of interest to better highlight the difference of signals of the two networks.

As a further independent test for tree similarity, we examined the phylogenetic compatibility within MAG and orgDNA tree sets (see Methods) by scoring the compatibility of each r-protein tree against all other trees from the same set^28^. Again, r-proteins from genomes of cultured organisms produced trees that are very similar to each other, as they should be^16–21^, while trees for r-proteins from MAGs were different from each other to an extent that is not observed for unbinned data (Fig. 4, Supplementary Fig. S8). The differences in cumulative distribution frequencies are obvious (Fig. 4) and highly significant (p = 10 ^−18^, two-tailed Kolmogorov-Smirnov test).

**Fig. 4.**
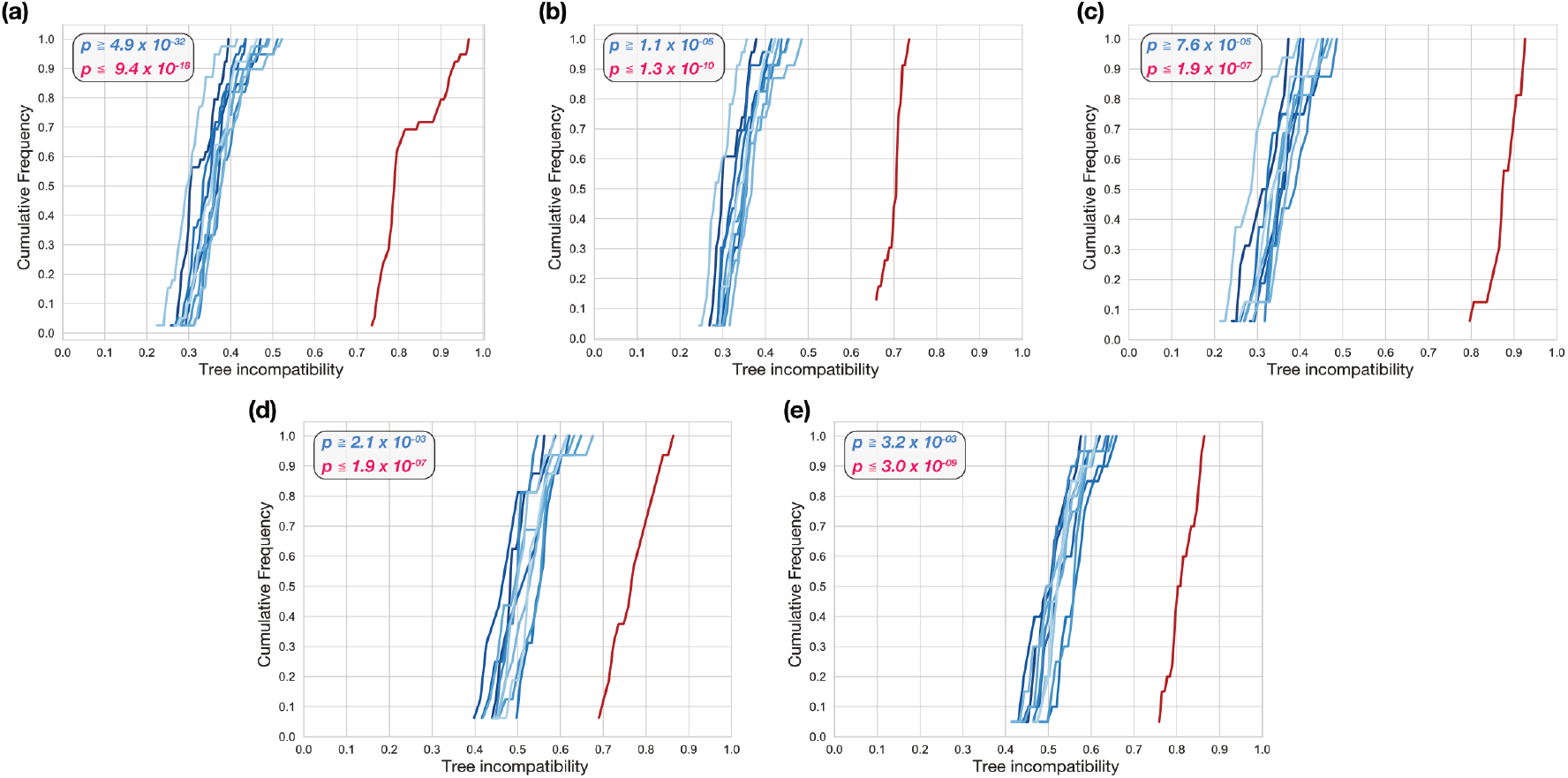
Tree compatibility scores for samples of tree reconstructed from orgDNA and metDNA. Cumulative distribution of tree incompatibility scores within sets of gene trees. In each case every curve represents a set of 30 organisms where the RefSeq samples are shown in shades of blue and the MAG sample is always shown in red **(a)** Trees for 39 universal proteins sampled from 10 archaeal RefSeq genomes vs Asgard MAGs, **(b)** Trees for a subset of 23 ribosomal proteins sampled from 10 archaeal RefSeq genomes vs Asgard MAGs **(c)** Trees for the complement set of 16 non-ribosomal proteins sampled from 10 archaeal RefSeq genomes vs Asgard MAGs. **(d)** Trees for 16 ribosomal proteins sampled from 10 bacterial RefSeq genomes vs CPR MAGs and **(e)** Trees for 20 ribosomal proteins sampled from 10 bacterial RefSeq genomes vs non-CPR bacterial MAGs.

Our analyses reveal that the new archaeal MAGs called Lokiarchaea and Asgard archaea, as well as the new CPR lineages, which are all represented by binned MAGs only, are binning artefacts for the proteins that — more so than any others known — should generate similar trees for prokaryotic lineages. Crucially, the name and rank of Asgard archaea and CPR taxa are defined by their r-protein phylogenies. These r-proteins are, however, not linked by common ancestry rather they are stitched together from sequences in same environment into computational bins that are assigned an organism-like taxonomic label. For non-ribosomal proteins the problem is much more severe than for r-proteins themselves. This has gone hitherto unnoticed because no one has tested whether the MAGs show natural phylogenetic properties like sequences from closed genomes do. A closed genome was recently reported for an anaerobic fermenting archaeon from sediment (sister to Crenarchaeoata termed Candidatus *Prometheoarchaeum syntrophicum* strain MK-D1) that branches as the sister to eukaryotes in the absence of MAG data; it is small (0.5 μm diameter), lacking endomembrane system, and not phagocytosing^29^. There have been challenges posed to individual protein sequences present in Asgard MAGs^5,6,30^. Our findings reveal that the Asgard data problem is deeply rooted and is furthermore systematic. There are real organisms in sediment^29^, but Asgard and CPR MAGs do not represent real organisms.

## Methods

### Identification of homologs of ribosomal proteins in RefSeq genomes

Ribosomal protein clusters for Archaea and Bacteria were retrieved from UniProt (Jan 2019). These clusters were used for a BLAST against the RefSeq 2016 database consisting of 5443 bacteria and 212 archaea with an identity threshold of 25% and an e-value cut-off of 10^−5^. The RefSeq dataset was further subsampled for groups of 30 organisms that reflect the full breadth of taxonomic distribution for the complete dataset for bacteria and archaea, respectively, to generate 10 reference samples each with 30 species or operational taxonomic units (OTUs) for archaea and 10 reference samples each with 30 species (OTUs) for bacteria. That is, the reference orgDNA samples were chosen to sample as much phylogenetic diversity, depth and breadth as prokaryotic RefSeq data have to offer (for more information see Supplementary Table 1 Supplementary Table 2; Supplementary Fig. S1; Supplementary Fig. S2).

### Phylogenetic tree construction

For each group (Archaea, Bacteria and CPR) matching sets of ribosomal proteins for each sample were chosen based on their universal presence in all 30 OTUs in the 10 reference sets as well as in the metagenomes. Maximum-likelihood trees were calculated using IQ-tree with the model set to the General matrix model by Le and Gascuel (LG) following an alignment performed using MAFFT (linsi)^31^

### Comparisons of phylogenetic trees

The pairwise distances between trees were calculated by 4 different tree comparison methods (ALIGN^32^, NODE^33^, RF^24^ and RFK^34^) (described in Kuhner and Yamato^35^) and the Kernel Density Estimate (KDE) of the histogram resulting from the pairwise distances between trees were plotted. Pairs of distributions were compared using a two-sample KS test to test if the two distributions are similar. The p-values for each set of comparisons were corrected for multiple comparisons using the Benjamin Hochberg procedure (Please see Supplementary Tables S3-5 for the full list of p-values for each comparison)

### Neighbour-Net analysis

Alignments of ribosomal proteins for each RefSeq sample and MAG sample were concatenated and used to draw a Neighbor-Net ^27^ using SplitsTree5 using the standard parameters. For the randomization, the 23 ribosomal proteins of the 30 organisms from the archaeal reference sample ARS3001 were scrambled such that each of the 30 organisms now had a random collection of the 23 ribosomal proteins. This results in each of the 30 organisms having a collection of ribosomal proteins from different organisms and therefore different evolutionary histories.

### BMGE based Trimming

In order to check if trimming the alignment to only retain sites with sufficient information Block Mapping and Gathering Entropy (BMGE)^36^ was used with the default settings with the BLOSUM30 substitution matrix. To ensure uniformity, all sequences of a protein (from each of the 10 reference samples and the MAGs) were combined together and aligned again with MAFFT (linsi). This combined alignment was then trimmed with BMGE and then separated into the 11 samples respectively. Trees were then drawn from the trimmed alignments as described before and the trees were compared.

### Incompatibility scores for a set of phylogenetic trees with equal OTUs

For a set of phylogenetic trees *T* we calculated compatibility scores for each tree *t* in the set as follows: Each *n* OTU tree in the set was decomposed into its (*n*-3) splits. A split, *s_1_* from tree *t_1_* is considered compatible with tree *t_2_* if *s_1_* is compatible with all splits present in *t_2_*. The compatibility of a split *s* with the complete set of trees *T* is defined as the fraction of trees in *T* that are compatible with *s*. Finally, the compatibility of a tree *t* with a reference set of trees *T* is defined as the mean compatibility observed among its splits. The differences in the distributions of tree compatibility scores for the two sets of trees was assessed using the two-tailed Kolmogorov–Smirnov test. (please see Supplementary Tables S6-8 which contain the p-values for each comparison).

## Supporting information

Supplementary Infromation

## Acknowledgements

This work was financially supported by the National Natural Science Foundation of China (91851210, 41530105 and 81774152), the European Research Council (ERC 666053), the Shenzhen Key Laboratory of Marine Archaea Geo-Omics, Southern University of Science and Technology, (ZDSYS201802081843490), the Shenzhen Science and Technology Innovation Commission (JCYJ20180305123458107), the VW foundation (93 046), and the Laboratory for Marine Geology, Qingdao National Laboratory for Marine Science and Technology (MGQNLM-TD201810).

---

**Extended Data 1.**
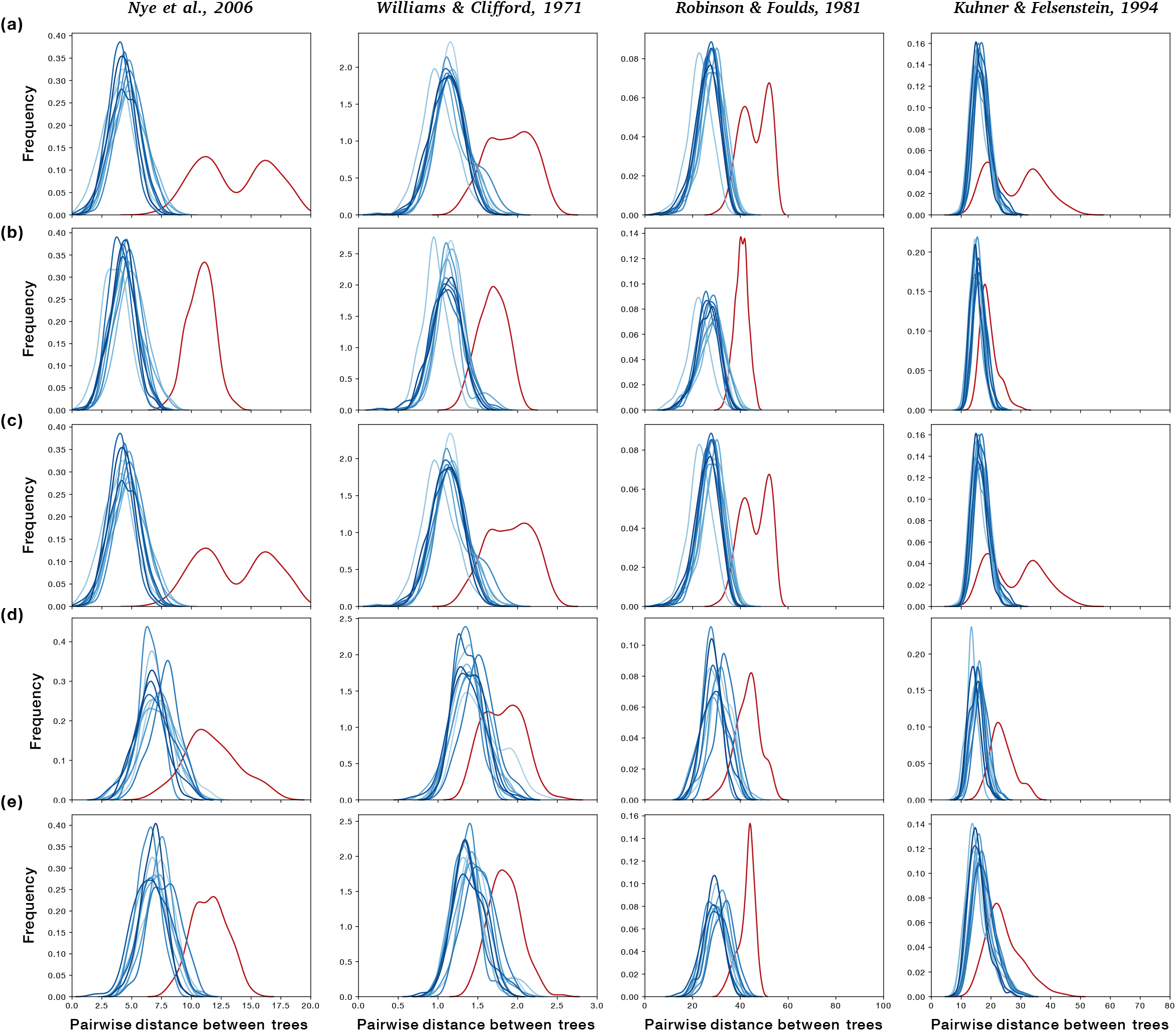
Distribution of pairwise tree-distances between MAGs and RefSeq after site exlusion: In order to test whether the topological inconsistency that we observe for MAG samples in Fig. 2 was due to phylogenetic effects stemming from highly variable sites, we trimmed sites from the alignment using BMGE^36^. In each case, alignments for a matched universal set of proteins from MAGs (shown in red) and 10 random samples of RefSeq (shown in shades of blue) were subjected to site exclusion, tree construction and comparison. Relative to Fig. 2, site-exclusion^36^ does not increase or detract from the topological consistency of trees within a set. **(a)** Trees for 39 universal proteins (site excluded alignments) from 30 Asgard archaeal MAGs are compared with those from 10 samples of 30 archaeal RefSeq genomes. **(b)** Trees for 23 ribosomal proteins (site excluded alignments) from Asgard archaeal MAGs are compared with those from archaeal RefSeq genomes. **(c)** Trees for 16 non-ribosomal proteins (site excluded alignments) from archaeal MAGs are compared with those from archaeal RefSeq genomes. **(d)** Trees for 16 ribosomal proteins (site excluded alignments) from CPR MAGs are compared with those from bacterial RefSeq genomes. **(e)** Trees for 20 ribosomal proteins from bacterial MAGs are compared with those from 10 samples of 30 RefSeq genomes. Note that the comparison of (r-protein or other) tree similarity within and between sets of genomes reported here is distinct from the inspection of individual genomes in which the position of one new set of (r-protein) sequences in a genome would be added to a reference system of closed genomes to be tested for statistical inconsistency in branching behaviour.

**Extended Data 2.**
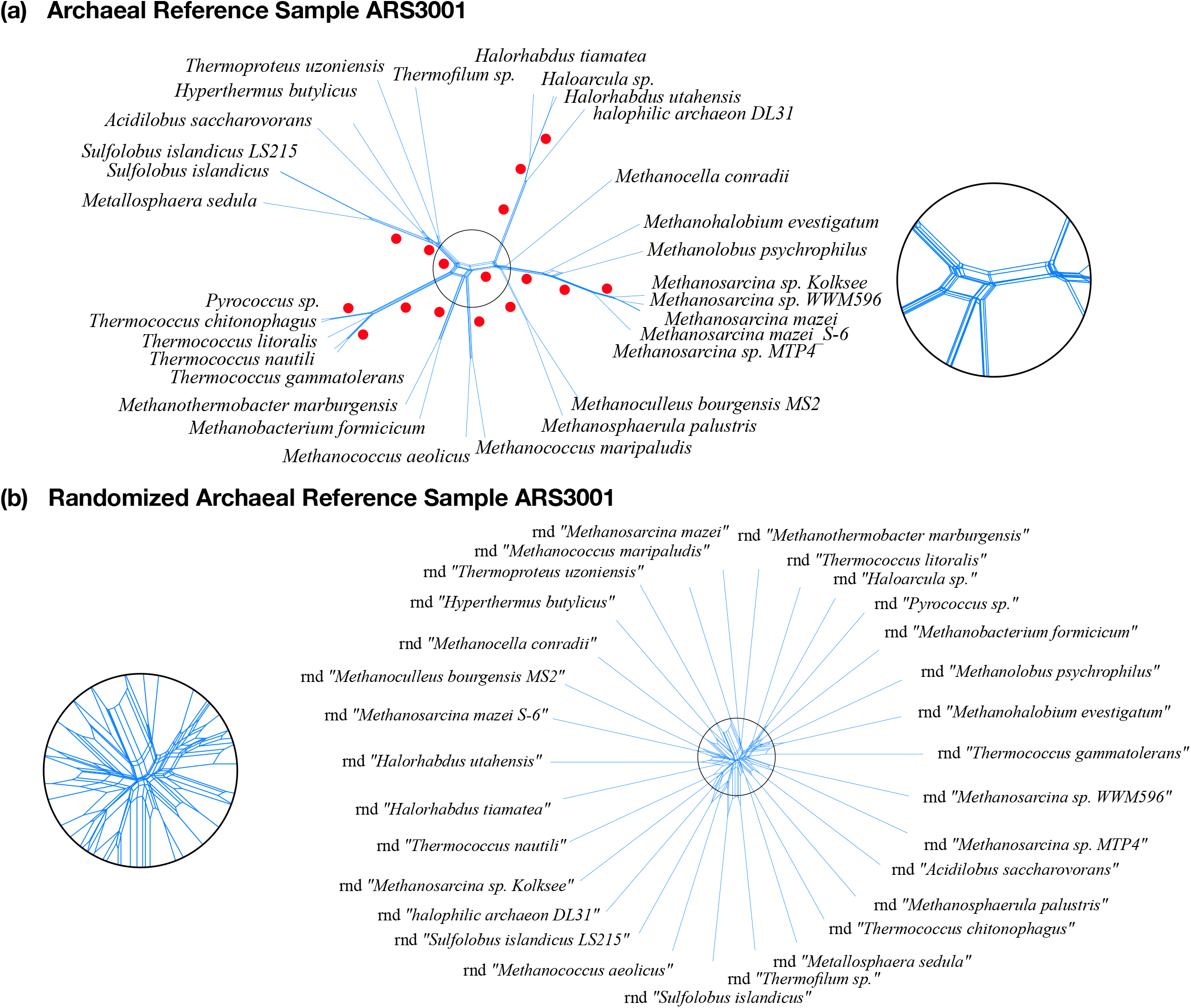
Neighbor-Nets reconstructed from concatenated alignments of 23 ribosomal proteins following randomization. **(a)** The Neighbor-Net of a concatenated alignment of 23 ribosomal proteins in the archaeal reference sample ARS3001 (Supplementary Fig. S1) shows very little conflict throughout, resulting in a tree-like network with 16 well supported splits (indicated with red dots). **(b)** Before generating a concatenated alignment, the 23 r-proteins from the 30 genomes in the RefSeq sample (file number ARS3001) were randomly redistributed. These scrambled genomes, indicated with the prefix rnd, were used to reconstruct a NNet, generated a star-like structure very similar to that of the Asgard archaeal MAG NNet in Fig. 3b.

**Extended Data 3.**
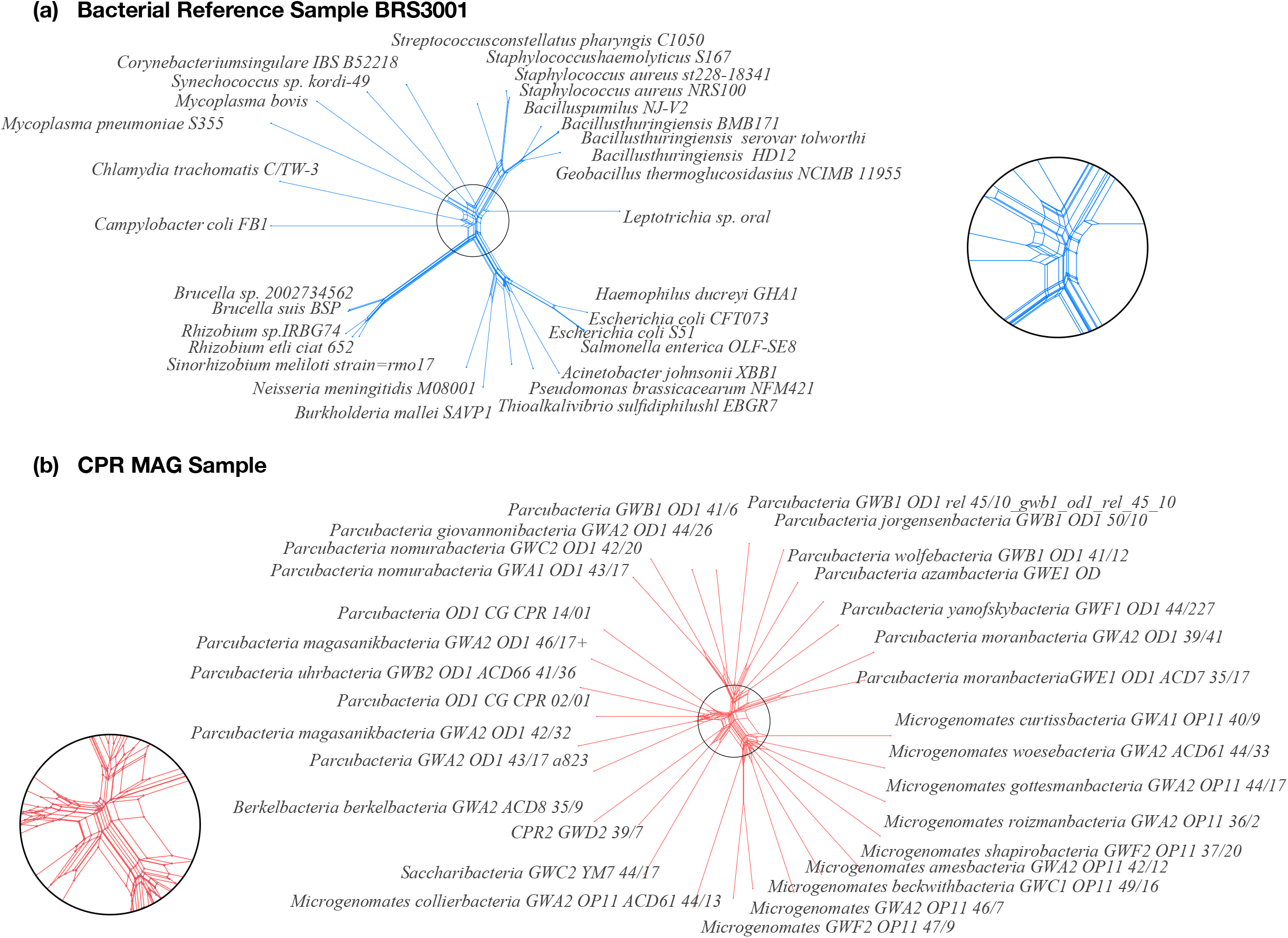
Neighbor-Nets reconstructed from concatenated alignments of 16 ribosomal proteins for bacterial reference samples and CPR MAGs. **(a)** The Neighbor-Net of a concatenated alignment of 16 ribosomal proteins in the bacterial reference sample BRS3001 shows very little conflict throughout, resulting in a tree-like network. **(b)** A Neighbor net drawn from a concatenated alignment of the same 16 ribosomal proteins from CPR MAGs results in a network with a star-like structure. The insets magnify the central area of interest to better highlight the difference between the two networks.

## References

1. Spang, A. et al. Complex archaea that bridge the gap between prokaryotes and eukaryotes. Nature 521, 173 (2015).

2. Zaremba-Niedzwiedzka, K. et al. Asgard archaea illuminate the origin of eukaryotic cellular complexity. Nature 541, 353–358

3. Akıl, C. & Robinson, R. C. Genomes of Asgard archaea encode profilins that regulate actin. Nature 562, 439–443 (2018).

4. Hug, L. A. et al. A new view of the tree of life. Nat Microbiol 1, 16048 (2016).

5. Cunha, V., Gaia, M., Nasir, A. & Forterre, P. Asgard archaea do not close the debate about the universal tree of life topology. PLOS Genet 14, e1007215 (2018).

6. Cunha, V., Gaia, M., Gadelle, D., Nasir, A. & Forterre, P. Lokiarchaea are close relatives of Euryarchaeota, not bridging the gap between prokaryotes and eukaryotes. PLOS Genet 13, e1006810 (2017).

7. Spang, A. et al. Asgard archaea are the closest prokaryotic relatives of eukaryotes. PLOS Genet 14, e1007080 (2018).

8. Penny D., Foulds L.R., Hendy M.D. Testing the theory of evolution by comparing phylogenetic trees constructed from five different protein sequences. Nature 297, 197–200 (1982).

9. Dell’Anno, A., Stefano, B. & Danovaro, R. Quantification, base composition, and fate of extracellular DNA in marine sediments. Limnol Oceanogr 47, 899–905 (2002).

10. Dell’Anno, A. & Danovaro, R. Extracellular DNA plays a key role in deep-sea ecosystem functioning. Science 309, 2179–2179 (2005).

11. Nagler, M., Insam, H., Pietramellara, G. & Ascher-Jenull, J. Extracellular DNA in natural environments: features, relevance and applications. Appl Microbiol Biot 102, 6343–6356 (2018).

12. Torti, A., Lever M. A. & Jørgensen B. B. Origin, dynamics, and implications of extracellular DNA pools in marine sediments. Mar Genomics 24, 185–196 (2015).

13. Orsi, W. D. et al. Climate oscillations reflected within the microbiome of Arabian Sea sediments. Sci Rep 7, 6040 (2017).

14. Vuillemin A. et al. Archaea dominate oxic subseafloor communities over multimillion-year time scales. Sci Adv 5, eaaw4108 (2019).

15. Stephen, J., McCaig, A., Smith, Z., Prosser, J. & Embley, T. M. Molecular diversity of soil and marine 16S rRNA gene sequences related to β-subgroup ammonia-oxidizing bacteria. Appl Environ Microbiol 62, 4147–54 (1996).

16. Martin, W. et al. Gene transfer to the nucleus and the evolution of chloroplasts. Nature 393, 162–165 (1998).

17. Hansmann, S. & Martin, W. Phylogeny of 33 ribosomal and six other proteins encoded in an ancient gene cluster that is conserved across prokaryotic genomes: influence of excluding poorly alignable sites from analysis. Int J Syst Evol Microbiol 50, 1655–1663 (2000).

18. Nesbø, C. L., Boucher, Y. & Doolittle, F. W. Defining the core of nontransferable prokaryotic genes: the euryarchaeal core. J Mol Evol 53, 340–350 (2001).

19. Daubin, V., Gouy, M. & Perrière, G. A Phylogenomic approach to bacterial phylogeny: evidence of a core of genes sharing a common history. Genome Res 12, 1080–1090 (2002).

20. Matte-Tailliez, O., Brochier, C., Forterre, P. & Philippe, H. Archaeal phylogeny based on ribosomal proteins. Mol Biol Evol 19, 631–639 (2002).

21. Brown, J. R., Douady, C. J., Italia, M. J., Marshall, W. E. & Stanhope, M. J. Universal trees based on large combined protein sequence data sets. Nature Genet 28, 281–285 (2001).

22. Shen, X.-X., Hittinger, C. & Rokas, A. Contentious relationships in phylogenomic studies can be driven by a handful of genes. Nature Ecol Evol 1, s41559-017–0126 (2017).

23. Steel, M., Huson, D. & Lockhart, P. J. Invariable sites models and their use in phylogeny reconstruction. Systematic Biol 49, 225–232 (2000).

24. Robinson, D. F. & Foulds, L. R. Comparison of phylogenetic trees. Mathematical Biosciences 53, 131–147 (1981).

25. Talavera, G. & Castresana, J. Improvement of phylogenies after removing divergent and ambiguously aligned blocks from protein sequence alignments. Systematic Biol 56, 564–577 (2007).

26. Fan, L. et al. Mitochondria branch within Alphaproteobacteria. Biorxiv 715870 (2019). doi:10.1101/715870

27. Bryant, D. & Moulton, V. Neighbor-Net: An agglomerative method for the construction of phylogenetic networks. Mol Biol Evol 21, 255–265 (2004).

28. Nelson-Sathi, S. et al. Acquisition of 1,000 eubacterial genes physiologically transformed a methanogen at the origin of Haloarchaea. Proc Natl Acad Sci USA 109, 20537–20542 (2012).

29. Imachi, H. et al. Isolation of an archaeon at the prokaryote-eukaryote interface. Biorxiv 726976 (2019). doi:10.1101/726976

30. Martin, W. F., Tielens, A. G. M., Mentel, M., Garg, S. G. & Gould, S. B. The physiology of phagocytosis in the context of mitochondrial origin. Microbiol Mol Biol R 81, e00008–17 (2017).

## Methods References

31. Katoh, K. & Standley, D. M. MAFFT Multiple sequence alignment software version 7: Improvements in performance and usability. Mol Biol Evol 30, 772–780 (2013).

32. Nye, T., Liò, P. & Gilks, W. R. A novel algorithm and web-based tool for comparing two alternative phylogenetic trees. Bioinformatics 22, 117–119 (2006).

33. Williams, W. & Clifford, H. On the comparison of two classifications of the same set of elements. Taxon 20, 519–522 (1971).

34. Kuhner, M. & Felsenstein, J. A simulation comparison of phylogeny algorithms under equal and unequal evolutionary rates. Mol Biol Evol 11, 459–68 (1994).

35. Kuhner, M. K. & Yamato, J. Practical performance of tree comparison metrics. Syst Biol 64, 205–214 (2015).

36. Criscuolo, A. & Gribaldo, S. BMGE (Block mapping and gathering with entropy): a new software for selection of phylogenetic informative regions from multiple sequence alignments. BMC Evol Biol 10, 210 (2010).

