## Supplementary Infromation for "Anomalous phylogenetic behavior of ribosomal proteins in metagenome assembled genomes"

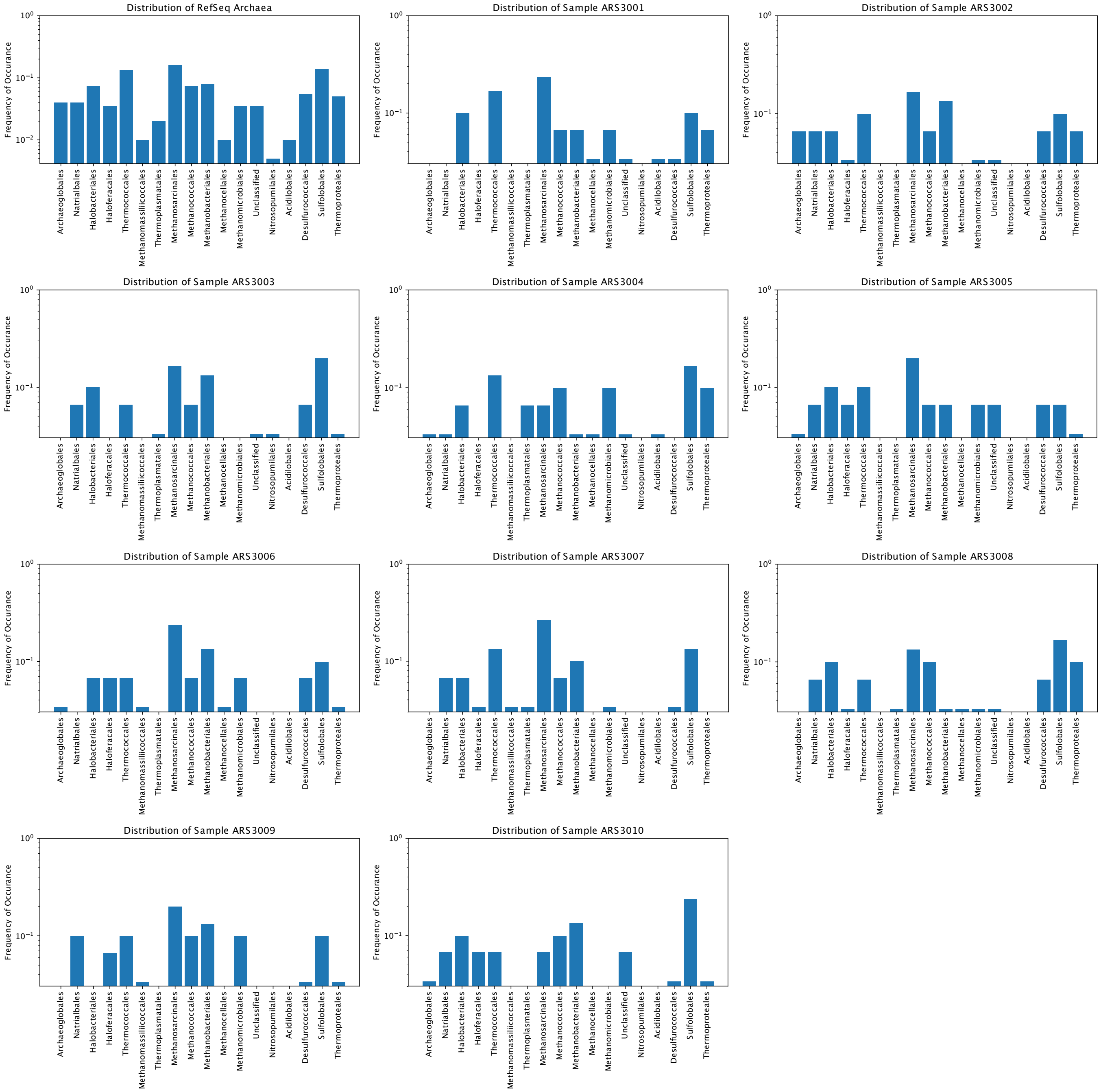


**Supplementary Fig. S1 | Distribution of archaeal genomes across taxonomic groups.** Each sample used in the study is represented by a panel, which indicates the taxonomic distribution of the organisms within that sample from across 212 completely sequenced archaeal RefSeq genomes. For each sample we chose 30 genomes such that they approximate the diversity across all 212 genomes.

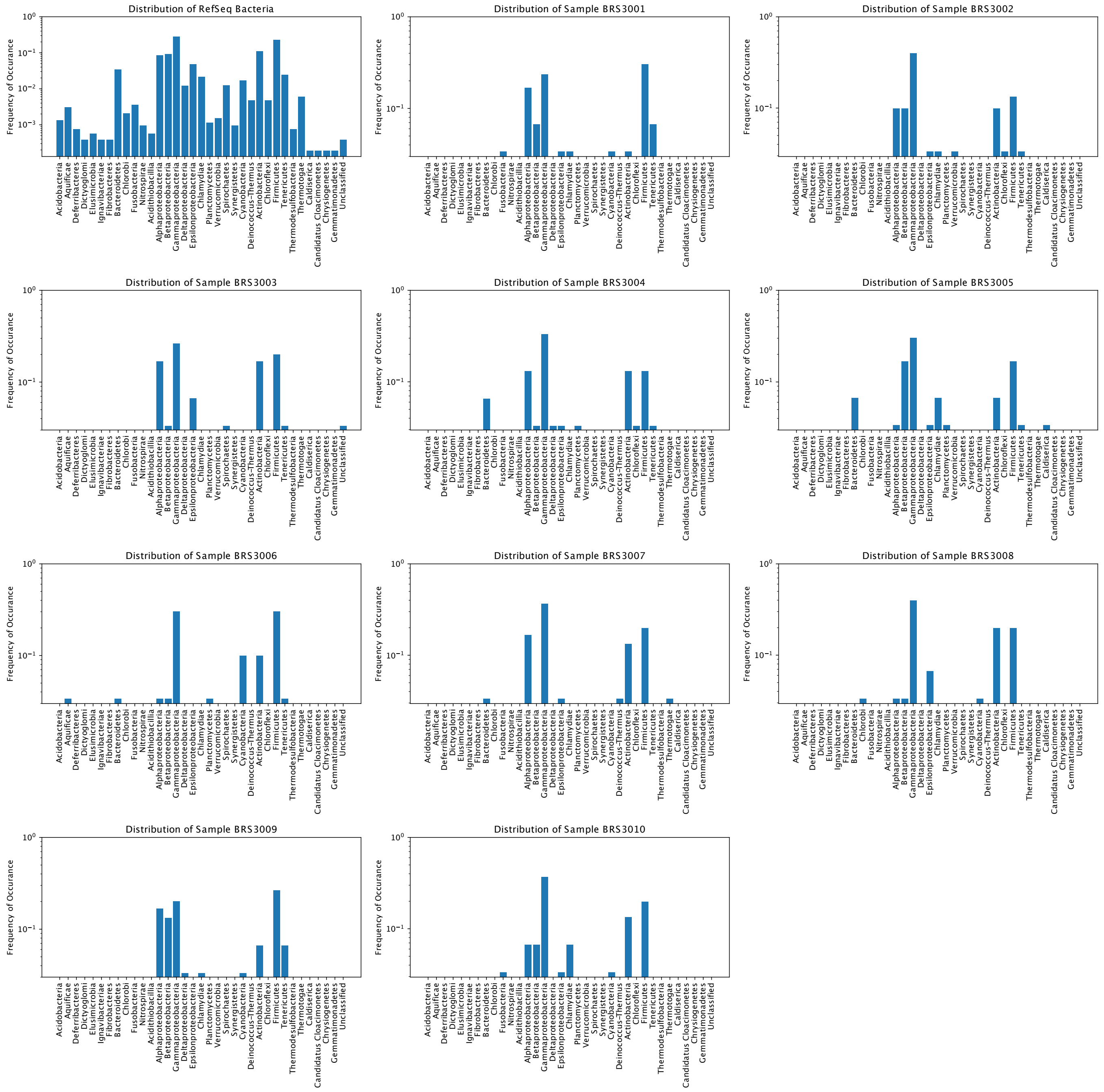


**Supplementary Fig. S2 | Distribution of bacterial genomes across taxonomic groups.** Each sample used in the study is represented by a panel which indicates the taxonomic distribution of the organisms within that sample from across 5443 completely sequenced bacterial RefSeq genomes. For each sample we chose 30 genomes such that they approximate the diversity and the representation across all 5443 genomes.

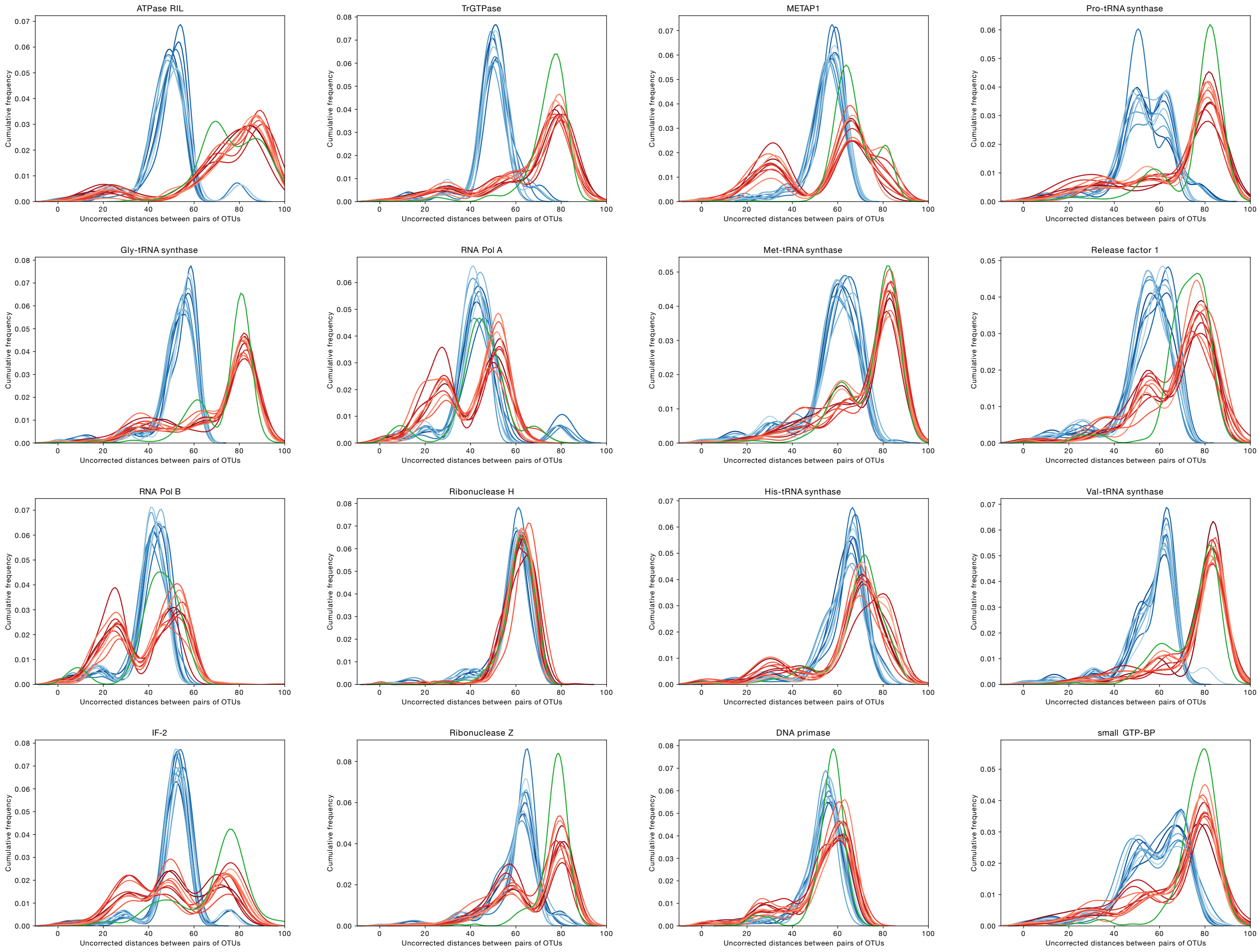


**Supplementary Fig. S3 | Uncorrected p-distances for 16 archaeal non-ribosomal proteins:** To examine the possibility that topological differences in trees could stem from the phylogenetic depth (sequence divergence) within the individual protein trees, we plotted uncorrected p-distances for each of the 16 universal non-ribosomal proteins from 10 archaeal RefSeq samples (blue), 10 non-asgard archaeal MAGs (red) and one asgard archaeal MAG sample (green), samples consisting of 30 organisms each.

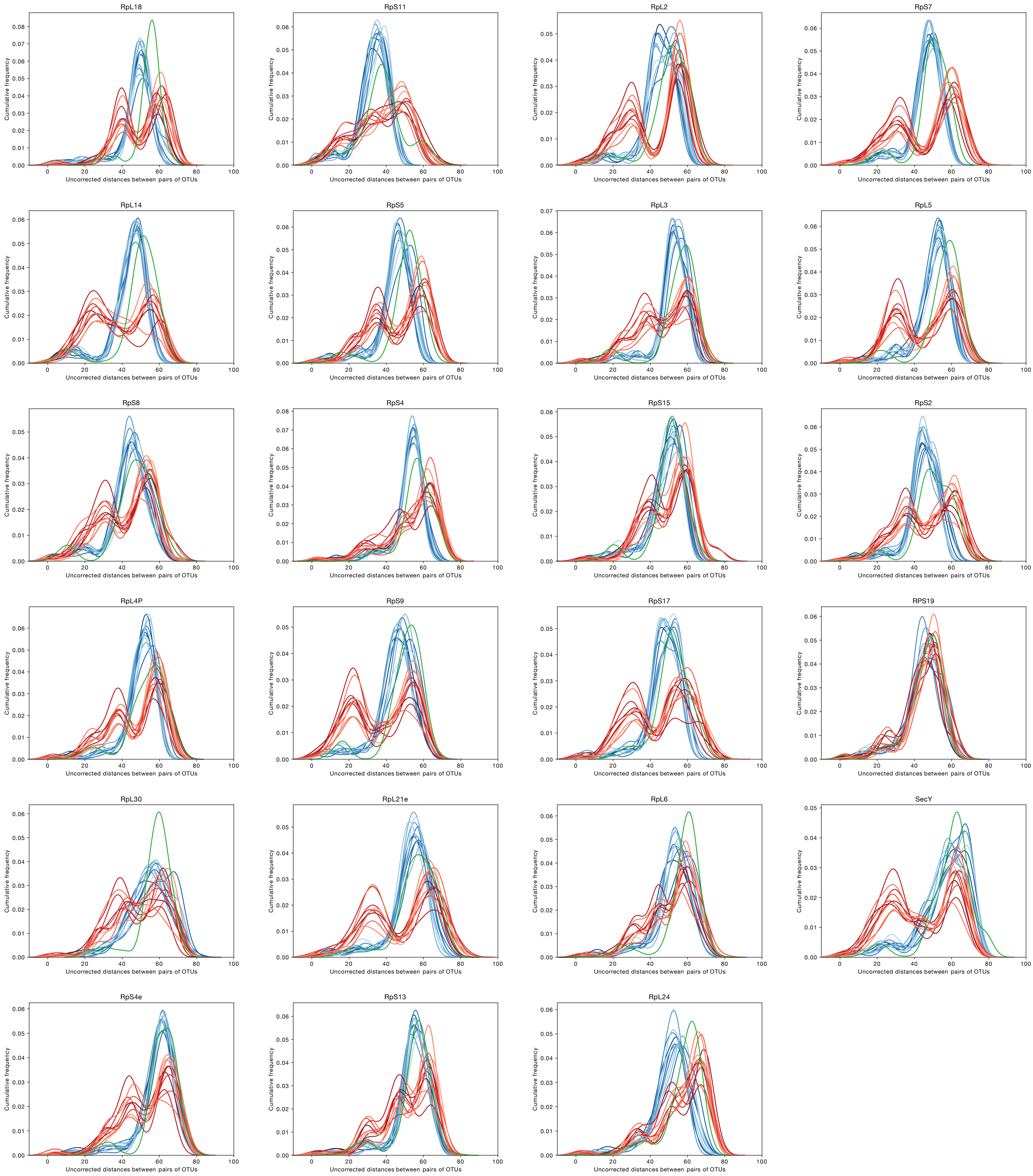


**Supplementary Fig. S4 | Uncorrected p-distances for 23 archaeal ribosomal proteins:** Same as in Supplemental Fig. S3, but for 23 ribosomal proteins from 10 archaeal RefSeq samples (blue), 10 non-asgard archaeal MAGs (red) and one asgard archaeal MAG sample (green), samples consisting of 30 organisms each. Note that p-distances for non-ribosomal proteins (Supplemental Fig. S3) of MAGs are often shifted towards higher divergence, which is a property of the MAGs themselves, while p-distances between 23 r-proteins are more uniform for the same samples, though often bimodal for non-asgard MAGs, but not bimodal for asgard MAGs. Note that the distribution pf p-distances for r-ptoeins from asgard MAGs closely follows the distribution for archaeal orgDNA.

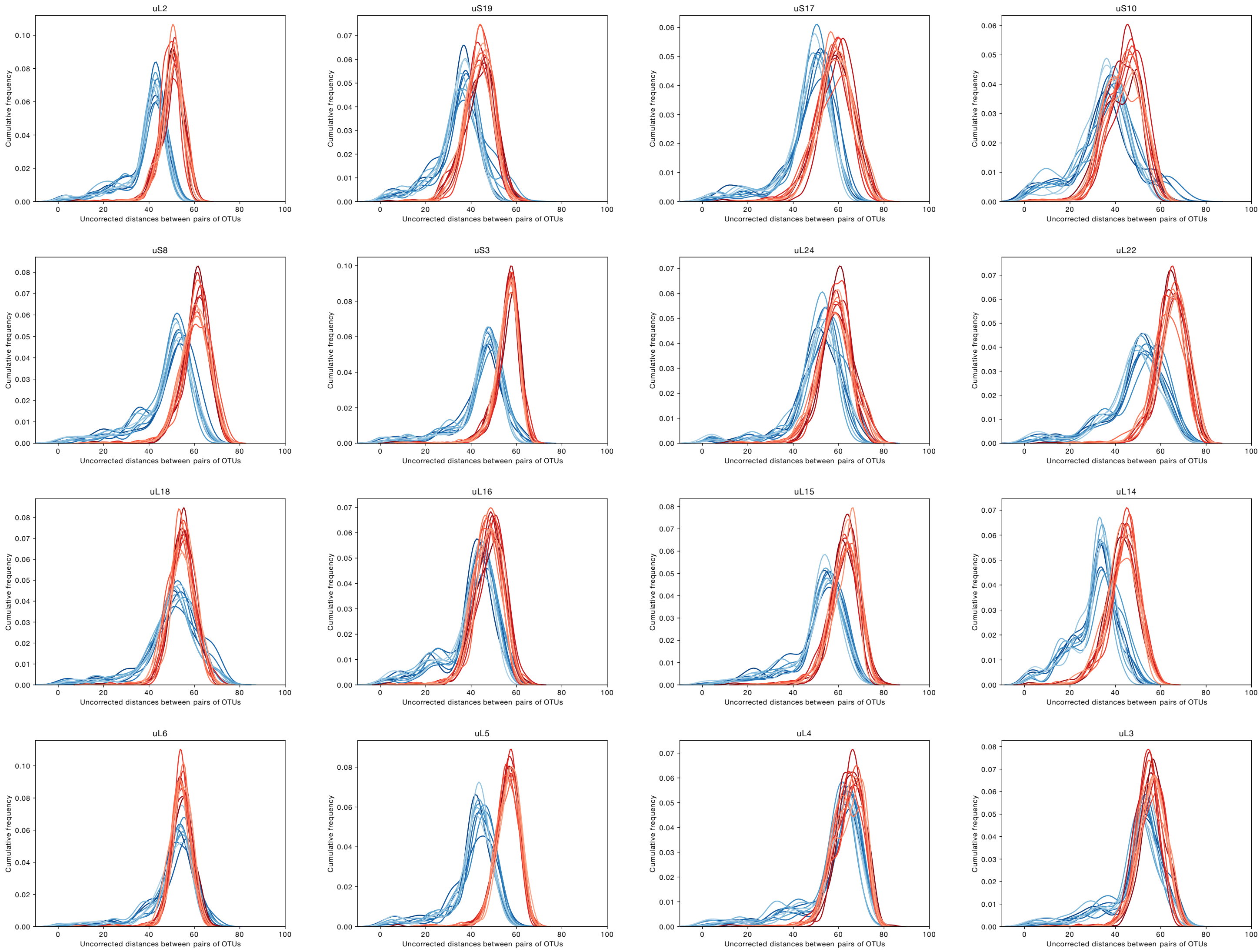


**Supplementary Fig. S5 | Uncorrected p-distances for 16 ribosomal proteins from bacterial RefSeq genomes and CPR MAGs:** The uncorrected p-distances for each of the 16 universal ribosomal proteins from 10 bacterial RefSeq samples (blue) and 11 CPR MAG samples (red) each consisting of 30 organisms each are plotted.

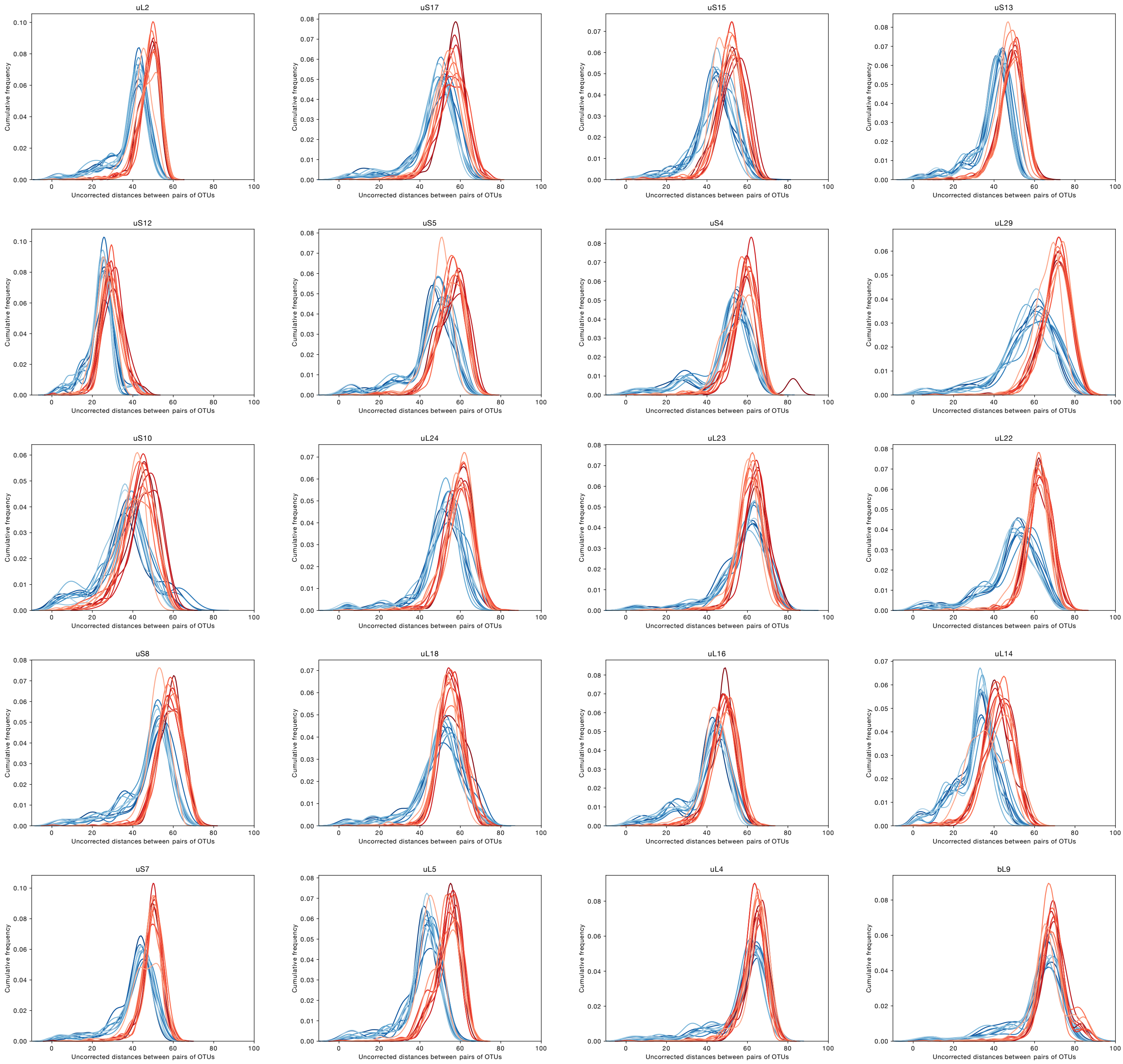


**Supplementary Fig. S6 | Uncorrected p-distances for 20 ribosomal proteins from bacteria:** The uncorrected p-distances for each of the 20 universal ribosomal proteins from 10 bacterial RefSeq samples (blue) and 10 bacterial non-CPR MAG samples (red) each consisting of 30 organisms each are plotted.

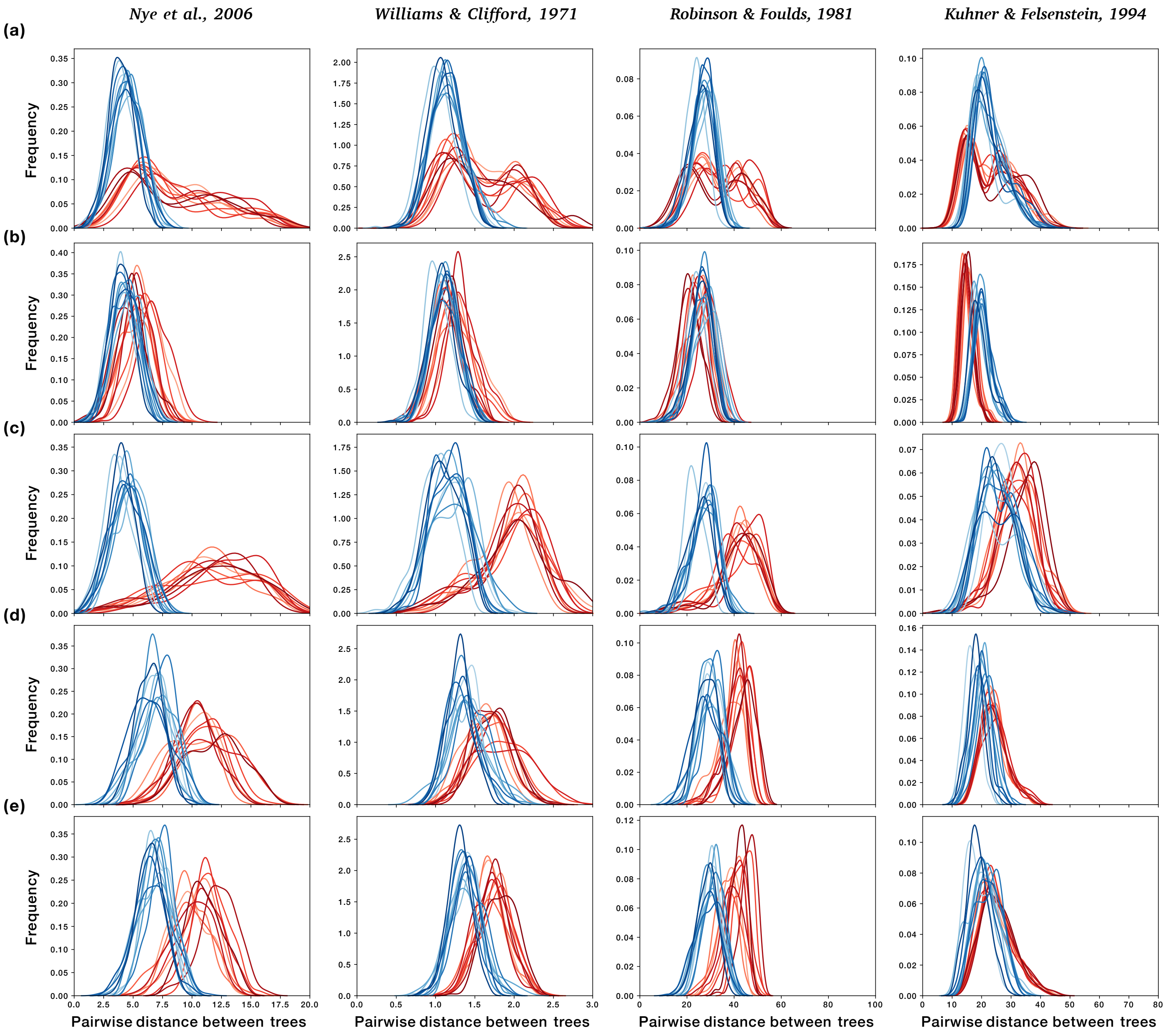


**Supplementary Fig. S7 | Distribution of pairwise tree-distances between RefSeq and metagenomes:** The pairwise comparisons of tree distances computed using four different metrics (see Methods) are shown. In each case, a matched set of proteins present in 10 MAG samples and 10 samples from RefSeq are taken to plot comparable distributions. The MAG sample is always shown in red. **(a)** Trees for 39 universal proteins from 10 samples of 30 non-asgard archaeal MAGs each are compared with trees for the 39 homologues from 10 samples of 30 archaeal RefSeq genomes **(b)** Trees for 23 ribosomal proteins from 10 samples of 30 non-asgard archaeal MAGs each are compared with those from 10 samples of 30 archaeal RefSeq genomes. **(c)** Trees for 16 non-ribosomal proteins from 30 archaeal MAGs are compared with those from 10 samples of 30 archaeal RefSeq genomes. **(d)** Trees for 16 ribosomal proteins from 30 candidate phyla radiation CPR MAGs from ref. 4 are compared with those from 10 samples of 30 bacterial RefSeq genomes. **(e)** Trees for 20 ribosomal proteins from non-CPR 30 bacterial MAGs are compared with those from 10 samples of 30 RefSeq genomes. In all panels, blue curves represent the 10 independent reference samples while the red curve represents the MAGs. Individual p-values for each comparison are given in Table S4.

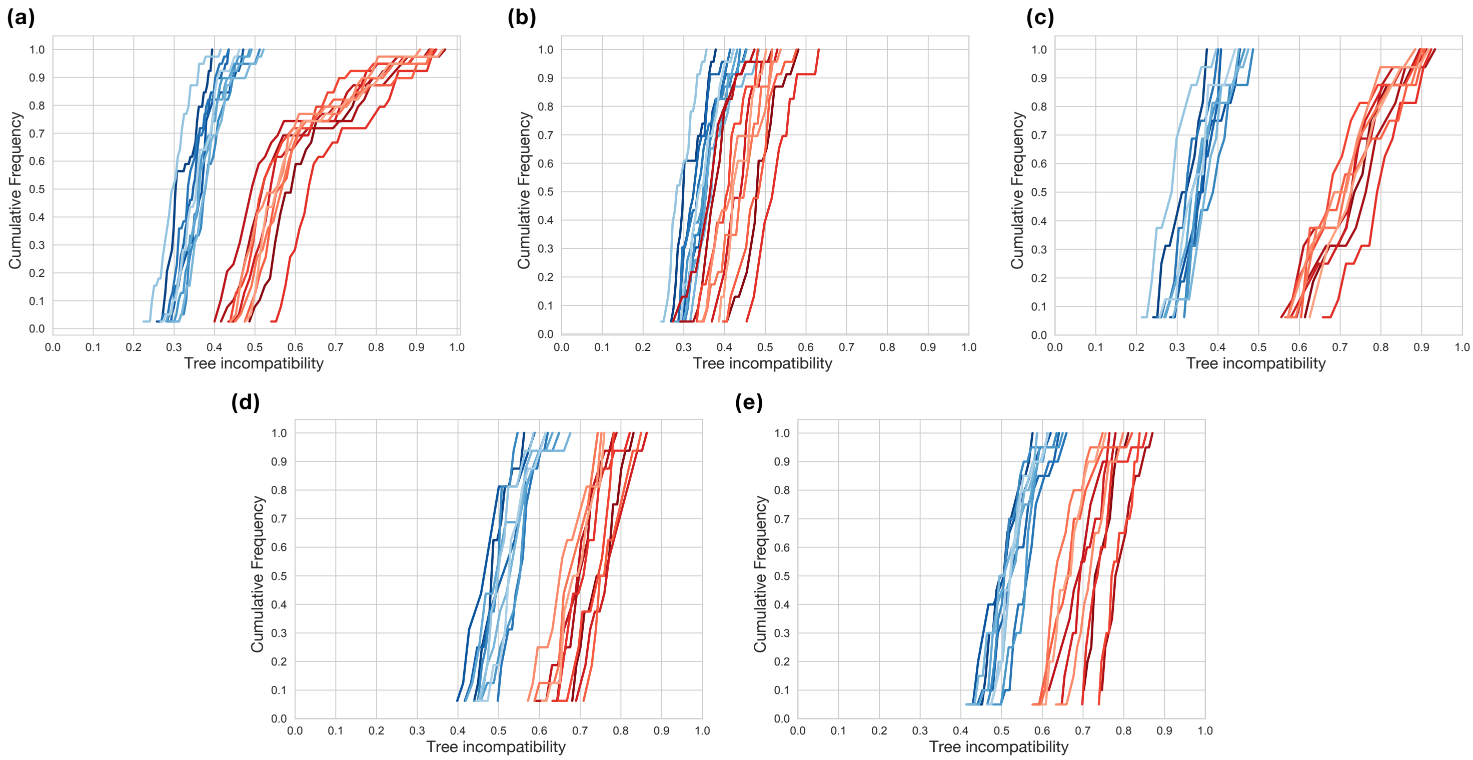


**Supplementary Fig. S8 | Tree compatibility scores for samples of tree reconstructed from orgDNA and metDNA.** Cumulative distribution of tree incompatibility scores within sets of gene trees. In each case every curve represents a set of 30 organisms where the RefSeq samples are shown in shades of blue, the MAG samples are shown in shades of red **(a)** Trees for 39 universal proteins sampled from 10 archaeal RefSeq genomes versus 10 non-asgard archaeal MAGs. **(b)** Trees for a subset of 23 ribosomal proteins sampled from 10 archaeal RefSeq genomes versus 10 non-asgard archaeal MAGs. Note that trees for ribosomal proteins from non-asgard archaeal MAGs (here) are more topologically similar to each other than trees for ribosomal proteins from asgard MAGs (main text, Fig. 4b). **(c)** Trees for the set of 16 non-ribosomal proteins sampled from 10 archaeal RefSeq genomes versus 10 non-asgard archaeal MAGs. **(d)** Trees for 16 ribosomal proteins sampled from 10 bacterial RefSeq genomes versus 10 CPR MAGs. **(e)** Trees for 20 ribosomal proteins sampled from 10 bacterial RefSeq genomes versus 10 non-CPR bacterial MAGs.

**Supplementary Table S3A:** Pairwise FDR corrected p-values of the Kolmogorow-Smirnow test on the distances between phylogenetic trees for 39 universal proteins in 10 samples of archaeal RefSeq genomes vs one sample containing archaeal MAGs (associated with main Fig. 2a)

Comparison of trees using the Nye et al., 2006 method

| | ARS3001 | ARS3002 | ARS3003 | ARS3004 | ARS3005 | ARS3006 | ARS3007 | ARS3008 | ARS3009 | ARS3010 | Metagenomes |
|---|---|---|---|---|---|---|---|---|---|---|---|
| ARS3001 | 1.00E+00 | 1.91E-14 | 3.26E-03 | 8.12E-06 | 1.76E-24 | 1.18E-08 | 6.93E-20 | 1.37E-22 | 2.91E-01 | 1.15E-04 | 0.00E+00 |
| ARS3002 | 1.91E-14 | 1.00E+00 | 9.73E-07 | 3.42E-04 | 1.57E-02 | 4.65E-03 | 1.67E-01 | 9.21E-02 | 1.30E-18 | 7.31E-05 | 0.00E+00 |
| ARS3003 | 3.26E-03 | 9.73E-07 | 1.00E+00 | 4.11E-01 | 5.36E-12 | 2.13E-02 | 1.18E-08 | 1.06E-10 | 1.68E-05 | 7.46E-01 | 0.00E+00 |
| ARS3004 | 8.12E-06 | 3.42E-04 | 4.11E-01 | 1.00E+00 | 7.49E-08 | 3.44E-01 | 2.92E-06 | 7.53E-07 | 1.18E-08 | 9.15E-01 | 0.00E+00 |
| ARS3005 | 1.76E-24 | 1.57E-02 | 5.36E-12 | 7.49E-08 | 1.00E+00 | 6.41E-06 | 7.10E-01 | 3.17E-01 | 9.58E-32 | 4.33E-10 | 0.00E+00 |
| ARS3006 | 1.18E-08 | 4.65E-03 | 2.13E-02 | 3.44E-01 | 6.41E-06 | 1.00E+00 | 2.24E-04 | 1.68E-05 | 5.15E-13 | 1.51E-01 | 0.00E+00 |
| ARS3007 | 6.93E-20 | 1.67E-01 | 1.18E-08 | 2.92E-06 | 7.10E-01 | 2.24E-04 | 1.00E+00 | 4.85E-01 | 6.57E-26 | 9.85E-08 | 0.00E+00 |
| ARS3008 | 1.37E-22 | 9.21E-02 | 1.06E-10 | 7.53E-07 | 3.17E-01 | 1.68E-05 | 4.85E-01 | 1.00E+00 | 6.08E-25 | 2.18E-08 | 0.00E+00 |
| ARS3009 | 2.91E-01 | 1.30E-18 | 1.68E-05 | 1.18E-08 | 9.58E-32 | 5.15E-13 | 6.57E-26 | 6.08E-25 | 1.00E+00 | 4.33E-07 | 0.00E+00 |
| ARS3010 | 1.15E-04 | 7.31E-05 | 7.46E-01 | 9.15E-01 | 4.33E-10 | 1.51E-01 | 9.85E-08 | 2.18E-08 | 4.33E-07 | 1.00E+00 | 0.00E+00 |
| Metagenomes | 0.00E+00 | 0.00E+00 | 0.00E+00 | 0.00E+00 | 0.00E+00 | 0.00E+00 | 0.00E+00 | 0.00E+00 | 0.00E+00 | 0.00E+00 | 1.00E+00 |

Comparison of trees using the Williams & Clifford, 1971 method

| | ARS3001 | ARS3002 | ARS3003 | ARS3004 | ARS3005 | ARS3006 | ARS3007 | ARS3008 | ARS3009 | ARS3010 | Metagenomes |
|---|---|---|---|---|---|---|---|---|---|---|---|
| ARS3001 | 1.00E+00 | 1.50E-07 | 1.37E-09 | 3.88E-05 | 3.40E-08 | 2.57E-08 | 7.86E-04 | 2.61E-09 | 1.15E-07 | 3.50E-10 | 1.73E-283 |
| ARS3002 | 1.50E-07 | 1.00E+00 | 9.90E-01 | 1.67E-01 | 1.26E-02 | 1.08E-01 | 9.87E-02 | 2.16E-03 | 4.61E-23 | 5.65E-01 | 2.68E-280 |
| ARS3003 | 1.37E-09 | 9.90E-01 | 1.00E+00 | 3.22E-02 | 2.47E-02 | 1.67E-01 | 1.87E-02 | 9.09E-03 | 2.04E-26 | 5.65E-01 | 6.66E-274 |
| ARS3004 | 3.88E-05 | 1.67E-01 | 3.22E-02 | 1.00E+00 | 3.67E-02 | 2.82E-02 | 5.39E-01 | 1.63E-02 | 1.50E-16 | 1.63E-02 | 9.89E-271 |
| ARS3005 | 3.40E-08 | 1.26E-02 | 2.47E-02 | 3.67E-02 | 1.00E+00 | 2.04E-01 | 1.45E-02 | 9.90E-01 | 6.44E-22 | 1.85E-01 | 3.01E-225 |
| ARS3006 | 2.57E-08 | 1.08E-01 | 1.67E-01 | 2.82E-02 | 2.04E-01 | 1.00E+00 | 2.47E-02 | 1.36E-01 | 6.06E-26 | 2.24E-01 | 1.65E-234 |
| ARS3007 | 7.86E-04 | 9.87E-02 | 1.87E-02 | 5.39E-01 | 1.45E-02 | 2.47E-02 | 1.00E+00 | 3.12E-03 | 5.59E-16 | 6.50E-03 | 4.28E-277 |
| ARS3008 | 2.61E-09 | 2.16E-03 | 9.09E-03 | 1.63E-02 | 9.90E-01 | 1.36E-01 | 3.12E-03 | 1.00E+00 | 9.45E-24 | 1.08E-01 | 3.59E-217 |
| ARS3009 | 1.15E-07 | 4.61E-23 | 2.04E-26 | 1.50E-16 | 6.44E-22 | 6.06E-26 | 5.59E-16 | 9.45E-24 | 1.00E+00 | 2.06E-27 | 0.00E+00 |
| ARS3010 | 3.50E-10 | 5.65E-01 | 5.65E-01 | 1.63E-02 | 1.85E-01 | 2.24E-01 | 6.50E-03 | 1.08E-01 | 2.06E-27 | 1.00E+00 | 2.90E-261 |
| Metagenomes | 1.73E-283 | 2.68E-280 | 6.66E-274 | 9.89E-271 | 3.01E-225 | 1.65E-234 | 4.28E-277 | 3.59E-217 | 0.00E+00 | 2.90E-261 | 1.00E+00 |

Comparison of trees using the Robinson & Foulds, 1981 method

| | ARS3001 | ARS3002 | ARS3003 | ARS3004 | ARS3005 | ARS3006 | ARS3007 | ARS3008 | ARS3009 | ARS3010 | Metagenomes |
|---|---|---|---|---|---|---|---|---|---|---|---|
| ARS3001 | 1.00E+00 | 2.45E-05 | 4.10E-06 | 5.14E-02 | 2.47E-07 | 3.98E-08 | 6.57E-15 | 1.37E-24 | 3.04E-11 | 5.59E-04 | 0.00E+00 |
| ARS3002 | 2.45E-05 | 1.00E+00 | 8.89E-01 | 3.99E-02 | 1.85E-03 | 2.50E-04 | 3.22E-07 | 3.20E-16 | 1.06E-29 | 8.82E-02 | 0.00E+00 |
| ARS3003 | 4.10E-06 | 8.89E-01 | 1.00E+00 | 3.94E-03 | 9.87E-02 | 2.62E-02 | 1.28E-04 | 1.01E-11 | 1.37E-33 | 8.10E-01 | 0.00E+00 |
| ARS3004 | 5.14E-02 | 3.99E-02 | 3.94E-03 | 1.00E+00 | 4.61E-04 | 1.55E-03 | 3.98E-08 | 1.84E-15 | 3.51E-20 | 1.61E-01 | 0.00E+00 |
| ARS3005 | 2.47E-07 | 1.85E-03 | 9.87E-02 | 4.61E-04 | 1.00E+00 | 1.00E+00 | 4.53E-02 | 5.21E-06 | 8.08E-37 | 4.03E-01 | 0.00E+00 |
| ARS3006 | 3.98E-08 | 2.50E-04 | 2.62E-02 | 1.55E-03 | 1.00E+00 | 1.00E+00 | 2.00E-01 | 1.28E-04 | 1.06E-37 | 2.74E-01 | 0.00E+00 |
| ARS3007 | 6.57E-15 | 3.22E-07 | 1.28E-04 | 3.98E-08 | 4.53E-02 | 2.00E-01 | 1.00E+00 | 3.99E-02 | 3.94E-52 | 3.04E-04 | 3.51E-315 |
| ARS3008 | 1.37E-24 | 3.20E-16 | 1.01E-11 | 1.84E-15 | 5.21E-06 | 1.28E-04 | 3.99E-02 | 1.00E+00 | 2.82E-61 | 4.33E-09 | 9.64E-289 |
| ARS3009 | 3.04E-11 | 1.06E-29 | 1.37E-33 | 3.51E-20 | 8.08E-37 | 1.06E-37 | 3.94E-52 | 2.82E-61 | 1.00E+00 | 7.98E-27 | 0.00E+00 |
| ARS3010 | 5.59E-04 | 8.82E-02 | 8.10E-01 | 1.61E-01 | 4.03E-01 | 2.74E-01 | 3.04E-04 | 4.33E-09 | 7.98E-27 | 1.00E+00 | 0.00E+00 |
| Metagenomes | 0.00E+00 | 0.00E+00 | 0.00E+00 | 0.00E+00 | 0.00E+00 | 0.00E+00 | 3.51E-315 | 9.64E-289 | 0.00E+00 | 0.00E+00 | 1.00E+00 |

Comparison of trees using the Kuhner & Felsenstein, 1994 method

| | ARS3001 | ARS3002 | ARS3003 | ARS3004 | ARS3005 | ARS3006 | ARS3007 | ARS3008 | ARS3009 | ARS3010 | Metagenomes |
|---|---|---|---|---|---|---|---|---|---|---|---|
| ARS3001 | 1.00E+00 | 2.75E-03 | 4.71E-09 | 4.71E-09 | 4.75E-05 | 8.13E-02 | 9.11E-02 | 2.63E-01 | 3.04E-04 | 1.85E-09 | 2.23E-136 |
| ARS3002 | 2.75E-03 | 1.00E+00 | 3.76E-03 | 3.76E-03 | 5.59E-04 | 3.76E-03 | 3.22E-06 | 3.21E-02 | 8.12E-11 | 6.22E-06 | 8.63E-109 |
| ARS3003 | 4.71E-09 | 3.76E-03 | 1.00E+00 | 6.97E-01 | 3.21E-02 | 3.59E-07 | 2.58E-15 | 3.96E-06 | 3.87E-21 | 8.13E-02 | 1.69E-128 |
| ARS3004 | 4.71E-09 | 3.76E-03 | 6.97E-01 | 1.00E+00 | 4.92E-02 | 4.65E-07 | 2.58E-15 | 5.04E-06 | 8.58E-22 | 7.38E-02 | 8.84E-127 |
| ARS3005 | 4.75E-05 | 5.59E-04 | 3.21E-02 | 4.92E-02 | 1.00E+00 | 3.26E-08 | 5.91E-08 | 8.06E-07 | 3.10E-14 | 2.45E-02 | 8.84E-127 |
| ARS3006 | 8.13E-02 | 3.76E-03 | 3.59E-07 | 4.65E-07 | 3.26E-08 | 1.00E+00 | 3.70E-04 | 7.32E-01 | 6.22E-06 | 3.61E-13 | 3.18E-135 |
| ARS3007 | 9.11E-02 | 3.22E-06 | 2.58E-15 | 2.58E-15 | 5.91E-08 | 3.70E-04 | 1.00E+00 | 1.27E-03 | 9.15E-03 | 1.40E-14 | 1.52E-136 |
| ARS3008 | 2.63E-01 | 3.21E-02 | 3.96E-06 | 5.04E-06 | 8.06E-07 | 7.32E-01 | 1.27E-03 | 1.00E+00 | 3.96E-06 | 6.79E-10 | 6.68E-131 |
| ARS3009 | 3.04E-04 | 8.12E-11 | 3.87E-21 | 8.58E-22 | 3.10E-14 | 6.22E-06 | 9.15E-03 | 3.96E-06 | 1.00E+00 | 5.49E-22 | 8.28E-121 |
| ARS3010 | 1.85E-09 | 6.22E-06 | 8.13E-02 | 7.38E-02 | 2.45E-02 | 3.61E-13 | 1.40E-14 | 6.79E-10 | 5.49E-22 | 1.00E+00 | 2.23E-136 |
| Metagenomes | 2.23E-136 | 8.63E-109 | 1.69E-128 | 8.84E-127 | 8.84E-127 | 3.18E-135 | 1.52E-136 | 6.68E-131 | 8.28E-121 | 2.23E-136 | 1.00E+00 |

**Supplementary Table S3B:** Pairwise FDR corrected p-values of the Kolmogorow-Smirnow test on the distances between phylogenetic trees for 23 ribosomal proteins (including SecY) in 10 samples of archaeal RefSeq genomes vs one sample containing archaeal MAGs. (associated with main Fig. 2b)

Comparison of trees using the Nye et al., 2006 method

| | ARS3001 | ARS3002 | ARS3003 | ARS3004 | ARS3005 | ARS3006 | ARS3007 | ARS3008 | ARS3009 | ARS3010 | Metagenomes |
|---|---|---|---|---|---|---|---|---|---|---|---|
| ARS3001 | 1.00E+00 | 4.01E-05 | 7.92E-01 | 1.33E-02 | 1.24E-04 | 6.60E-02 | 5.54E-09 | 5.89E-05 | 3.71E-01 | 2.11E-02 | 5.20E-145 |
| ARS3002 | 4.01E-05 | 1.00E+00 | 3.51E-04 | 1.01E-02 | 7.20E-01 | 2.11E-02 | 9.89E-02 | 9.82E-01 | 9.21E-09 | 9.89E-02 | 4.89E-141 |
| ARS3003 | 7.92E-01 | 3.51E-04 | 1.00E+00 | 2.71E-02 | 1.01E-03 | 2.70E-01 | 4.75E-08 | 4.97E-04 | 1.86E-01 | 6.60E-02 | 4.12E-147 |
| ARS3004 | 1.33E-02 | 1.01E-02 | 2.71E-02 | 1.00E+00 | 2.11E-02 | 7.20E-01 | 2.46E-04 | 5.55E-02 | 1.79E-04 | 7.20E-01 | 4.12E-147 |
| ARS3005 | 1.24E-04 | 7.20E-01 | 1.01E-03 | 2.11E-02 | 1.00E+00 | 5.55E-02 | 2.11E-02 | 5.04E-01 | 1.33E-07 | 3.71E-01 | 5.83E-143 |
| ARS3006 | 6.60E-02 | 2.11E-02 | 2.70E-01 | 7.20E-01 | 5.55E-02 | 1.00E+00 | 1.24E-04 | 6.60E-02 | 5.53E-03 | 8.59E-01 | 5.83E-143 |
| ARS3007 | 5.54E-09 | 9.89E-02 | 4.75E-08 | 2.46E-04 | 2.11E-02 | 1.24E-04 | 1.00E+00 | 9.89E-02 | 3.12E-13 | 2.46E-04 | 4.89E-141 |
| ARS3008 | 5.89E-05 | 9.82E-01 | 4.97E-04 | 5.55E-02 | 5.04E-01 | 6.60E-02 | 9.89E-02 | 1.00E+00 | 3.62E-07 | 4.52E-02 | 4.41E-139 |
| ARS3009 | 3.71E-01 | 9.21E-09 | 1.86E-01 | 1.79E-04 | 1.33E-07 | 5.53E-03 | 3.12E-13 | 3.62E-07 | 1.00E+00 | 4.01E-05 | 5.20E-145 |
| ARS3010 | 2.11E-02 | 9.89E-02 | 6.60E-02 | 7.20E-01 | 3.71E-01 | 8.59E-01 | 2.46E-04 | 4.52E-02 | 4.01E-05 | 1.00E+00 | 4.89E-141 |
| Metagenomes | 5.20E-145 | 4.89E-141 | 4.12E-147 | 4.12E-147 | 5.83E-143 | 5.83E-143 | 4.89E-141 | 4.41E-139 | 5.20E-145 | 4.89E-141 | 1.00E+00 |

Comparison of trees using the Williams & Clifford, 1971 method

| | ARS3001 | ARS3002 | ARS3003 | ARS3004 | ARS3005 | ARS3006 | ARS3007 | ARS3008 | ARS3009 | ARS3010 | Metagenomes |
|---|---|---|---|---|---|---|---|---|---|---|---|
| ARS3001 | 1.00E+00 | 1.87E-03 | 9.45E-04 | 1.66E-02 | 2.86E-02 | 5.82E-02 | 3.36E-01 | 1.66E-02 | 3.11E-04 | 2.55E-03 | 3.10E-97 |
| ARS3002 | 1.87E-03 | 1.00E+00 | 8.56E-01 | 3.74E-01 | 9.71E-01 | 3.36E-01 | 5.82E-02 | 1.00E+00 | 3.38E-11 | 1.00E+00 | 3.10E-97 |
| ARS3003 | 9.45E-04 | 8.56E-01 | 1.00E+00 | 3.74E-01 | 8.56E-01 | 3.74E-01 | 4.80E-02 | 4.79E-01 | 4.83E-12 | 1.00E+00 | 8.83E-85 |
| ARS3004 | 1.66E-02 | 3.74E-01 | 3.74E-01 | 1.00E+00 | 9.71E-01 | 3.36E-01 | 2.70E-01 | 4.79E-01 | 2.01E-07 | 3.10E-01 | 9.54E-87 |
| ARS3005 | 2.86E-02 | 9.71E-01 | 8.56E-01 | 9.71E-01 | 1.00E+00 | 3.36E-01 | 4.35E-01 | 8.56E-01 | 7.98E-09 | 8.13E-01 | 7.51E-91 |
| ARS3006 | 5.82E-02 | 3.36E-01 | 3.74E-01 | 3.36E-01 | 3.36E-01 | 1.00E+00 | 4.79E-01 | 7.32E-01 | 2.37E-08 | 1.82E-01 | 6.49E-104 |
| ARS3007 | 3.36E-01 | 5.82E-02 | 4.80E-02 | 2.70E-01 | 4.35E-01 | 4.79E-01 | 1.00E+00 | 3.10E-01 | 2.45E-06 | 7.27E-02 | 3.48E-96 |
| ARS3008 | 1.66E-02 | 1.00E+00 | 4.79E-01 | 4.79E-01 | 8.56E-01 | 7.32E-01 | 3.10E-01 | 1.00E+00 | 4.14E-10 | 9.28E-01 | 1.74E-100 |
| ARS3009 | 3.11E-04 | 3.38E-11 | 4.83E-12 | 2.01E-07 | 7.98E-09 | 2.37E-08 | 2.45E-06 | 4.14E-10 | 1.00E+00 | 1.31E-12 | 1.12E-122 |
| ARS3010 | 2.55E-03 | 1.00E+00 | 1.00E+00 | 3.10E-01 | 8.13E-01 | 1.82E-01 | 7.27E-02 | 9.28E-01 | 1.31E-12 | 1.00E+00 | 6.13E-93 |
| Metagenomes | 3.10E-97 | 3.10E-97 | 8.83E-85 | 9.54E-87 | 7.51E-91 | 6.49E-104 | 3.48E-96 | 1.74E-100 | 1.12E-122 | 6.13E-93 | 1.00E+00 |

Comparison of trees using the Robinson & Foulds, 1981 method

| | ARS3001 | ARS3002 | ARS3003 | ARS3004 | ARS3005 | ARS3006 | ARS3007 | ARS3008 | ARS3009 | ARS3010 | Metagenomes |
|---|---|---|---|---|---|---|---|---|---|---|---|
| ARS3001 | 1.00E+00 | 6.59E-01 | 6.59E-01 | 9.09E-01 | 2.89E-01 | 1.15E-01 | 3.14E-05 | 8.56E-07 | 4.47E-05 | 3.57E-02 | 1.84E-121 |
| ARS3002 | 6.59E-01 | 1.00E+00 | 1.00E+00 | 4.40E-02 | 1.00E+00 | 3.25E-01 | 1.94E-03 | 2.21E-06 | 7.04E-09 | 2.89E-01 | 9.24E-115 |
| ARS3003 | 6.59E-01 | 1.00E+00 | 1.00E+00 | 4.40E-02 | 1.00E+00 | 6.59E-01 | 1.94E-03 | 4.47E-05 | 7.04E-09 | 3.25E-01 | 9.24E-115 |
| ARS3004 | 9.09E-01 | 4.40E-02 | 4.40E-02 | 1.00E+00 | 9.45E-03 | 3.57E-02 | 3.27E-06 | 1.11E-07 | 4.79E-04 | 9.45E-03 | 9.24E-115 |
| ARS3005 | 2.89E-01 | 1.00E+00 | 1.00E+00 | 9.45E-03 | 1.00E+00 | 3.25E-01 | 3.38E-04 | 3.27E-06 | 2.20E-10 | 1.36E-01 | 2.95E-117 |
| ARS3006 | 1.15E-01 | 3.25E-01 | 6.59E-01 | 3.57E-02 | 3.25E-01 | 1.00E+00 | 2.04E-01 | 1.24E-02 | 6.53E-11 | 1.00E+00 | 6.53E-101 |
| ARS3007 | 3.14E-05 | 1.94E-03 | 1.94E-03 | 3.27E-06 | 3.38E-04 | 2.04E-01 | 1.00E+00 | 4.49E-01 | 2.14E-18 | 1.36E-01 | 3.04E-95 |
| ARS3008 | 8.56E-07 | 2.21E-06 | 4.47E-05 | 1.11E-07 | 3.27E-06 | 1.24E-02 | 4.49E-01 | 1.00E+00 | 1.08E-17 | 1.62E-02 | 8.16E-88 |
| ARS3009 | 4.47E-05 | 7.04E-09 | 7.04E-09 | 4.79E-04 | 2.20E-10 | 6.53E-11 | 2.14E-18 | 1.08E-17 | 1.00E+00 | 1.33E-09 | 8.40E-126 |
| ARS3010 | 3.57E-02 | 2.89E-01 | 3.25E-01 | 9.45E-03 | 1.36E-01 | 1.00E+00 | 1.36E-01 | 1.62E-02 | 1.33E-09 | 1.00E+00 | 8.50E-111 |
| Metagenomes | 1.84E-121 | 9.24E-115 | 9.24E-115 | 9.24E-115 | 2.95E-117 | 6.53E-101 | 3.04E-95 | 8.16E-88 | 8.40E-126 | 8.50E-111 | 1.00E+00 |

Comparison of trees using the Kuhner & Felsenstein, 1994 method

| | ARS3001 | ARS3002 | ARS3003 | ARS3004 | ARS3005 | ARS3006 | ARS3007 | ARS3008 | ARS3009 | ARS3010 | Metagenomes |
|---|---|---|---|---|---|---|---|---|---|---|---|
| ARS3001 | 1.00E+00 | 4.06E-07 | 2.03E-09 | 6.75E-10 | 1.97E-04 | 4.17E-04 | 2.76E-01 | 8.56E-04 | 6.10E-01 | 2.58E-05 | 4.53E-54 |
| ARS3002 | 4.06E-07 | 1.00E+00 | 6.10E-01 | 1.04E-01 | 3.71E-01 | 4.43E-03 | 8.20E-12 | 3.18E-01 | 1.13E-08 | 6.10E-01 | 4.09E-36 |
| ARS3003 | 2.03E-09 | 6.10E-01 | 1.00E+00 | 5.74E-01 | 1.04E-01 | 1.70E-03 | 1.38E-13 | 5.40E-02 | 2.05E-10 | 2.33E-01 | 1.70E-32 |
| ARS3004 | 6.75E-10 | 1.04E-01 | 5.74E-01 | 1.00E+00 | 1.07E-02 | 1.77E-06 | 1.57E-15 | 1.88E-02 | 2.23E-12 | 1.58E-01 | 3.13E-26 |
| ARS3005 | 1.97E-04 | 3.71E-01 | 1.04E-01 | 1.07E-02 | 1.00E+00 | 1.07E-02 | 9.50E-08 | 6.80E-01 | 1.72E-05 | 4.30E-01 | 3.71E-37 |
| ARS3006 | 4.17E-04 | 4.43E-03 | 1.70E-03 | 1.77E-06 | 1.07E-02 | 1.00E+00 | 4.06E-07 | 6.76E-02 | 5.92E-05 | 2.33E-03 | 1.39E-57 |
| ARS3007 | 2.76E-01 | 8.20E-12 | 1.38E-13 | 1.57E-15 | 9.50E-08 | 4.06E-07 | 1.00E+00 | 1.55E-07 | 3.18E-01 | 2.03E-09 | 6.60E-59 |
| ARS3008 | 8.56E-04 | 3.18E-01 | 5.40E-02 | 1.88E-02 | 6.80E-01 | 6.76E-02 | 1.55E-07 | 1.00E+00 | 1.72E-05 | 6.10E-01 | 1.57E-40 |
| ARS3009 | 6.10E-01 | 1.13E-08 | 2.05E-10 | 2.23E-12 | 1.72E-05 | 5.92E-05 | 3.18E-01 | 1.72E-05 | 1.00E+00 | 5.78E-08 | 6.59E-61 |
| ARS3010 | 2.58E-05 | 6.10E-01 | 2.33E-01 | 1.58E-01 | 4.30E-01 | 2.33E-03 | 2.03E-09 | 6.10E-01 | 5.78E-08 | 1.00E+00 | 5.17E-34 |
| Metagenomes | 4.53E-54 | 4.09E-36 | 1.70E-32 | 3.13E-26 | 3.71E-37 | 1.39E-57 | 6.60E-59 | 1.57E-40 | 6.59E-61 | 5.17E-34 | 1.00E+00 |

**Supplementary Table S3C:** Pairwise FDR corrected p-values of the Kolmogorow-Smirnow test on the distances between phylogenetic trees for 16 non-ribosomal proteins in 10 samples of archaeal RefSeq genomes vs one sample containing archaeal MAGs (associated with main Fig. 2c)

Comparison of trees using the Nye et al., 2006 method

| | ARS3001 | ARS3002 | ARS3003 | ARS3004 | ARS3005 | ARS3006 | ARS3007 | ARS3008 | ARS3009 | ARS3010 | Metagenomes |
|---|---|---|---|---|---|---|---|---|---|---|---|
| ARS3001 | 1.00E+00 | 4.21E-03 | 2.04E-02 | 2.81E-02 | 1.24E-06 | 1.65E-03 | 4.21E-03 | 1.40E-07 | 7.64E-02 | 8.81E-01 | 7.99E-68 |
| ARS3002 | 4.21E-03 | 1.00E+00 | 5.01E-01 | 2.48E-01 | 1.42E-01 | 8.81E-01 | 3.19E-01 | 2.81E-02 | 1.86E-05 | 2.04E-02 | 3.82E-66 |
| ARS3003 | 2.04E-02 | 5.01E-01 | 1.00E+00 | 6.98E-01 | 5.76E-02 | 5.97E-01 | 8.11E-01 | 1.41E-02 | 1.65E-03 | 1.89E-01 | 7.99E-68 |
| ARS3004 | 2.81E-02 | 2.48E-01 | 6.98E-01 | 1.00E+00 | 1.41E-02 | 6.98E-01 | 5.01E-01 | 6.31E-03 | 1.65E-03 | 5.76E-02 | 7.99E-68 |
| ARS3005 | 1.24E-06 | 1.42E-01 | 5.76E-02 | 1.41E-02 | 1.00E+00 | 1.05E-01 | 9.68E-03 | 8.81E-01 | 5.19E-09 | 6.16E-05 | 3.82E-66 |
| ARS3006 | 1.65E-03 | 8.81E-01 | 5.97E-01 | 6.98E-01 | 1.05E-01 | 1.00E+00 | 5.97E-01 | 7.64E-02 | 6.16E-05 | 6.31E-03 | 3.82E-66 |
| ARS3007 | 4.21E-03 | 3.19E-01 | 8.11E-01 | 5.01E-01 | 9.68E-03 | 5.97E-01 | 1.00E+00 | 1.11E-03 | 1.00E-05 | 2.81E-02 | 3.82E-66 |
| ARS3008 | 1.40E-07 | 2.81E-02 | 1.41E-02 | 6.31E-03 | 8.81E-01 | 7.64E-02 | 1.11E-03 | 1.00E+00 | 2.32E-09 | 6.24E-07 | 3.82E-66 |
| ARS3009 | 7.64E-02 | 1.86E-05 | 1.65E-03 | 1.65E-03 | 5.19E-09 | 6.16E-05 | 1.00E-05 | 2.32E-09 | 1.00E+00 | 7.64E-02 | 7.99E-68 |
| ARS3010 | 8.81E-01 | 2.04E-02 | 1.89E-01 | 5.76E-02 | 6.16E-05 | 6.31E-03 | 2.81E-02 | 6.24E-07 | 7.64E-02 | 1.00E+00 | 3.82E-66 |
| Metagenomes | 7.99E-68 | 3.82E-66 | 7.99E-68 | 7.99E-68 | 3.82E-66 | 3.82E-66 | 3.82E-66 | 3.82E-66 | 7.99E-68 | 3.82E-66 | 1.00E+00 |

Comparison of trees using the Williams & Clifford, 1971 method

| | ARS3001 | ARS3002 | ARS3003 | ARS3004 | ARS3005 | ARS3006 | ARS3007 | ARS3008 | ARS3009 | ARS3010 | Metagenomes |
|---|---|---|---|---|---|---|---|---|---|---|---|
| ARS3001 | 1.00E+00 | 1.23E-01 | 2.14E-01 | 4.12E-01 | 1.57E-02 | 4.78E-03 | 5.85E-01 | 6.88E-05 | 1.23E-01 | 5.04E-02 | 8.97E-62 |
| ARS3002 | 1.23E-01 | 1.00E+00 | 6.98E-01 | 7.81E-01 | 3.51E-02 | 9.55E-02 | 4.12E-01 | 2.11E-04 | 6.88E-05 | 4.12E-01 | 2.12E-60 |
| ARS3003 | 2.14E-01 | 6.98E-01 | 1.00E+00 | 3.42E-01 | 7.10E-02 | 3.42E-01 | 3.42E-01 | 1.57E-02 | 2.11E-04 | 5.85E-01 | 5.53E-59 |
| ARS3004 | 4.12E-01 | 7.81E-01 | 3.42E-01 | 1.00E+00 | 1.23E-01 | 1.61E-01 | 7.81E-01 | 7.40E-03 | 3.62E-04 | 5.85E-01 | 2.12E-60 |
| ARS3005 | 1.57E-02 | 3.51E-02 | 7.10E-02 | 1.23E-01 | 1.00E+00 | 9.79E-01 | 1.08E-02 | 5.85E-01 | 1.43E-06 | 5.85E-01 | 2.18E-48 |
| ARS3006 | 4.78E-03 | 9.55E-02 | 3.42E-01 | 1.61E-01 | 9.79E-01 | 1.00E+00 | 1.08E-02 | 2.79E-01 | 1.52E-07 | 7.81E-01 | 9.25E-55 |
| ARS3007 | 5.85E-01 | 4.12E-01 | 3.42E-01 | 7.81E-01 | 1.08E-02 | 1.08E-02 | 1.00E+00 | 6.88E-05 | 4.78E-03 | 1.61E-01 | 5.53E-59 |
| ARS3008 | 6.88E-05 | 2.11E-04 | 1.57E-02 | 7.40E-03 | 5.85E-01 | 2.79E-01 | 6.88E-05 | 1.00E+00 | 3.18E-08 | 9.55E-02 | 9.25E-55 |
| ARS3009 | 1.23E-01 | 6.88E-05 | 2.11E-04 | 3.62E-04 | 1.43E-06 | 1.52E-07 | 4.78E-03 | 3.18E-08 | 1.00E+00 | 4.39E-05 | 3.82E-65 |
| ARS3010 | 5.04E-02 | 4.12E-01 | 5.85E-01 | 5.85E-01 | 5.85E-01 | 7.81E-01 | 1.61E-01 | 9.55E-02 | 4.39E-05 | 1.00E+00 | 9.25E-55 |
| Metagenomes | 8.97E-62 | 2.12E-60 | 5.53E-59 | 2.12E-60 | 2.18E-48 | 9.25E-55 | 5.53E-59 | 9.25E-55 | 3.82E-65 | 9.25E-55 | 1.00E+00 |

Comparison of trees using the Robinson & Foulds, 1981 method

| | ARS3001 | ARS3002 | ARS3003 | ARS3004 | ARS3005 | ARS3006 | ARS3007 | ARS3008 | ARS3009 | ARS3010 | Metagenomes |
|---|---|---|---|---|---|---|---|---|---|---|---|
| ARS3001 | 1.00E+00 | 9.86E-02 | 5.22E-02 | 3.94E-02 | 1.18E-02 | 1.27E-01 | 7.35E-02 | 3.62E-04 | 3.04E-03 | 1.00E+00 | 4.77E-66 |
| ARS3002 | 9.86E-02 | 1.00E+00 | 2.73E-01 | 1.27E-01 | 5.22E-02 | 5.22E-02 | 4.42E-01 | 3.62E-04 | 2.69E-08 | 6.65E-01 | 4.77E-66 |
| ARS3003 | 5.22E-02 | 2.73E-01 | 1.00E+00 | 1.00E+00 | 9.89E-01 | 1.00E+00 | 1.00E+00 | 2.73E-01 | 5.75E-08 | 2.73E-01 | 4.77E-66 |
| ARS3004 | 3.94E-02 | 1.27E-01 | 1.00E+00 | 1.00E+00 | 1.00E+00 | 1.00E+00 | 9.89E-01 | 2.20E-01 | 1.16E-06 | 2.20E-01 | 4.77E-66 |
| ARS3005 | 1.18E-02 | 5.22E-02 | 9.89E-01 | 1.00E+00 | 1.00E+00 | 9.89E-01 | 9.89E-01 | 5.47E-01 | 5.19E-09 | 9.86E-02 | 8.47E-61 |
| ARS3006 | 1.27E-01 | 5.22E-02 | 1.00E+00 | 1.00E+00 | 9.89E-01 | 1.00E+00 | 7.91E-01 | 3.94E-02 | 1.21E-07 | 2.20E-01 | 4.77E-66 |
| ARS3007 | 7.35E-02 | 4.42E-01 | 1.00E+00 | 9.89E-01 | 9.89E-01 | 7.91E-01 | 1.00E+00 | 1.27E-01 | 2.29E-06 | 3.50E-01 | 4.77E-66 |
| ARS3008 | 3.62E-04 | 3.62E-04 | 2.73E-01 | 2.20E-01 | 5.47E-01 | 3.94E-02 | 1.27E-01 | 1.00E+00 | 9.76E-14 | 4.78E-03 | 1.99E-62 |
| ARS3009 | 3.04E-03 | 2.69E-08 | 5.75E-08 | 1.16E-06 | 5.19E-09 | 1.21E-07 | 2.29E-06 | 9.76E-14 | 1.00E+00 | 1.21E-04 | 3.19E-67 |
| ARS3010 | 1.00E+00 | 6.65E-01 | 2.73E-01 | 2.20E-01 | 9.86E-02 | 2.20E-01 | 3.50E-01 | 4.78E-03 | 1.21E-04 | 1.00E+00 | 4.77E-66 |
| Metagenomes | 4.77E-66 | 4.77E-66 | 4.77E-66 | 4.77E-66 | 8.47E-61 | 4.77E-66 | 4.77E-66 | 1.99E-62 | 3.19E-67 | 4.77E-66 | 1.00E+00 |

Comparison of trees using the Kuhner & Felsenstein, 1994 method

| | ARS3001 | ARS3002 | ARS3003 | ARS3004 | ARS3005 | ARS3006 | ARS3007 | ARS3008 | ARS3009 | ARS3010 | Metagenomes |
|---|---|---|---|---|---|---|---|---|---|---|---|
| ARS3001 | 1.00E+00 | 9.69E-02 | 7.95E-01 | 4.42E-01 | 2.33E-01 | 7.95E-01 | 7.95E-01 | 7.95E-01 | 1.81E-01 | 3.62E-02 | 7.57E-64 |
| ARS3002 | 9.69E-02 | 1.00E+00 | 1.81E-01 | 4.42E-01 | 4.42E-01 | 2.65E-02 | 1.18E-01 | 9.69E-02 | 1.81E-01 | 2.65E-02 | 1.06E-60 |
| ARS3003 | 7.95E-01 | 1.81E-01 | 1.00E+00 | 8.81E-01 | 2.33E-01 | 3.88E-01 | 7.57E-01 | 4.42E-01 | 5.18E-02 | 3.62E-02 | 3.19E-67 |
| ARS3004 | 4.42E-01 | 4.42E-01 | 8.81E-01 | 1.00E+00 | 7.95E-01 | 3.88E-01 | 7.95E-01 | 5.35E-01 | 1.54E-01 | 1.81E-01 | 3.23E-56 |
| ARS3005 | 2.33E-01 | 4.42E-01 | 2.33E-01 | 7.95E-01 | 1.00E+00 | 1.18E-01 | 7.57E-01 | 1.18E-01 | 1.94E-02 | 4.42E-01 | 1.06E-60 |
| ARS3006 | 7.95E-01 | 2.65E-02 | 3.88E-01 | 3.88E-01 | 1.18E-01 | 1.00E+00 | 6.50E-01 | 8.81E-01 | 7.30E-02 | 1.31E-02 | 1.91E-65 |
| ARS3007 | 7.95E-01 | 1.18E-01 | 7.57E-01 | 7.95E-01 | 7.57E-01 | 6.50E-01 | 1.00E+00 | 5.35E-01 | 1.18E-01 | 1.54E-01 | 2.99E-62 |
| ARS3008 | 7.95E-01 | 9.69E-02 | 4.42E-01 | 5.35E-01 | 1.18E-01 | 8.81E-01 | 5.35E-01 | 1.00E+00 | 1.81E-01 | 1.31E-02 | 7.57E-64 |
| ARS3009 | 1.81E-01 | 1.81E-01 | 5.18E-02 | 1.54E-01 | 1.94E-02 | 7.30E-02 | 1.18E-01 | 1.81E-01 | 1.00E+00 | 3.65E-04 | 3.23E-56 |
| ARS3010 | 3.62E-02 | 2.65E-02 | 3.62E-02 | 1.81E-01 | 4.42E-01 | 1.31E-02 | 1.54E-01 | 1.31E-02 | 3.65E-04 | 1.00E+00 | 2.99E-62 |
| Metagenomes | 7.57E-64 | 1.06E-60 | 3.19E-67 | 3.23E-56 | 1.06E-60 | 1.91E-65 | 2.99E-62 | 7.57E-64 | 3.23E-56 | 2.99E-62 | 1.00E+00 |

**Supplementary Table S3D:** Pairwise FDR corrected p-values of the Kolmogorow-Smirnow test on the distances between phylogenetic trees for ribosomal proteins in 10 samples of bacterial RefSeq genomes vs one sample containing CPR MAGs (associated with main Fig. 2d)

Comparison of trees using the Nye et al., 2006 method

| | BRS3001 | BRS3002 | BRS3003 | BRS3004 | BRS3005 | BRS3006 | BRS3007 | BRS3008 | BRS3009 | BRS3010 | Meta_CPR3004 |
|---|---|---|---|---|---|---|---|---|---|---|---|
| BRS3001 | 1.00E+00 | 1.39E-01 | 1.81E-01 | 6.71E-08 | 2.76E-01 | 1.29E-04 | 2.01E-04 | 6.08E-03 | 1.81E-01 | 4.30E-02 | 9.62E-53 |
| BRS3002 | 1.39E-01 | 1.00E+00 | 5.92E-02 | 5.66E-09 | 1.52E-03 | 1.15E-05 | 4.10E-05 | 2.01E-04 | 6.08E-03 | 7.31E-05 | 3.23E-55 |
| BRS3003 | 1.81E-01 | 5.92E-02 | 1.00E+00 | 2.01E-04 | 2.27E-01 | 1.41E-02 | 3.07E-02 | 1.11E-01 | 2.76E-01 | 1.39E-01 | 8.36E-37 |
| BRS3004 | 6.71E-08 | 5.66E-09 | 2.01E-04 | 1.00E+00 | 1.43E-06 | 2.76E-01 | 2.27E-01 | 4.04E-03 | 5.82E-04 | 9.34E-03 | 1.47E-42 |
| BRS3005 | 2.76E-01 | 1.52E-03 | 2.27E-01 | 1.43E-06 | 1.00E+00 | 3.44E-04 | 1.52E-03 | 4.30E-02 | 5.42E-01 | 1.11E-01 | 3.36E-49 |
| BRS3006 | 1.29E-04 | 1.15E-05 | 1.41E-02 | 2.76E-01 | 3.44E-04 | 1.00E+00 | 7.66E-01 | 1.39E-01 | 1.52E-03 | 1.11E-01 | 8.36E-37 |
| BRS3007 | 2.01E-04 | 4.10E-05 | 3.07E-02 | 2.27E-01 | 1.52E-03 | 7.66E-01 | 1.00E+00 | 5.92E-02 | 3.07E-02 | 2.27E-01 | 6.22E-36 |
| BRS3008 | 6.08E-03 | 2.01E-04 | 1.11E-01 | 4.04E-03 | 4.30E-02 | 1.39E-01 | 5.92E-02 | 1.00E+00 | 1.39E-01 | 4.44E-01 | 1.39E-41 |
| BRS3009 | 1.81E-01 | 6.08E-03 | 2.76E-01 | 5.82E-04 | 5.42E-01 | 1.52E-03 | 3.07E-02 | 1.39E-01 | 1.00E+00 | 2.76E-01 | 2.56E-50 |
| BRS3010 | 4.30E-02 | 7.31E-05 | 1.39E-01 | 9.34E-03 | 1.11E-01 | 1.11E-01 | 2.27E-01 | 4.44E-01 | 2.76E-01 | 1.00E+00 | 9.25E-46 |
| Meta_CPR3004 | 9.62E-53 | 3.23E-55 | 8.36E-37 | 1.47E-42 | 3.36E-49 | 8.36E-37 | 6.22E-36 | 1.39E-41 | 2.56E-50 | 9.25E-46 | 1.00E+00 |

Comparison of trees using the Williams & Clifford, 1971 method

| | BRS3001 | BRS3002 | BRS3003 | BRS3004 | BRS3005 | BRS3006 | BRS3007 | BRS3008 | BRS3009 | BRS3010 | Meta_CPR3004 |
|---|---|---|---|---|---|---|---|---|---|---|---|
| BRS3001 | 1.00E+00 | 2.99E-01 | 6.71E-01 | 2.32E-09 | 2.42E-01 | 4.39E-05 | 1.11E-03 | 1.04E-02 | 5.01E-03 | 2.36E-04 | 1.93E-27 |
| BRS3002 | 2.99E-01 | 1.00E+00 | 5.63E-01 | 7.80E-05 | 3.71E-01 | 1.04E-02 | 1.46E-01 | 1.94E-01 | 3.71E-01 | 8.49E-02 | 4.86E-23 |
| BRS3003 | 6.71E-01 | 5.63E-01 | 1.00E+00 | 1.55E-06 | 2.99E-01 | 1.11E-03 | 1.04E-02 | 8.49E-02 | 1.11E-01 | 1.04E-02 | 6.68E-22 |
| BRS3004 | 2.32E-09 | 7.80E-05 | 1.55E-06 | 1.00E+00 | 1.37E-04 | 2.99E-01 | 1.11E-01 | 2.25E-02 | 4.39E-05 | 4.02E-04 | 1.07E-13 |
| BRS3005 | 2.42E-01 | 3.71E-01 | 2.99E-01 | 1.37E-04 | 1.00E+00 | 7.74E-03 | 8.49E-02 | 1.46E-01 | 2.42E-01 | 8.49E-02 | 4.86E-23 |
| BRS3006 | 4.39E-05 | 1.04E-02 | 1.11E-03 | 2.99E-01 | 7.74E-03 | 1.00E+00 | 7.81E-01 | 4.61E-01 | 1.51E-02 | 4.71E-02 | 1.29E-15 |
| BRS3007 | 1.11E-03 | 1.46E-01 | 1.04E-02 | 1.11E-01 | 8.49E-02 | 7.81E-01 | 1.00E+00 | 1.00E+00 | 1.51E-02 | 1.11E-01 | 3.87E-14 |
| BRS3008 | 1.04E-02 | 1.94E-01 | 8.49E-02 | 2.25E-02 | 1.46E-01 | 4.61E-01 | 1.00E+00 | 1.00E+00 | 8.49E-02 | 2.42E-01 | 1.29E-15 |
| BRS3009 | 5.01E-03 | 3.71E-01 | 1.11E-01 | 4.39E-05 | 2.42E-01 | 1.51E-02 | 1.51E-02 | 8.49E-02 | 1.00E+00 | 3.28E-02 | 6.68E-22 |
| BRS3010 | 2.36E-04 | 8.49E-02 | 1.04E-02 | 4.02E-04 | 8.49E-02 | 4.71E-02 | 1.11E-01 | 2.42E-01 | 3.28E-02 | 1.00E+00 | 2.05E-23 |
| Meta_CPR3004 | 1.93E-27 | 4.86E-23 | 6.68E-22 | 1.07E-13 | 4.86E-23 | 1.29E-15 | 3.87E-14 | 1.29E-15 | 6.68E-22 | 2.05E-23 | 1.00E+00 |

Comparison of trees using the Robinson & Foulds, 1981 method

| | BRS3001 | BRS3002 | BRS3003 | BRS3004 | BRS3005 | BRS3006 | BRS3007 | BRS3008 | BRS3009 | BRS3010 | Meta_CPR3004 |
|---|---|---|---|---|---|---|---|---|---|---|---|
| BRS3001 | 1.00E+00 | 2.60E-01 | 6.27E-02 | 7.80E-05 | 1.00E+00 | 7.53E-04 | 2.03E-01 | 4.71E-02 | 8.43E-01 | 4.71E-02 | 2.01E-51 |
| BRS3002 | 2.60E-01 | 1.00E+00 | 4.71E-02 | 1.36E-08 | 8.73E-02 | 6.64E-06 | 6.27E-02 | 7.40E-03 | 3.94E-02 | 1.37E-04 | 8.22E-47 |
| BRS3003 | 6.27E-02 | 4.71E-02 | 1.00E+00 | 2.00E-03 | 4.71E-02 | 4.71E-02 | 1.00E+00 | 9.50E-01 | 2.60E-01 | 4.22E-01 | 1.46E-40 |
| BRS3004 | 7.80E-05 | 1.36E-08 | 2.00E-03 | 1.00E+00 | 4.39E-05 | 7.26E-01 | 5.01E-03 | 6.27E-02 | 2.00E-03 | 1.20E-01 | 1.46E-40 |
| BRS3005 | 1.00E+00 | 8.73E-02 | 4.71E-02 | 4.39E-05 | 1.00E+00 | 4.39E-05 | 4.71E-02 | 7.40E-03 | 1.00E+00 | 3.94E-02 | 2.01E-51 |
| BRS3006 | 7.53E-04 | 6.64E-06 | 4.71E-02 | 7.26E-01 | 4.39E-05 | 1.00E+00 | 1.53E-01 | 6.10E-01 | 5.01E-03 | 5.12E-01 | 9.68E-38 |
| BRS3007 | 2.03E-01 | 6.27E-02 | 1.00E+00 | 5.01E-03 | 4.71E-02 | 1.53E-01 | 1.00E+00 | 1.00E+00 | 5.12E-01 | 1.53E-01 | 1.20E-38 |
| BRS3008 | 4.71E-02 | 7.40E-03 | 9.50E-01 | 6.27E-02 | 7.40E-03 | 6.10E-01 | 1.00E+00 | 1.00E+00 | 1.53E-01 | 6.10E-01 | 3.75E-34 |
| BRS3009 | 8.43E-01 | 3.94E-02 | 2.60E-01 | 2.00E-03 | 1.00E+00 | 5.01E-03 | 5.12E-01 | 1.53E-01 | 1.00E+00 | 3.34E-01 | 8.22E-47 |
| BRS3010 | 4.71E-02 | 1.37E-04 | 4.22E-01 | 1.20E-01 | 3.94E-02 | 5.12E-01 | 1.53E-01 | 6.10E-01 | 3.34E-01 | 1.00E+00 | 1.51E-43 |
| Meta_CPR3004 | 2.01E-51 | 8.22E-47 | 1.46E-40 | 1.46E-40 | 2.01E-51 | 9.68E-38 | 1.20E-38 | 3.75E-34 | 8.22E-47 | 1.51E-43 | 1.00E+00 |

Comparison of trees using the Kuhner & Felsenstein, 1994 method

| | BRS3001 | BRS3002 | BRS3003 | BRS3004 | BRS3005 | BRS3006 | BRS3007 | BRS3008 | BRS3009 | BRS3010 | Meta_CPR3004 |
|---|---|---|---|---|---|---|---|---|---|---|---|
| BRS3001 | 1.00E+00 | 3.32E-02 | 3.64E-07 | 1.03E-15 | 3.65E-11 | 4.03E-08 | 1.75E-07 | 7.81E-01 | 1.72E-03 | 3.77E-04 | 5.58E-25 |
| BRS3002 | 3.32E-02 | 1.00E+00 | 1.06E-02 | 4.03E-08 | 3.77E-04 | 4.37E-03 | 1.59E-02 | 6.37E-02 | 2.18E-01 | 8.66E-08 | 2.63E-18 |
| BRS3003 | 3.64E-07 | 1.06E-02 | 1.00E+00 | 2.33E-04 | 4.74E-02 | 7.81E-01 | 9.79E-01 | 2.05E-05 | 6.37E-02 | 2.85E-15 | 1.03E-15 |
| BRS3004 | 1.03E-15 | 4.03E-08 | 2.33E-04 | 1.00E+00 | 3.32E-02 | 6.36E-04 | 2.05E-05 | 1.84E-12 | 2.95E-06 | 5.58E-25 | 7.44E-07 |
| BRS3005 | 3.65E-11 | 3.77E-04 | 4.74E-02 | 3.32E-02 | 1.00E+00 | 2.81E-01 | 8.80E-02 | 1.75E-07 | 4.37E-03 | 2.63E-18 | 3.46E-09 |
| BRS3006 | 4.03E-08 | 4.37E-03 | 7.81E-01 | 6.36E-04 | 2.81E-01 | 1.00E+00 | 5.63E-01 | 1.50E-06 | 2.34E-02 | 2.63E-18 | 6.83E-13 |
| BRS3007 | 1.75E-07 | 1.59E-02 | 9.79E-01 | 2.05E-05 | 8.80E-02 | 5.63E-01 | 1.00E+00 | 2.05E-05 | 6.37E-02 | 2.85E-15 | 1.03E-15 |
| BRS3008 | 7.81E-01 | 6.37E-02 | 2.05E-05 | 1.84E-12 | 1.75E-07 | 1.50E-06 | 2.05E-05 | 1.00E+00 | 1.72E-03 | 3.77E-04 | 2.10E-24 |
| BRS3009 | 1.72E-03 | 2.18E-01 | 6.37E-02 | 2.95E-06 | 4.37E-03 | 2.34E-02 | 6.37E-02 | 1.72E-03 | 1.00E+00 | 1.50E-09 | 2.49E-13 |
| BRS3010 | 3.77E-04 | 8.66E-08 | 2.85E-15 | 5.58E-25 | 2.63E-18 | 2.63E-18 | 2.85E-15 | 3.77E-04 | 1.50E-09 | 1.00E+00 | 3.75E-33 |
| Meta_CPR3004 | 5.58E-25 | 2.63E-18 | 1.03E-15 | 7.44E-07 | 3.46E-09 | 6.83E-13 | 1.03E-15 | 2.10E-24 | 2.49E-13 | 3.75E-33 | 1.00E+00 |

**Supplementary Table S3E:** Pairwise FDR corrected p-values of the Kolmogorow-Smirnow test on the distances between phylogenetic trees for 20 ribosomal proteins in 10 samples of bacterial RefSeq genomes vs one sample containing bacterial MAGs (associated with main Fig. 2e)

Comparison of trees using the Nye et al., 2006 method

| | BRS3001 | BRS3002 | BRS3003 | BRS3004 | BRS3005 | BRS3006 | BRS3007 | BRS3008 | BRS3009 | BRS3010 | Meta_BMS3002 |
|---|---|---|---|---|---|---|---|---|---|---|---|
| BRS3001 | 1.00E+00 | 5.63E-01 | 1.65E-02 | 1.53E-12 | 2.83E-01 | 2.45E-06 | 4.64E-04 | 1.59E-01 | 8.47E-03 | 4.16E-03 | 1.29E-97 |
| BRS3002 | 5.63E-01 | 1.00E+00 | 2.24E-02 | 3.16E-13 | 1.33E-01 | 2.60E-07 | 4.64E-04 | 1.05E-01 | 1.44E-03 | 4.64E-04 | 2.73E-90 |
| BRS3003 | 1.65E-02 | 2.24E-02 | 1.00E+00 | 7.08E-06 | 4.99E-02 | 2.92E-03 | 5.63E-01 | 2.83E-01 | 8.20E-02 | 8.20E-02 | 6.00E-89 |
| BRS3004 | 1.53E-12 | 3.16E-13 | 7.08E-06 | 1.00E+00 | 2.16E-08 | 3.89E-02 | 1.44E-03 | 8.15E-07 | 1.15E-05 | 2.13E-04 | 1.42E-87 |
| BRS3005 | 2.83E-01 | 1.33E-01 | 4.99E-02 | 2.16E-08 | 1.00E+00 | 1.44E-03 | 2.92E-03 | 2.38E-01 | 1.59E-01 | 2.93E-02 | 1.29E-97 |
| BRS3006 | 2.45E-06 | 2.60E-07 | 2.92E-03 | 3.89E-02 | 1.44E-03 | 1.00E+00 | 4.99E-02 | 1.39E-04 | 2.93E-02 | 1.59E-01 | 2.73E-90 |
| BRS3007 | 4.64E-04 | 4.64E-04 | 5.63E-01 | 1.44E-03 | 2.92E-03 | 4.99E-02 | 1.00E+00 | 8.20E-02 | 1.93E-01 | 1.93E-01 | 6.00E-89 |
| BRS3008 | 1.59E-01 | 1.05E-01 | 2.83E-01 | 8.15E-07 | 2.38E-01 | 1.39E-04 | 8.20E-02 | 1.00E+00 | 1.07E-03 | 2.10E-03 | 4.01E-96 |
| BRS3009 | 8.47E-03 | 1.44E-03 | 8.20E-02 | 1.15E-05 | 1.59E-01 | 2.93E-02 | 1.93E-01 | 1.07E-03 | 1.00E+00 | 5.63E-01 | 3.69E-93 |
| BRS3010 | 4.16E-03 | 4.64E-04 | 8.20E-02 | 2.13E-04 | 2.93E-02 | 1.59E-01 | 1.93E-01 | 2.10E-03 | 5.63E-01 | 1.00E+00 | 1.24E-94 |
| Meta_BMS3002 | 1.29E-97 | 2.73E-90 | 6.00E-89 | 1.42E-87 | 1.29E-97 | 2.73E-90 | 6.00E-89 | 4.01E-96 | 3.69E-93 | 1.24E-94 | 1.00E+00 |

Comparison of trees using the Williams & Clifford, 1971 method

| | BRS3001 | BRS3002 | BRS3003 | BRS3004 | BRS3005 | BRS3006 | BRS3007 | BRS3008 | BRS3009 | BRS3010 | Meta_BMS3002 |
|---|---|---|---|---|---|---|---|---|---|---|---|
| BRS3001 | 1.00E+00 | 2.80E-06 | 5.89E-02 | 3.85E-10 | 1.08E-02 | 1.69E-06 | 8.74E-04 | 2.72E-02 | 8.63E-08 | 1.38E-05 | 3.46E-67 |
| BRS3002 | 2.80E-06 | 1.00E+00 | 3.63E-02 | 1.08E-02 | 1.78E-01 | 4.38E-01 | 4.38E-01 | 7.65E-02 | 3.28E-01 | 6.55E-01 | 1.08E-47 |
| BRS3003 | 5.89E-02 | 3.63E-02 | 1.00E+00 | 1.65E-04 | 5.73E-01 | 5.61E-03 | 1.46E-01 | 4.95E-01 | 5.61E-03 | 9.83E-02 | 2.73E-53 |
| BRS3004 | 3.85E-10 | 1.08E-02 | 1.65E-04 | 1.00E+00 | 3.76E-04 | 2.20E-01 | 5.61E-03 | 3.76E-04 | 5.89E-02 | 5.61E-03 | 6.99E-28 |
| BRS3005 | 1.08E-02 | 1.78E-01 | 5.73E-01 | 3.76E-04 | 1.00E+00 | 1.46E-01 | 1.46E-01 | 1.46E-01 | 5.89E-02 | 2.70E-01 | 9.12E-56 |
| BRS3006 | 1.69E-06 | 4.38E-01 | 5.61E-03 | 2.20E-01 | 1.46E-01 | 1.00E+00 | 1.46E-01 | 2.72E-02 | 4.38E-01 | 4.95E-01 | 9.75E-40 |
| BRS3007 | 8.74E-04 | 4.38E-01 | 1.46E-01 | 5.61E-03 | 1.46E-01 | 1.46E-01 | 1.00E+00 | 4.95E-01 | 1.78E-01 | 3.93E-01 | 9.75E-40 |
| BRS3008 | 2.72E-02 | 7.65E-02 | 4.95E-01 | 3.76E-04 | 1.46E-01 | 2.72E-02 | 4.95E-01 | 1.00E+00 | 2.10E-02 | 5.89E-02 | 6.60E-44 |
| BRS3009 | 8.63E-08 | 3.28E-01 | 5.61E-03 | 5.89E-02 | 5.89E-02 | 4.38E-01 | 1.78E-01 | 2.10E-02 | 1.00E+00 | 4.38E-01 | 3.15E-43 |
| BRS3010 | 1.38E-05 | 6.55E-01 | 9.83E-02 | 5.61E-03 | 2.70E-01 | 4.95E-01 | 3.93E-01 | 5.89E-02 | 4.38E-01 | 1.00E+00 | 4.99E-50 |
| Meta_BMS3002 | 3.46E-67 | 1.08E-47 | 2.73E-53 | 6.99E-28 | 9.12E-56 | 9.75E-40 | 9.75E-40 | 6.60E-44 | 3.15E-43 | 4.99E-50 | 1.00E+00 |

Comparison of trees using the Robinson & Foulds, 1981 method

| | BRS3001 | BRS3002 | BRS3003 | BRS3004 | BRS3005 | BRS3006 | BRS3007 | BRS3008 | BRS3009 | BRS3010 | Meta_BMS3002 |
|---|---|---|---|---|---|---|---|---|---|---|---|
| BRS3001 | 1.00E+00 | 8.09E-01 | 1.28E-03 | 2.55E-08 | 1.00E+00 | 1.63E-07 | 1.25E-01 | 3.28E-01 | 1.01E-01 | 6.08E-02 | 1.03E-102 |
| BRS3002 | 8.09E-01 | 1.00E+00 | 2.93E-02 | 4.93E-06 | 2.83E-01 | 2.20E-05 | 6.19E-01 | 1.00E+00 | 4.80E-02 | 4.80E-02 | 2.63E-93 |
| BRS3003 | 1.28E-03 | 2.93E-02 | 1.00E+00 | 4.80E-02 | 1.55E-02 | 1.01E-01 | 3.28E-01 | 1.54E-01 | 6.08E-02 | 2.37E-01 | 3.45E-86 |
| BRS3004 | 2.55E-08 | 4.93E-06 | 4.80E-02 | 1.00E+00 | 1.69E-06 | 3.28E-01 | 6.14E-03 | 3.99E-04 | 1.28E-03 | 1.55E-02 | 3.45E-86 |
| BRS3005 | 1.00E+00 | 2.83E-01 | 1.55E-02 | 1.69E-06 | 1.00E+00 | 8.02E-06 | 4.47E-01 | 3.77E-01 | 1.25E-01 | 2.83E-01 | 3.01E-96 |
| BRS3006 | 1.63E-07 | 2.20E-05 | 1.01E-01 | 3.28E-01 | 8.02E-06 | 1.00E+00 | 1.55E-02 | 9.23E-04 | 8.55E-03 | 3.90E-02 | 2.63E-93 |
| BRS3007 | 1.25E-01 | 6.19E-01 | 3.28E-01 | 6.14E-03 | 4.47E-01 | 1.55E-02 | 1.00E+00 | 1.00E+00 | 3.77E-01 | 3.77E-01 | 7.75E-92 |
| BRS3008 | 3.28E-01 | 1.00E+00 | 1.54E-01 | 3.99E-04 | 3.77E-01 | 9.23E-04 | 1.00E+00 | 1.00E+00 | 1.92E-01 | 1.54E-01 | 9.94E-95 |
| BRS3009 | 1.01E-01 | 4.80E-02 | 6.08E-02 | 1.28E-03 | 1.25E-01 | 8.55E-03 | 3.77E-01 | 1.92E-01 | 1.00E+00 | 1.00E+00 | 3.01E-96 |
| BRS3010 | 6.08E-02 | 4.80E-02 | 2.37E-01 | 1.55E-02 | 2.83E-01 | 3.90E-02 | 3.77E-01 | 1.54E-01 | 1.00E+00 | 1.00E+00 | 1.29E-97 |
| Meta_BMS3002 | 1.03E-102 | 2.63E-93 | 3.45E-86 | 3.45E-86 | 3.01E-96 | 2.63E-93 | 7.75E-92 | 9.94E-95 | 3.01E-96 | 1.29E-97 | 1.00E+00 |

Comparison of trees using the Kuhner & Felsenstein, 1994 method

| | BRS3001 | BRS3002 | BRS3003 | BRS3004 | BRS3005 | BRS3006 | BRS3007 | BRS3008 | BRS3009 | BRS3010 | Meta_BMS3002 |
|---|---|---|---|---|---|---|---|---|---|---|---|
| BRS3001 | 1.00E+00 | 5.70E-03 | 6.75E-10 | 8.04E-13 | 4.97E-13 | 4.85E-09 | 1.65E-08 | 1.07E-02 | 9.76E-08 | 1.91E-02 | 3.60E-29 |
| BRS3002 | 5.70E-03 | 1.00E+00 | 5.54E-04 | 1.05E-06 | 1.53E-05 | 7.98E-03 | 1.44E-02 | 3.20E-01 | 1.91E-02 | 3.53E-05 | 1.37E-17 |
| BRS3003 | 6.75E-10 | 5.54E-04 | 1.00E+00 | 4.29E-01 | 6.93E-01 | 4.29E-01 | 4.29E-01 | 3.53E-05 | 4.29E-01 | 8.04E-13 | 5.47E-06 |
| BRS3004 | 8.04E-13 | 1.05E-06 | 4.29E-01 | 1.00E+00 | 2.64E-01 | 2.51E-02 | 1.44E-02 | 5.45E-08 | 2.51E-02 | 7.25E-15 | 3.53E-05 |
| BRS3005 | 4.97E-13 | 1.53E-05 | 6.93E-01 | 2.64E-01 | 1.00E+00 | 3.85E-01 | 8.01E-02 | 3.53E-05 | 2.15E-01 | 1.38E-15 | 2.47E-05 |
| BRS3006 | 4.85E-09 | 7.98E-03 | 4.29E-01 | 2.51E-02 | 3.85E-01 | 1.00E+00 | 1.00E+00 | 5.70E-03 | 7.79E-01 | 8.04E-13 | 3.01E-08 |
| BRS3007 | 1.65E-08 | 1.44E-02 | 4.29E-01 | 1.44E-02 | 8.01E-02 | 1.00E+00 | 1.00E+00 | 5.70E-03 | 9.46E-01 | 3.77E-12 | 4.85E-09 |
| BRS3008 | 1.07E-02 | 3.20E-01 | 3.53E-05 | 5.45E-08 | 3.53E-05 | 5.70E-03 | 5.70E-03 | 1.00E+00 | 1.07E-02 | 5.60E-05 | 3.02E-19 |
| BRS3009 | 9.76E-08 | 1.91E-02 | 4.29E-01 | 2.51E-02 | 2.15E-01 | 7.79E-01 | 9.46E-01 | 1.07E-02 | 1.00E+00 | 3.53E-10 | 1.65E-08 |
| BRS3010 | 1.91E-02 | 3.53E-05 | 8.04E-13 | 7.25E-15 | 1.38E-15 | 8.04E-13 | 3.77E-12 | 5.60E-05 | 3.53E-10 | 1.00E+00 | 6.88E-29 |
| Meta_BMS3002 | 3.60E-29 | 1.37E-17 | 5.47E-06 | 3.53E-05 | 2.47E-05 | 3.01E-08 | 4.85E-09 | 3.02E-19 | 1.65E-08 | 6.88E-29 | 1.00E+00 |

**Supplementary Table S4A:** Pairwise FDR corrected p-values of the Kolmogorow-Smirnow test on the distances between phylogenetic trees for 39 universal proteins in 10 samples of archaeal RefSeq genomes vs 10 samples of archaeal MAGs (associated with Supplementary Fig. S7a)

Comparison of trees using the Nye et al., 2006 method

| | ARS3001 | ARS3002 | ARS3003 | ARS3004 | ARS3005 | ARS3006 | ARS3007 | ARS3008 | ARS3009 | ARS3010 | Meta_AMS3002 | Meta_AMS3003 | Meta_AMS3004 | Meta_AMS3005 | Meta_AMS3006 | Meta_AMS3007 | Meta_AMS3008 | Meta_AMS3009 | Meta_AMS3010 | Meta_AMS3011 |
|---|---|---|---|---|---|---|---|---|---|---|---|---|---|---|---|---|---|---|---|---|
| ARS3001 | 1.00E+00 | 9.88E-15 | 2.74E-03 | 6.09E-06 | 7.75E-25 | 7.55E-09 | 3.36E-20 | 6.35E-23 | 2.52E-01 | 9.30E-05 | 8.82E-112 | 3.64E-93 | 5.45E-146 | 9.73E-255 | 5.92E-150 | 1.62E-167 | 4.04E-140 | 5.45E-146 | 5.81E-169 | 3.81E-211 |
| ARS3002 | 9.88E-15 | 1.00E+00 | 6.97E-07 | 2.85E-04 | 1.33E-02 | 3.96E-03 | 1.44E-01 | 7.78E-02 | 6.62E-19 | 5.81E-05 | 7.09E-85 | 4.21E-62 | 1.15E-93 | 3.23E-180 | 1.11E-91 | 1.11E-95 | 2.65E-86 | 4.74E-82 | 2.89E-98 | 1.12E-152 |
| ARS3003 | 2.74E-03 | 6.97E-07 | 1.00E+00 | 3.72E-01 | 2.98E-12 | 1.80E-02 | 7.55E-09 | 6.00E-11 | 1.32E-05 | 7.04E-01 | 2.36E-100 | 1.62E-82 | 5.84E-127 | 9.74E-231 | 4.98E-130 | 5.06E-144 | 3.79E-122 | 9.44E-126 | 1.29E-148 | 1.48E-189 |
| ARS3004 | 6.09E-06 | 2.85E-04 | 3.72E-01 | 1.00E+00 | 4.95E-08 | 3.07E-01 | 2.13E-06 | 5.26E-07 | 7.55E-09 | 8.76E-01 | 3.68E-94 | 8.93E-75 | 2.50E-116 | 2.38E-216 | 1.25E-118 | 4.98E-130 | 4.13E-110 | 1.82E-113 | 5.37E-135 | 3.23E-180 |
| ARS3005 | 7.75E-25 | 1.33E-02 | 2.98E-12 | 4.95E-08 | 1.00E+00 | 4.72E-06 | 6.62E-01 | 2.79E-01 | 3.45E-32 | 2.54E-10 | 7.85E-86 | 6.49E-61 | 3.61E-92 | 2.07E-173 | 9.90E-89 | 3.27E-89 | 1.62E-82 | 2.26E-78 | 1.11E-91 | 5.65E-148 |
| ARS3006 | 7.55E-09 | 3.96E-03 | 1.80E-02 | 3.07E-01 | 4.72E-06 | 1.00E+00 | 1.83E-04 | 1.32E-05 | 2.74E-13 | 1.28E-01 | 9.90E-89 | 6.10E-65 | 1.52E-103 | 7.72E-197 | 1.05E-105 | 4.60E-118 | 8.80E-99 | 7.08E-101 | 5.73E-121 | 8.83E-162 |
| ARS3007 | 3.36E-20 | 1.44E-01 | 7.55E-09 | 2.13E-06 | 6.62E-01 | 1.83E-04 | 1.00E+00 | 4.46E-01 | 2.53E-26 | 6.66E-08 | 7.85E-86 | 6.49E-61 | 3.45E-90 | 3.57E-177 | 3.27E-89 | 3.61E-92 | 1.89E-83 | 1.38E-81 | 1.15E-93 | 5.65E-148 |
| ARS3008 | 6.35E-23 | 7.78E-02 | 6.00E-11 | 5.26E-07 | 2.79E-01 | 1.32E-05 | 4.46E-01 | 1.00E+00 | 2.51E-25 | 1.42E-08 | 9.55E-80 | 2.23E-54 | 3.95E-81 | 3.94E-156 | 5.19E-77 | 3.95E-81 | 3.90E-72 | 2.84E-68 | 1.14E-80 | 1.76E-132 |
| ARS3009 | 2.52E-01 | 6.62E-19 | 1.32E-05 | 7.55E-09 | 3.45E-32 | 2.74E-13 | 2.53E-26 | 2.51E-25 | 1.00E+00 | 2.96E-07 | 2.42E-107 | 9.34E-88 | 4.54E-142 | 5.43E-245 | 2.45E-145 | 6.42E-164 | 1.28E-135 | 4.54E-142 | 1.28E-164 | 6.91E-207 |
| ARS3010 | 9.30E-05 | 5.81E-05 | 7.04E-01 | 8.76E-01 | 2.54E-10 | 1.28E-01 | 6.66E-08 | 1.42E-08 | 2.96E-07 | 1.00E+00 | 3.56E-95 | 3.23E-75 | 4.60E-118 | 3.82E-217 | 8.52E-120 | 1.25E-130 | 4.13E-110 | 8.82E-112 | 7.26E-132 | 1.09E-181 |
| Meta_AMS3002 | 8.82E-112 | 7.09E-85 | 2.36E-100 | 3.68E-94 | 7.85E-86 | 9.90E-89 | 7.85E-86 | 9.55E-80 | 2.42E-107 | 3.56E-95 | 1.00E+00 | 3.30E-03 | 8.07E-03 | 3.73E-23 | 6.66E-08 | 4.72E-06 | 1.13E-02 | 4.73E-03 | 6.09E-06 | 1.22E-13 |
| Meta_AMS3003 | 3.64E-93 | 4.21E-62 | 1.62E-82 | 8.93E-75 | 6.49E-61 | 6.10E-65 | 6.49E-61 | 2.23E-54 | 9.34E-88 | 3.23E-75 | 3.30E-03 | 1.00E+00 | 2.67E-08 | 6.51E-32 | 3.64E-08 | 9.34E-12 | 2.23E-07 | 2.78E-06 | 6.42E-12 | 3.36E-20 |
| Meta_AMS3004 | 5.45E-146 | 1.15E-93 | 5.84E-127 | 2.50E-116 | 3.61E-92 | 1.52E-103 | 3.45E-90 | 3.95E-81 | 4.54E-142 | 4.60E-118 | 8.07E-03 | 2.67E-08 | 1.00E+00 | 1.22E-13 | 1.83E-04 | 1.53E-03 | 1.01E-01 | 1.81E-01 | 1.56E-02 | 3.95E-07 |
| Meta_AMS3005 | 9.73E-255 | 3.23E-180 | 9.74E-231 | 2.38E-216 | 2.07E-173 | 7.72E-197 | 3.57E-177 | 3.94E-156 | 5.43E-245 | 3.82E-217 | 3.73E-23 | 6.51E-32 | 1.22E-13 | 1.00E+00 | 1.18E-17 | 2.95E-16 | 2.71E-15 | 1.89E-16 | 1.98E-11 | 1.03E-08 |
| Meta_AMS3006 | 5.92E-150 | 1.11E-91 | 4.98E-130 | 1.25E-118 | 9.90E-89 | 1.05E-105 | 3.27E-89 | 5.19E-77 | 2.45E-145 | 8.52E-120 | 6.66E-08 | 3.64E-08 | 1.83E-04 | 1.18E-17 | 1.00E+00 | 5.81E-05 | 2.88E-02 | 1.80E-02 | 9.55E-03 | 2.96E-07 |
| Meta_AMS3007 | 1.62E-167 | 1.11E-95 | 5.06E-144 | 4.98E-130 | 3.27E-89 | 4.60E-118 | 3.61E-92 | 3.95E-81 | 6.42E-164 | 1.25E-130 | 4.72E-06 | 9.34E-12 | 1.53E-03 | 2.95E-16 | 5.81E-05 | 1.00E+00 | 6.83E-02 | 1.47E-04 | 1.14E-01 | 2.84E-09 |
| Meta_AMS3008 | 4.04E-140 | 2.65E-86 | 3.79E-122 | 4.13E-110 | 1.62E-82 | 8.80E-99 | 1.89E-83 | 3.90E-72 | 1.28E-135 | 4.13E-110 | 1.13E-02 | 2.23E-07 | 1.01E-01 | 2.71E-15 | 2.88E-02 | 6.83E-02 | 1.00E+00 | 1.80E-02 | 1.28E-01 | 1.03E-08 |
| Meta_AMS3009 | 5.45E-146 | 4.74E-82 | 9.44E-126 | 1.82E-113 | 2.26E-78 | 7.08E-101 | 1.38E-81 | 2.84E-68 | 4.54E-142 | 8.82E-112 | 4.73E-03 | 2.78E-06 | 1.81E-01 | 1.89E-16 | 1.80E-02 | 1.47E-04 | 1.80E-02 | 1.00E+00 | 6.83E-02 | 7.26E-10 |
| Meta_AMS3010 | 5.81E-169 | 2.89E-98 | 1.29E-148 | 5.37E-135 | 1.11E-91 | 5.73E-121 | 1.15E-93 | 1.14E-80 | 1.28E-164 | 7.26E-132 | 6.09E-06 | 6.42E-12 | 1.56E-02 | 1.98E-11 | 9.55E-03 | 1.14E-01 | 1.28E-01 | 6.83E-02 | 1.00E+00 | 2.13E-06 |
| Meta_AMS3011 | 3.81E-211 | 1.12E-152 | 1.48E-189 | 3.23E-180 | 5.65E-148 | 8.83E-162 | 5.65E-148 | 1.76E-132 | 6.91E-207 | 1.09E-181 | 1.22E-13 | 3.36E-20 | 3.95E-07 | 1.03E-08 | 2.96E-07 | 2.84E-09 | 1.03E-08 | 7.26E-10 | 2.13E-06 | 1.00E+00 |

Comparison of trees using the Williams & Clifford, 1971 method

| | ARS3001 | ARS3002 | ARS3003 | ARS3004 | ARS3005 | ARS3006 | ARS3007 | ARS3008 | ARS3009 | ARS3010 | Meta_AMS3002 | Meta_AMS3003 | Meta_AMS3004 | Meta_AMS3005 | Meta_AMS3006 | Meta_AMS3007 | Meta_AMS3008 | Meta_AMS3009 | Meta_AMS3010 | Meta_AMS3011 |
|---|---|---|---|---|---|---|---|---|---|---|---|---|---|---|---|---|---|---|---|---|
| ARS3001 | 1.00E+00 | 9.48E-08 | 7.38E-10 | 2.38E-05 | 2.02E-08 | 1.47E-08 | 4.84E-04 | 1.46E-09 | 7.02E-08 | 1.83E-10 | 4.12E-73 | 3.05E-54 | 9.28E-94 | 7.37E-136 | 7.03E-62 | 9.30E-67 | 1.02E-60 | 9.28E-94 | 5.34E-102 | 9.01E-124 |
| ARS3002 | 9.48E-08 | 1.00E+00 | 9.47E-01 | 1.45E-01 | 8.37E-03 | 8.90E-02 | 7.87E-02 | 1.34E-03 | 2.16E-23 | 5.29E-01 | 1.11E-72 | 7.00E-54 | 2.87E-80 | 1.77E-121 | 1.02E-60 | 1.92E-46 | 1.02E-60 | 2.86E-88 | 1.49E-77 | 1.32E-114 |
| ARS3003 | 7.38E-10 | 9.47E-01 | 1.00E+00 | 2.49E-02 | 1.83E-02 | 1.45E-01 | 1.37E-02 | 6.01E-03 | 7.92E-27 | 5.29E-01 | 1.50E-70 | 2.41E-51 | 1.49E-77 | 2.16E-117 | 2.17E-58 | 9.17E-45 | 1.25E-57 | 6.26E-85 | 5.51E-74 | 1.18E-110 |
| ARS3004 | 2.38E-05 | 1.45E-01 | 2.49E-02 | 1.00E+00 | 2.90E-02 | 2.14E-02 | 4.92E-01 | 1.16E-02 | 7.55E-17 | 1.16E-02 | 5.29E-68 | 6.48E-50 | 2.59E-75 | 1.32E-114 | 1.25E-57 | 1.90E-43 | 1.00E-55 | 1.42E-82 | 1.18E-76 | 1.78E-108 |
| ARS3005 | 2.02E-08 | 8.37E-03 | 1.83E-02 | 2.90E-02 | 1.00E+00 | 1.81E-01 | 9.91E-03 | 9.47E-01 | 3.18E-22 | 1.62E-01 | 1.06E-51 | 8.01E-42 | 3.78E-59 | 9.28E-94 | 8.01E-42 | 6.85E-34 | 9.08E-44 | 1.12E-62 | 1.74E-61 | 5.50E-84 |
| ARS3006 | 1.47E-08 | 8.90E-02 | 1.45E-01 | 2.14E-02 | 1.81E-01 | 1.00E+00 | 1.83E-02 | 1.15E-01 | 2.53E-26 | 2.02E-01 | 7.00E-54 | 4.28E-45 | 2.74E-64 | 1.45E-103 | 4.28E-45 | 3.45E-41 | 1.90E-43 | 2.86E-69 | 2.13E-71 | 1.00E-94 |
| ARS3007 | 4.84E-04 | 7.87E-02 | 1.37E-02 | 4.92E-01 | 9.91E-03 | 1.83E-02 | 1.00E+00 | 1.98E-03 | 2.95E-16 | 4.22E-03 | 1.50E-70 | 4.65E-52 | 1.97E-78 | 1.26E-122 | 4.28E-61 | 1.45E-49 | 2.17E-58 | 2.39E-86 | 2.87E-80 | 1.09E-115 |
| ARS3008 | 1.46E-09 | 1.34E-03 | 6.01E-03 | 1.16E-02 | 9.47E-01 | 1.15E-01 | 1.98E-03 | 1.00E+00 | 4.20E-24 | 8.90E-02 | 1.80E-47 | 3.45E-41 | 2.36E-55 | 2.63E-87 | 2.28E-38 | 2.55E-33 | 1.49E-40 | 9.14E-59 | 1.31E-54 | 1.04E-80 |
| ARS3009 | 7.02E-08 | 2.16E-23 | 7.92E-27 | 7.55E-17 | 3.18E-22 | 2.53E-26 | 2.95E-16 | 4.20E-24 | 1.00E+00 | 7.43E-28 | 8.25E-98 | 3.61E-67 | 1.91E-125 | 8.93E-174 | 2.14E-85 | 8.25E-98 | 8.15E-72 | 6.45E-125 | 2.42E-135 | 4.37E-160 |
| ARS3010 | 1.83E-10 | 5.29E-01 | 5.29E-01 | 1.16E-02 | 1.62E-01 | 2.02E-01 | 4.22E-03 | 8.90E-02 | 7.43E-28 | 1.00E+00 | 6.90E-64 | 1.80E-47 | 2.13E-71 | 1.78E-108 | 2.03E-52 | 6.33E-40 | 8.85E-53 | 5.50E-78 | 6.23E-66 | 1.64E-102 |
| Meta_AMS3002 | 4.12E-73 | 1.11E-72 | 1.50E-70 | 5.29E-68 | 1.06E-51 | 7.00E-54 | 1.50E-70 | 1.80E-47 | 8.25E-98 | 6.90E-64 | 1.00E+00 | 7.95E-05 | 7.10E-03 | 2.60E-10 | 1.29E-01 | 7.32E-04 | 7.10E-03 | 1.60E-02 | 1.10E-03 | 5.18E-08 |
| Meta_AMS3003 | 3.05E-54 | 7.00E-54 | 2.41E-51 | 6.48E-50 | 8.01E-42 | 4.28E-45 | 4.65E-52 | 3.45E-41 | 3.61E-67 | 1.80E-47 | 7.95E-05 | 1.00E+00 | 4.44E-12 | 7.25E-24 | 3.35E-02 | 9.76E-07 | 6.01E-03 | 7.38E-10 | 5.35E-14 | 1.21E-20 |
| Meta_AMS3004 | 9.28E-94 | 2.87E-80 | 1.49E-77 | 2.59E-75 | 3.78E-59 | 2.74E-64 | 1.97E-78 | 2.36E-55 | 1.91E-125 | 2.13E-71 | 7.10E-03 | 4.44E-12 | 1.00E+00 | 2.38E-05 | 3.89E-06 | 6.30E-05 | 4.01E-09 | 5.70E-01 | 1.83E-02 | 5.96E-04 |
| Meta_AMS3005 | 7.37E-136 | 1.77E-121 | 2.16E-117 | 1.32E-114 | 9.28E-94 | 1.45E-103 | 1.26E-122 | 2.63E-87 | 8.93E-174 | 1.78E-108 | 2.60E-10 | 7.25E-24 | 2.38E-05 | 1.00E+00 | 2.71E-15 | 3.01E-17 | 5.36E-22 | 3.10E-07 | 1.72E-07 | 1.63E-03 |
| Meta_AMS3006 | 7.03E-62 | 1.02E-60 | 2.17E-58 | 1.25E-57 | 8.01E-42 | 4.28E-45 | 4.28E-61 | 2.28E-38 | 2.14E-85 | 2.03E-52 | 1.29E-01 | 3.35E-02 | 3.89E-06 | 2.71E-15 | 1.00E+00 | 1.83E-02 | 3.42E-01 | 1.27E-04 | 1.72E-07 | 1.38E-12 |
| Meta_AMS3007 | 9.30E-67 | 1.92E-46 | 9.17E-45 | 1.90E-43 | 6.85E-34 | 3.45E-41 | 1.45E-49 | 2.55E-33 | 8.25E-98 | 6.33E-40 | 7.32E-04 | 9.76E-07 | 6.30E-05 | 3.01E-17 | 1.83E-02 | 1.00E+00 | 4.22E-03 | 7.38E-07 | 1.88E-05 | 2.32E-14 |
| Meta_AMS3008 | 1.02E-60 | 1.02E-60 | 1.25E-57 | 1.00E-55 | 9.08E-44 | 1.90E-43 | 2.17E-58 | 1.49E-40 | 8.15E-72 | 8.85E-53 | 7.10E-03 | 6.01E-03 | 4.01E-09 | 5.36E-22 | 3.42E-01 | 4.22E-03 | 1.00E+00 | 5.55E-09 | 2.95E-11 | 1.75E-18 |
| Meta_AMS3009 | 9.28E-94 | 2.86E-88 | 6.26E-85 | 1.42E-82 | 1.12E-62 | 2.86E-69 | 2.39E-86 | 9.14E-59 | 6.45E-125 | 5.50E-78 | 1.60E-02 | 7.38E-10 | 5.70E-01 | 3.10E-07 | 1.27E-04 | 7.38E-07 | 5.55E-09 | 1.00E+00 | 8.37E-03 | 1.83E-02 |
| Meta_AMS3010 | 5.34E-102 | 1.49E-77 | 5.51E-74 | 1.18E-76 | 1.74E-61 | 2.13E-71 | 2.87E-80 | 1.31E-54 | 2.42E-135 | 6.23E-66 | 1.10E-03 | 5.35E-14 | 1.83E-02 | 1.72E-07 | 1.72E-07 | 1.88E-05 | 2.95E-11 | 8.37E-03 | 1.00E+00 | 3.04E-05 |
| Meta_AMS3011 | 9.01E-124 | 1.32E-114 | 1.18E-110 | 1.78E-108 | 5.50E-84 | 1.00E-94 | 1.09E-115 | 1.04E-80 | 4.37E-160 | 1.64E-102 | 5.18E-08 | 1.21E-20 | 5.96E-04 | 1.63E-03 | 1.38E-12 | 2.32E-14 | 1.75E-18 | 1.83E-02 | 3.04E-05 | 1.00E+00 |

Comparison of trees using the Robinson & Foulds, 1981 method

| | ARS3001 | ARS3002 | ARS3003 | ARS3004 | ARS3005 | ARS3006 | ARS3007 | ARS3008 | ARS3009 | ARS3010 | Meta_AMS3002 | Meta_AMS3003 | Meta_AMS3004 | Meta_AMS3005 | Meta_AMS3006 | Meta_AMS3007 | Meta_AMS3008 | Meta_AMS3009 | Meta_AMS3010 | Meta_AMS3011 |
|---|---|---|---|---|---|---|---|---|---|---|---|---|---|---|---|---|---|---|---|---|
| ARS3001 | 1.00E+00 | 1.74E-05 | 2.86E-06 | 4.41E-02 | 1.65E-07 | 2.61E-08 | 4.05E-15 | 7.33E-25 | 1.98E-11 | 4.33E-04 | 1.23E-70 | 4.64E-54 | 3.41E-63 | 5.12E-107 | 6.91E-51 | 7.96E-50 | 1.20E-61 | 2.74E-52 | 2.01E-54 | 1.98E-74 |
| ARS3002 | 1.74E-05 | 1.00E+00 | 8.39E-01 | 3.32E-02 | 1.48E-03 | 1.84E-04 | 2.22E-07 | 1.82E-16 | 5.36E-30 | 7.70E-02 | 8.27E-70 | 2.28E-53 | 1.20E-61 | 5.06E-104 | 8.78E-49 | 9.30E-48 | 2.39E-59 | 1.40E-51 | 1.03E-53 | 7.45E-72 |
| ARS3003 | 2.86E-06 | 8.39E-01 | 1.00E+00 | 3.16E-03 | 8.75E-02 | 2.10E-02 | 9.30E-05 | 6.37E-12 | 6.65E-34 | 7.55E-01 | 8.27E-70 | 2.28E-53 | 1.20E-61 | 2.73E-99 | 2.25E-45 | 1.05E-44 | 8.45E-58 | 1.40E-51 | 1.03E-53 | 5.00E-71 |
| ARS3004 | 4.41E-02 | 3.32E-02 | 3.16E-03 | 1.00E+00 | 3.53E-04 | 1.21E-03 | 2.61E-08 | 1.09E-15 | 1.95E-20 | 1.44E-01 | 2.13E-69 | 5.24E-53 | 2.85E-61 | 7.68E-99 | 9.62E-44 | 2.18E-44 | 1.96E-57 | 3.11E-51 | 2.28E-53 | 1.23E-70 |
| ARS3005 | 1.65E-07 | 1.48E-03 | 8.75E-02 | 3.53E-04 | 1.00E+00 | 1.00E+00 | 3.83E-02 | 3.71E-06 | 3.90E-37 | 3.70E-01 | 8.39E-63 | 4.58E-44 | 1.40E-51 | 8.35E-86 | 2.34E-35 | 1.95E-37 | 4.35E-48 | 2.18E-44 | 2.18E-44 | 4.10E-60 |
| ARS3006 | 2.61E-08 | 1.84E-04 | 2.10E-02 | 1.21E-03 | 1.00E+00 | 1.00E+00 | 1.80E-01 | 9.30E-05 | 4.83E-38 | 2.49E-01 | 1.20E-61 | 9.62E-44 | 3.11E-51 | 2.27E-85 | 4.58E-35 | 1.53E-36 | 9.30E-48 | 2.04E-43 | 4.58E-44 | 9.91E-60 |
| ARS3007 | 4.05E-15 | 2.22E-07 | 9.30E-05 | 2.61E-08 | 3.83E-02 | 1.80E-01 | 1.00E+00 | 3.32E-02 | 2.74E-52 | 2.29E-04 | 6.76E-61 | 1.78E-41 | 1.75E-49 | 1.55E-83 | 4.78E-33 | 6.03E-36 | 2.25E-45 | 9.14E-43 | 4.06E-42 | 4.65E-57 |
| ARS3008 | 7.33E-25 | 1.82E-16 | 6.37E-12 | 1.09E-15 | 3.71E-06 | 9.30E-05 | 3.32E-02 | 1.00E+00 | 2.85E-61 | 2.75E-09 | 1.10E-56 | 7.74E-37 | 1.06E-45 | 1.89E-78 | 4.20E-25 | 9.12E-33 | 2.84E-39 | 2.84E-39 | 1.18E-38 | 6.91E-51 |
| ARS3009 | 1.98E-11 | 5.36E-30 | 6.65E-34 | 1.95E-20 | 3.90E-37 | 4.83E-38 | 2.74E-52 | 2.85E-61 | 1.00E+00 | 4.19E-27 | 1.19E-76 | 6.76E-61 | 3.79E-82 | 2.88E-139 | 4.01E-73 | 1.93E-71 | 6.20E-85 | 8.10E-65 | 7.45E-72 | 1.22E-106 |
| ARS3010 | 4.33E-04 | 7.70E-02 | 7.55E-01 | 1.44E-01 | 3.70E-01 | 2.49E-01 | 2.29E-04 | 2.75E-09 | 4.19E-27 | 1.00E+00 | 1.83E-66 | 1.75E-49 | 1.96E-57 | 1.02E-93 | 3.65E-41 | 3.65E-41 | 1.03E-53 | 9.30E-48 | 7.96E-50 | 1.83E-66 |
| Meta_AMS3002 | 1.23E-70 | 8.27E-70 | 8.27E-70 | 2.13E-69 | 8.39E-63 | 1.20E-61 | 6.76E-61 | 1.10E-56 | 1.19E-76 | 1.83E-66 | 1.00E+00 | 7.74E-03 | 5.85E-05 | 4.20E-25 | 1.40E-09 | 5.11E-10 | 1.66E-06 | 8.15E-06 | 1.93E-08 | 3.42E-14 |
| Meta_AMS3003 | 4.64E-54 | 2.28E-53 | 2.28E-53 | 5.24E-53 | 4.58E-44 | 9.62E-44 | 1.78E-41 | 7.74E-37 | 6.76E-61 | 1.75E-49 | 7.74E-03 | 1.00E+00 | 5.33E-07 | 6.28E-29 | 4.65E-05 | 6.06E-13 | 1.42E-08 | 9.06E-08 | 4.15E-11 | 7.13E-18 |
| Meta_AMS3004 | 3.41E-63 | 1.20E-61 | 1.20E-61 | 2.85E-61 | 1.40E-51 | 3.11E-51 | 1.75E-49 | 1.06E-45 | 3.79E-82 | 1.96E-57 | 5.85E-05 | 5.33E-07 | 1.00E+00 | 1.82E-17 | 1.35E-05 | 1.05E-05 | 1.44E-01 | 2.49E-01 | 1.61E-01 | 5.85E-05 |
| Meta_AMS3005 | 5.12E-107 | 5.06E-104 | 2.73E-99 | 7.68E-99 | 8.35E-86 | 2.27E-85 | 1.55E-83 | 1.89E-78 | 2.88E-139 | 1.02E-93 | 4.20E-25 | 6.28E-29 | 1.82E-17 | 1.00E+00 | 4.31E-26 | 2.54E-10 | 1.40E-09 | 6.07E-23 | 1.02E-09 | 1.40E-09 |
| Meta_AMS3006 | 6.91E-51 | 8.78E-49 | 2.25E-45 | 9.62E-44 | 2.34E-35 | 4.58E-35 | 4.78E-33 | 4.20E-25 | 4.01E-73 | 3.65E-41 | 1.40E-09 | 4.65E-05 | 1.35E-05 | 4.31E-26 | 1.00E+00 | 3.71E-06 | 1.17E-04 | 5.34E-04 | 2.22E-05 | 1.93E-08 |
| Meta_AMS3007 | 7.96E-50 | 9.30E-48 | 1.05E-44 | 2.18E-44 | 1.95E-37 | 1.53E-36 | 6.03E-36 | 9.12E-33 | 1.93E-71 | 3.65E-41 | 5.11E-10 | 6.06E-13 | 1.05E-05 | 2.54E-10 | 3.71E-06 | 1.00E+00 | 6.52E-03 | 7.43E-09 | 2.85E-04 | 1.21E-03 |
| Meta_AMS3008 | 1.20E-61 | 2.39E-59 | 8.45E-58 | 1.96E-57 | 4.35E-48 | 9.30E-48 | 2.25E-45 | 2.84E-39 | 6.20E-85 | 1.03E-53 | 1.66E-06 | 1.42E-08 | 1.44E-01 | 1.40E-09 | 1.17E-04 | 6.52E-03 | 1.00E+00 | 2.18E-03 | 3.06E-01 | 6.52E-03 |
| Meta_AMS3009 | 2.74E-52 | 1.40E-51 | 1.40E-51 | 3.11E-51 | 2.18E-44 | 2.04E-43 | 9.14E-43 | 2.84E-39 | 8.10E-65 | 9.30E-48 | 8.15E-06 | 9.06E-08 | 2.49E-01 | 6.07E-23 | 5.34E-04 | 7.43E-09 | 2.18E-03 | 1.00E+00 | 2.63E-03 | 8.15E-06 |
| Meta_AMS3010 | 2.01E-54 | 1.03E-53 | 1.03E-53 | 2.28E-53 | 2.18E-44 | 4.58E-44 | 4.06E-42 | 1.18E-38 | 7.45E-72 | 7.96E-50 | 1.93E-08 | 4.15E-11 | 1.61E-01 | 1.02E-09 | 2.22E-05 | 2.85E-04 | 3.06E-01 | 2.63E-03 | 1.00E+00 | 4.33E-04 |
| Meta_AMS3011 | 1.98E-74 | 7.45E-72 | 5.00E-71 | 1.23E-70 | 4.10E-60 | 9.91E-60 | 4.65E-57 | 6.91E-51 | 1.22E-106 | 1.83E-66 | 3.42E-14 | 7.13E-18 | 5.85E-05 | 1.40E-09 | 1.93E-08 | 1.21E-03 | 6.52E-03 | 8.15E-06 | 4.33E-04 | 1.00E+00 |

Comparison of trees using the Kuhner & Felsenstein, 1994 method

| | ARS3001 | ARS3002 | ARS3003 | ARS3004 | ARS3005 | ARS3006 | ARS3007 | ARS3008 | ARS3009 | ARS3010 | Meta_AMS3002 | Meta_AMS3003 | Meta_AMS3004 | Meta_AMS3005 | Meta_AMS3006 | Meta_AMS3007 | Meta_AMS3008 | Meta_AMS3009 | Meta_AMS3010 | Meta_AMS3011 |
|---|---|---|---|---|---|---|---|---|---|---|---|---|---|---|---|---|---|---|---|---|
| ARS3001 | 1.00E+00 | 2.26E-03 | 3.06E-09 | 3.06E-09 | 3.89E-05 | 7.70E-02 | 8.75E-02 | 2.49E-01 | 2.46E-04 | 1.12E-09 | 4.67E-23 | 1.65E-16 | 7.90E-23 | 1.52E-11 | 1.04E-18 | 6.56E-19 | 1.06E-16 | 1.07E-24 | 2.28E-12 | 3.92E-16 |
| ARS3002 | 2.26E-03 | 1.00E+00 | 3.26E-03 | 3.26E-03 | 4.62E-04 | 3.26E-03 | 2.34E-06 | 2.90E-02 | 4.59E-11 | 5.07E-06 | 2.50E-27 | 6.54E-25 | 3.50E-31 | 1.08E-17 | 3.25E-24 | 8.03E-28 | 9.80E-14 | 1.20E-30 | 4.81E-13 | 2.13E-25 |
| ARS3003 | 3.06E-09 | 3.26E-03 | 1.00E+00 | 6.62E-01 | 2.90E-02 | 2.45E-07 | 1.44E-15 | 3.03E-06 | 2.77E-21 | 7.70E-02 | 6.34E-38 | 1.96E-32 | 3.28E-39 | 1.23E-25 | 1.07E-31 | 1.13E-36 | 1.08E-17 | 8.70E-38 | 2.77E-21 | 6.23E-36 |
| ARS3004 | 3.06E-09 | 3.26E-03 | 6.62E-01 | 1.00E+00 | 4.51E-02 | 3.27E-07 | 1.44E-15 | 3.92E-06 | 6.19E-22 | 6.83E-02 | 3.48E-33 | 2.44E-29 | 3.71E-36 | 1.75E-21 | 7.78E-29 | 1.96E-32 | 1.44E-15 | 2.15E-35 | 7.01E-18 | 1.07E-31 |
| ARS3005 | 3.89E-05 | 4.62E-04 | 2.90E-02 | 4.51E-02 | 1.00E+00 | 2.13E-08 | 4.00E-08 | 5.74E-07 | 1.84E-14 | 2.15E-02 | 4.27E-27 | 1.23E-25 | 3.28E-32 | 2.72E-17 | 2.50E-27 | 1.39E-28 | 9.51E-11 | 6.16E-33 | 3.24E-13 | 1.23E-25 |
| ARS3006 | 7.70E-02 | 3.26E-03 | 2.45E-07 | 3.27E-07 | 2.13E-08 | 1.00E+00 | 3.06E-04 | 7.01E-01 | 5.07E-06 | 2.17E-13 | 4.15E-26 | 2.75E-23 | 1.37E-29 | 1.08E-17 | 1.69E-23 | 4.15E-26 | 2.77E-21 | 4.36E-29 | 3.92E-16 | 2.75E-23 |
| ARS3007 | 8.75E-02 | 2.34E-06 | 1.44E-15 | 1.44E-15 | 4.00E-08 | 3.06E-04 | 1.00E+00 | 1.05E-03 | 8.02E-03 | 8.02E-15 | 1.07E-24 | 1.55E-12 | 6.56E-19 | 1.05E-12 | 1.84E-14 | 2.21E-15 | 4.28E-17 | 2.10E-20 | 1.05E-12 | 7.20E-12 |
| ARS3008 | 2.49E-01 | 2.90E-02 | 3.03E-06 | 3.92E-06 | 5.74E-07 | 7.01E-01 | 1.05E-03 | 1.00E+00 | 3.03E-06 | 3.98E-10 | 3.77E-22 | 1.04E-18 | 1.07E-24 | 4.40E-18 | 2.10E-20 | 6.19E-22 | 9.20E-20 | 4.15E-26 | 8.02E-15 | 9.20E-20 |
| ARS3009 | 2.46E-04 | 4.59E-11 | 2.77E-21 | 6.19E-22 | 1.84E-14 | 5.07E-06 | 8.02E-03 | 3.03E-06 | 1.00E+00 | 3.77E-22 | 2.77E-21 | 6.08E-16 | 2.72E-17 | 2.75E-23 | 2.17E-13 | 1.84E-14 | 2.29E-22 | 3.44E-20 | 2.55E-16 | 2.55E-16 |
| ARS3010 | 1.12E-09 | 5.07E-06 | 7.70E-02 | 6.83E-02 | 2.15E-02 | 2.17E-13 | 8.02E-15 | 3.98E-10 | 3.77E-22 | 1.00E+00 | 3.28E-32 | 4.54E-28 | 2.93E-34 | 4.60E-21 | 4.27E-27 | 3.50E-31 | 1.47E-13 | 1.01E-33 | 1.06E-16 | 1.37E-29 |
| Meta_AMS3002 | 4.67E-23 | 2.50E-27 | 6.34E-38 | 3.48E-33 | 4.27E-27 | 4.15E-26 | 1.07E-24 | 3.77E-22 | 2.77E-21 | 3.28E-32 | 1.00E+00 | 2.46E-04 | 4.35E-07 | 1.01E-06 | 4.59E-11 | 1.00E-07 | 6.21E-05 | 1.05E-11 | 3.92E-06 | 7.20E-12 |
| Meta_AMS3003 | 1.65E-16 | 6.54E-25 | 1.96E-32 | 2.44E-29 | 1.23E-25 | 2.75E-23 | 1.55E-12 | 1.04E-18 | 6.08E-16 | 4.54E-28 | 2.46E-04 | 1.00E+00 | 9.86E-02 | 1.11E-01 | 4.62E-04 | 9.86E-02 | 3.76E-04 | 1.05E-03 | 6.76E-03 | 1.28E-03 |
| Meta_AMS3004 | 7.90E-23 | 3.50E-31 | 3.28E-39 | 3.71E-36 | 3.28E-32 | 1.37E-29 | 6.56E-19 | 1.07E-24 | 2.72E-17 | 2.93E-34 | 4.35E-07 | 9.86E-02 | 1.00E+00 | 3.76E-04 | 2.26E-03 | 7.01E-01 | 1.56E-08 | 1.28E-03 | 2.45E-07 | 1.84E-02 |
| Meta_AMS3005 | 1.52E-11 | 1.08E-17 | 1.23E-25 | 1.75E-21 | 2.72E-17 | 1.08E-17 | 1.05E-12 | 4.40E-18 | 2.75E-23 | 4.60E-21 | 1.01E-06 | 1.11E-01 | 3.76E-04 | 1.00E+00 | 5.74E-07 | 2.26E-03 | 7.70E-02 | 1.00E-07 | 2.49E-01 | 2.51E-02 |
| Meta_AMS3006 | 1.04E-18 | 3.25E-24 | 1.07E-31 | 7.78E-29 | 2.50E-27 | 1.69E-23 | 1.84E-14 | 2.10E-20 | 2.17E-13 | 4.27E-27 | 4.59E-11 | 4.62E-04 | 2.26E-03 | 5.74E-07 | 1.00E+00 | 1.58E-02 | 1.14E-08 | 3.06E-01 | 1.43E-05 | 1.55E-03 |
| Meta_AMS3007 | 6.56E-19 | 8.03E-28 | 1.13E-36 | 1.96E-32 | 1.39E-28 | 4.15E-26 | 2.21E-15 | 6.19E-22 | 1.84E-14 | 3.50E-31 | 1.00E-07 | 9.86E-02 | 7.01E-01 | 2.26E-03 | 1.58E-02 | 1.00E+00 | 8.56E-06 | 5.98E-02 | 4.91E-05 | 2.26E-01 |
| Meta_AMS3008 | 1.06E-16 | 9.80E-14 | 1.08E-17 | 1.44E-15 | 9.51E-11 | 2.77E-21 | 4.28E-17 | 9.20E-20 | 2.29E-22 | 1.47E-13 | 6.21E-05 | 3.76E-04 | 1.56E-08 | 7.70E-02 | 1.14E-08 | 8.56E-06 | 1.00E+00 | 7.96E-10 | 4.07E-01 | 4.91E-05 |
| Meta_AMS3009 | 1.07E-24 | 1.20E-30 | 8.70E-38 | 2.15E-35 | 6.16E-33 | 4.36E-29 | 2.10E-20 | 4.15E-26 | 3.44E-20 | 1.01E-33 | 1.05E-11 | 1.05E-03 | 1.28E-03 | 1.00E-07 | 3.06E-01 | 5.98E-02 | 7.96E-10 | 1.00E+00 | 8.25E-09 | 1.99E-04 |
| Meta_AMS3010 | 2.28E-12 | 4.81E-13 | 2.77E-21 | 7.01E-18 | 3.24E-13 | 3.92E-16 | 1.05E-12 | 8.02E-15 | 2.55E-16 | 1.06E-16 | 3.92E-06 | 6.76E-03 | 2.45E-07 | 2.49E-01 | 1.43E-05 | 4.91E-05 | 4.07E-01 | 8.25E-09 | 1.00E+00 | 1.05E-03 |
| Meta_AMS3011 | 3.92E-16 | 2.13E-25 | 6.23E-36 | 1.07E-31 | 1.23E-25 | 2.75E-23 | 7.20E-12 | 9.20E-20 | 2.55E-16 | 1.37E-29 | 7.20E-12 | 1.28E-03 | 1.84E-02 | 2.51E-02 | 1.55E-03 | 2.26E-01 | 4.91E-05 | 1.99E-04 | 1.05E-03 | 1.00E+00 |

**Supplementary Table S4B:** Pairwise FDR corrected p-values of the Kolmogorow-Smirnow test on the distances between phylogenetic trees for 23 ribosomal proteins (including SecY) in 10 samples of archaeal RefSeq genomes vs 10 samples of archaeal MAGs (associated with Supplementary Fig. S7b)

Comparison of trees using the Nye et al., 2006 method

| | ARS3001 | ARS3002 | ARS3003 | ARS3004 | ARS3005 | ARS3006 | ARS3007 | ARS3008 | ARS3009 | ARS3010 | Meta_AMS3002 | Meta_AMS3003 | Meta_AMS3004 | Meta_AMS3005 | Meta_AMS3006 | Meta_AMS3007 | Meta_AMS3008 | Meta_AMS3009 | Meta_AMS3010 | Meta_AMS3011 |
|---|---|---|---|---|---|---|---|---|---|---|---|---|---|---|---|---|---|---|---|---|
| ARS3001 | 1.00E+00 | 1.97E-05 | 7.46E-01 | 9.01E-03 | 6.75E-05 | 5.42E-02 | 2.41E-09 | 3.01E-05 | 3.33E-01 | 1.56E-02 | 8.31E-17 | 1.32E-06 | 1.54E-33 | 2.33E-79 | 4.02E-24 | 7.17E-50 | 7.34E-23 | 4.24E-20 | 5.48E-31 | 3.70E-64 |
| ARS3002 | 1.97E-05 | 1.00E+00 | 2.12E-04 | 6.72E-03 | 6.69E-01 | 1.56E-02 | 8.57E-02 | 9.45E-01 | 4.21E-09 | 8.57E-02 | 1.00E-04 | 4.51E-01 | 4.63E-12 | 1.64E-50 | 1.30E-08 | 2.65E-26 | 1.32E-06 | 6.47E-08 | 1.69E-11 | 3.34E-37 |
| ARS3003 | 7.46E-01 | 2.12E-04 | 1.00E+00 | 2.01E-02 | 6.51E-04 | 2.37E-01 | 2.24E-08 | 3.10E-04 | 1.62E-01 | 5.42E-02 | 4.05E-15 | 4.54E-05 | 5.08E-30 | 7.32E-70 | 7.41E-21 | 1.30E-43 | 2.30E-19 | 1.81E-16 | 1.14E-27 | 8.44E-57 |
| ARS3004 | 9.01E-03 | 6.72E-03 | 2.01E-02 | 1.00E+00 | 1.56E-02 | 6.69E-01 | 1.47E-04 | 4.33E-02 | 1.00E-04 | 6.69E-01 | 1.09E-07 | 2.63E-02 | 2.30E-19 | 1.17E-62 | 7.71E-14 | 4.16E-35 | 2.39E-12 | 6.24E-13 | 1.62E-17 | 5.71E-48 |
| ARS3005 | 6.75E-05 | 6.69E-01 | 6.51E-04 | 1.56E-02 | 1.00E+00 | 4.33E-02 | 1.56E-02 | 4.51E-01 | 6.47E-08 | 3.33E-01 | 1.47E-04 | 2.84E-01 | 3.98E-16 | 5.99E-54 | 1.14E-10 | 1.52E-29 | 4.21E-09 | 4.21E-09 | 3.79E-14 | 1.24E-40 |
| ARS3006 | 5.42E-02 | 1.56E-02 | 2.37E-01 | 6.69E-01 | 4.33E-02 | 1.00E+00 | 6.75E-05 | 5.42E-02 | 3.58E-03 | 8.20E-01 | 1.34E-09 | 1.56E-02 | 1.81E-20 | 1.62E-60 | 1.82E-14 | 4.16E-35 | 1.53E-13 | 4.63E-12 | 1.00E-19 | 4.44E-46 |
| ARS3007 | 2.41E-09 | 8.57E-02 | 2.24E-08 | 1.47E-04 | 1.56E-02 | 6.75E-05 | 1.00E+00 | 8.57E-02 | 1.53E-13 | 1.47E-04 | 1.07E-01 | 1.32E-01 | 3.82E-08 | 7.79E-45 | 2.12E-04 | 1.00E-19 | 6.75E-05 | 1.97E-05 | 2.10E-06 | 1.37E-28 |
| ARS3008 | 3.01E-05 | 9.45E-01 | 3.10E-04 | 4.33E-02 | 4.51E-01 | 5.42E-02 | 8.57E-02 | 1.00E+00 | 1.81E-07 | 3.39E-02 | 2.12E-04 | 9.45E-01 | 1.53E-13 | 1.07E-46 | 2.24E-08 | 4.02E-24 | 1.09E-07 | 3.30E-06 | 8.88E-12 | 1.54E-33 |
| ARS3009 | 3.33E-01 | 4.21E-09 | 1.62E-01 | 1.00E-04 | 6.47E-08 | 3.58E-03 | 1.53E-13 | 1.81E-07 | 1.00E+00 | 1.97E-05 | 4.24E-20 | 1.30E-08 | 1.11E-36 | 7.82E-77 | 1.14E-27 | 5.71E-48 | 1.54E-24 | 4.24E-20 | 4.16E-35 | 3.11E-61 |
| ARS3010 | 1.56E-02 | 8.57E-02 | 5.42E-02 | 6.69E-01 | 3.33E-01 | 8.20E-01 | 1.47E-04 | 3.39E-02 | 1.97E-05 | 1.00E+00 | 3.82E-08 | 3.39E-02 | 2.30E-19 | 6.01E-62 | 7.71E-14 | 1.34E-35 | 1.23E-12 | 1.23E-12 | 3.05E-18 | 5.71E-48 |
| Meta_AMS3002 | 8.31E-17 | 1.00E-04 | 4.05E-15 | 1.09E-07 | 1.47E-04 | 1.34E-09 | 1.07E-01 | 2.12E-04 | 4.24E-20 | 3.82E-08 | 1.00E+00 | 2.12E-04 | 1.30E-05 | 1.11E-36 | 1.32E-03 | 1.81E-16 | 1.20E-02 | 3.58E-03 | 4.54E-05 | 1.54E-24 |
| Meta_AMS3003 | 1.32E-06 | 4.51E-01 | 4.54E-05 | 2.63E-02 | 2.84E-01 | 1.56E-02 | 1.32E-01 | 9.45E-01 | 1.30E-08 | 3.39E-02 | 2.12E-04 | 1.00E+00 | 8.88E-12 | 2.08E-42 | 1.81E-07 | 4.78E-22 | 3.30E-06 | 5.24E-06 | 3.19E-11 | 1.68E-30 |
| Meta_AMS3004 | 1.54E-33 | 4.63E-12 | 5.08E-30 | 2.30E-19 | 3.98E-16 | 1.81E-20 | 3.82E-08 | 1.53E-13 | 1.11E-36 | 2.30E-19 | 1.30E-05 | 8.88E-12 | 1.00E+00 | 1.20E-21 | 5.42E-02 | 3.01E-05 | 2.01E-02 | 9.30E-04 | 3.33E-01 | 3.19E-11 |
| Meta_AMS3005 | 2.33E-79 | 1.64E-50 | 7.32E-70 | 1.17E-62 | 5.99E-54 | 1.62E-60 | 7.79E-45 | 1.07E-46 | 7.82E-77 | 6.01E-62 | 1.11E-36 | 2.08E-42 | 1.20E-21 | 1.00E+00 | 1.06E-23 | 3.01E-07 | 4.78E-22 | 7.41E-21 | 2.65E-26 | 2.01E-02 |
| Meta_AMS3006 | 4.02E-24 | 1.30E-08 | 7.41E-21 | 7.71E-14 | 1.14E-10 | 1.82E-14 | 2.12E-04 | 2.24E-08 | 1.14E-27 | 7.71E-14 | 1.32E-03 | 1.81E-07 | 5.42E-02 | 1.06E-23 | 1.00E+00 | 3.30E-06 | 1.00E+00 | 4.51E-01 | 3.92E-01 | 6.24E-13 |
| Meta_AMS3007 | 7.17E-50 | 2.65E-26 | 1.30E-43 | 4.16E-35 | 1.52E-29 | 4.16E-35 | 1.00E-19 | 4.02E-24 | 5.71E-48 | 1.34E-35 | 1.81E-16 | 4.78E-22 | 3.01E-05 | 3.01E-07 | 3.30E-06 | 1.00E+00 | 8.24E-07 | 1.32E-06 | 2.10E-06 | 2.01E-02 |
| Meta_AMS3008 | 7.34E-23 | 1.32E-06 | 2.30E-19 | 2.39E-12 | 4.21E-09 | 1.53E-13 | 6.75E-05 | 1.09E-07 | 1.54E-24 | 1.23E-12 | 1.20E-02 | 3.30E-06 | 2.01E-02 | 4.78E-22 | 1.00E+00 | 8.24E-07 | 1.00E+00 | 6.69E-01 | 2.37E-01 | 2.39E-12 |
| Meta_AMS3009 | 4.24E-20 | 6.47E-08 | 1.81E-16 | 6.24E-13 | 4.21E-09 | 4.63E-12 | 1.97E-05 | 3.30E-06 | 4.24E-20 | 1.23E-12 | 3.58E-03 | 5.24E-06 | 9.30E-04 | 7.41E-21 | 4.51E-01 | 1.32E-06 | 6.69E-01 | 1.00E+00 | 9.01E-03 | 1.69E-11 |
| Meta_AMS3010 | 5.48E-31 | 1.69E-11 | 1.14E-27 | 1.62E-17 | 3.79E-14 | 1.00E-19 | 2.10E-06 | 8.88E-12 | 4.16E-35 | 3.05E-18 | 4.54E-05 | 3.19E-11 | 3.33E-01 | 2.65E-26 | 3.92E-01 | 2.10E-06 | 2.37E-01 | 9.01E-03 | 1.00E+00 | 1.89E-15 |
| Meta_AMS3011 | 3.70E-64 | 3.34E-37 | 8.44E-57 | 5.71E-48 | 1.24E-40 | 4.44E-46 | 1.37E-28 | 1.54E-33 | 3.11E-61 | 5.71E-48 | 1.54E-24 | 1.68E-30 | 3.19E-11 | 2.01E-02 | 6.24E-13 | 2.01E-02 | 2.39E-12 | 1.69E-11 | 1.89E-15 | 1.00E+00 |

Comparison of trees using the Williams & Clifford, 1971 method

| | ARS3001 | ARS3002 | ARS3003 | ARS3004 | ARS3005 | ARS3006 | ARS3007 | ARS3008 | ARS3009 | ARS3010 | Meta_AMS3002 | Meta_AMS3003 | Meta_AMS3004 | Meta_AMS3005 | Meta_AMS3006 | Meta_AMS3007 | Meta_AMS3008 | Meta_AMS3009 | Meta_AMS3010 | Meta_AMS3011 |
|---|---|---|---|---|---|---|---|---|---|---|---|---|---|---|---|---|---|---|---|---|
| ARS3001 | 1.00E+00 | 1.16E-03 | 5.78E-04 | 1.09E-02 | 1.91E-02 | 4.11E-02 | 2.79E-01 | 1.09E-02 | 1.86E-04 | 1.64E-03 | 1.79E-08 | 4.11E-02 | 2.07E-31 | 4.17E-41 | 1.45E-02 | 1.21E-15 | 6.04E-03 | 1.21E-15 | 2.60E-15 | 1.47E-36 |
| ARS3002 | 1.16E-03 | 1.00E+00 | 7.88E-01 | 3.23E-01 | 9.07E-01 | 2.79E-01 | 4.11E-02 | 9.94E-01 | 2.59E-11 | 9.94E-01 | 6.73E-02 | 1.99E-01 | 3.48E-19 | 7.30E-27 | 4.92E-01 | 7.85E-08 | 8.42E-01 | 5.85E-10 | 7.85E-08 | 2.98E-22 |
| ARS3003 | 5.78E-04 | 7.88E-01 | 1.00E+00 | 3.23E-01 | 7.88E-01 | 3.23E-01 | 3.28E-02 | 4.28E-01 | 3.68E-12 | 9.94E-01 | 4.11E-02 | 1.65E-01 | 3.48E-19 | 3.03E-27 | 8.42E-01 | 7.85E-08 | 7.23E-01 | 2.85E-08 | 2.85E-08 | 4.74E-21 |
| ARS3004 | 1.09E-02 | 3.23E-01 | 3.23E-01 | 1.00E+00 | 9.07E-01 | 2.79E-01 | 1.99E-01 | 4.28E-01 | 1.33E-07 | 2.40E-01 | 8.20E-03 | 8.42E-01 | 1.04E-17 | 1.09E-24 | 9.65E-01 | 2.85E-08 | 5.65E-01 | 7.85E-08 | 2.20E-07 | 6.73E-20 |
| ARS3005 | 1.91E-02 | 9.07E-01 | 7.88E-01 | 9.07E-01 | 1.00E+00 | 2.79E-01 | 3.73E-01 | 7.88E-01 | 6.09E-09 | 7.23E-01 | 3.28E-02 | 4.92E-01 | 1.94E-18 | 4.13E-25 | 1.00E+00 | 3.54E-07 | 9.94E-01 | 2.85E-08 | 9.47E-07 | 1.93E-21 |
| ARS3006 | 4.11E-02 | 2.79E-01 | 3.23E-01 | 2.79E-01 | 2.79E-01 | 1.00E+00 | 4.28E-01 | 6.45E-01 | 1.79E-08 | 1.34E-01 | 1.27E-04 | 2.79E-01 | 2.53E-24 | 6.59E-33 | 3.23E-01 | 4.84E-11 | 3.73E-01 | 2.32E-13 | 4.84E-11 | 4.78E-28 |
| ARS3007 | 2.79E-01 | 4.11E-02 | 3.28E-02 | 1.99E-01 | 3.73E-01 | 4.28E-01 | 1.00E+00 | 2.40E-01 | 1.52E-06 | 5.28E-02 | 2.71E-04 | 3.73E-01 | 1.54E-25 | 1.94E-34 | 3.23E-01 | 7.16E-12 | 2.79E-01 | 3.68E-12 | 1.38E-11 | 4.78E-28 |
| ARS3008 | 1.09E-02 | 9.94E-01 | 4.28E-01 | 4.28E-01 | 7.88E-01 | 6.45E-01 | 2.40E-01 | 1.00E+00 | 3.19E-10 | 8.42E-01 | 1.09E-02 | 3.73E-01 | 1.51E-19 | 7.30E-27 | 4.28E-01 | 2.85E-08 | 7.88E-01 | 9.10E-11 | 4.84E-08 | 1.18E-22 |
| ARS3009 | 1.86E-04 | 2.59E-11 | 3.68E-12 | 1.33E-07 | 6.09E-09 | 1.79E-08 | 1.52E-06 | 3.19E-10 | 1.00E+00 | 9.49E-13 | 5.28E-17 | 3.87E-06 | 8.25E-43 | 5.99E-53 | 1.07E-08 | 1.19E-27 | 3.49E-09 | 2.53E-24 | 5.74E-26 | 4.30E-49 |
| ARS3010 | 1.64E-03 | 9.94E-01 | 9.94E-01 | 2.40E-01 | 7.23E-01 | 1.34E-01 | 5.28E-02 | 8.42E-01 | 9.49E-13 | 1.00E+00 | 1.99E-01 | 4.11E-02 | 2.64E-16 | 2.53E-24 | 3.73E-01 | 3.54E-07 | 4.28E-01 | 6.09E-09 | 9.47E-07 | 2.80E-20 |
| Meta_AMS3002 | 1.79E-08 | 6.73E-02 | 4.11E-02 | 8.20E-03 | 3.28E-02 | 1.27E-04 | 2.71E-04 | 1.09E-02 | 5.28E-17 | 1.99E-01 | 1.00E+00 | 2.30E-03 | 2.59E-11 | 2.36E-17 | 1.91E-02 | 1.16E-03 | 3.17E-03 | 1.52E-06 | 1.27E-04 | 1.21E-15 |
| Meta_AMS3003 | 4.11E-02 | 1.99E-01 | 1.65E-01 | 8.42E-01 | 4.92E-01 | 2.79E-01 | 3.73E-01 | 3.73E-01 | 3.87E-06 | 4.11E-02 | 2.30E-03 | 1.00E+00 | 1.51E-19 | 3.03E-27 | 7.88E-01 | 3.49E-09 | 3.23E-01 | 1.79E-08 | 2.85E-08 | 1.18E-22 |
| Meta_AMS3004 | 2.07E-31 | 3.48E-19 | 3.48E-19 | 1.04E-17 | 1.94E-18 | 2.53E-24 | 1.54E-25 | 1.51E-19 | 8.25E-43 | 2.64E-16 | 2.59E-11 | 1.51E-19 | 1.00E+00 | 2.71E-04 | 1.51E-19 | 8.31E-04 | 1.16E-20 | 3.17E-03 | 1.16E-03 | 3.66E-05 |
| Meta_AMS3005 | 4.17E-41 | 7.30E-27 | 3.03E-27 | 1.09E-24 | 4.13E-25 | 6.59E-33 | 1.94E-34 | 7.30E-27 | 5.99E-53 | 2.53E-24 | 2.36E-17 | 3.03E-27 | 2.71E-04 | 1.00E+00 | 7.30E-27 | 3.54E-07 | 1.19E-27 | 2.45E-06 | 9.47E-07 | 5.65E-01 |
| Meta_AMS3006 | 1.45E-02 | 4.92E-01 | 8.42E-01 | 9.65E-01 | 1.00E+00 | 3.23E-01 | 3.23E-01 | 4.28E-01 | 1.07E-08 | 3.73E-01 | 1.91E-02 | 7.88E-01 | 1.51E-19 | 7.30E-27 | 1.00E+00 | 3.54E-07 | 8.42E-01 | 4.84E-08 | 2.20E-07 | 4.80E-23 |
| Meta_AMS3007 | 1.21E-15 | 7.85E-08 | 7.85E-08 | 2.85E-08 | 3.54E-07 | 4.84E-11 | 7.16E-12 | 2.85E-08 | 1.19E-27 | 3.54E-07 | 1.16E-03 | 3.49E-09 | 8.31E-04 | 3.54E-07 | 3.54E-07 | 1.00E+00 | 7.85E-08 | 4.92E-01 | 8.42E-01 | 3.66E-05 |
| Meta_AMS3008 | 6.04E-03 | 8.42E-01 | 7.23E-01 | 5.65E-01 | 9.94E-01 | 3.73E-01 | 2.79E-01 | 7.88E-01 | 3.49E-09 | 4.28E-01 | 3.17E-03 | 3.23E-01 | 1.16E-20 | 1.19E-27 | 8.42E-01 | 7.85E-08 | 1.00E+00 | 1.79E-08 | 2.85E-08 | 2.53E-24 |
| Meta_AMS3009 | 1.21E-15 | 5.85E-10 | 2.85E-08 | 7.85E-08 | 2.85E-08 | 2.32E-13 | 3.68E-12 | 9.10E-11 | 2.53E-24 | 6.09E-09 | 1.52E-06 | 1.79E-08 | 3.17E-03 | 2.45E-06 | 4.84E-08 | 4.92E-01 | 1.79E-08 | 1.00E+00 | 1.07E-01 | 1.86E-04 |
| Meta_AMS3010 | 2.60E-15 | 7.85E-08 | 2.85E-08 | 2.20E-07 | 9.47E-07 | 4.84E-11 | 1.38E-11 | 4.84E-08 | 5.74E-26 | 9.47E-07 | 1.27E-04 | 2.85E-08 | 1.16E-03 | 9.47E-07 | 2.20E-07 | 8.42E-01 | 2.85E-08 | 1.07E-01 | 1.00E+00 | 3.87E-06 |
| Meta_AMS3011 | 1.47E-36 | 2.98E-22 | 4.74E-21 | 6.73E-20 | 1.93E-21 | 4.78E-28 | 4.78E-28 | 1.18E-22 | 4.30E-49 | 2.80E-20 | 1.21E-15 | 1.18E-22 | 3.66E-05 | 5.65E-01 | 4.80E-23 | 3.66E-05 | 2.53E-24 | 1.86E-04 | 3.87E-06 | 1.00E+00 |

Comparison of trees using the Robinson & Foulds, 1981 method

| | ARS3001 | ARS3002 | ARS3003 | ARS3004 | ARS3005 | ARS3006 | ARS3007 | ARS3008 | ARS3009 | ARS3010 | Meta_AMS3002 | Meta_AMS3003 | Meta_AMS3004 | Meta_AMS3005 | Meta_AMS3006 | Meta_AMS3007 | Meta_AMS3008 | Meta_AMS3009 | Meta_AMS3010 | Meta_AMS3011 |
|---|---|---|---|---|---|---|---|---|---|---|---|---|---|---|---|---|---|---|---|---|
| ARS3001 | 1.00E+00 | 6.19E-01 | 6.19E-01 | 8.52E-01 | 2.52E-01 | 9.66E-02 | 2.54E-05 | 6.89E-07 | 3.85E-05 | 2.95E-02 | 7.56E-17 | 3.70E-20 | 2.95E-02 | 3.66E-09 | 2.59E-07 | 1.76E-01 | 4.10E-01 | 1.63E-03 | 3.74E-02 | 2.32E-02 |
| ARS3002 | 6.19E-01 | 1.00E+00 | 1.00E+00 | 3.74E-02 | 1.00E+00 | 2.98E-01 | 1.63E-03 | 1.76E-06 | 6.38E-09 | 2.52E-01 | 6.56E-23 | 1.03E-26 | 1.92E-04 | 2.59E-07 | 8.08E-12 | 7.79E-01 | 2.32E-02 | 2.73E-06 | 5.73E-04 | 2.11E-01 |
| ARS3003 | 6.19E-01 | 1.00E+00 | 1.00E+00 | 3.74E-02 | 1.00E+00 | 6.19E-01 | 1.63E-03 | 3.85E-05 | 6.38E-09 | 2.98E-01 | 2.59E-23 | 4.71E-27 | 1.92E-04 | 1.09E-06 | 8.08E-12 | 8.52E-01 | 7.68E-03 | 1.09E-06 | 5.73E-04 | 4.10E-01 |
| ARS3004 | 8.52E-01 | 3.74E-02 | 3.74E-02 | 1.00E+00 | 7.68E-03 | 2.95E-02 | 2.73E-06 | 9.56E-08 | 4.03E-04 | 7.68E-03 | 2.18E-12 | 3.57E-15 | 9.66E-02 | 2.91E-11 | 2.54E-05 | 6.17E-02 | 7.79E-01 | 2.95E-02 | 1.19E-01 | 4.34E-03 |
| ARS3005 | 2.52E-01 | 1.00E+00 | 1.00E+00 | 7.68E-03 | 1.00E+00 | 2.98E-01 | 2.76E-04 | 2.73E-06 | 1.85E-10 | 1.19E-01 | 1.44E-24 | 5.98E-28 | 1.68E-05 | 9.56E-08 | 1.44E-13 | 4.75E-01 | 7.68E-03 | 6.89E-07 | 5.79E-05 | 2.11E-01 |
| ARS3006 | 9.66E-02 | 2.98E-01 | 6.19E-01 | 2.95E-02 | 2.98E-01 | 1.00E+00 | 1.76E-01 | 1.04E-02 | 5.39E-11 | 1.00E+00 | 1.65E-22 | 7.95E-26 | 6.89E-07 | 5.73E-04 | 5.55E-13 | 1.00E+00 | 1.36E-02 | 6.89E-06 | 1.09E-06 | 1.00E+00 |
| ARS3007 | 2.54E-05 | 1.63E-03 | 1.63E-03 | 2.73E-06 | 2.76E-04 | 1.76E-01 | 1.00E+00 | 4.10E-01 | 2.78E-18 | 1.19E-01 | 1.03E-26 | 1.33E-30 | 1.44E-13 | 1.36E-02 | 6.39E-21 | 9.66E-02 | 1.09E-06 | 2.09E-09 | 2.84E-13 | 2.52E-01 |
| ARS3008 | 6.89E-07 | 1.76E-06 | 3.85E-05 | 9.56E-08 | 2.73E-06 | 1.04E-02 | 4.10E-01 | 1.00E+00 | 1.48E-17 | 1.36E-02 | 2.26E-28 | 5.51E-31 | 8.08E-16 | 3.74E-02 | 3.70E-20 | 1.18E-03 | 3.43E-08 | 3.42E-10 | 1.73E-15 | 1.79E-02 |
| ARS3009 | 3.85E-05 | 6.38E-09 | 6.38E-09 | 4.03E-04 | 1.85E-10 | 5.39E-11 | 2.78E-18 | 1.48E-17 | 1.00E+00 | 1.16E-09 | 1.08E-05 | 9.56E-08 | 3.53E-01 | 1.03E-26 | 4.75E-01 | 1.85E-10 | 2.28E-03 | 9.66E-02 | 1.46E-01 | 5.55E-13 |
| ARS3010 | 2.95E-02 | 2.52E-01 | 2.98E-01 | 7.68E-03 | 1.19E-01 | 1.00E+00 | 1.19E-01 | 1.36E-02 | 1.16E-09 | 1.00E+00 | 4.12E-22 | 1.44E-24 | 5.88E-08 | 2.76E-04 | 8.08E-12 | 1.00E+00 | 4.34E-03 | 2.54E-05 | 9.56E-08 | 1.00E+00 |
| Meta_AMS3002 | 7.56E-17 | 6.56E-23 | 2.59E-23 | 2.18E-12 | 1.44E-24 | 1.65E-22 | 1.03E-26 | 2.26E-28 | 1.08E-05 | 4.12E-22 | 1.00E+00 | 6.19E-01 | 3.66E-09 | 1.93E-46 | 7.68E-03 | 2.63E-21 | 5.39E-11 | 2.59E-07 | 1.85E-10 | 2.10E-25 |
| Meta_AMS3003 | 3.70E-20 | 1.03E-26 | 4.71E-27 | 3.57E-15 | 5.98E-28 | 7.95E-26 | 1.33E-30 | 5.51E-31 | 9.56E-08 | 1.44E-24 | 6.19E-01 | 1.00E+00 | 1.55E-11 | 3.75E-50 | 2.76E-04 | 1.02E-23 | 1.44E-13 | 1.12E-08 | 2.84E-13 | 8.62E-29 |
| Meta_AMS3004 | 2.95E-02 | 1.92E-04 | 1.92E-04 | 9.66E-02 | 1.68E-05 | 6.89E-07 | 1.44E-13 | 8.08E-16 | 3.53E-01 | 5.88E-08 | 3.66E-09 | 1.55E-11 | 1.00E+00 | 8.80E-20 | 7.68E-03 | 2.73E-06 | 1.46E-01 | 2.95E-02 | 1.00E+00 | 1.61E-07 |
| Meta_AMS3005 | 3.66E-09 | 2.59E-07 | 1.09E-06 | 2.91E-11 | 9.56E-08 | 5.73E-04 | 1.36E-02 | 3.74E-02 | 1.03E-26 | 2.76E-04 | 1.93E-46 | 3.75E-50 | 8.80E-20 | 1.00E+00 | 1.07E-29 | 1.92E-04 | 2.91E-11 | 8.08E-16 | 2.07E-19 | 3.17E-03 |
| Meta_AMS3006 | 2.59E-07 | 8.08E-12 | 8.08E-12 | 2.54E-05 | 1.44E-13 | 5.55E-13 | 6.39E-21 | 3.70E-20 | 4.75E-01 | 8.08E-12 | 7.68E-03 | 2.76E-04 | 7.68E-03 | 1.07E-29 | 1.00E+00 | 2.18E-12 | 1.92E-04 | 1.79E-02 | 7.68E-03 | 3.57E-15 |
| Meta_AMS3007 | 1.76E-01 | 7.79E-01 | 8.52E-01 | 6.17E-02 | 4.75E-01 | 1.00E+00 | 9.66E-02 | 1.18E-03 | 1.85E-10 | 1.00E+00 | 2.63E-21 | 1.02E-23 | 2.73E-06 | 1.92E-04 | 2.18E-12 | 1.00E+00 | 2.32E-02 | 5.79E-05 | 4.35E-06 | 9.87E-01 |
| Meta_AMS3008 | 4.10E-01 | 2.32E-02 | 7.68E-03 | 7.79E-01 | 7.68E-03 | 1.36E-02 | 1.09E-06 | 3.43E-08 | 2.28E-03 | 4.34E-03 | 5.39E-11 | 1.44E-13 | 1.46E-01 | 2.91E-11 | 1.92E-04 | 2.32E-02 | 1.00E+00 | 2.11E-01 | 1.76E-01 | 1.63E-03 |
| Meta_AMS3009 | 1.63E-03 | 2.73E-06 | 1.09E-06 | 2.95E-02 | 6.89E-07 | 6.89E-06 | 2.09E-09 | 3.42E-10 | 9.66E-02 | 2.54E-05 | 2.59E-07 | 1.12E-08 | 2.95E-02 | 8.08E-16 | 1.79E-02 | 5.79E-05 | 2.11E-01 | 1.00E+00 | 5.94E-03 | 2.59E-07 |
| Meta_AMS3010 | 3.74E-02 | 5.73E-04 | 5.73E-04 | 1.19E-01 | 5.79E-05 | 1.09E-06 | 2.84E-13 | 1.73E-15 | 1.46E-01 | 9.56E-08 | 1.85E-10 | 2.84E-13 | 1.00E+00 | 2.07E-19 | 7.68E-03 | 4.35E-06 | 1.76E-01 | 5.94E-03 | 1.00E+00 | 4.30E-07 |
| Meta_AMS3011 | 2.32E-02 | 2.11E-01 | 4.10E-01 | 4.34E-03 | 2.11E-01 | 1.00E+00 | 2.52E-01 | 1.79E-02 | 5.55E-13 | 1.00E+00 | 2.10E-25 | 8.62E-29 | 1.61E-07 | 3.17E-03 | 3.57E-15 | 9.87E-01 | 1.63E-03 | 2.59E-07 | 4.30E-07 | 1.00E+00 |

Comparison of trees using the Kuhner & Felsenstein, 1994 method

| | ARS3001 | ARS3002 | ARS3003 | ARS3004 | ARS3005 | ARS3006 | ARS3007 | ARS3008 | ARS3009 | ARS3010 | Meta_AMS3002 | Meta_AMS3003 | Meta_AMS3004 | Meta_AMS3005 | Meta_AMS3006 | Meta_AMS3007 | Meta_AMS3008 | Meta_AMS3009 | Meta_AMS3010 | Meta_AMS3011 |
|---|---|---|---|---|---|---|---|---|---|---|---|---|---|---|---|---|---|---|---|---|
| ARS3001 | 1.00E+00 | 2.54E-07 | 1.03E-09 | 3.11E-10 | 1.41E-04 | 2.99E-04 | 2.37E-01 | 6.20E-04 | 5.86E-01 | 1.80E-05 | 9.31E-38 | 9.60E-42 | 1.50E-48 | 6.81E-25 | 1.38E-40 | 9.12E-45 | 5.65E-16 | 3.41E-61 | 7.93E-19 | 6.06E-32 |
| ARS3002 | 2.54E-07 | 1.00E+00 | 5.86E-01 | 8.52E-02 | 3.31E-01 | 3.35E-03 | 3.48E-12 | 2.81E-01 | 5.89E-09 | 5.86E-01 | 7.88E-70 | 3.20E-68 | 1.74E-77 | 6.94E-52 | 1.29E-66 | 3.79E-73 | 1.92E-39 | 9.71E-85 | 1.52E-43 | 3.41E-61 |
| ARS3003 | 1.03E-09 | 5.86E-01 | 1.00E+00 | 5.24E-01 | 8.52E-02 | 1.25E-03 | 5.26E-14 | 4.30E-02 | 9.09E-11 | 1.98E-01 | 3.79E-73 | 7.88E-70 | 3.04E-78 | 4.49E-56 | 2.09E-67 | 1.74E-77 | 9.12E-45 | 1.42E-85 | 2.34E-45 | 3.03E-64 |
| ARS3004 | 3.11E-10 | 8.52E-02 | 5.24E-01 | 1.00E+00 | 8.32E-03 | 1.13E-06 | 5.65E-16 | 1.49E-02 | 8.87E-13 | 1.32E-01 | 1.29E-70 | 1.29E-70 | 4.67E-80 | 1.69E-57 | 3.20E-68 | 3.79E-73 | 9.12E-45 | 2.50E-87 | 1.30E-46 | 1.83E-63 |
| ARS3005 | 1.41E-04 | 3.31E-01 | 8.52E-02 | 8.32E-03 | 1.00E+00 | 8.32E-03 | 5.36E-08 | 6.61E-01 | 1.16E-05 | 3.90E-01 | 1.03E-62 | 3.41E-61 | 1.29E-70 | 1.52E-43 | 8.95E-57 | 1.29E-66 | 6.06E-32 | 1.74E-77 | 4.96E-35 | 6.00E-54 |
| ARS3006 | 2.99E-04 | 3.35E-03 | 1.25E-03 | 1.13E-06 | 8.32E-03 | 1.00E+00 | 2.54E-07 | 5.45E-02 | 4.14E-05 | 1.76E-03 | 1.03E-62 | 5.29E-65 | 7.90E-75 | 9.12E-45 | 1.89E-60 | 5.05E-69 | 1.91E-31 | 6.39E-81 | 1.46E-35 | 1.20E-54 |
| ARS3007 | 2.37E-01 | 3.48E-12 | 5.26E-14 | 5.65E-16 | 5.36E-08 | 2.54E-07 | 1.00E+00 | 9.11E-08 | 2.81E-01 | 1.03E-09 | 1.83E-33 | 2.58E-38 | 5.55E-46 | 1.53E-21 | 7.08E-39 | 3.66E-41 | 2.22E-13 | 1.01E-59 | 2.66E-15 | 1.74E-29 |
| ARS3008 | 6.20E-04 | 2.81E-01 | 4.30E-02 | 1.49E-02 | 6.61E-01 | 5.45E-02 | 9.11E-08 | 1.00E+00 | 1.16E-05 | 5.86E-01 | 1.01E-59 | 4.49E-56 | 3.03E-64 | 3.66E-41 | 2.96E-53 | 1.03E-62 | 5.75E-30 | 7.88E-70 | 1.83E-33 | 3.29E-51 |
| ARS3009 | 5.86E-01 | 5.89E-09 | 9.09E-11 | 8.87E-13 | 1.16E-05 | 4.14E-05 | 2.81E-01 | 1.16E-05 | 1.00E+00 | 3.13E-08 | 5.14E-40 | 1.38E-40 | 1.50E-48 | 6.81E-25 | 5.14E-40 | 9.12E-45 | 5.65E-16 | 5.62E-59 | 7.93E-19 | 5.89E-33 |
| ARS3010 | 1.80E-05 | 5.86E-01 | 1.98E-01 | 1.32E-01 | 3.90E-01 | 1.76E-03 | 1.03E-09 | 5.86E-01 | 3.13E-08 | 1.00E+00 | 1.89E-60 | 1.01E-59 | 5.05E-69 | 9.60E-42 | 4.49E-56 | 3.03E-64 | 5.89E-33 | 1.08E-75 | 1.67E-34 | 6.94E-52 |
| Meta_AMS3002 | 9.31E-38 | 7.88E-70 | 3.79E-73 | 1.29E-70 | 1.03E-62 | 1.03E-62 | 1.83E-33 | 1.01E-59 | 5.14E-40 | 1.89E-60 | 1.00E+00 | 8.52E-02 | 6.26E-05 | 6.86E-02 | 8.85E-04 | 8.32E-03 | 2.74E-05 | 9.09E-11 | 2.99E-04 | 8.11E-01 |
| Meta_AMS3003 | 9.60E-42 | 3.20E-68 | 7.88E-70 | 1.29E-70 | 3.41E-61 | 5.29E-65 | 2.58E-38 | 4.49E-56 | 1.38E-40 | 1.01E-59 | 8.52E-02 | 1.00E+00 | 6.86E-02 | 3.35E-03 | 2.37E-01 | 5.86E-01 | 3.13E-08 | 2.95E-06 | 2.54E-07 | 5.45E-02 |
| Meta_AMS3004 | 1.50E-48 | 1.74E-77 | 3.04E-78 | 4.67E-80 | 1.29E-70 | 7.90E-75 | 5.55E-46 | 3.03E-64 | 1.50E-48 | 5.05E-69 | 6.26E-05 | 6.86E-02 | 1.00E+00 | 1.13E-06 | 5.24E-01 | 3.31E-01 | 4.48E-13 | 2.56E-02 | 8.87E-13 | 2.74E-05 |
| Meta_AMS3005 | 6.81E-25 | 6.94E-52 | 4.49E-56 | 1.69E-57 | 1.52E-43 | 9.12E-45 | 1.53E-21 | 3.66E-41 | 6.81E-25 | 9.60E-42 | 6.86E-02 | 3.35E-03 | 1.13E-06 | 1.00E+00 | 4.14E-05 | 2.06E-04 | 6.25E-03 | 2.54E-14 | 2.56E-02 | 2.81E-01 |
| Meta_AMS3006 | 1.38E-40 | 1.29E-66 | 2.09E-67 | 3.20E-68 | 8.95E-57 | 1.89E-60 | 7.08E-39 | 2.96E-53 | 5.14E-40 | 4.49E-56 | 8.85E-04 | 2.37E-01 | 5.24E-01 | 4.14E-05 | 1.00E+00 | 7.38E-01 | 9.09E-11 | 3.35E-03 | 1.68E-10 | 6.20E-04 |
| Meta_AMS3007 | 9.12E-45 | 3.79E-73 | 1.74E-77 | 3.79E-73 | 1.29E-66 | 5.05E-69 | 3.66E-41 | 1.03E-62 | 9.12E-45 | 3.03E-64 | 8.32E-03 | 5.86E-01 | 3.31E-01 | 2.06E-04 | 7.38E-01 | 1.00E+00 | 1.03E-09 | 2.06E-04 | 5.89E-09 | 6.25E-03 |
| Meta_AMS3008 | 5.65E-16 | 1.92E-39 | 9.12E-45 | 9.12E-45 | 6.06E-32 | 1.91E-31 | 2.22E-13 | 5.75E-30 | 5.65E-16 | 5.89E-33 | 2.74E-05 | 3.13E-08 | 4.48E-13 | 6.25E-03 | 9.09E-11 | 1.03E-09 | 1.00E+00 | 9.11E-23 | 1.00E+00 | 6.20E-04 |
| Meta_AMS3009 | 3.41E-61 | 9.71E-85 | 1.42E-85 | 2.50E-87 | 1.74E-77 | 6.39E-81 | 1.01E-59 | 7.88E-70 | 5.62E-59 | 1.08E-75 | 9.09E-11 | 2.95E-06 | 2.56E-02 | 2.54E-14 | 3.35E-03 | 2.06E-04 | 9.11E-23 | 1.00E+00 | 3.51E-23 | 1.68E-10 |
| Meta_AMS3010 | 7.93E-19 | 1.52E-43 | 2.34E-45 | 1.30E-46 | 4.96E-35 | 1.46E-35 | 2.66E-15 | 1.83E-33 | 7.93E-19 | 1.67E-34 | 2.99E-04 | 2.54E-07 | 8.87E-13 | 2.56E-02 | 1.68E-10 | 5.89E-09 | 1.00E+00 | 3.51E-23 | 1.00E+00 | 3.35E-03 |
| Meta_AMS3011 | 6.06E-32 | 3.41E-61 | 3.03E-64 | 1.83E-63 | 6.00E-54 | 1.20E-54 | 1.74E-29 | 3.29E-51 | 5.89E-33 | 6.94E-52 | 8.11E-01 | 5.45E-02 | 2.74E-05 | 2.81E-01 | 6.20E-04 | 6.25E-03 | 6.20E-04 | 1.68E-10 | 3.35E-03 | 1.00E+00 |

**Supplementary Table S4C:** Pairwise FDR corrected p-values of the Kolmogorow-Smirnow test on the distances between phylogenetic trees for 16 non-ribosomal proteins in 10 samples of archaeal RefSeq genomes vs 10 samples of archaeal MAGs (associated with Supplementary Fig. S7c)

Comparison of trees using the Nye et al., 2006 method

| | ARS3001 | ARS3002 | ARS3003 | ARS3004 | ARS3005 | ARS3006 | ARS3007 | ARS3008 | ARS3009 | ARS3010 | Meta_AMS3002 | Meta_AMS3003 | Meta_AMS3004 | Meta_AMS3005 | Meta_AMS3006 | Meta_AMS3007 | Meta_AMS3008 | Meta_AMS3009 | Meta_AMS3010 | Meta_AMS3011 |
|---|---|---|---|---|---|---|---|---|---|---|---|---|---|---|---|---|---|---|---|---|
| ARS3001 | 1.00E+00 | 2.90E-03 | 1.59E-02 | 2.36E-02 | 5.86E-07 | 1.07E-03 | 2.90E-03 | 5.83E-08 | 6.78E-02 | 8.48E-01 | 1.99E-53 | 2.55E-45 | 4.58E-57 | 2.80E-59 | 1.20E-54 | 1.20E-54 | 1.16E-48 | 1.99E-53 | 3.66E-58 | 4.58E-57 |
| ARS3002 | 2.90E-03 | 1.00E+00 | 4.39E-01 | 2.22E-01 | 1.27E-01 | 8.48E-01 | 2.87E-01 | 2.36E-02 | 9.70E-06 | 1.59E-02 | 5.64E-51 | 3.46E-42 | 5.64E-51 | 1.20E-54 | 3.35E-43 | 2.51E-37 | 2.07E-46 | 3.35E-43 | 1.60E-47 | 1.16E-48 |
| ARS3003 | 1.59E-02 | 4.39E-01 | 1.00E+00 | 6.50E-01 | 4.86E-02 | 5.40E-01 | 7.53E-01 | 1.06E-02 | 1.07E-03 | 1.69E-01 | 1.99E-53 | 2.55E-45 | 7.12E-56 | 7.12E-56 | 8.24E-50 | 3.35E-43 | 1.16E-48 | 8.24E-50 | 1.20E-54 | 3.32E-52 |
| ARS3004 | 2.36E-02 | 2.22E-01 | 6.50E-01 | 1.00E+00 | 1.06E-02 | 6.50E-01 | 4.39E-01 | 4.47E-03 | 1.07E-03 | 4.86E-02 | 1.99E-53 | 2.55E-45 | 1.20E-54 | 7.12E-56 | 2.07E-46 | 3.46E-42 | 1.16E-48 | 1.60E-47 | 1.20E-54 | 5.64E-51 |
| ARS3005 | 5.86E-07 | 1.27E-01 | 4.86E-02 | 1.06E-02 | 1.00E+00 | 9.37E-02 | 7.05E-03 | 8.48E-01 | 2.02E-09 | 3.48E-05 | 8.24E-50 | 3.52E-41 | 8.24E-50 | 1.99E-53 | 3.46E-42 | 2.06E-36 | 3.04E-44 | 3.52E-41 | 1.60E-47 | 1.16E-48 |
| ARS3006 | 1.07E-03 | 8.48E-01 | 5.40E-01 | 6.50E-01 | 9.37E-02 | 1.00E+00 | 5.40E-01 | 6.78E-02 | 3.48E-05 | 4.47E-03 | 3.32E-52 | 3.35E-43 | 5.64E-51 | 7.12E-56 | 2.07E-46 | 3.28E-39 | 2.07E-46 | 3.04E-44 | 8.24E-50 | 1.16E-48 |
| ARS3007 | 2.90E-03 | 2.87E-01 | 7.53E-01 | 4.39E-01 | 7.05E-03 | 5.40E-01 | 1.00E+00 | 6.46E-04 | 4.96E-06 | 2.36E-02 | 3.32E-52 | 3.35E-43 | 1.99E-53 | 7.12E-56 | 1.60E-47 | 3.46E-42 | 2.07E-46 | 1.16E-48 | 3.32E-52 | 3.32E-52 |
| ARS3008 | 5.83E-08 | 2.36E-02 | 1.06E-02 | 4.47E-03 | 8.48E-01 | 6.78E-02 | 6.46E-04 | 1.00E+00 | 8.35E-10 | 2.78E-07 | 8.24E-50 | 3.52E-41 | 8.24E-50 | 1.99E-53 | 3.46E-42 | 2.51E-37 | 2.55E-45 | 3.46E-42 | 1.16E-48 | 1.60E-47 |
| ARS3009 | 6.78E-02 | 9.70E-06 | 1.07E-03 | 1.07E-03 | 2.02E-09 | 3.48E-05 | 4.96E-06 | 8.35E-10 | 1.00E+00 | 6.78E-02 | 1.99E-53 | 2.55E-45 | 4.58E-57 | 3.66E-58 | 3.32E-52 | 3.32E-52 | 8.24E-50 | 1.99E-53 | 4.58E-57 | 4.58E-57 |
| ARS3010 | 8.48E-01 | 1.59E-02 | 1.69E-01 | 4.86E-02 | 3.48E-05 | 4.47E-03 | 2.36E-02 | 2.78E-07 | 6.78E-02 | 1.00E+00 | 3.32E-52 | 3.04E-44 | 7.12E-56 | 7.12E-56 | 8.24E-50 | 2.07E-46 | 1.60E-47 | 5.64E-51 | 1.20E-54 | 1.20E-54 |
| Meta_AMS3002 | 1.99E-53 | 5.64E-51 | 1.99E-53 | 1.99E-53 | 8.24E-50 | 3.32E-52 | 3.32E-52 | 8.24E-50 | 1.99E-53 | 3.32E-52 | 1.00E+00 | 1.27E-01 | 7.53E-01 | 6.78E-02 | 4.47E-03 | 1.69E-01 | 2.87E-01 | 4.47E-03 | 4.39E-01 | 3.65E-01 |
| Meta_AMS3003 | 2.55E-45 | 3.46E-42 | 2.55E-45 | 2.55E-45 | 3.52E-41 | 3.35E-43 | 3.35E-43 | 3.52E-41 | 2.55E-45 | 3.04E-44 | 1.27E-01 | 1.00E+00 | 4.39E-01 | 3.75E-04 | 1.06E-02 | 4.47E-03 | 2.22E-01 | 4.86E-02 | 2.22E-01 | 4.39E-01 |
| Meta_AMS3004 | 4.58E-57 | 5.64E-51 | 7.12E-56 | 1.20E-54 | 8.24E-50 | 5.64E-51 | 1.99E-53 | 8.24E-50 | 4.58E-57 | 7.12E-56 | 7.53E-01 | 4.39E-01 | 1.00E+00 | 1.06E-02 | 1.59E-02 | 2.87E-01 | 4.39E-01 | 9.37E-02 | 8.48E-01 | 9.37E-01 |
| Meta_AMS3005 | 2.80E-59 | 1.20E-54 | 7.12E-56 | 7.12E-56 | 1.99E-53 | 7.12E-56 | 7.12E-56 | 1.99E-53 | 3.66E-58 | 7.12E-56 | 6.78E-02 | 3.75E-04 | 1.06E-02 | 1.00E+00 | 1.21E-06 | 1.06E-02 | 3.75E-04 | 1.86E-05 | 3.42E-02 | 1.79E-03 |
| Meta_AMS3006 | 1.20E-54 | 3.35E-43 | 8.24E-50 | 2.07E-46 | 3.46E-42 | 2.07E-46 | 1.60E-47 | 3.46E-42 | 3.32E-52 | 8.24E-50 | 4.47E-03 | 1.06E-02 | 1.59E-02 | 1.21E-06 | 1.00E+00 | 3.42E-02 | 4.39E-01 | 3.42E-02 | 4.86E-02 | 1.69E-01 |
| Meta_AMS3007 | 1.20E-54 | 2.51E-37 | 3.35E-43 | 3.46E-42 | 2.06E-36 | 3.28E-39 | 3.46E-42 | 2.51E-37 | 3.32E-52 | 2.07E-46 | 1.69E-01 | 4.47E-03 | 2.87E-01 | 1.06E-02 | 3.42E-02 | 1.00E+00 | 4.39E-01 | 4.47E-03 | 3.65E-01 | 1.27E-01 |
| Meta_AMS3008 | 1.16E-48 | 2.07E-46 | 1.16E-48 | 1.16E-48 | 3.04E-44 | 2.07E-46 | 2.07E-46 | 2.55E-45 | 8.24E-50 | 1.60E-47 | 2.87E-01 | 2.22E-01 | 4.39E-01 | 3.75E-04 | 4.39E-01 | 4.39E-01 | 1.00E+00 | 9.37E-02 | 4.39E-01 | 7.53E-01 |
| Meta_AMS3009 | 1.99E-53 | 3.35E-43 | 8.24E-50 | 1.60E-47 | 3.52E-41 | 3.04E-44 | 1.16E-48 | 3.46E-42 | 1.99E-53 | 5.64E-51 | 4.47E-03 | 4.86E-02 | 9.37E-02 | 1.86E-05 | 3.42E-02 | 4.47E-03 | 9.37E-02 | 1.00E+00 | 1.69E-01 | 3.65E-01 |
| Meta_AMS3010 | 3.66E-58 | 1.60E-47 | 1.20E-54 | 1.20E-54 | 1.60E-47 | 8.24E-50 | 3.32E-52 | 1.16E-48 | 4.58E-57 | 1.20E-54 | 4.39E-01 | 2.22E-01 | 8.48E-01 | 3.42E-02 | 4.86E-02 | 3.65E-01 | 4.39E-01 | 1.69E-01 | 1.00E+00 | 7.53E-01 |
| Meta_AMS3011 | 4.58E-57 | 1.16E-48 | 3.32E-52 | 5.64E-51 | 1.16E-48 | 1.16E-48 | 3.32E-52 | 1.60E-47 | 4.58E-57 | 1.20E-54 | 3.65E-01 | 4.39E-01 | 9.37E-01 | 1.79E-03 | 1.69E-01 | 1.27E-01 | 7.53E-01 | 3.65E-01 | 7.53E-01 | 1.00E+00 |

Comparison of trees using the Williams & Clifford, 1971 method

| | ARS3001 | ARS3002 | ARS3003 | ARS3004 | ARS3005 | ARS3006 | ARS3007 | ARS3008 | ARS3009 | ARS3010 | Meta_AMS3002 | Meta_AMS3003 | Meta_AMS3004 | Meta_AMS3005 | Meta_AMS3006 | Meta_AMS3007 | Meta_AMS3008 | Meta_AMS3009 | Meta_AMS3010 | Meta_AMS3011 |
|---|---|---|---|---|---|---|---|---|---|---|---|---|---|---|---|---|---|---|---|---|
| ARS3001 | 1.00E+00 | 1.00E-01 | 1.78E-01 | 3.56E-01 | 1.10E-02 | 2.90E-03 | 5.34E-01 | 3.52E-05 | 1.00E-01 | 3.69E-02 | 7.64E-46 | 5.23E-38 | 6.01E-48 | 4.23E-50 | 4.15E-37 | 4.15E-37 | 8.71E-42 | 7.25E-47 | 7.85E-41 | 7.64E-46 |
| ARS3002 | 1.00E-01 | 1.00E+00 | 6.39E-01 | 7.37E-01 | 2.50E-02 | 7.38E-02 | 3.56E-01 | 1.16E-04 | 3.52E-05 | 3.56E-01 | 8.60E-44 | 5.23E-38 | 6.01E-48 | 4.23E-50 | 3.16E-36 | 2.20E-35 | 8.71E-42 | 7.64E-46 | 5.23E-38 | 7.64E-46 |
| ARS3003 | 1.78E-01 | 6.39E-01 | 1.00E+00 | 2.84E-01 | 5.30E-02 | 2.84E-01 | 2.84E-01 | 1.10E-02 | 1.16E-04 | 5.34E-01 | 7.85E-41 | 3.16E-36 | 8.60E-44 | 6.01E-48 | 1.55E-34 | 7.59E-33 | 6.24E-39 | 7.85E-41 | 2.20E-35 | 8.57E-43 |
| ARS3004 | 3.56E-01 | 7.37E-01 | 2.84E-01 | 1.00E+00 | 1.00E-01 | 1.35E-01 | 7.37E-01 | 4.61E-03 | 2.06E-04 | 5.34E-01 | 8.57E-43 | 4.15E-37 | 7.64E-46 | 5.56E-49 | 2.20E-35 | 1.55E-34 | 7.85E-41 | 8.39E-45 | 3.16E-36 | 8.39E-45 |
| ARS3005 | 1.10E-02 | 2.50E-02 | 5.30E-02 | 1.00E-01 | 1.00E+00 | 9.37E-01 | 7.17E-03 | 5.34E-01 | 5.97E-07 | 5.34E-01 | 3.10E-31 | 1.12E-29 | 1.55E-34 | 6.24E-39 | 3.67E-28 | 3.67E-28 | 1.87E-30 | 1.55E-34 | 5.54E-26 | 5.02E-32 |
| ARS3006 | 2.90E-03 | 7.38E-02 | 2.84E-01 | 1.35E-01 | 9.37E-01 | 1.00E+00 | 7.17E-03 | 2.28E-01 | 5.89E-08 | 7.37E-01 | 2.20E-35 | 1.55E-34 | 4.15E-37 | 8.57E-43 | 3.10E-31 | 3.10E-31 | 1.09E-33 | 6.24E-39 | 1.12E-29 | 2.20E-35 |
| ARS3007 | 5.34E-01 | 3.56E-01 | 2.84E-01 | 7.37E-01 | 7.17E-03 | 7.17E-03 | 1.00E+00 | 3.52E-05 | 2.90E-03 | 1.35E-01 | 8.71E-42 | 3.16E-36 | 8.39E-45 | 6.01E-48 | 2.20E-35 | 1.09E-33 | 7.30E-40 | 8.60E-44 | 4.15E-37 | 8.60E-44 |
| ARS3008 | 3.52E-05 | 1.16E-04 | 1.10E-02 | 4.61E-03 | 5.34E-01 | 2.28E-01 | 3.52E-05 | 1.00E+00 | 1.15E-08 | 7.38E-02 | 2.20E-35 | 1.55E-34 | 7.30E-40 | 8.60E-44 | 3.10E-31 | 1.87E-30 | 2.20E-35 | 6.24E-39 | 1.87E-30 | 5.23E-38 |
| ARS3009 | 1.00E-01 | 3.52E-05 | 1.16E-04 | 2.06E-04 | 5.97E-07 | 5.89E-08 | 2.90E-03 | 1.15E-08 | 1.00E+00 | 1.90E-05 | 4.43E-51 | 7.85E-41 | 5.56E-49 | 6.36E-52 | 8.57E-43 | 8.57E-43 | 7.25E-47 | 4.43E-51 | 7.25E-47 | 4.23E-50 |
| ARS3010 | 3.69E-02 | 3.56E-01 | 5.34E-01 | 5.34E-01 | 5.34E-01 | 7.37E-01 | 1.35E-01 | 7.38E-02 | 1.90E-05 | 1.00E+00 | 3.16E-36 | 1.09E-33 | 7.85E-41 | 8.39E-45 | 3.10E-31 | 5.02E-32 | 2.20E-35 | 7.30E-40 | 1.87E-30 | 6.24E-39 |
| Meta_AMS3002 | 7.64E-46 | 8.60E-44 | 7.85E-41 | 8.57E-43 | 3.10E-31 | 2.20E-35 | 8.71E-42 | 2.20E-35 | 4.43E-51 | 3.16E-36 | 1.00E+00 | 7.38E-02 | 2.28E-01 | 1.78E-01 | 2.84E-01 | 4.45E-01 | 7.38E-02 | 1.78E-01 | 6.45E-05 | 7.37E-01 |
| Meta_AMS3003 | 5.23E-38 | 5.23E-38 | 3.16E-36 | 4.15E-37 | 1.12E-29 | 1.55E-34 | 3.16E-36 | 1.55E-34 | 7.85E-41 | 1.09E-33 | 7.38E-02 | 1.00E+00 | 4.45E-01 | 6.29E-04 | 2.28E-01 | 2.50E-02 | 2.84E-01 | 2.28E-01 | 2.50E-02 | 1.35E-01 |
| Meta_AMS3004 | 6.01E-48 | 6.01E-48 | 8.60E-44 | 7.64E-46 | 1.55E-34 | 4.15E-37 | 8.39E-45 | 7.30E-40 | 5.56E-49 | 7.85E-41 | 2.28E-01 | 4.45E-01 | 1.00E+00 | 1.00E-01 | 6.39E-01 | 2.28E-01 | 4.45E-01 | 1.78E-01 | 7.17E-03 | 3.56E-01 |
| Meta_AMS3005 | 4.23E-50 | 4.23E-50 | 6.01E-48 | 5.56E-49 | 6.24E-39 | 8.57E-43 | 6.01E-48 | 8.60E-44 | 6.36E-52 | 8.39E-45 | 1.78E-01 | 6.29E-04 | 1.00E-01 | 1.00E+00 | 1.00E-01 | 2.84E-01 | 5.30E-02 | 4.61E-03 | 2.51E-06 | 2.28E-01 |
| Meta_AMS3006 | 4.15E-37 | 3.16E-36 | 1.55E-34 | 2.20E-35 | 3.67E-28 | 3.10E-31 | 2.20E-35 | 3.10E-31 | 8.57E-43 | 3.10E-31 | 2.84E-01 | 2.28E-01 | 6.39E-01 | 1.00E-01 | 1.00E+00 | 1.35E-01 | 3.56E-01 | 2.84E-01 | 2.90E-03 | 7.37E-01 |
| Meta_AMS3007 | 4.15E-37 | 2.20E-35 | 7.59E-33 | 1.55E-34 | 3.67E-28 | 3.10E-31 | 1.09E-33 | 1.87E-30 | 8.57E-43 | 5.02E-32 | 4.45E-01 | 2.50E-02 | 2.28E-01 | 2.84E-01 | 1.35E-01 | 1.00E+00 | 2.84E-01 | 2.28E-01 | 2.51E-06 | 5.34E-01 |
| Meta_AMS3008 | 8.71E-42 | 8.71E-42 | 6.24E-39 | 7.85E-41 | 1.87E-30 | 1.09E-33 | 7.30E-40 | 2.20E-35 | 7.25E-47 | 2.20E-35 | 7.38E-02 | 2.84E-01 | 4.45E-01 | 5.30E-02 | 3.56E-01 | 2.84E-01 | 1.00E+00 | 6.39E-01 | 6.45E-05 | 1.78E-01 |
| Meta_AMS3009 | 7.25E-47 | 7.64E-46 | 7.85E-41 | 8.39E-45 | 1.55E-34 | 6.24E-39 | 8.60E-44 | 6.24E-39 | 4.43E-51 | 7.30E-40 | 1.78E-01 | 2.28E-01 | 1.78E-01 | 4.61E-03 | 2.84E-01 | 2.28E-01 | 6.39E-01 | 1.00E+00 | 1.90E-05 | 2.84E-01 |
| Meta_AMS3010 | 7.85E-41 | 5.23E-38 | 2.20E-35 | 3.16E-36 | 5.54E-26 | 1.12E-29 | 4.15E-37 | 1.87E-30 | 7.25E-47 | 1.87E-30 | 6.45E-05 | 2.50E-02 | 7.17E-03 | 2.51E-06 | 2.90E-03 | 2.51E-06 | 6.45E-05 | 1.90E-05 | 1.00E+00 | 2.06E-04 |
| Meta_AMS3011 | 7.64E-46 | 7.64E-46 | 8.57E-43 | 8.39E-45 | 5.02E-32 | 2.20E-35 | 8.60E-44 | 5.23E-38 | 4.23E-50 | 6.24E-39 | 7.37E-01 | 1.35E-01 | 3.56E-01 | 2.28E-01 | 7.37E-01 | 5.34E-01 | 1.78E-01 | 2.84E-01 | 2.06E-04 | 1.00E+00 |

Comparison of trees using the Robinson & Foulds, 1981 method

| | ARS3001 | ARS3002 | ARS3003 | ARS3004 | ARS3005 | ARS3006 | ARS3007 | ARS3008 | ARS3009 | ARS3010 | Meta_AMS3002 | Meta_AMS3003 | Meta_AMS3004 | Meta_AMS3005 | Meta_AMS3006 | Meta_AMS3007 | Meta_AMS3008 | Meta_AMS3009 | Meta_AMS3010 | Meta_AMS3011 |
|---|---|---|---|---|---|---|---|---|---|---|---|---|---|---|---|---|---|---|---|---|
| ARS3001 | 1.00E+00 | 7.07E-02 | 3.52E-02 | 2.45E-02 | 6.94E-03 | 9.56E-02 | 5.03E-02 | 2.08E-04 | 1.76E-03 | 9.63E-01 | 7.46E-45 | 5.54E-38 | 9.40E-43 | 3.17E-50 | 1.63E-34 | 1.18E-29 | 3.07E-36 | 3.07E-36 | 4.15E-37 | 9.40E-43 |
| ARS3002 | 7.07E-02 | 1.00E+00 | 2.24E-01 | 9.56E-02 | 3.52E-02 | 3.52E-02 | 3.69E-01 | 2.08E-04 | 1.10E-08 | 5.59E-01 | 6.01E-48 | 8.88E-41 | 8.49E-46 | 2.12E-52 | 2.23E-35 | 1.18E-29 | 6.77E-39 | 4.15E-37 | 7.81E-40 | 8.49E-46 |
| ARS3003 | 3.52E-02 | 2.24E-01 | 1.00E+00 | 1.00E+00 | 8.85E-01 | 1.00E+00 | 1.00E+00 | 2.24E-01 | 2.51E-08 | 2.24E-01 | 7.25E-47 | 7.81E-40 | 7.46E-45 | 2.66E-51 | 1.15E-33 | 6.78E-29 | 5.54E-38 | 3.07E-36 | 6.77E-39 | 7.46E-45 |
| ARS3004 | 2.45E-02 | 9.56E-02 | 1.00E+00 | 1.00E+00 | 1.00E+00 | 1.00E+00 | 8.85E-01 | 1.71E-01 | 5.70E-07 | 1.71E-01 | 8.60E-44 | 4.15E-37 | 8.71E-42 | 3.17E-50 | 3.28E-31 | 3.82E-28 | 2.23E-35 | 3.07E-36 | 3.07E-36 | 8.71E-42 |
| ARS3005 | 6.94E-03 | 3.52E-02 | 8.85E-01 | 1.00E+00 | 1.00E+00 | 8.85E-01 | 8.85E-01 | 4.58E-01 | 2.02E-09 | 7.07E-02 | 4.15E-37 | 3.28E-31 | 2.23E-35 | 8.60E-44 | 5.65E-26 | 6.43E-24 | 1.18E-29 | 3.28E-31 | 2.02E-30 | 2.23E-35 |
| ARS3006 | 9.56E-02 | 3.52E-02 | 1.00E+00 | 1.00E+00 | 8.85E-01 | 1.00E+00 | 6.72E-01 | 2.45E-02 | 5.62E-08 | 1.71E-01 | 8.71E-42 | 2.23E-35 | 7.81E-40 | 6.01E-48 | 1.18E-29 | 1.10E-26 | 1.15E-33 | 1.63E-34 | 1.63E-34 | 7.81E-40 |
| ARS3007 | 5.03E-02 | 3.69E-01 | 1.00E+00 | 8.85E-01 | 8.85E-01 | 6.72E-01 | 1.00E+00 | 9.56E-02 | 1.18E-06 | 2.89E-01 | 7.25E-47 | 7.81E-40 | 7.46E-45 | 2.66E-51 | 1.15E-33 | 6.78E-29 | 5.54E-38 | 3.07E-36 | 6.77E-39 | 7.46E-45 |
| ARS3008 | 2.08E-04 | 2.08E-04 | 2.24E-01 | 1.71E-01 | 4.58E-01 | 2.45E-02 | 9.56E-02 | 1.00E+00 | 3.51E-14 | 2.78E-03 | 6.77E-39 | 7.87E-33 | 4.15E-37 | 8.49E-46 | 2.08E-27 | 2.81E-25 | 3.28E-31 | 7.87E-33 | 5.27E-32 | 4.15E-37 |
| ARS3009 | 1.76E-03 | 1.10E-08 | 2.51E-08 | 5.70E-07 | 2.02E-09 | 5.62E-08 | 1.18E-06 | 3.51E-14 | 1.00E+00 | 6.51E-05 | 2.12E-52 | 7.46E-45 | 4.94E-49 | 2.75E-53 | 7.46E-45 | 4.15E-37 | 7.46E-45 | 8.71E-42 | 7.25E-47 | 6.01E-48 |
| ARS3010 | 9.63E-01 | 5.59E-01 | 2.24E-01 | 1.71E-01 | 7.07E-02 | 1.71E-01 | 2.89E-01 | 2.78E-03 | 6.51E-05 | 1.00E+00 | 7.46E-45 | 5.54E-38 | 8.71E-42 | 3.17E-50 | 2.23E-35 | 2.02E-30 | 4.15E-37 | 3.07E-36 | 5.54E-38 | 8.71E-42 |
| Meta_AMS3002 | 7.46E-45 | 6.01E-48 | 7.25E-47 | 8.60E-44 | 4.15E-37 | 8.71E-42 | 7.25E-47 | 6.77E-39 | 2.12E-52 | 7.46E-45 | 1.00E+00 | 2.89E-01 | 1.71E-01 | 1.71E-01 | 1.76E-03 | 9.56E-02 | 2.89E-01 | 1.18E-04 | 6.72E-01 | 3.69E-01 |
| Meta_AMS3003 | 5.54E-38 | 8.88E-41 | 7.81E-40 | 4.15E-37 | 3.28E-31 | 2.23E-35 | 7.81E-40 | 7.87E-33 | 7.46E-45 | 5.54E-38 | 2.89E-01 | 1.00E+00 | 8.85E-01 | 2.08E-04 | 1.30E-01 | 1.07E-02 | 4.58E-01 | 7.07E-02 | 7.92E-01 | 4.58E-01 |
| Meta_AMS3004 | 9.40E-43 | 8.49E-46 | 7.46E-45 | 8.71E-42 | 2.23E-35 | 7.81E-40 | 7.46E-45 | 4.15E-37 | 4.94E-49 | 8.71E-42 | 1.71E-01 | 8.85E-01 | 1.00E+00 | 6.29E-04 | 2.78E-03 | 1.07E-02 | 9.63E-01 | 2.24E-01 | 4.58E-01 | 5.59E-01 |
| Meta_AMS3005 | 3.17E-50 | 2.12E-52 | 2.66E-51 | 3.17E-50 | 8.60E-44 | 6.01E-48 | 2.66E-51 | 8.49E-46 | 2.75E-53 | 3.17E-50 | 1.71E-01 | 2.08E-04 | 6.29E-04 | 1.00E+00 | 1.10E-08 | 6.94E-03 | 1.76E-03 | 4.79E-09 | 2.78E-03 | 2.78E-03 |
| Meta_AMS3006 | 1.63E-34 | 2.23E-35 | 1.15E-33 | 3.28E-31 | 5.65E-26 | 1.18E-29 | 1.15E-33 | 2.08E-27 | 7.46E-45 | 2.23E-35 | 1.76E-03 | 1.30E-01 | 2.78E-03 | 1.10E-08 | 1.00E+00 | 6.94E-03 | 9.56E-02 | 6.94E-03 | 2.78E-03 | 3.65E-04 |
| Meta_AMS3007 | 1.18E-29 | 1.18E-29 | 6.78E-29 | 3.82E-28 | 6.43E-24 | 1.10E-26 | 6.78E-29 | 2.81E-25 | 4.15E-37 | 2.02E-30 | 9.56E-02 | 1.07E-02 | 1.07E-02 | 6.94E-03 | 6.94E-03 | 1.00E+00 | 5.03E-02 | 6.51E-05 | 5.03E-02 | 1.30E-01 |
| Meta_AMS3008 | 3.07E-36 | 6.77E-39 | 5.54E-38 | 2.23E-35 | 1.18E-29 | 1.15E-33 | 5.54E-38 | 3.28E-31 | 7.46E-45 | 4.15E-37 | 2.89E-01 | 4.58E-01 | 9.63E-01 | 1.76E-03 | 9.56E-02 | 5.03E-02 | 1.00E+00 | 1.30E-01 | 9.63E-01 | 5.59E-01 |
| Meta_AMS3009 | 3.07E-36 | 4.15E-37 | 3.07E-36 | 3.07E-36 | 3.28E-31 | 1.63E-34 | 3.07E-36 | 7.87E-33 | 8.71E-42 | 3.07E-36 | 1.18E-04 | 7.07E-02 | 2.24E-01 | 4.79E-09 | 6.94E-03 | 6.51E-05 | 1.30E-01 | 1.00E+00 | 3.52E-02 | 9.56E-02 |
| Meta_AMS3010 | 4.15E-37 | 7.81E-40 | 6.77E-39 | 3.07E-36 | 2.02E-30 | 1.63E-34 | 6.77E-39 | 5.27E-32 | 7.25E-47 | 5.54E-38 | 6.72E-01 | 7.92E-01 | 4.58E-01 | 2.78E-03 | 2.78E-03 | 5.03E-02 | 9.63E-01 | 3.52E-02 | 1.00E+00 | 9.63E-01 |
| Meta_AMS3011 | 9.40E-43 | 8.49E-46 | 7.46E-45 | 8.71E-42 | 2.23E-35 | 7.81E-40 | 7.46E-45 | 4.15E-37 | 6.01E-48 | 8.71E-42 | 3.69E-01 | 4.58E-01 | 5.59E-01 | 2.78E-03 | 3.65E-04 | 1.30E-01 | 5.59E-01 | 9.56E-02 | 9.63E-01 | 1.00E+00 |

Comparison of trees using the Kuhner & Felsenstein, 1994 method

| | ARS3001 | ARS3002 | ARS3003 | ARS3004 | ARS3005 | ARS3006 | ARS3007 | ARS3008 | ARS3009 | ARS3010 | Meta_AMS3002 | Meta_AMS3003 | Meta_AMS3004 | Meta_AMS3005 | Meta_AMS3006 | Meta_AMS3007 | Meta_AMS3008 | Meta_AMS3009 | Meta_AMS3010 | Meta_AMS3011 |
|---|---|---|---|---|---|---|---|---|---|---|---|---|---|---|---|---|---|---|---|---|
| ARS3001 | 1.00E+00 | 5.38E-02 | 7.57E-01 | 3.67E-01 | 1.76E-01 | 7.57E-01 | 7.57E-01 | 7.57E-01 | 1.31E-01 | 1.74E-02 | 9.87E-22 | 5.75E-14 | 1.70E-11 | 1.54E-13 | 3.27E-09 | 2.61E-10 | 5.75E-14 | 1.65E-08 | 2.44E-15 | 2.61E-10 |
| ARS3002 | 5.38E-02 | 1.00E+00 | 1.31E-01 | 3.67E-01 | 3.67E-01 | 1.18E-02 | 7.27E-02 | 5.38E-02 | 1.31E-01 | 1.18E-02 | 3.15E-12 | 3.50E-08 | 6.69E-07 | 3.50E-08 | 1.99E-05 | 5.41E-06 | 1.65E-08 | 1.24E-04 | 3.50E-08 | 6.69E-07 |
| ARS3003 | 7.57E-01 | 1.31E-01 | 1.00E+00 | 8.52E-01 | 1.76E-01 | 2.95E-01 | 6.68E-01 | 3.67E-01 | 2.58E-02 | 1.74E-02 | 4.89E-20 | 7.36E-12 | 7.36E-09 | 1.70E-11 | 3.28E-07 | 1.65E-08 | 4.20E-11 | 6.69E-07 | 1.54E-13 | 7.36E-09 |
| ARS3004 | 3.67E-01 | 3.67E-01 | 8.52E-01 | 1.00E+00 | 7.57E-01 | 2.95E-01 | 7.57E-01 | 4.58E-01 | 9.96E-02 | 1.31E-01 | 2.52E-17 | 1.70E-11 | 1.56E-07 | 1.07E-10 | 1.99E-05 | 1.56E-07 | 7.36E-12 | 1.05E-05 | 7.36E-12 | 1.56E-07 |
| ARS3005 | 1.76E-01 | 3.67E-01 | 1.76E-01 | 7.57E-01 | 1.00E+00 | 7.27E-02 | 6.68E-01 | 7.27E-02 | 7.72E-03 | 3.67E-01 | 8.23E-17 | 6.15E-10 | 7.61E-08 | 6.15E-10 | 5.41E-06 | 7.61E-08 | 1.70E-11 | 5.41E-06 | 4.20E-11 | 1.56E-07 |
| ARS3006 | 7.57E-01 | 1.18E-02 | 2.95E-01 | 2.95E-01 | 7.27E-02 | 1.00E+00 | 5.62E-01 | 8.52E-01 | 3.77E-02 | 4.94E-03 | 5.54E-24 | 8.43E-16 | 3.15E-12 | 8.43E-16 | 3.27E-09 | 7.36E-12 | 2.52E-17 | 3.27E-09 | 2.52E-17 | 7.36E-12 |
| ARS3007 | 7.57E-01 | 7.27E-02 | 6.68E-01 | 7.57E-01 | 6.68E-01 | 5.62E-01 | 1.00E+00 | 4.58E-01 | 7.27E-02 | 9.96E-02 | 3.31E-21 | 2.15E-14 | 6.15E-10 | 1.22E-12 | 6.69E-07 | 1.43E-09 | 2.15E-14 | 3.28E-07 | 2.44E-15 | 3.27E-09 |
| ARS3008 | 7.57E-01 | 5.38E-02 | 3.67E-01 | 4.58E-01 | 7.27E-02 | 8.52E-01 | 4.58E-01 | 1.00E+00 | 1.31E-01 | 4.94E-03 | 9.87E-22 | 1.54E-13 | 1.07E-10 | 3.15E-12 | 7.36E-09 | 1.43E-09 | 5.75E-14 | 7.61E-08 | 7.53E-15 | 1.43E-09 |
| ARS3009 | 1.31E-01 | 1.31E-01 | 2.58E-02 | 9.96E-02 | 7.72E-03 | 3.77E-02 | 7.27E-02 | 1.31E-01 | 1.00E+00 | 1.24E-04 | 4.46E-13 | 1.70E-11 | 1.43E-09 | 1.22E-12 | 3.50E-08 | 7.36E-09 | 1.70E-11 | 1.56E-07 | 1.70E-11 | 6.15E-10 |
| ARS3010 | 1.74E-02 | 1.18E-02 | 1.74E-02 | 1.31E-01 | 3.67E-01 | 4.94E-03 | 9.96E-02 | 4.94E-03 | 1.24E-04 | 1.00E+00 | 2.52E-17 | 4.20E-11 | 3.50E-08 | 2.61E-10 | 1.24E-04 | 3.28E-07 | 4.20E-11 | 1.05E-05 | 7.36E-12 | 2.80E-06 |
| Meta_AMS3002 | 9.87E-22 | 3.15E-12 | 4.89E-20 | 2.52E-17 | 8.23E-17 | 5.54E-24 | 3.31E-21 | 9.87E-22 | 4.46E-13 | 2.52E-17 | 1.00E+00 | 7.27E-02 | 5.41E-06 | 3.92E-04 | 2.80E-06 | 3.76E-05 | 3.77E-02 | 3.50E-08 | 1.24E-04 | 1.56E-07 |
| Meta_AMS3003 | 5.75E-14 | 3.50E-08 | 7.36E-12 | 1.70E-11 | 6.15E-10 | 8.43E-16 | 2.15E-14 | 1.54E-13 | 1.70E-11 | 4.20E-11 | 7.27E-02 | 1.00E+00 | 1.74E-02 | 2.95E-01 | 1.74E-02 | 9.96E-02 | 7.57E-01 | 1.14E-03 | 1.31E-01 | 4.94E-03 |
| Meta_AMS3004 | 1.70E-11 | 6.69E-07 | 7.36E-09 | 1.56E-07 | 7.61E-08 | 3.15E-12 | 6.15E-10 | 1.07E-10 | 1.43E-09 | 3.50E-08 | 5.41E-06 | 1.74E-02 | 1.00E+00 | 2.95E-01 | 2.32E-01 | 1.00E+00 | 2.58E-02 | 7.57E-01 | 1.31E-01 | 9.42E-01 |
| Meta_AMS3005 | 1.54E-13 | 3.50E-08 | 1.70E-11 | 1.07E-10 | 6.15E-10 | 8.43E-16 | 1.22E-12 | 3.15E-12 | 1.22E-12 | 2.61E-10 | 3.92E-04 | 2.95E-01 | 2.95E-01 | 1.00E+00 | 5.38E-02 | 4.58E-01 | 3.67E-01 | 7.27E-02 | 5.62E-01 | 2.32E-01 |
| Meta_AMS3006 | 3.27E-09 | 1.99E-05 | 3.28E-07 | 1.99E-05 | 5.41E-06 | 3.27E-09 | 6.69E-07 | 7.36E-09 | 3.50E-08 | 1.24E-04 | 2.80E-06 | 1.74E-02 | 2.32E-01 | 5.38E-02 | 1.00E+00 | 2.95E-01 | 1.18E-02 | 8.52E-01 | 7.72E-03 | 6.68E-01 |
| Meta_AMS3007 | 2.61E-10 | 5.41E-06 | 1.65E-08 | 1.56E-07 | 7.61E-08 | 7.36E-12 | 1.43E-09 | 1.43E-09 | 7.36E-09 | 3.28E-07 | 3.76E-05 | 9.96E-02 | 1.00E+00 | 4.58E-01 | 2.95E-01 | 1.00E+00 | 7.27E-02 | 2.95E-01 | 1.31E-01 | 6.68E-01 |
| Meta_AMS3008 | 5.75E-14 | 1.65E-08 | 4.20E-11 | 7.36E-12 | 1.70E-11 | 2.52E-17 | 2.15E-14 | 5.75E-14 | 1.70E-11 | 4.20E-11 | 3.77E-02 | 7.57E-01 | 2.58E-02 | 3.67E-01 | 1.18E-02 | 7.27E-02 | 1.00E+00 | 1.14E-03 | 9.96E-02 | 4.94E-03 |
| Meta_AMS3009 | 1.65E-08 | 1.24E-04 | 6.69E-07 | 1.05E-05 | 5.41E-06 | 3.27E-09 | 3.28E-07 | 7.61E-08 | 1.56E-07 | 1.05E-05 | 3.50E-08 | 1.14E-03 | 7.57E-01 | 7.27E-02 | 8.52E-01 | 2.95E-01 | 1.14E-03 | 1.00E+00 | 1.74E-02 | 8.52E-01 |
| Meta_AMS3010 | 2.44E-15 | 3.50E-08 | 1.54E-13 | 7.36E-12 | 4.20E-11 | 2.52E-17 | 2.44E-15 | 7.53E-15 | 1.70E-11 | 7.36E-12 | 1.24E-04 | 1.31E-01 | 1.31E-01 | 5.62E-01 | 7.72E-03 | 1.31E-01 | 9.96E-02 | 1.74E-02 | 1.00E+00 | 7.27E-02 |
| Meta_AMS3011 | 2.61E-10 | 6.69E-07 | 7.36E-09 | 1.56E-07 | 1.56E-07 | 7.36E-12 | 3.27E-09 | 1.43E-09 | 6.15E-10 | 2.80E-06 | 1.56E-07 | 4.94E-03 | 9.42E-01 | 2.32E-01 | 6.68E-01 | 6.68E-01 | 4.94E-03 | 8.52E-01 | 7.27E-02 | 1.00E+00 |

**Supplementary Table S4D:** Pairwise FDR corrected p-values of the Kolmogorow-Smirnow test on the distances between phylogenetic trees for ribosomal proteins in 10 samples of bacterial RefSeq genomes vs 10 samples of CPR MAGs (associated with Supplementary Fig. S7d)

Comparison of trees using the Nye et al., 2006 method

| | BRS3001 | BRS3002 | BRS3003 | BRS3004 | BRS3005 | BRS3006 | BRS3007 | BRS3008 | BRS3009 | BRS3010 | Meta_CPR3001 | Meta_CPR3002 | Meta_CPR3003 | Meta_CPR3004 | Meta_CPR3005 | Meta_CPR3006 | Meta_CPR3007 | Meta_CPR3008 | Meta_CPR3009 | Meta_CPR3010 |
|---|---|---|---|---|---|---|---|---|---|---|---|---|---|---|---|---|---|---|---|---|
| BRS3001 | 1.00E+00 | 1.18E-01 | 1.58E-01 | 2.49E-08 | 2.65E-01 | 5.83E-05 | 1.03E-04 | 3.99E-03 | 1.58E-01 | 3.11E-02 | 6.88E-54 | 6.37E-46 | 9.20E-42 | 9.08E-53 | 7.85E-41 | 4.51E-48 | 3.56E-55 | 2.81E-35 | 6.36E-26 | 9.20E-42 |
| BRS3002 | 1.18E-01 | 1.00E+00 | 4.49E-02 | 2.00E-09 | 9.24E-04 | 4.57E-06 | 1.69E-05 | 1.03E-04 | 3.99E-03 | 3.17E-05 | 9.08E-53 | 1.48E-51 | 7.46E-45 | 3.56E-55 | 6.37E-46 | 2.11E-50 | 3.67E-56 | 5.88E-38 | 1.46E-29 | 8.60E-44 |
| BRS3003 | 1.58E-01 | 4.49E-02 | 1.00E+00 | 1.03E-04 | 2.07E-01 | 9.73E-03 | 2.15E-02 | 8.79E-02 | 2.65E-01 | 1.18E-01 | 5.88E-38 | 3.86E-31 | 4.77E-28 | 4.78E-37 | 2.53E-27 | 2.00E-34 | 7.85E-41 | 3.22E-23 | 3.49E-16 | 4.77E-28 |
| BRS3004 | 2.49E-08 | 2.00E-09 | 1.03E-04 | 1.00E+00 | 5.44E-07 | 2.65E-01 | 2.07E-01 | 2.52E-03 | 3.21E-04 | 6.31E-03 | 7.46E-45 | 1.37E-33 | 1.37E-33 | 9.11E-43 | 9.24E-33 | 7.85E-41 | 6.04E-47 | 3.09E-25 | 2.36E-18 | 1.37E-33 |
| BRS3005 | 2.65E-01 | 9.24E-04 | 2.07E-01 | 5.44E-07 | 1.00E+00 | 1.84E-04 | 9.24E-04 | 3.11E-02 | 5.23E-01 | 8.79E-02 | 2.97E-49 | 6.37E-46 | 7.30E-40 | 2.97E-49 | 7.85E-41 | 6.37E-46 | 1.48E-51 | 2.81E-35 | 3.09E-25 | 7.85E-41 |
| BRS3006 | 5.83E-05 | 4.57E-06 | 9.73E-03 | 2.65E-01 | 1.84E-04 | 1.00E+00 | 7.41E-01 | 1.18E-01 | 9.24E-04 | 8.79E-02 | 3.81E-36 | 2.53E-27 | 6.36E-26 | 4.78E-37 | 1.27E-26 | 2.00E-34 | 7.30E-40 | 2.60E-21 | 3.16E-13 | 1.27E-26 |
| BRS3007 | 1.03E-04 | 1.69E-05 | 2.15E-02 | 2.07E-01 | 9.24E-04 | 7.41E-01 | 1.00E+00 | 4.49E-02 | 2.15E-02 | 2.07E-01 | 4.78E-37 | 2.53E-27 | 1.27E-26 | 3.81E-36 | 6.36E-26 | 9.24E-33 | 7.85E-41 | 2.60E-21 | 3.59E-14 | 1.27E-26 |
| BRS3008 | 3.99E-03 | 1.03E-04 | 8.79E-02 | 2.52E-03 | 3.11E-02 | 1.18E-01 | 4.49E-02 | 1.00E+00 | 1.18E-01 | 4.25E-01 | 9.11E-43 | 6.17E-32 | 3.86E-31 | 9.20E-42 | 3.86E-31 | 7.30E-40 | 7.46E-45 | 1.49E-24 | 3.01E-17 | 9.24E-33 |
| BRS3009 | 1.58E-01 | 3.99E-03 | 2.65E-01 | 3.21E-04 | 5.23E-01 | 9.24E-04 | 2.15E-02 | 1.18E-01 | 1.00E+00 | 2.65E-01 | 2.11E-50 | 8.60E-44 | 7.85E-41 | 2.11E-50 | 7.85E-41 | 6.37E-46 | 9.08E-53 | 1.37E-33 | 3.09E-25 | 6.77E-39 |
| BRS3010 | 3.11E-02 | 3.17E-05 | 1.18E-01 | 6.31E-03 | 8.79E-02 | 8.79E-02 | 2.07E-01 | 4.25E-01 | 2.65E-01 | 1.00E+00 | 6.04E-47 | 2.81E-35 | 2.00E-34 | 6.37E-46 | 2.00E-34 | 9.11E-43 | 2.97E-49 | 1.27E-26 | 1.68E-19 | 2.81E-35 |
| Meta_CPR3001 | 6.88E-54 | 9.08E-53 | 5.88E-38 | 7.46E-45 | 2.97E-49 | 3.81E-36 | 4.78E-37 | 9.11E-43 | 2.11E-50 | 6.04E-47 | 1.00E+00 | 5.51E-08 | 2.49E-08 | 6.29E-01 | 1.69E-05 | 3.11E-02 | 8.48E-01 | 1.30E-10 | 5.44E-07 | 1.13E-06 |
| Meta_CPR3002 | 6.37E-46 | 1.48E-51 | 3.86E-31 | 1.37E-33 | 6.37E-46 | 2.53E-27 | 2.53E-27 | 6.17E-32 | 8.60E-44 | 2.81E-35 | 5.51E-08 | 1.00E+00 | 9.37E-01 | 3.17E-05 | 1.58E-01 | 1.54E-03 | 2.49E-08 | 4.25E-01 | 1.46E-02 | 6.29E-01 |
| Meta_CPR3003 | 9.20E-42 | 7.46E-45 | 4.77E-28 | 1.37E-33 | 7.30E-40 | 6.36E-26 | 1.27E-26 | 3.86E-31 | 7.85E-41 | 2.00E-34 | 2.49E-08 | 9.37E-01 | 1.00E+00 | 8.96E-06 | 6.40E-02 | 5.53E-04 | 4.74E-09 | 4.25E-01 | 3.11E-02 | 2.65E-01 |
| Meta_CPR3004 | 9.08E-53 | 3.56E-55 | 4.78E-37 | 9.11E-43 | 2.97E-49 | 4.78E-37 | 3.81E-36 | 9.20E-42 | 2.11E-50 | 6.37E-46 | 6.29E-01 | 3.17E-05 | 8.96E-06 | 1.00E+00 | 3.21E-04 | 2.15E-02 | 5.23E-01 | 1.21E-07 | 1.69E-05 | 1.03E-04 |
| Meta_CPR3005 | 7.85E-41 | 6.37E-46 | 2.53E-27 | 9.24E-33 | 7.85E-41 | 1.27E-26 | 6.36E-26 | 3.86E-31 | 7.85E-41 | 2.00E-34 | 1.69E-05 | 1.58E-01 | 6.40E-02 | 3.21E-04 | 1.00E+00 | 8.79E-02 | 4.57E-06 | 1.46E-02 | 1.46E-02 | 4.25E-01 |
| Meta_CPR3006 | 4.51E-48 | 2.11E-50 | 2.00E-34 | 7.85E-41 | 6.37E-46 | 2.00E-34 | 9.24E-33 | 7.30E-40 | 6.37E-46 | 9.11E-43 | 3.11E-02 | 1.54E-03 | 5.53E-04 | 2.15E-02 | 8.79E-02 | 1.00E+00 | 3.99E-03 | 5.83E-05 | 1.03E-04 | 2.15E-02 |
| Meta_CPR3007 | 3.56E-55 | 3.67E-56 | 7.85E-41 | 6.04E-47 | 1.48E-51 | 7.30E-40 | 7.85E-41 | 7.46E-45 | 9.08E-53 | 2.97E-49 | 8.48E-01 | 2.49E-08 | 4.74E-09 | 5.23E-01 | 4.57E-06 | 3.99E-03 | 1.00E+00 | 5.07E-11 | 5.51E-08 | 5.44E-07 |
| Meta_CPR3008 | 2.81E-35 | 5.88E-38 | 3.22E-23 | 3.09E-25 | 2.81E-35 | 2.60E-21 | 2.60E-21 | 1.49E-24 | 1.37E-33 | 1.27E-26 | 1.30E-10 | 4.25E-01 | 4.25E-01 | 1.21E-07 | 1.46E-02 | 5.83E-05 | 5.07E-11 | 1.00E+00 | 2.07E-01 | 1.18E-01 |
| Meta_CPR3009 | 6.36E-26 | 1.46E-29 | 3.49E-16 | 2.36E-18 | 3.09E-25 | 3.16E-13 | 3.59E-14 | 3.01E-17 | 3.09E-25 | 1.68E-19 | 5.44E-07 | 1.46E-02 | 3.11E-02 | 1.69E-05 | 1.46E-02 | 1.03E-04 | 5.51E-08 | 2.07E-01 | 1.00E+00 | 9.73E-03 |
| Meta_CPR3010 | 9.20E-42 | 8.60E-44 | 4.77E-28 | 1.37E-33 | 7.85E-41 | 1.27E-26 | 1.27E-26 | 9.24E-33 | 6.77E-39 | 2.81E-35 | 1.13E-06 | 6.29E-01 | 2.65E-01 | 1.03E-04 | 4.25E-01 | 2.15E-02 | 5.44E-07 | 1.18E-01 | 9.73E-03 | 1.00E+00 |

Comparison of trees using the Williams & Clifford, 1971 method

| | BRS3001 | BRS3002 | BRS3003 | BRS3004 | BRS3005 | BRS3006 | BRS3007 | BRS3008 | BRS3009 | BRS3010 | Meta_CPR3001 | Meta_CPR3002 | Meta_CPR3003 | Meta_CPR3004 | Meta_CPR3005 | Meta_CPR3006 | Meta_CPR3007 | Meta_CPR3008 | Meta_CPR3009 | Meta_CPR3010 |
|---|---|---|---|---|---|---|---|---|---|---|---|---|---|---|---|---|---|---|---|---|
| BRS3001 | 1.00E+00 | 2.79E-01 | 6.43E-01 | 9.06E-10 | 2.18E-01 | 1.85E-05 | 5.98E-04 | 6.74E-03 | 2.72E-03 | 1.12E-04 | 3.02E-32 | 1.08E-30 | 2.59E-27 | 5.79E-28 | 8.22E-23 | 1.63E-33 | 3.02E-32 | 2.14E-23 | 4.87E-17 | 5.79E-28 |
| BRS3002 | 2.79E-01 | 1.00E+00 | 5.40E-01 | 3.36E-05 | 3.52E-01 | 6.74E-03 | 1.23E-01 | 1.66E-01 | 3.52E-01 | 6.60E-02 | 2.59E-27 | 8.22E-23 | 8.22E-23 | 2.14E-23 | 3.39E-19 | 3.56E-29 | 2.59E-27 | 4.87E-17 | 8.59E-12 | 8.22E-23 |
| BRS3003 | 6.43E-01 | 5.40E-01 | 1.00E+00 | 6.21E-07 | 2.79E-01 | 5.98E-04 | 6.74E-03 | 6.60E-02 | 9.07E-02 | 6.74E-03 | 5.22E-24 | 5.22E-24 | 1.37E-21 | 3.31E-22 | 4.27E-18 | 1.25E-26 | 2.77E-25 | 1.19E-18 | 3.22E-12 | 3.31E-22 |
| BRS3004 | 9.06E-10 | 3.36E-05 | 6.21E-07 | 1.00E+00 | 6.23E-05 | 2.79E-01 | 9.07E-02 | 1.54E-02 | 1.85E-05 | 2.01E-04 | 5.75E-11 | 6.26E-08 | 3.22E-12 | 4.73E-14 | 2.61E-06 | 4.87E-17 | 3.22E-12 | 1.85E-05 | 1.54E-02 | 5.19E-09 |
| BRS3005 | 2.18E-01 | 3.52E-01 | 2.79E-01 | 6.23E-05 | 1.00E+00 | 4.36E-03 | 6.60E-02 | 1.23E-01 | 2.18E-01 | 6.60E-02 | 1.25E-26 | 1.26E-24 | 8.22E-23 | 2.14E-23 | 1.19E-18 | 1.55E-28 | 5.79E-28 | 1.44E-17 | 1.18E-12 | 1.26E-24 |
| BRS3006 | 1.85E-05 | 6.74E-03 | 5.98E-04 | 2.79E-01 | 4.36E-03 | 1.00E+00 | 7.53E-01 | 4.39E-01 | 1.03E-02 | 3.33E-02 | 1.58E-14 | 2.27E-11 | 4.73E-14 | 5.34E-16 | 2.19E-09 | 9.03E-20 | 1.64E-15 | 2.98E-07 | 3.53E-04 | 8.59E-12 |
| BRS3007 | 5.98E-04 | 1.23E-01 | 6.74E-03 | 9.07E-02 | 6.60E-02 | 7.53E-01 | 1.00E+00 | 1.00E+00 | 1.03E-02 | 9.07E-02 | 1.63E-16 | 1.64E-15 | 1.58E-14 | 1.58E-14 | 5.75E-11 | 3.39E-19 | 1.44E-17 | 1.46E-10 | 2.61E-06 | 1.41E-13 |
| BRS3008 | 6.74E-03 | 1.66E-01 | 6.60E-02 | 1.54E-02 | 1.23E-01 | 4.39E-01 | 1.00E+00 | 1.00E+00 | 6.60E-02 | 2.18E-01 | 1.19E-18 | 1.63E-16 | 1.64E-15 | 5.34E-16 | 8.59E-12 | 2.35E-20 | 9.03E-20 | 5.75E-11 | 6.21E-07 | 1.64E-15 |
| BRS3009 | 2.72E-03 | 3.52E-01 | 9.07E-02 | 1.85E-05 | 2.18E-01 | 1.03E-02 | 1.03E-02 | 6.60E-02 | 1.00E+00 | 2.29E-02 | 2.77E-25 | 1.44E-17 | 8.22E-23 | 3.31E-22 | 1.19E-18 | 5.79E-28 | 2.77E-25 | 5.21E-15 | 1.46E-10 | 1.37E-21 |
| BRS3010 | 1.12E-04 | 6.60E-02 | 6.74E-03 | 2.01E-04 | 6.60E-02 | 3.33E-02 | 9.07E-02 | 2.18E-01 | 2.29E-02 | 1.00E+00 | 5.22E-24 | 1.44E-17 | 2.14E-23 | 5.22E-24 | 4.27E-18 | 1.55E-28 | 6.29E-26 | 1.64E-15 | 5.75E-11 | 3.31E-22 |
| Meta_CPR3001 | 3.02E-32 | 2.59E-27 | 5.22E-24 | 5.75E-11 | 1.25E-26 | 1.58E-14 | 1.63E-16 | 1.19E-18 | 2.77E-25 | 5.22E-24 | 1.00E+00 | 2.79E-01 | 6.43E-01 | 1.01E-03 | 1.23E-01 | 5.98E-04 | 7.53E-01 | 6.60E-02 | 1.85E-05 | 8.52E-01 |
| Meta_CPR3002 | 1.08E-30 | 8.22E-23 | 5.22E-24 | 6.26E-08 | 1.26E-24 | 2.27E-11 | 1.64E-15 | 1.63E-16 | 1.44E-17 | 1.44E-17 | 2.79E-01 | 1.00E+00 | 2.79E-01 | 3.36E-05 | 3.52E-01 | 3.36E-05 | 9.07E-02 | 4.39E-01 | 1.54E-02 | 8.52E-01 |
| Meta_CPR3003 | 2.59E-27 | 8.22E-23 | 1.37E-21 | 3.22E-12 | 8.22E-23 | 4.73E-14 | 1.58E-14 | 1.64E-15 | 8.22E-23 | 2.14E-23 | 6.43E-01 | 2.79E-01 | 1.00E+00 | 2.72E-03 | 1.66E-01 | 1.67E-03 | 6.43E-01 | 6.60E-02 | 3.36E-05 | 8.52E-01 |
| Meta_CPR3004 | 5.79E-28 | 2.14E-23 | 3.31E-22 | 4.73E-14 | 2.14E-23 | 5.34E-16 | 1.58E-14 | 5.34E-16 | 3.31E-22 | 5.22E-24 | 1.01E-03 | 3.36E-05 | 2.72E-03 | 1.00E+00 | 1.85E-05 | 6.43E-01 | 1.03E-02 | 9.98E-06 | 9.06E-10 | 1.12E-04 |
| Meta_CPR3005 | 8.22E-23 | 3.39E-19 | 4.27E-18 | 2.61E-06 | 1.19E-18 | 2.19E-09 | 5.75E-11 | 8.59E-12 | 1.19E-18 | 4.27E-18 | 1.23E-01 | 3.52E-01 | 1.66E-01 | 1.85E-05 | 1.00E+00 | 9.98E-06 | 3.33E-02 | 1.00E+00 | 3.33E-02 | 5.40E-01 |
| Meta_CPR3006 | 1.63E-33 | 3.56E-29 | 1.25E-26 | 4.87E-17 | 1.55E-28 | 9.03E-20 | 3.39E-19 | 2.35E-20 | 5.79E-28 | 1.55E-28 | 5.98E-04 | 3.36E-05 | 1.67E-03 | 6.43E-01 | 9.98E-06 | 1.00E+00 | 1.54E-02 | 5.15E-06 | 2.27E-11 | 3.36E-05 |
| Meta_CPR3007 | 3.02E-32 | 2.59E-27 | 2.77E-25 | 3.22E-12 | 5.79E-28 | 1.64E-15 | 1.44E-17 | 9.03E-20 | 2.77E-25 | 6.29E-26 | 7.53E-01 | 9.07E-02 | 6.43E-01 | 1.03E-02 | 3.33E-02 | 1.54E-02 | 1.00E+00 | 2.29E-02 | 5.15E-06 | 4.39E-01 |
| Meta_CPR3008 | 2.14E-23 | 4.87E-17 | 1.19E-18 | 1.85E-05 | 1.44E-17 | 2.98E-07 | 1.46E-10 | 5.75E-11 | 5.21E-15 | 1.64E-15 | 6.60E-02 | 4.39E-01 | 6.60E-02 | 9.98E-06 | 1.00E+00 | 5.15E-06 | 2.29E-02 | 1.00E+00 | 4.83E-02 | 3.52E-01 |
| Meta_CPR3009 | 4.87E-17 | 8.59E-12 | 3.22E-12 | 1.54E-02 | 1.18E-12 | 3.53E-04 | 2.61E-06 | 6.21E-07 | 1.46E-10 | 5.75E-11 | 1.85E-05 | 1.54E-02 | 3.36E-05 | 9.06E-10 | 3.33E-02 | 2.27E-11 | 5.15E-06 | 4.83E-02 | 1.00E+00 | 1.67E-03 |
| Meta_CPR3010 | 5.79E-28 | 8.22E-23 | 3.31E-22 | 5.19E-09 | 1.26E-24 | 8.59E-12 | 1.41E-13 | 1.64E-15 | 1.37E-21 | 3.31E-22 | 8.52E-01 | 8.52E-01 | 8.52E-01 | 1.12E-04 | 5.40E-01 | 3.36E-05 | 4.39E-01 | 3.52E-01 | 1.67E-03 | 1.00E+00 |

Comparison of trees using the Robinson-Foulds method

| | BRS3001 | BRS3002 | BRS3003 | BRS3004 | BRS3005 | BRS3006 | BRS3007 | BRS3008 | BRS3009 | BRS3010 | Meta_CPR3001 | Meta_CPR3002 | Meta_CPR3003 | Meta_CPR3004 | Meta_CPR3005 | Meta_CPR3006 | Meta_CPR3007 | Meta_CPR3008 | Meta_CPR3009 | Meta_CPR3010 |
|---|---|---|---|---|---|---|---|---|---|---|---|---|---|---|---|---|---|---|---|---|
| BRS3001 | 1.00E+00 | 2.17E-01 | 4.57E-02 | 3.12E-05 | 1.00E+00 | 3.31E-04 | 1.64E-01 | 3.22E-02 | 7.74E-01 | 3.22E-02 | 1.06E-52 | 7.28E-46 | 7.24E-39 | 1.33E-51 | 1.02E-40 | 2.67E-55 | 1.83E-56 | 3.14E-35 | 2.60E-27 | 7.24E-39 |
| BRS3002 | 2.17E-01 | 1.00E+00 | 3.22E-02 | 4.40E-09 | 6.52E-02 | 2.23E-06 | 4.57E-02 | 4.11E-03 | 2.27E-02 | 5.74E-05 | 4.81E-48 | 9.33E-40 | 2.21E-34 | 6.39E-47 | 3.14E-35 | 4.81E-48 | 1.33E-51 | 6.75E-32 | 3.31E-25 | 1.56E-33 |
| BRS3003 | 4.57E-02 | 3.22E-02 | 1.00E+00 | 9.59E-04 | 3.22E-02 | 3.22E-02 | 1.00E+00 | 8.80E-01 | 2.17E-01 | 3.63E-01 | 9.33E-40 | 2.65E-30 | 2.60E-27 | 1.02E-40 | 2.60E-27 | 6.26E-38 | 7.28E-46 | 3.29E-23 | 3.49E-16 | 1.58E-24 |
| BRS3004 | 3.12E-05 | 4.40E-09 | 9.59E-04 | 1.00E+00 | 1.65E-05 | 6.64E-01 | 2.59E-03 | 4.57E-02 | 9.59E-04 | 8.96E-02 | 1.19E-41 | 1.06E-32 | 1.57E-29 | 1.02E-40 | 1.57E-29 | 1.02E-40 | 7.28E-46 | 3.31E-25 | 1.04E-16 | 1.34E-26 |
| BRS3005 | 1.00E+00 | 6.52E-02 | 3.22E-02 | 1.65E-05 | 1.00E+00 | 1.65E-05 | 3.22E-02 | 4.11E-03 | 9.73E-01 | 2.27E-02 | 1.06E-52 | 7.28E-46 | 7.24E-39 | 1.33E-51 | 1.02E-40 | 2.67E-55 | 1.83E-56 | 7.24E-39 | 4.27E-31 | 7.24E-39 |
| BRS3006 | 3.31E-04 | 2.23E-06 | 3.22E-02 | 6.64E-01 | 1.65E-05 | 1.00E+00 | 1.21E-01 | 5.53E-01 | 2.59E-03 | 4.52E-01 | 5.29E-37 | 8.98E-29 | 6.75E-26 | 6.26E-38 | 6.75E-26 | 4.20E-36 | 1.12E-42 | 6.21E-22 | 1.16E-14 | 3.29E-23 |
| BRS3007 | 1.64E-01 | 4.57E-02 | 1.00E+00 | 2.59E-03 | 3.22E-02 | 1.21E-01 | 1.00E+00 | 1.00E+00 | 4.52E-01 | 1.21E-01 | 9.33E-40 | 6.75E-32 | 4.97E-28 | 7.24E-39 | 4.97E-28 | 7.24E-39 | 9.98E-44 | 7.39E-24 | 8.58E-18 | 1.34E-26 |
| BRS3008 | 3.22E-02 | 4.11E-03 | 8.80E-01 | 4.57E-02 | 4.11E-03 | 5.53E-01 | 1.00E+00 | 1.00E+00 | 1.21E-01 | 5.53E-01 | 1.56E-33 | 1.34E-26 | 1.46E-22 | 2.21E-34 | 3.29E-23 | 1.06E-32 | 7.24E-39 | 1.69E-19 | 3.55E-14 | 6.21E-22 |
| BRS3009 | 7.74E-01 | 2.27E-02 | 2.17E-01 | 9.59E-04 | 9.73E-01 | 2.59E-03 | 4.52E-01 | 1.21E-01 | 1.00E+00 | 2.84E-01 | 4.81E-48 | 1.02E-40 | 2.21E-34 | 6.39E-47 | 4.20E-36 | 4.04E-49 | 1.33E-51 | 4.27E-31 | 1.58E-24 | 2.21E-34 |
| BRS3010 | 3.22E-02 | 5.74E-05 | 3.63E-01 | 8.96E-02 | 2.27E-02 | 4.52E-01 | 1.21E-01 | 5.53E-01 | 2.84E-01 | 1.00E+00 | 9.15E-45 | 3.14E-35 | 6.75E-32 | 9.98E-44 | 6.75E-32 | 9.98E-44 | 4.81E-48 | 2.60E-27 | 2.38E-18 | 8.98E-29 |
| Meta_CPR3001 | 1.06E-52 | 4.81E-48 | 9.33E-40 | 1.19E-41 | 1.06E-52 | 5.29E-37 | 9.33E-40 | 1.56E-33 | 4.81E-48 | 9.15E-45 | 1.00E+00 | 1.85E-09 | 1.01E-08 | 1.00E+00 | 1.65E-05 | 5.70E-04 | 1.00E+00 | 1.89E-11 | 7.88E-10 | 1.85E-09 |
| Meta_CPR3002 | 7.28E-46 | 9.33E-40 | 2.65E-30 | 1.06E-32 | 7.28E-46 | 8.98E-29 | 6.75E-32 | 1.34E-26 | 1.02E-40 | 3.14E-35 | 1.85E-09 | 1.00E+00 | 8.80E-01 | 3.19E-10 | 6.64E-01 | 4.57E-02 | 1.01E-08 | 2.27E-02 | 4.11E-03 | 7.74E-01 |
| Meta_CPR3003 | 7.24E-39 | 2.21E-34 | 2.60E-27 | 1.57E-29 | 7.24E-39 | 6.75E-26 | 4.97E-28 | 1.46E-22 | 2.21E-34 | 6.75E-32 | 1.01E-08 | 8.80E-01 | 1.00E+00 | 1.85E-09 | 7.74E-01 | 3.22E-02 | 5.18E-08 | 1.21E-01 | 8.96E-02 | 1.00E+00 |
| Meta_CPR3004 | 1.33E-51 | 6.39E-47 | 1.02E-40 | 1.02E-40 | 1.33E-51 | 6.26E-38 | 7.24E-39 | 2.21E-34 | 6.39E-47 | 9.98E-44 | 1.00E+00 | 3.19E-10 | 1.85E-09 | 1.00E+00 | 4.42E-06 | 1.88E-04 | 1.00E+00 | 2.55E-12 | 1.28E-10 | 3.19E-10 |
| Meta_CPR3005 | 1.02E-40 | 3.14E-35 | 2.60E-27 | 1.57E-29 | 1.02E-40 | 6.75E-26 | 4.97E-28 | 3.29E-23 | 4.20E-36 | 6.75E-32 | 1.65E-05 | 6.64E-01 | 7.74E-01 | 4.42E-06 | 1.00E+00 | 3.63E-01 | 5.74E-05 | 1.54E-02 | 6.45E-03 | 4.52E-01 |
| Meta_CPR3006 | 2.67E-55 | 4.81E-48 | 6.26E-38 | 1.02E-40 | 2.67E-55 | 4.20E-36 | 7.24E-39 | 1.06E-32 | 4.04E-49 | 9.98E-44 | 5.70E-04 | 4.57E-02 | 3.22E-02 | 1.88E-04 | 3.63E-01 | 1.00E+00 | 1.60E-03 | 4.42E-06 | 2.23E-06 | 6.45E-03 |
| Meta_CPR3007 | 1.83E-56 | 1.33E-51 | 7.28E-46 | 7.28E-46 | 1.83E-56 | 1.12E-42 | 9.98E-44 | 7.24E-39 | 1.33E-51 | 4.81E-48 | 1.00E+00 | 1.01E-08 | 5.18E-08 | 1.00E+00 | 5.74E-05 | 1.60E-03 | 1.00E+00 | 1.28E-10 | 1.85E-09 | 1.01E-08 |
| Meta_CPR3008 | 3.14E-35 | 6.75E-32 | 3.29E-23 | 3.31E-25 | 7.24E-39 | 6.21E-22 | 7.39E-24 | 1.69E-19 | 4.27E-31 | 2.60E-27 | 1.89E-11 | 2.27E-02 | 1.21E-01 | 2.55E-12 | 1.54E-02 | 4.42E-06 | 1.28E-10 | 1.00E+00 | 8.96E-02 | 5.53E-01 |
| Meta_CPR3009 | 2.60E-27 | 3.31E-25 | 3.49E-16 | 1.04E-16 | 4.27E-31 | 1.16E-14 | 8.58E-18 | 3.55E-14 | 1.58E-24 | 2.38E-18 | 7.88E-10 | 4.11E-03 | 8.96E-02 | 1.28E-10 | 6.45E-03 | 2.23E-06 | 1.85E-09 | 8.96E-02 | 1.00E+00 | 8.96E-02 |
| Meta_CPR3010 | 7.24E-39 | 1.56E-33 | 1.58E-24 | 1.34E-26 | 7.24E-39 | 3.29E-23 | 1.34E-26 | 6.21E-22 | 2.21E-34 | 8.98E-29 | 1.85E-09 | 7.74E-01 | 1.00E+00 | 3.19E-10 | 4.52E-01 | 6.45E-03 | 1.01E-08 | 5.53E-01 | 8.96E-02 | 1.00E+00 |

Comparison of trees using the Branch-length aware Robinson-Foulds method

| | BRS3001 | BRS3002 | BRS3003 | BRS3004 | BRS3005 | BRS3006 | BRS3007 | BRS3008 | BRS3009 | BRS3010 | Meta_CPR3001 | Meta_CPR3002 | Meta_CPR3003 | Meta_CPR3004 | Meta_CPR3005 | Meta_CPR3006 | Meta_CPR3007 | Meta_CPR3008 | Meta_CPR3009 | Meta_CPR3010 |
|---|---|---|---|---|---|---|---|---|---|---|---|---|---|---|---|---|---|---|---|---|
| BRS3001 | 1.00E+00 | 3.38E-02 | 2.60E-07 | 6.22E-16 | 2.51E-11 | 2.66E-08 | 1.23E-07 | 7.74E-01 | 1.60E-03 | 3.31E-04 | 3.56E-55 | 1.69E-23 | 3.56E-26 | 1.73E-25 | 3.56E-26 | 3.67E-24 | 4.71E-29 | 8.18E-25 | 1.34E-21 | 3.67E-24 |
| BRS3002 | 3.38E-02 | 1.00E+00 | 1.02E-02 | 2.66E-08 | 3.31E-04 | 4.20E-03 | 1.55E-02 | 6.69E-02 | 2.24E-01 | 5.95E-08 | 4.23E-50 | 1.60E-13 | 1.91E-15 | 1.22E-18 | 5.63E-14 | 6.08E-15 | 4.44E-18 | 6.22E-16 | 1.89E-14 | 1.60E-13 |
| BRS3003 | 2.60E-07 | 1.02E-02 | 1.00E+00 | 1.91E-04 | 4.79E-02 | 7.74E-01 | 9.63E-01 | 1.72E-05 | 6.69E-02 | 1.91E-15 | 4.23E-50 | 9.26E-10 | 1.23E-12 | 6.22E-16 | 9.26E-10 | 6.32E-11 | 4.51E-13 | 1.23E-12 | 1.18E-08 | 1.18E-08 |
| BRS3004 | 6.22E-16 | 2.66E-08 | 1.91E-04 | 1.00E+00 | 3.38E-02 | 5.70E-04 | 1.72E-05 | 1.23E-12 | 2.31E-06 | 1.73E-25 | 2.50E-43 | 1.02E-02 | 1.07E-04 | 5.49E-07 | 2.31E-02 | 9.67E-04 | 1.07E-04 | 1.07E-04 | 4.79E-02 | 6.59E-03 |
| BRS3005 | 2.51E-11 | 3.31E-04 | 4.79E-02 | 3.38E-02 | 1.00E+00 | 2.87E-01 | 9.25E-02 | 1.23E-07 | 4.20E-03 | 1.22E-18 | 2.50E-43 | 3.22E-05 | 1.23E-07 | 2.19E-09 | 9.11E-06 | 1.23E-07 | 2.19E-09 | 2.60E-07 | 3.22E-05 | 5.97E-05 |
| BRS3006 | 2.66E-08 | 4.20E-03 | 7.74E-01 | 5.70E-04 | 2.87E-01 | 1.00E+00 | 5.62E-01 | 1.14E-06 | 2.31E-02 | 1.22E-18 | 9.02E-48 | 1.23E-07 | 6.32E-11 | 4.51E-13 | 2.60E-07 | 3.89E-10 | 3.44E-12 | 3.89E-10 | 1.23E-07 | 1.23E-07 |
| BRS3007 | 1.23E-07 | 1.55E-02 | 9.63E-01 | 1.72E-05 | 9.25E-02 | 5.62E-01 | 1.00E+00 | 1.72E-05 | 6.69E-02 | 1.91E-15 | 6.88E-54 | 3.89E-10 | 4.51E-13 | 6.22E-16 | 6.32E-11 | 1.23E-12 | 1.91E-15 | 5.63E-14 | 1.60E-10 | 9.26E-10 |
| BRS3008 | 7.74E-01 | 6.69E-02 | 1.72E-05 | 1.23E-12 | 1.23E-07 | 1.14E-06 | 1.72E-05 | 1.00E+00 | 1.60E-03 | 3.31E-04 | 1.40E-59 | 2.15E-20 | 7.22E-23 | 8.18E-25 | 2.15E-20 | 1.34E-21 | 3.56E-26 | 7.22E-23 | 5.56E-21 | 2.15E-20 |
| BRS3009 | 1.60E-03 | 2.24E-01 | 6.69E-02 | 2.31E-06 | 4.20E-03 | 2.31E-02 | 6.69E-02 | 1.60E-03 | 1.00E+00 | 9.26E-10 | 9.02E-48 | 2.19E-09 | 2.51E-11 | 1.60E-13 | 1.60E-10 | 3.89E-10 | 1.60E-13 | 6.32E-11 | 1.18E-08 | 5.19E-09 |
| BRS3010 | 3.31E-04 | 5.95E-08 | 1.91E-15 | 1.73E-25 | 1.22E-18 | 1.22E-18 | 1.91E-15 | 3.31E-04 | 9.26E-10 | 1.00E+00 | 5.93E-61 | 1.34E-30 | 8.86E-34 | 8.86E-34 | 1.36E-34 | 3.75E-32 | 2.62E-37 | 6.04E-33 | 2.40E-31 | 3.75E-32 |
| Meta_CPR3001 | 3.56E-55 | 4.23E-50 | 4.23E-50 | 2.50E-43 | 2.50E-43 | 9.02E-48 | 6.88E-54 | 1.40E-59 | 9.02E-48 | 5.93E-61 | 1.00E+00 | 2.55E-28 | 3.67E-24 | 7.22E-23 | 7.41E-27 | 2.55E-28 | 1.34E-30 | 8.08E-30 | 1.34E-30 | 7.41E-27 |
| Meta_CPR3002 | 1.69E-23 | 1.60E-13 | 9.26E-10 | 1.02E-02 | 3.22E-05 | 1.23E-07 | 3.89E-10 | 2.15E-20 | 2.19E-09 | 1.34E-30 | 2.55E-28 | 1.00E+00 | 5.62E-01 | 1.27E-01 | 1.00E+00 | 1.00E+00 | 3.69E-01 | 6.76E-01 | 5.62E-01 | 1.00E+00 |
| Meta_CPR3003 | 3.56E-26 | 1.91E-15 | 1.23E-12 | 1.07E-04 | 1.23E-07 | 6.32E-11 | 4.51E-13 | 7.22E-23 | 2.51E-11 | 8.86E-34 | 3.67E-24 | 5.62E-01 | 1.00E+00 | 1.70E-01 | 4.66E-01 | 1.00E+00 | 7.74E-01 | 8.76E-01 | 4.79E-02 | 5.62E-01 |
| Meta_CPR3004 | 1.73E-25 | 1.22E-18 | 6.22E-16 | 5.49E-07 | 2.19E-09 | 4.51E-13 | 6.22E-16 | 8.18E-25 | 1.60E-13 | 8.86E-34 | 7.22E-23 | 1.27E-01 | 1.70E-01 | 1.00E+00 | 4.79E-02 | 1.70E-01 | 1.70E-01 | 2.24E-01 | 6.59E-03 | 9.25E-02 |
| Meta_CPR3005 | 3.56E-26 | 5.63E-14 | 9.26E-10 | 2.31E-02 | 9.11E-06 | 2.60E-07 | 6.32E-11 | 2.15E-20 | 1.60E-10 | 1.36E-34 | 7.41E-27 | 1.00E+00 | 4.66E-01 | 4.79E-02 | 1.00E+00 | 7.74E-01 | 2.87E-01 | 6.76E-01 | 7.74E-01 | 1.00E+00 |
| Meta_CPR3006 | 3.67E-24 | 6.08E-15 | 6.32E-11 | 9.67E-04 | 1.23E-07 | 3.89E-10 | 1.23E-12 | 1.34E-21 | 3.89E-10 | 3.75E-32 | 2.55E-28 | 1.00E+00 | 1.00E+00 | 1.70E-01 | 7.74E-01 | 1.00E+00 | 7.74E-01 | 8.76E-01 | 2.87E-01 | 8.76E-01 |
| Meta_CPR3007 | 4.71E-29 | 4.44E-18 | 4.51E-13 | 1.07E-04 | 2.19E-09 | 3.44E-12 | 1.91E-15 | 3.56E-26 | 1.60E-13 | 2.62E-37 | 1.34E-30 | 3.69E-01 | 7.74E-01 | 1.70E-01 | 2.87E-01 | 7.74E-01 | 1.00E+00 | 9.63E-01 | 6.69E-02 | 2.24E-01 |
| Meta_CPR3008 | 8.18E-25 | 6.22E-16 | 1.23E-12 | 1.07E-04 | 2.60E-07 | 3.89E-10 | 5.63E-14 | 7.22E-23 | 6.32E-11 | 6.04E-33 | 8.08E-30 | 6.76E-01 | 8.76E-01 | 2.24E-01 | 6.76E-01 | 8.76E-01 | 9.63E-01 | 1.00E+00 | 9.25E-02 | 5.62E-01 |
| Meta_CPR3009 | 1.34E-21 | 1.89E-14 | 1.18E-08 | 4.79E-02 | 3.22E-05 | 1.23E-07 | 1.60E-10 | 5.56E-21 | 1.18E-08 | 2.40E-31 | 1.34E-30 | 5.62E-01 | 4.79E-02 | 6.59E-03 | 7.74E-01 | 2.87E-01 | 6.69E-02 | 9.25E-02 | 1.00E+00 | 2.87E-01 |
| Meta_CPR3010 | 3.67E-24 | 1.60E-13 | 1.18E-08 | 6.59E-03 | 5.97E-05 | 1.23E-07 | 9.26E-10 | 2.15E-20 | 5.19E-09 | 3.75E-32 | 7.41E-27 | 1.00E+00 | 5.62E-01 | 9.25E-02 | 1.00E+00 | 8.76E-01 | 2.24E-01 | 5.62E-01 | 2.87E-01 | 1.00E+00 |

**Supplementary Table S4E:** Pairwise FDR corrected p-values of the Kolmogorow-Smirnow test on the distances between phylogenetic trees for 20 ribosomal proteins in 10 samples of bacterial RefSeq genomes vs 10 samples of bacterial MAGs (associated with Supplementary Fig. S7e)

Comparison of trees using the Nye et al.,2006 method

| | BRS3001 | BRS3002 | BRS3003 | BRS3004 | BRS3005 | BRS3006 | BRS3007 | BRS3008 | BRS3009 | BRS3010 | Meta_BMS3001 | Meta_BMS3002 | Meta_BMS3003 | Meta_BMS3004 | Meta_BMS3005 | Meta_BMS3006 | Meta_BMS3007 | Meta_BMS3008 | Meta_BMS3009 | Meta_BMS3010 |
|---|---|---|---|---|---|---|---|---|---|---|---|---|---|---|---|---|---|---|---|---|
| BRS3001 | 1.00E+00 | 5.42E-01 | 1.16E-02 | 5.28E-13 | 2.68E-01 | 9.98E-07 | 2.48E-04 | 1.40E-01 | 5.78E-03 | 2.79E-03 | 7.14E-89 | 1.07E-97 | 1.29E-57 | 2.72E-68 | 4.06E-93 | 1.43E-106 | 9.16E-51 | 2.57E-61 | 2.38E-69 | 4.30E-54 |
| BRS3002 | 5.42E-01 | 1.00E+00 | 1.59E-02 | 1.01E-13 | 1.12E-01 | 9.55E-08 | 2.48E-04 | 8.71E-02 | 8.76E-04 | 2.48E-04 | 1.67E-87 | 3.00E-90 | 8.03E-56 | 3.00E-67 | 1.67E-87 | 1.07E-97 | 2.35E-48 | 1.92E-59 | 2.38E-69 | 2.06E-52 |
| BRS3003 | 1.16E-02 | 1.59E-02 | 1.00E+00 | 3.01E-06 | 3.87E-02 | 1.90E-03 | 5.42E-01 | 2.68E-01 | 6.69E-02 | 6.69E-02 | 3.16E-65 | 7.14E-89 | 3.26E-40 | 5.98E-50 | 1.35E-72 | 7.14E-89 | 1.52E-35 | 2.50E-42 | 3.04E-53 | 7.65E-37 |
| BRS3004 | 5.28E-13 | 1.01E-13 | 3.01E-06 | 1.00E+00 | 7.61E-09 | 2.92E-02 | 8.76E-04 | 3.18E-07 | 5.15E-06 | 1.02E-04 | 3.20E-66 | 1.67E-87 | 1.68E-37 | 3.77E-49 | 1.35E-72 | 1.21E-91 | 4.74E-33 | 7.76E-39 | 5.95E-55 | 4.42E-30 |
| BRS3005 | 2.68E-01 | 1.12E-01 | 3.87E-02 | 7.61E-09 | 1.00E+00 | 8.76E-04 | 1.90E-03 | 2.18E-01 | 1.40E-01 | 2.15E-02 | 3.00E-90 | 1.07E-97 | 1.92E-59 | 1.68E-71 | 4.06E-93 | 1.43E-106 | 9.16E-51 | 2.81E-62 | 1.35E-72 | 2.06E-52 |
| BRS3006 | 9.98E-07 | 9.55E-08 | 1.90E-03 | 2.92E-02 | 8.76E-04 | 1.00E+00 | 3.87E-02 | 6.43E-05 | 2.15E-02 | 1.40E-01 | 2.38E-69 | 3.00E-90 | 6.53E-41 | 2.06E-52 | 3.38E-77 | 3.98E-96 | 3.66E-38 | 2.84E-45 | 1.29E-57 | 2.79E-34 |
| BRS3007 | 2.48E-04 | 2.48E-04 | 5.42E-01 | 8.76E-04 | 1.90E-03 | 3.87E-02 | 1.00E+00 | 6.69E-02 | 1.76E-01 | 1.76E-01 | 3.16E-65 | 7.14E-89 | 1.68E-37 | 1.44E-47 | 1.68E-71 | 4.06E-93 | 2.79E-34 | 6.53E-41 | 4.30E-54 | 1.18E-30 |
| BRS3008 | 1.40E-01 | 8.71E-02 | 2.68E-01 | 3.18E-07 | 2.18E-01 | 6.43E-05 | 6.69E-02 | 1.00E+00 | 5.87E-04 | 1.29E-03 | 3.63E-82 | 3.98E-96 | 1.41E-51 | 3.06E-63 | 1.21E-91 | 8.49E-103 | 8.48E-47 | 5.95E-55 | 3.16E-65 | 8.89E-44 |
| BRS3009 | 5.78E-03 | 8.76E-04 | 6.69E-02 | 5.15E-06 | 1.40E-01 | 2.15E-02 | 1.76E-01 | 5.87E-04 | 1.00E+00 | 5.42E-01 | 7.59E-75 | 4.06E-93 | 8.48E-47 | 1.29E-57 | 3.63E-82 | 5.11E-101 | 1.30E-41 | 3.77E-49 | 2.27E-60 | 2.50E-42 |
| BRS3010 | 2.79E-03 | 2.48E-04 | 6.69E-02 | 1.02E-04 | 2.15E-02 | 1.40E-01 | 1.76E-01 | 1.29E-03 | 5.42E-01 | 1.00E+00 | 3.63E-82 | 1.49E-94 | 9.16E-51 | 2.81E-62 | 3.00E-90 | 1.20E-104 | 2.84E-45 | 3.04E-53 | 3.16E-64 | 2.50E-42 |
| Meta_BMS3001 | 7.14E-89 | 1.67E-87 | 3.16E-65 | 3.20E-66 | 3.00E-90 | 2.38E-69 | 3.16E-65 | 3.63E-82 | 7.59E-75 | 3.63E-82 | 1.00E+00 | 1.18E-12 | 1.02E-04 | 1.12E-01 | 1.59E-02 | 1.61E-04 | 2.00E-09 | 5.66E-11 | 3.25E-01 | 5.66E-11 |
| Meta_BMS3002 | 1.07E-97 | 3.00E-90 | 7.14E-89 | 1.67E-87 | 1.07E-97 | 3.00E-90 | 7.14E-89 | 3.98E-96 | 4.06E-93 | 1.49E-94 | 1.18E-12 | 1.00E+00 | 3.95E-24 | 8.09E-17 | 7.61E-09 | 1.29E-03 | 4.42E-30 | 1.15E-33 | 1.30E-15 | 1.15E-33 |
| Meta_BMS3003 | 1.29E-57 | 8.03E-56 | 3.26E-40 | 1.68E-37 | 1.92E-59 | 6.53E-41 | 1.68E-37 | 1.41E-51 | 8.48E-47 | 9.16E-51 | 1.02E-04 | 3.95E-24 | 1.00E+00 | 1.59E-02 | 2.00E-09 | 1.83E-14 | 2.18E-01 | 2.15E-02 | 5.78E-03 | 6.69E-02 |
| Meta_BMS3004 | 2.72E-68 | 3.00E-67 | 5.98E-50 | 3.77E-49 | 1.68E-71 | 2.06E-52 | 1.44E-47 | 3.06E-63 | 1.29E-57 | 2.81E-62 | 1.12E-01 | 8.09E-17 | 1.59E-02 | 1.00E+00 | 1.29E-03 | 1.74E-06 | 1.45E-05 | 9.55E-08 | 8.07E-01 | 5.69E-07 |
| Meta_BMS3005 | 4.06E-93 | 1.67E-87 | 1.35E-72 | 1.35E-72 | 4.06E-93 | 3.38E-77 | 1.68E-71 | 1.21E-91 | 3.63E-82 | 3.00E-90 | 1.59E-02 | 7.61E-09 | 2.00E-09 | 1.29E-03 | 1.00E+00 | 3.87E-02 | 3.16E-15 | 4.50E-18 | 5.87E-04 | 1.68E-18 |
| Meta_BMS3006 | 1.43E-106 | 1.07E-97 | 7.14E-89 | 1.21E-91 | 1.43E-106 | 3.98E-96 | 4.06E-93 | 8.49E-103 | 5.11E-101 | 1.20E-104 | 1.61E-04 | 1.29E-03 | 1.83E-14 | 1.74E-06 | 3.87E-02 | 1.00E+00 | 1.11E-21 | 1.14E-25 | 9.98E-07 | 1.23E-24 |
| Meta_BMS3007 | 9.16E-51 | 2.35E-48 | 1.52E-35 | 4.74E-33 | 9.16E-51 | 3.66E-38 | 2.79E-34 | 8.48E-47 | 1.30E-41 | 2.84E-45 | 2.00E-09 | 4.42E-30 | 2.18E-01 | 1.45E-05 | 3.16E-15 | 1.11E-21 | 1.00E+00 | 3.91E-01 | 8.69E-06 | 5.42E-01 |
| Meta_BMS3008 | 2.57E-61 | 1.92E-59 | 2.50E-42 | 7.76E-39 | 2.81E-62 | 2.84E-45 | 6.53E-41 | 5.95E-55 | 3.77E-49 | 3.04E-53 | 5.66E-11 | 1.15E-33 | 2.15E-02 | 9.55E-08 | 4.50E-18 | 1.14E-25 | 3.91E-01 | 1.00E+00 | 5.21E-08 | 3.25E-01 |
| Meta_BMS3009 | 2.38E-69 | 2.38E-69 | 3.04E-53 | 5.95E-55 | 1.35E-72 | 1.29E-57 | 4.30E-54 | 3.16E-65 | 2.27E-60 | 3.16E-64 | 3.25E-01 | 1.30E-15 | 5.78E-03 | 8.07E-01 | 5.87E-04 | 9.98E-07 | 8.69E-06 | 5.21E-08 | 1.00E+00 | 2.78E-08 |
| Meta_BMS3010 | 4.30E-54 | 2.06E-52 | 7.65E-37 | 4.42E-30 | 2.06E-52 | 2.79E-34 | 1.18E-30 | 8.89E-44 | 2.50E-42 | 2.50E-42 | 5.66E-11 | 1.15E-33 | 6.69E-02 | 5.69E-07 | 1.68E-18 | 1.23E-24 | 5.42E-01 | 3.25E-01 | 2.78E-08 | 1.00E+00 |

Comparison of trees using the Williams & Clifford, 1971 method

| | BRS3001 | BRS3002 | BRS3003 | BRS3004 | BRS3005 | BRS3006 | BRS3007 | BRS3008 | BRS3009 | BRS3010 | Meta_BMS3001 | Meta_BMS3002 | Meta_BMS3003 | Meta_BMS3004 | Meta_BMS3005 | Meta_BMS3006 | Meta_BMS3007 | Meta_BMS3008 | Meta_BMS3009 | Meta_BMS3010 |
|---|---|---|---|---|---|---|---|---|---|---|---|---|---|---|---|---|---|---|---|---|
| BRS3001 | 1.00E+00 | 1.21E-06 | 4.32E-02 | 1.39E-10 | 6.59E-03 | 6.87E-07 | 4.43E-04 | 1.79E-02 | 3.29E-08 | 6.22E-06 | 1.68E-74 | 5.72E-67 | 3.69E-47 | 7.47E-57 | 3.03E-63 | 2.59E-54 | 3.69E-47 | 1.91E-60 | 4.69E-65 | 2.50E-62 |
| BRS3002 | 1.21E-06 | 1.00E+00 | 2.45E-02 | 6.59E-03 | 1.46E-01 | 4.02E-01 | 4.02E-01 | 5.62E-02 | 2.80E-01 | 6.37E-01 | 3.65E-55 | 7.15E-48 | 9.32E-28 | 1.10E-40 | 3.69E-47 | 3.19E-44 | 5.35E-30 | 2.48E-39 | 4.86E-51 | 8.84E-43 |
| BRS3003 | 4.32E-02 | 2.45E-02 | 1.00E+00 | 7.47E-05 | 5.53E-01 | 3.23E-03 | 1.17E-01 | 4.71E-01 | 3.23E-03 | 7.29E-02 | 1.17E-58 | 1.69E-53 | 5.29E-38 | 5.91E-45 | 4.86E-51 | 1.66E-43 | 2.06E-35 | 2.03E-46 | 1.69E-53 | 7.15E-48 |
| BRS3004 | 1.39E-10 | 6.59E-03 | 7.47E-05 | 1.00E+00 | 1.85E-04 | 1.82E-01 | 3.23E-03 | 1.85E-04 | 4.32E-02 | 3.23E-03 | 5.29E-38 | 2.63E-28 | 3.43E-17 | 4.29E-25 | 9.58E-32 | 2.45E-32 | 3.43E-17 | 1.34E-22 | 5.35E-30 | 1.42E-23 |
| BRS3005 | 6.59E-03 | 1.46E-01 | 5.53E-01 | 1.85E-04 | 1.00E+00 | 1.17E-01 | 1.17E-01 | 1.17E-01 | 4.32E-02 | 2.27E-01 | 3.71E-64 | 5.48E-56 | 2.45E-32 | 5.91E-45 | 8.25E-52 | 3.69E-47 | 1.52E-33 | 5.91E-45 | 3.65E-55 | 1.13E-45 |
| BRS3006 | 6.87E-07 | 4.02E-01 | 3.23E-03 | 1.82E-01 | 1.17E-01 | 1.00E+00 | 1.17E-01 | 1.79E-02 | 4.02E-01 | 4.71E-01 | 2.03E-46 | 5.18E-40 | 1.22E-21 | 9.58E-32 | 1.16E-38 | 2.40E-37 | 4.41E-23 | 2.45E-32 | 1.10E-40 | 1.52E-33 |
| BRS3007 | 4.43E-04 | 4.02E-01 | 1.17E-01 | 3.23E-03 | 1.17E-01 | 1.17E-01 | 1.00E+00 | 4.71E-01 | 1.46E-01 | 3.39E-01 | 2.03E-46 | 5.18E-40 | 1.30E-25 | 2.45E-32 | 5.29E-38 | 3.76E-34 | 1.34E-22 | 1.52E-33 | 5.18E-40 | 2.06E-35 |
| BRS3008 | 1.79E-02 | 5.62E-02 | 4.71E-01 | 1.85E-04 | 1.17E-01 | 1.79E-02 | 4.71E-01 | 1.00E+00 | 1.31E-02 | 4.32E-02 | 4.86E-51 | 3.19E-44 | 3.74E-31 | 4.78E-36 | 4.53E-42 | 1.16E-38 | 2.63E-28 | 2.40E-37 | 1.66E-43 | 2.48E-39 |
| BRS3009 | 3.29E-08 | 2.80E-01 | 3.23E-03 | 4.32E-02 | 4.32E-02 | 4.02E-01 | 1.46E-01 | 1.31E-02 | 1.00E+00 | 4.02E-01 | 2.87E-50 | 1.66E-43 | 1.39E-24 | 4.78E-36 | 4.53E-42 | 5.18E-40 | 3.89E-26 | 3.76E-34 | 1.13E-45 | 4.78E-36 |
| BRS3010 | 6.22E-06 | 6.37E-01 | 7.29E-02 | 3.23E-03 | 2.27E-01 | 4.71E-01 | 3.39E-01 | 4.32E-02 | 4.02E-01 | 1.00E+00 | 1.17E-58 | 2.87E-50 | 7.37E-29 | 1.10E-40 | 3.69E-47 | 5.91E-45 | 5.35E-30 | 1.10E-40 | 2.87E-50 | 1.10E-40 |
| Meta_BMS3001 | 1.68E-74 | 3.65E-55 | 1.17E-58 | 5.29E-38 | 3.71E-64 | 2.03E-46 | 2.03E-46 | 4.86E-51 | 2.87E-50 | 1.17E-58 | 1.00E+00 | 4.43E-04 | 1.18E-09 | 1.03E-03 | 4.32E-02 | 1.17E-01 | 1.03E-05 | 4.62E-09 | 1.79E-02 | 6.15E-08 |
| Meta_BMS3002 | 5.72E-67 | 7.15E-48 | 1.69E-53 | 2.63E-28 | 5.48E-56 | 5.18E-40 | 5.18E-40 | 3.19E-44 | 1.66E-43 | 2.87E-50 | 4.43E-04 | 1.00E+00 | 1.54E-03 | 4.71E-01 | 3.39E-01 | 7.29E-02 | 6.59E-03 | 3.23E-03 | 9.37E-02 | 1.31E-02 |
| Meta_BMS3003 | 3.69E-47 | 9.32E-28 | 5.29E-38 | 3.43E-17 | 2.45E-32 | 1.22E-21 | 1.30E-25 | 3.74E-31 | 1.39E-24 | 7.37E-29 | 1.18E-09 | 1.54E-03 | 1.00E+00 | 5.62E-02 | 1.03E-05 | 2.11E-06 | 1.82E-01 | 9.37E-02 | 1.03E-05 | 4.32E-02 |
| Meta_BMS3004 | 7.47E-57 | 1.10E-40 | 5.91E-45 | 4.29E-25 | 5.91E-45 | 9.58E-32 | 2.45E-32 | 4.78E-36 | 4.78E-36 | 1.10E-40 | 1.03E-03 | 4.71E-01 | 5.62E-02 | 1.00E+00 | 3.33E-02 | 5.62E-02 | 1.82E-01 | 5.62E-02 | 9.37E-02 | 1.46E-01 |
| Meta_BMS3005 | 3.03E-63 | 3.69E-47 | 4.86E-51 | 9.58E-32 | 8.25E-52 | 1.16E-38 | 5.29E-38 | 4.53E-42 | 4.53E-42 | 3.69E-47 | 4.32E-02 | 3.39E-01 | 1.03E-05 | 3.33E-02 | 1.00E+00 | 8.93E-01 | 1.85E-04 | 2.84E-05 | 7.26E-01 | 2.89E-04 |
| Meta_BMS3006 | 2.59E-54 | 3.19E-44 | 1.66E-43 | 2.45E-32 | 3.69E-47 | 2.40E-37 | 3.76E-34 | 1.16E-38 | 5.18E-40 | 5.91E-45 | 1.17E-01 | 7.29E-02 | 2.11E-06 | 5.62E-02 | 8.93E-01 | 1.00E+00 | 3.23E-03 | 6.22E-06 | 6.37E-01 | 7.47E-05 |
| Meta_BMS3007 | 3.69E-47 | 5.35E-30 | 2.06E-35 | 3.43E-17 | 1.52E-33 | 4.41E-23 | 1.34E-22 | 2.63E-28 | 3.89E-26 | 5.35E-30 | 1.03E-05 | 6.59E-03 | 1.82E-01 | 1.82E-01 | 1.85E-04 | 3.23E-03 | 1.00E+00 | 1.46E-01 | 6.59E-03 | 7.29E-02 |
| Meta_BMS3008 | 1.91E-60 | 2.48E-39 | 2.03E-46 | 1.34E-22 | 5.91E-45 | 2.45E-32 | 1.52E-33 | 2.40E-37 | 3.76E-34 | 1.10E-40 | 4.62E-09 | 3.23E-03 | 9.37E-02 | 5.62E-02 | 2.84E-05 | 6.22E-06 | 1.46E-01 | 1.00E+00 | 1.72E-05 | 9.57E-01 |
| Meta_BMS3009 | 4.69E-65 | 4.86E-51 | 1.69E-53 | 5.35E-30 | 3.65E-55 | 1.10E-40 | 5.18E-40 | 1.66E-43 | 1.13E-45 | 2.87E-50 | 1.79E-02 | 9.37E-02 | 1.03E-05 | 9.37E-02 | 7.26E-01 | 6.37E-01 | 6.59E-03 | 1.72E-05 | 1.00E+00 | 1.85E-04 |
| Meta_BMS3010 | 2.50E-62 | 8.84E-43 | 7.15E-48 | 1.42E-23 | 1.13E-45 | 1.52E-33 | 2.06E-35 | 2.48E-39 | 4.78E-36 | 1.10E-40 | 6.15E-08 | 1.31E-02 | 4.32E-02 | 1.46E-01 | 2.89E-04 | 7.47E-05 | 7.29E-02 | 9.57E-01 | 1.85E-04 | 1.00E+00 |

Comparison of trees using the Robinson-Foulds method

| | BRS3001 | BRS3002 | BRS3003 | BRS3004 | BRS3005 | BRS3006 | BRS3007 | BRS3008 | BRS3009 | BRS3010 | Meta_BMS3001 | Meta_BMS3002 | Meta_BMS3003 | Meta_BMS3004 | Meta_BMS3005 | Meta_BMS3006 | Meta_BMS3007 | Meta_BMS3008 | Meta_BMS3009 | Meta_BMS3010 |
|---|---|---|---|---|---|---|---|---|---|---|---|---|---|---|---|---|---|---|---|---|
| BRS3001 | 1.00E+00 | 7.42E-01 | 6.04E-04 | 7.55E-09 | 9.67E-01 | 5.21E-08 | 9.08E-02 | 2.77E-01 | 7.02E-02 | 4.10E-02 | 1.58E-91 | 1.70E-102 | 6.98E-55 | 3.52E-66 | 9.32E-81 | 3.60E-104 | 5.85E-43 | 1.98E-37 | 8.08E-75 | 3.03E-48 |
| BRS3002 | 7.42E-01 | 1.00E+00 | 1.67E-02 | 1.74E-06 | 2.27E-01 | 8.76E-06 | 5.59E-01 | 9.67E-01 | 3.07E-02 | 3.07E-02 | 2.54E-83 | 5.54E-93 | 1.10E-43 | 3.65E-53 | 1.08E-73 | 1.83E-94 | 1.76E-35 | 1.29E-30 | 2.62E-60 | 4.03E-40 |
| BRS3003 | 6.04E-04 | 1.67E-02 | 1.00E+00 | 3.07E-02 | 8.53E-03 | 7.02E-02 | 2.77E-01 | 1.15E-01 | 4.10E-02 | 1.83E-01 | 2.79E-69 | 5.71E-86 | 1.29E-30 | 3.09E-42 | 3.03E-61 | 1.49E-79 | 1.12E-21 | 2.04E-16 | 3.03E-48 | 1.17E-25 |
| BRS3004 | 7.55E-09 | 1.74E-06 | 3.07E-02 | 1.00E+00 | 5.73E-07 | 2.77E-01 | 2.96E-03 | 1.66E-04 | 6.04E-04 | 8.53E-03 | 3.18E-68 | 5.71E-86 | 1.04E-26 | 1.57E-41 | 2.62E-60 | 1.49E-79 | 1.65E-18 | 5.70E-12 | 1.84E-47 | 1.12E-21 |
| BRS3005 | 9.67E-01 | 2.27E-01 | 8.53E-03 | 5.73E-07 | 1.00E+00 | 3.01E-06 | 4.02E-01 | 3.34E-01 | 9.08E-02 | 2.27E-01 | 2.54E-83 | 6.63E-96 | 6.20E-46 | 9.48E-56 | 1.08E-73 | 1.83E-94 | 8.88E-37 | 8.72E-32 | 3.57E-63 | 1.57E-41 |
| BRS3006 | 5.21E-08 | 8.76E-06 | 7.02E-02 | 2.77E-01 | 3.01E-06 | 1.00E+00 | 8.53E-03 | 3.98E-04 | 4.26E-03 | 2.29E-02 | 8.08E-75 | 5.54E-93 | 1.98E-37 | 6.20E-46 | 3.52E-66 | 5.71E-86 | 3.02E-27 | 7.96E-20 | 2.50E-52 | 8.72E-32 |
| BRS3007 | 9.08E-02 | 5.59E-01 | 2.77E-01 | 2.96E-03 | 4.02E-01 | 8.53E-03 | 1.00E+00 | 1.00E+00 | 3.34E-01 | 3.34E-01 | 5.81E-76 | 1.58E-91 | 1.98E-37 | 1.10E-46 | 3.40E-67 | 5.71E-86 | 4.84E-30 | 1.17E-25 | 3.65E-53 | 3.22E-34 |
| BRS3008 | 2.77E-01 | 9.67E-01 | 1.15E-01 | 1.66E-04 | 3.34E-01 | 3.98E-04 | 1.00E+00 | 1.00E+00 | 1.45E-01 | 1.15E-01 | 5.81E-76 | 1.83E-94 | 5.85E-43 | 2.50E-52 | 3.40E-67 | 3.12E-87 | 2.22E-32 | 8.64E-28 | 2.25E-59 | 8.88E-37 |
| BRS3009 | 7.02E-02 | 3.07E-02 | 4.10E-02 | 6.04E-04 | 9.08E-02 | 4.26E-03 | 3.34E-01 | 1.45E-01 | 1.00E+00 | 1.00E+00 | 1.49E-79 | 6.63E-96 | 6.20E-46 | 9.48E-56 | 2.37E-70 | 4.50E-90 | 7.60E-35 | 4.84E-30 | 3.57E-63 | 1.97E-39 |
| BRS3010 | 4.10E-02 | 3.07E-02 | 1.83E-01 | 8.53E-03 | 2.27E-01 | 2.29E-02 | 3.34E-01 | 1.15E-01 | 1.00E+00 | 1.00E+00 | 5.71E-86 | 2.13E-97 | 3.54E-45 | 6.98E-55 | 5.81E-76 | 2.13E-97 | 1.33E-33 | 3.54E-26 | 3.34E-62 | 9.44E-39 |
| Meta_BMS3001 | 1.58E-91 | 2.54E-83 | 2.79E-69 | 3.18E-68 | 2.54E-83 | 8.08E-75 | 5.81E-76 | 5.81E-76 | 1.49E-79 | 5.71E-86 | 1.00E+00 | 2.20E-19 | 9.63E-08 | 1.37E-03 | 8.25E-01 | 3.21E-07 | 1.17E-12 | 1.29E-23 | 7.02E-02 | 4.27E-14 |
| Meta_BMS3002 | 1.70E-102 | 5.54E-93 | 5.71E-86 | 5.71E-86 | 6.63E-96 | 5.54E-93 | 1.58E-91 | 1.83E-94 | 6.63E-96 | 2.13E-97 | 2.20E-19 | 1.00E+00 | 1.33E-33 | 1.29E-30 | 1.83E-14 | 4.26E-03 | 4.39E-38 | 1.56E-57 | 4.10E-24 | 1.16E-50 |
| Meta_BMS3003 | 6.98E-55 | 1.10E-43 | 1.29E-30 | 1.04E-26 | 6.20E-46 | 1.98E-37 | 1.98E-37 | 5.85E-43 | 6.20E-46 | 3.54E-45 | 9.63E-08 | 1.33E-33 | 1.00E+00 | 1.15E-01 | 1.74E-06 | 1.18E-17 | 4.76E-01 | 2.59E-04 | 2.96E-03 | 1.15E-01 |
| Meta_BMS3004 | 3.52E-66 | 3.65E-53 | 3.09E-42 | 1.57E-41 | 9.48E-56 | 6.20E-46 | 1.10E-46 | 2.50E-52 | 9.48E-56 | 6.98E-55 | 1.37E-03 | 1.29E-30 | 1.15E-01 | 1.00E+00 | 6.10E-03 | 4.27E-14 | 3.98E-04 | 7.55E-09 | 8.25E-01 | 9.13E-04 |
| Meta_BMS3005 | 9.32E-81 | 1.08E-73 | 3.03E-61 | 2.62E-60 | 1.08E-73 | 3.52E-66 | 3.40E-67 | 3.40E-67 | 2.37E-70 | 5.81E-76 | 8.25E-01 | 1.83E-14 | 1.74E-06 | 6.10E-03 | 1.00E+00 | 1.66E-04 | 5.56E-11 | 2.20E-19 | 1.45E-01 | 5.56E-11 |
| Meta_BMS3006 | 3.60E-104 | 1.83E-94 | 1.49E-79 | 1.49E-79 | 1.83E-94 | 5.71E-86 | 5.71E-86 | 3.12E-87 | 4.50E-90 | 2.13E-97 | 3.21E-07 | 4.26E-03 | 1.18E-17 | 4.27E-14 | 1.66E-04 | 1.00E+00 | 4.03E-23 | 4.39E-38 | 2.40E-10 | 1.80E-29 |
| Meta_BMS3007 | 5.85E-43 | 1.76E-35 | 1.12E-21 | 1.65E-18 | 8.88E-37 | 3.02E-27 | 4.84E-30 | 2.22E-32 | 7.60E-35 | 1.33E-33 | 1.17E-12 | 4.39E-38 | 4.76E-01 | 3.98E-04 | 5.56E-11 | 4.03E-23 | 1.00E+00 | 5.44E-02 | 1.74E-06 | 4.76E-01 |
| Meta_BMS3008 | 1.98E-37 | 1.29E-30 | 2.04E-16 | 5.70E-12 | 8.72E-32 | 7.96E-20 | 1.17E-25 | 8.64E-28 | 4.84E-30 | 3.54E-26 | 1.29E-23 | 1.56E-57 | 2.59E-04 | 7.55E-09 | 2.20E-19 | 4.39E-38 | 5.44E-02 | 1.00E+00 | 1.17E-12 | 7.02E-02 |
| Meta_BMS3009 | 8.08E-75 | 2.62E-60 | 3.03E-48 | 1.84E-47 | 3.57E-63 | 2.50E-52 | 3.65E-53 | 2.25E-59 | 3.57E-63 | 3.34E-62 | 7.02E-02 | 4.10E-24 | 2.96E-03 | 8.25E-01 | 1.45E-01 | 2.40E-10 | 1.74E-06 | 1.17E-12 | 1.00E+00 | 1.74E-06 |
| Meta_BMS3010 | 3.03E-48 | 4.03E-40 | 1.17E-25 | 1.12E-21 | 1.57E-41 | 8.72E-32 | 3.22E-34 | 8.88E-37 | 1.97E-39 | 9.44E-39 | 4.27E-14 | 1.16E-50 | 1.15E-01 | 9.13E-04 | 5.56E-11 | 1.80E-29 | 4.76E-01 | 7.02E-02 | 1.74E-06 | 1.00E+00 |

Comparison of trees using the Branch-length aware Robinson-Foulds method

| | BRS3001 | BRS3002 | BRS3003 | BRS3004 | BRS3005 | BRS3006 | BRS3007 | BRS3008 | BRS3009 | BRS3010 | Meta_BMS3001 | Meta_BMS3002 | Meta_BMS3003 | Meta_BMS3004 | Meta_BMS3005 | Meta_BMS3006 | Meta_BMS3007 | Meta_BMS3008 | Meta_BMS3009 | Meta_BMS3010 |
|---|---|---|---|---|---|---|---|---|---|---|---|---|---|---|---|---|---|---|---|---|
| BRS3001 | 1.00E+00 | 5.45E-03 | 6.45E-10 | 6.82E-13 | 3.19E-13 | 5.12E-09 | 1.90E-08 | 1.07E-02 | 1.17E-07 | 1.98E-02 | 1.18E-24 | 1.19E-28 | 2.07E-26 | 3.71E-16 | 3.89E-25 | 3.58E-24 | 8.10E-22 | 1.40E-13 | 2.07E-26 | 1.03E-22 |
| BRS3002 | 5.45E-03 | 1.00E+00 | 5.44E-04 | 1.09E-06 | 1.51E-05 | 7.73E-03 | 1.46E-02 | 3.14E-01 | 1.98E-02 | 3.84E-05 | 6.09E-14 | 1.01E-17 | 2.06E-15 | 1.17E-07 | 2.69E-14 | 2.69E-14 | 1.45E-12 | 3.17E-06 | 2.06E-15 | 1.45E-12 |
| BRS3003 | 6.45E-10 | 5.44E-04 | 1.00E+00 | 4.41E-01 | 6.89E-01 | 4.41E-01 | 4.41E-01 | 3.84E-05 | 4.41E-01 | 6.82E-13 | 3.67E-04 | 5.30E-06 | 1.55E-04 | 6.23E-02 | 2.38E-04 | 2.38E-04 | 3.67E-04 | 3.52E-02 | 1.00E-04 | 1.85E-03 |
| BRS3004 | 6.82E-13 | 1.09E-06 | 4.41E-01 | 1.00E+00 | 2.60E-01 | 2.64E-02 | 1.46E-02 | 6.58E-08 | 2.64E-02 | 4.96E-15 | 5.44E-04 | 3.84E-05 | 1.24E-03 | 1.03E-01 | 5.45E-03 | 5.44E-04 | 1.55E-04 | 1.67E-01 | 2.68E-03 | 5.45E-03 |
| BRS3005 | 3.19E-13 | 1.51E-05 | 6.89E-01 | 2.60E-01 | 1.00E+00 | 3.78E-01 | 8.01E-02 | 3.84E-05 | 2.09E-01 | 8.77E-16 | 1.55E-04 | 2.48E-05 | 2.38E-04 | 2.64E-02 | 5.44E-04 | 1.55E-04 | 8.21E-04 | 8.01E-02 | 5.44E-04 | 3.91E-03 |
| BRS3006 | 5.12E-09 | 7.73E-03 | 4.41E-01 | 2.64E-02 | 3.78E-01 | 1.00E+00 | 1.00E+00 | 5.45E-03 | 7.66E-01 | 6.82E-13 | 3.17E-06 | 3.52E-08 | 6.42E-07 | 1.98E-02 | 3.17E-06 | 3.17E-06 | 3.84E-05 | 1.03E-01 | 6.42E-07 | 3.84E-05 |
| BRS3007 | 1.90E-08 | 1.46E-02 | 4.41E-01 | 1.46E-02 | 8.01E-02 | 1.00E+00 | 1.00E+00 | 5.45E-03 | 9.23E-01 | 3.19E-12 | 6.42E-07 | 5.12E-09 | 1.17E-07 | 2.68E-03 | 3.76E-07 | 1.09E-06 | 1.51E-05 | 2.64E-02 | 2.15E-07 | 5.30E-06 |
| BRS3008 | 1.07E-02 | 3.14E-01 | 3.84E-05 | 6.58E-08 | 3.84E-05 | 5.45E-03 | 5.45E-03 | 1.00E+00 | 1.07E-02 | 6.25E-05 | 1.18E-14 | 1.76E-19 | 2.33E-17 | 3.52E-08 | 8.77E-16 | 5.93E-17 | 6.09E-14 | 6.42E-07 | 3.71E-16 | 1.45E-12 |
| BRS3009 | 1.17E-07 | 1.98E-02 | 4.41E-01 | 2.64E-02 | 2.09E-01 | 7.66E-01 | 9.23E-01 | 1.07E-02 | 1.00E+00 | 3.18E-10 | 1.09E-06 | 1.90E-08 | 3.76E-07 | 2.68E-03 | 1.91E-06 | 1.09E-06 | 5.30E-06 | 2.64E-02 | 3.76E-07 | 2.48E-05 |
| BRS3010 | 1.98E-02 | 3.84E-05 | 6.82E-13 | 4.96E-15 | 8.77E-16 | 6.82E-13 | 3.19E-12 | 6.25E-05 | 3.18E-10 | 1.00E+00 | 1.08E-23 | 2.27E-28 | 5.87E-26 | 2.33E-17 | 3.89E-25 | 3.89E-25 | 2.79E-22 | 2.33E-17 | 1.70E-25 | 2.79E-22 |
| Meta_BMS3001 | 1.18E-24 | 6.09E-14 | 3.67E-04 | 5.44E-04 | 1.55E-04 | 3.17E-06 | 6.42E-07 | 1.18E-14 | 1.09E-06 | 1.08E-23 | 1.00E+00 | 5.25E-01 | 9.77E-01 | 1.67E-01 | 6.89E-01 | 9.77E-01 | 1.00E+00 | 3.52E-02 | 9.23E-01 | 3.14E-01 |
| Meta_BMS3002 | 1.19E-28 | 1.01E-17 | 5.30E-06 | 3.84E-05 | 2.48E-05 | 3.52E-08 | 5.12E-09 | 1.76E-19 | 1.90E-08 | 2.27E-28 | 5.25E-01 | 1.00E+00 | 9.23E-01 | 1.07E-02 | 3.14E-01 | 8.57E-01 | 4.41E-01 | 8.21E-04 | 6.13E-01 | 1.32E-01 |
| Meta_BMS3003 | 2.07E-26 | 2.06E-15 | 1.55E-04 | 1.24E-03 | 2.38E-04 | 6.42E-07 | 1.17E-07 | 2.33E-17 | 3.76E-07 | 5.87E-26 | 9.77E-01 | 9.23E-01 | 1.00E+00 | 3.52E-02 | 7.66E-01 | 1.00E+00 | 6.89E-01 | 7.73E-03 | 6.89E-01 | 3.78E-01 |
| Meta_BMS3004 | 3.71E-16 | 1.17E-07 | 6.23E-02 | 1.03E-01 | 2.64E-02 | 1.98E-02 | 2.68E-03 | 3.52E-08 | 2.68E-03 | 2.33E-17 | 1.67E-01 | 1.07E-02 | 3.52E-02 | 1.00E+00 | 6.23E-02 | 4.73E-02 | 2.09E-01 | 9.77E-01 | 8.01E-02 | 3.78E-01 |
| Meta_BMS3005 | 3.89E-25 | 2.69E-14 | 2.38E-04 | 5.45E-03 | 5.44E-04 | 3.17E-06 | 3.76E-07 | 8.77E-16 | 1.91E-06 | 3.89E-25 | 6.89E-01 | 3.14E-01 | 7.66E-01 | 6.23E-02 | 1.00E+00 | 6.89E-01 | 7.66E-01 | 1.46E-02 | 1.00E+00 | 4.41E-01 |
| Meta_BMS3006 | 3.58E-24 | 2.69E-14 | 2.38E-04 | 5.44E-04 | 1.55E-04 | 3.17E-06 | 1.09E-06 | 5.93E-17 | 1.09E-06 | 3.89E-25 | 9.77E-01 | 8.57E-01 | 1.00E+00 | 4.73E-02 | 6.89E-01 | 1.00E+00 | 9.23E-01 | 1.46E-02 | 6.89E-01 | 3.14E-01 |
| Meta_BMS3007 | 8.10E-22 | 1.45E-12 | 3.67E-04 | 1.55E-04 | 8.21E-04 | 3.84E-05 | 1.51E-05 | 6.09E-14 | 5.30E-06 | 2.79E-22 | 1.00E+00 | 4.41E-01 | 6.89E-01 | 2.09E-01 | 7.66E-01 | 9.23E-01 | 1.00E+00 | 1.03E-01 | 7.66E-01 | 7.66E-01 |
| Meta_BMS3008 | 1.40E-13 | 3.17E-06 | 3.52E-02 | 1.67E-01 | 8.01E-02 | 1.03E-01 | 2.64E-02 | 6.42E-07 | 2.64E-02 | 2.33E-17 | 3.52E-02 | 8.21E-04 | 7.73E-03 | 9.77E-01 | 1.46E-02 | 1.46E-02 | 1.03E-01 | 1.00E+00 | 1.07E-02 | 8.01E-02 |
| Meta_BMS3009 | 2.07E-26 | 2.06E-15 | 1.00E-04 | 2.68E-03 | 5.44E-04 | 6.42E-07 | 2.15E-07 | 3.71E-16 | 3.76E-07 | 1.70E-25 | 9.23E-01 | 6.13E-01 | 6.89E-01 | 8.01E-02 | 1.00E+00 | 6.89E-01 | 7.66E-01 | 1.07E-02 | 1.00E+00 | 6.13E-01 |
| Meta_BMS3010 | 1.03E-22 | 1.45E-12 | 1.85E-03 | 5.45E-03 | 3.91E-03 | 3.84E-05 | 5.30E-06 | 1.45E-12 | 2.48E-05 | 2.79E-22 | 3.14E-01 | 1.32E-01 | 3.78E-01 | 3.78E-01 | 4.41E-01 | 3.14E-01 | 7.66E-01 | 8.01E-02 | 6.13E-01 | 1.00E+00 |

**Supplementary Table S3A:** Pairwise FDR corrected p-values of the Kolmogorow-Smirnow test on the distances between phylogenetic trees for 39 universal proteins in 10 samples of archaeal RefSeq genomes vs one sample containing archaeal MAGs (associated with Extended Data 1a)

Comparison of trees using the Nye et al., 2006 method

| | ARS3001 | ARS3002 | ARS3003 | ARS3004 | ARS3005 | ARS3006 | ARS3007 | ARS3008 | ARS3009 | ARS3010 | Metagenomes |
|---|---|---|---|---|---|---|---|---|---|---|---|
| ARS3001 | 1.00E+00 | 1.91E-05 | 6.79E-04 | 1.25E-04 | 1.56E-21 | 4.22E-08 | 1.61E-15 | 5.72E-22 | 3.16E-08 | 6.84E-01 | 0.00E+00 |
| ARS3002 | 1.91E-05 | 1.00E+00 | 1.22E-11 | 6.82E-02 | 1.81E-12 | 1.85E-03 | 6.10E-07 | 4.25E-10 | 1.73E-11 | 2.38E-04 | 0.00E+00 |
| ARS3003 | 6.79E-04 | 1.22E-11 | 1.00E+00 | 3.77E-13 | 3.20E-39 | 8.75E-20 | 2.11E-29 | 3.25E-36 | 1.01E-04 | 3.63E-04 | 0.00E+00 |
| ARS3004 | 1.25E-04 | 6.82E-02 | 3.77E-13 | 1.00E+00 | 3.54E-11 | 6.05E-02 | 2.39E-06 | 3.07E-10 | 2.78E-16 | 8.19E-05 | 0.00E+00 |
| ARS3005 | 1.56E-21 | 1.81E-12 | 3.20E-39 | 3.54E-11 | 1.00E+00 | 2.39E-06 | 4.08E-02 | 2.58E-01 | 4.31E-41 | 5.72E-22 | 0.00E+00 |
| ARS3006 | 4.22E-08 | 1.85E-03 | 8.75E-20 | 6.05E-02 | 2.39E-06 | 1.00E+00 | 2.38E-04 | 2.37E-08 | 2.51E-21 | 4.68E-07 | 0.00E+00 |
| ARS3007 | 1.61E-15 | 6.10E-07 | 2.11E-29 | 2.39E-06 | 4.08E-02 | 2.38E-04 | 1.00E+00 | 4.63E-02 | 3.68E-29 | 1.37E-14 | 0.00E+00 |
| ARS3008 | 5.72E-22 | 4.25E-10 | 3.25E-36 | 3.07E-10 | 2.58E-01 | 2.37E-08 | 4.63E-02 | 1.00E+00 | 1.75E-36 | 2.28E-19 | 0.00E+00 |
| ARS3009 | 3.16E-08 | 1.73E-11 | 1.01E-04 | 2.78E-16 | 4.31E-41 | 2.51E-21 | 3.68E-29 | 1.75E-36 | 1.00E+00 | 9.00E-06 | 0.00E+00 |
| ARS3010 | 6.84E-01 | 2.38E-04 | 3.63E-04 | 8.19E-05 | 5.72E-22 | 4.68E-07 | 1.37E-14 | 2.28E-19 | 9.00E-06 | 1.00E+00 | 0.00E+00 |
| Metagenomes | 0.00E+00 | 0.00E+00 | 0.00E+00 | 0.00E+00 | 0.00E+00 | 0.00E+00 | 0.00E+00 | 0.00E+00 | 0.00E+00 | 0.00E+00 | 1.00E+00 |

Comparison of trees using the Williams & Clifford, 1971 method

| | ARS3001 | ARS3002 | ARS3003 | ARS3004 | ARS3005 | ARS3006 | ARS3007 | ARS3008 | ARS3009 | ARS3010 | Metagenomes |
|---|---|---|---|---|---|---|---|---|---|---|---|
| ARS3001 | 1.00E+00 | 2.19E-01 | 2.93E-01 | 2.26E-03 | 1.71E-11 | 4.24E-06 | 2.96E-04 | 6.42E-08 | 5.32E-17 | 8.42E-08 | 0.00E+00 |
| ARS3002 | 2.19E-01 | 1.00E+00 | 5.09E-01 | 2.68E-01 | 1.44E-07 | 1.55E-03 | 1.77E-02 | 1.01E-04 | 3.14E-21 | 1.21E-04 | 8.96E-305 |
| ARS3003 | 2.93E-01 | 5.09E-01 | 1.00E+00 | 1.77E-02 | 2.61E-09 | 8.57E-04 | 5.73E-03 | 2.52E-06 | 4.99E-21 | 1.46E-05 | 1.11E-303 |
| ARS3004 | 2.26E-03 | 2.68E-01 | 1.77E-02 | 1.00E+00 | 1.46E-05 | 5.46E-02 | 1.57E-02 | 1.57E-02 | 8.30E-28 | 2.69E-02 | 8.86E-298 |
| ARS3005 | 1.71E-11 | 1.44E-07 | 2.61E-09 | 1.46E-05 | 1.00E+00 | 1.21E-04 | 3.51E-09 | 2.39E-02 | 7.40E-49 | 1.12E-06 | 2.78E-209 |
| ARS3006 | 4.24E-06 | 1.55E-03 | 8.57E-04 | 5.46E-02 | 1.21E-04 | 1.00E+00 | 6.43E-05 | 5.46E-02 | 1.99E-37 | 6.36E-01 | 2.25E-287 |
| ARS3007 | 2.96E-04 | 1.77E-02 | 5.73E-03 | 1.57E-02 | 3.51E-09 | 6.43E-05 | 1.00E+00 | 1.44E-07 | 2.19E-17 | 7.05E-06 | 3.09E-257 |
| ARS3008 | 6.42E-08 | 1.01E-04 | 2.52E-06 | 1.57E-02 | 2.39E-02 | 5.46E-02 | 1.44E-07 | 1.00E+00 | 1.88E-41 | 2.69E-02 | 3.26E-258 |
| ARS3009 | 5.32E-17 | 3.14E-21 | 4.99E-21 | 8.30E-28 | 7.40E-49 | 1.99E-37 | 2.19E-17 | 1.88E-41 | 1.00E+00 | 7.64E-41 | 0.00E+00 |
| ARS3010 | 8.42E-08 | 1.21E-04 | 1.46E-05 | 2.69E-02 | 1.12E-06 | 6.36E-01 | 7.05E-06 | 2.69E-02 | 7.64E-41 | 1.00E+00 | 3.64E-285 |
| Metagenomes | 0.00E+00 | 8.96E-305 | 1.11E-303 | 8.86E-298 | 2.78E-209 | 2.25E-287 | 3.09E-257 | 3.26E-258 | 0.00E+00 | 3.64E-285 | 1.00E+00 |

Comparison of trees using the Robinson & Foulds, 1981 method

| | ARS3001 | ARS3002 | ARS3003 | ARS3004 | ARS3005 | ARS3006 | ARS3007 | ARS3008 | ARS3009 | ARS3010 | Metagenomes |
|---|---|---|---|---|---|---|---|---|---|---|---|
| ARS3001 | 1.00E+00 | 2.45E-02 | 9.59E-03 | 5.81E-02 | 1.29E-09 | 6.91E-10 | 1.20E-20 | 9.22E-22 | 4.50E-18 | 1.49E-04 | 0.00E+00 |
| ARS3002 | 2.45E-02 | 1.00E+00 | 1.00E+00 | 1.00E+00 | 8.35E-04 | 3.81E-05 | 2.50E-13 | 3.51E-14 | 1.69E-32 | 9.67E-02 | 0.00E+00 |
| ARS3003 | 9.59E-03 | 1.00E+00 | 1.00E+00 | 7.10E-01 | 4.49E-04 | 1.89E-05 | 7.50E-14 | 6.57E-15 | 3.37E-34 | 6.55E-02 | 0.00E+00 |
| ARS3004 | 5.81E-02 | 1.00E+00 | 7.10E-01 | 1.00E+00 | 4.75E-05 | 1.49E-04 | 2.68E-12 | 5.14E-14 | 9.45E-30 | 2.00E-01 | 0.00E+00 |
| ARS3005 | 1.29E-09 | 8.35E-04 | 4.49E-04 | 4.75E-05 | 1.00E+00 | 6.73E-01 | 2.69E-03 | 3.81E-05 | 9.26E-55 | 5.14E-02 | 0.00E+00 |
| ARS3006 | 6.91E-10 | 3.81E-05 | 1.89E-05 | 1.49E-04 | 6.73E-01 | 1.00E+00 | 2.13E-02 | 1.24E-03 | 4.17E-51 | 1.85E-02 | 0.00E+00 |
| ARS3007 | 1.20E-20 | 2.50E-13 | 7.50E-14 | 2.68E-12 | 2.69E-03 | 2.13E-02 | 1.00E+00 | 2.74E-01 | 4.73E-67 | 3.96E-06 | 6.42E-323 |
| ARS3008 | 9.22E-22 | 3.51E-14 | 6.57E-15 | 5.14E-14 | 3.81E-05 | 1.24E-03 | 2.74E-01 | 1.00E+00 | 6.36E-74 | 1.29E-09 | 9.08E-318 |
| ARS3009 | 4.50E-18 | 1.69E-32 | 3.37E-34 | 9.45E-30 | 9.26E-55 | 4.17E-51 | 4.73E-67 | 6.36E-74 | 1.00E+00 | 1.85E-37 | 0.00E+00 |
| ARS3010 | 1.49E-04 | 9.67E-02 | 6.55E-02 | 2.00E-01 | 5.14E-02 | 1.85E-02 | 3.96E-06 | 1.29E-09 | 1.85E-37 | 1.00E+00 | 0.00E+00 |
| Metagenomes | 0.00E+00 | 0.00E+00 | 0.00E+00 | 0.00E+00 | 0.00E+00 | 0.00E+00 | 6.42E-323 | 9.08E-318 | 0.00E+00 | 0.00E+00 | 1.00E+00 |

Comparison of trees using the Kuhner & Felsenstein, 1994 method

| | ARS3001 | ARS3002 | ARS3003 | ARS3004 | ARS3005 | ARS3006 | ARS3007 | ARS3008 | ARS3009 | ARS3010 | Metagenomes |
|---|---|---|---|---|---|---|---|---|---|---|---|
| ARS3001 | 1.00E+00 | 5.83E-09 | 1.17E-21 | 2.32E-17 | 1.72E-06 | 1.78E-04 | 1.81E-01 | 4.16E-02 | 3.21E-02 | 5.30E-16 | 1.19E-154 |
| ARS3002 | 5.83E-09 | 1.00E+00 | 1.45E-04 | 1.32E-02 | 3.66E-02 | 3.26E-03 | 7.84E-09 | 7.67E-06 | 1.72E-14 | 1.77E-02 | 1.05E-101 |
| ARS3003 | 1.17E-21 | 1.45E-04 | 1.00E+00 | 1.48E-01 | 2.92E-06 | 2.79E-08 | 3.09E-24 | 2.51E-14 | 1.86E-33 | 1.64E-01 | 8.07E-114 |
| ARS3004 | 2.32E-17 | 1.32E-02 | 1.48E-01 | 1.00E+00 | 2.74E-04 | 6.22E-06 | 8.43E-19 | 4.43E-11 | 3.44E-27 | 4.73E-02 | 1.62E-119 |
| ARS3005 | 1.72E-06 | 3.66E-02 | 2.92E-06 | 2.74E-04 | 1.00E+00 | 2.80E-02 | 1.34E-06 | 1.64E-05 | 4.66E-12 | 3.85E-03 | 1.69E-128 |
| ARS3006 | 1.78E-04 | 3.26E-03 | 2.79E-08 | 6.22E-06 | 2.80E-02 | 1.00E+00 | 7.67E-06 | 6.17E-02 | 3.81E-10 | 2.38E-07 | 1.95E-138 |
| ARS3007 | 1.81E-01 | 7.84E-09 | 3.09E-24 | 8.43E-19 | 1.34E-06 | 7.67E-06 | 1.00E+00 | 1.77E-02 | 1.18E-01 | 5.30E-16 | 3.56E-152 |
| ARS3008 | 4.16E-02 | 7.67E-06 | 2.51E-14 | 4.43E-11 | 1.64E-05 | 6.17E-02 | 1.77E-02 | 1.00E+00 | 6.22E-06 | 1.96E-13 | 1.10E-134 |
| ARS3009 | 3.21E-02 | 1.72E-14 | 1.86E-33 | 3.44E-27 | 4.66E-12 | 3.81E-10 | 1.18E-01 | 6.22E-06 | 1.00E+00 | 3.43E-25 | 2.90E-140 |
| ARS3010 | 5.30E-16 | 1.77E-02 | 1.64E-01 | 4.73E-02 | 3.85E-03 | 2.38E-07 | 5.30E-16 | 1.96E-13 | 3.43E-25 | 1.00E+00 | 1.19E-116 |
| Metagenomes | 1.19E-154 | 1.05E-101 | 8.07E-114 | 1.62E-119 | 1.69E-128 | 1.95E-138 | 3.56E-152 | 1.10E-134 | 2.90E-140 | 1.19E-116 | 1.00E+00 |

**Supplementary Table S3B:** Pairwise FDR corrected p-values of the Kolmogorow-Smirnow test on the distances between phylogenetic trees for 23 ribosomal proteins (including SecY) in 10 samples of archaeal RefSeq genomes vs one sample containing archaeal MAGs (associated with Extended Data 1b)

Comparison of trees using the Nye et al., 2006 method

| | ARS3001 | ARS3002 | ARS3003 | ARS3004 | ARS3005 | ARS3006 | ARS3007 | ARS3008 | ARS3009 | ARS3010 | Metagenoms |
|---|---|---|---|---|---|---|---|---|---|---|---|
| ARS3001 | 1.00E+00 | 7.20E-01 | 9.94E-03 | 4.14E-01 | 3.86E-05 | 1.47E-01 | 3.80E-04 | 4.99E-11 | 2.41E-07 | 4.14E-01 | 2.72E-150 |
| ARS3002 | 7.20E-01 | 1.00E+00 | 5.48E-03 | 8.59E-01 | 3.86E-05 | 9.67E-02 | 3.80E-04 | 6.30E-09 | 6.26E-07 | 9.18E-01 | 2.72E-150 |
| ARS3003 | 9.94E-03 | 5.48E-03 | 1.00E+00 | 2.83E-04 | 1.41E-11 | 6.87E-06 | 1.71E-10 | 7.60E-19 | 2.83E-04 | 5.25E-04 | 2.72E-150 |
| ARS3004 | 4.14E-01 | 8.59E-01 | 2.83E-04 | 1.00E+00 | 3.80E-04 | 3.11E-01 | 1.06E-03 | 1.07E-08 | 1.80E-08 | 7.92E-01 | 2.72E-150 |
| ARS3005 | 3.86E-05 | 3.86E-05 | 1.41E-11 | 3.80E-04 | 1.00E+00 | 1.31E-02 | 4.14E-01 | 2.21E-02 | 3.43E-19 | 5.25E-04 | 2.72E-150 |
| ARS3006 | 1.47E-01 | 9.67E-02 | 6.87E-06 | 3.11E-01 | 1.31E-02 | 1.00E+00 | 4.79E-02 | 6.26E-07 | 3.86E-12 | 3.11E-01 | 1.03E-147 |
| ARS3007 | 3.80E-04 | 3.80E-04 | 1.71E-10 | 1.06E-03 | 4.14E-01 | 4.79E-02 | 1.00E+00 | 2.21E-02 | 2.14E-17 | 5.48E-03 | 4.89E-141 |
| ARS3008 | 4.99E-11 | 6.30E-09 | 7.60E-19 | 1.07E-08 | 2.21E-02 | 6.26E-07 | 2.21E-02 | 1.00E+00 | 9.90E-27 | 1.48E-07 | 4.41E-139 |
| ARS3009 | 2.41E-07 | 6.26E-07 | 2.83E-04 | 1.80E-08 | 3.43E-19 | 3.86E-12 | 2.14E-17 | 9.90E-27 | 1.00E+00 | 3.02E-08 | 2.72E-150 |
| ARS3010 | 4.14E-01 | 9.18E-01 | 5.25E-04 | 7.92E-01 | 5.25E-04 | 3.11E-01 | 5.48E-03 | 1.48E-07 | 3.02E-08 | 1.00E+00 | 1.03E-147 |
| Metagenoms | 2.72E-150 | 2.72E-150 | 2.72E-150 | 2.72E-150 | 2.72E-150 | 1.03E-147 | 4.89E-141 | 4.41E-139 | 2.72E-150 | 1.03E-147 | 1.00E+00 |

Comparison of trees using the Williams & Clifford, 1971 method

| | ARS3001 | ARS3002 | ARS3003 | ARS3004 | ARS3005 | ARS3006 | ARS3007 | ARS3008 | ARS3009 | ARS3010 | Metagenoms |
|---|---|---|---|---|---|---|---|---|---|---|---|
| ARS3001 | 1.00E+00 | 8.59E-01 | 1.00E+00 | 3.18E-01 | 1.94E-02 | 1.36E-01 | 4.30E-01 | 1.46E-03 | 1.02E-15 | 6.90E-03 | 3.85E-108 |
| ARS3002 | 8.59E-01 | 1.00E+00 | 4.94E-01 | 5.52E-01 | 1.46E-03 | 1.21E-02 | 3.18E-01 | 3.61E-05 | 3.05E-12 | 3.82E-04 | 3.99E-109 |
| ARS3003 | 1.00E+00 | 4.94E-01 | 1.00E+00 | 7.92E-01 | 2.50E-02 | 3.18E-01 | 1.99E-01 | 5.17E-04 | 2.34E-16 | 2.84E-03 | 4.25E-102 |
| ARS3004 | 3.18E-01 | 5.52E-01 | 7.92E-01 | 1.00E+00 | 1.94E-02 | 1.65E-01 | 1.58E-02 | 5.17E-04 | 1.02E-15 | 9.15E-03 | 4.29E-93 |
| ARS3005 | 1.94E-02 | 1.46E-03 | 2.50E-02 | 1.94E-02 | 1.00E+00 | 5.34E-03 | 3.82E-04 | 4.30E-01 | 2.89E-27 | 3.18E-01 | 3.04E-95 |
| ARS3006 | 1.36E-01 | 1.21E-02 | 3.18E-01 | 1.65E-01 | 5.34E-03 | 1.00E+00 | 2.38E-01 | 1.94E-02 | 2.69E-23 | 1.36E-01 | 1.41E-111 |
| ARS3007 | 4.30E-01 | 3.18E-01 | 1.99E-01 | 1.58E-02 | 3.82E-04 | 2.38E-01 | 1.00E+00 | 1.88E-04 | 2.10E-15 | 6.90E-03 | 1.08E-105 |
| ARS3008 | 1.46E-03 | 3.61E-05 | 5.17E-04 | 5.17E-04 | 4.30E-01 | 1.94E-02 | 1.88E-04 | 1.00E+00 | 4.54E-32 | 5.52E-01 | 3.85E-108 |
| ARS3009 | 1.02E-15 | 3.05E-12 | 2.34E-16 | 1.02E-15 | 2.89E-27 | 2.69E-23 | 2.10E-15 | 4.54E-32 | 1.00E+00 | 1.21E-28 | 4.41E-138 |
| ARS3010 | 6.90E-03 | 3.82E-04 | 2.84E-03 | 9.15E-03 | 3.18E-01 | 1.36E-01 | 6.90E-03 | 5.52E-01 | 1.21E-28 | 1.00E+00 | 3.85E-108 |
| Metagenoms | 3.85E-108 | 3.99E-109 | 4.25E-102 | 4.29E-93 | 3.04E-95 | 1.41E-111 | 1.08E-105 | 3.85E-108 | 4.41E-138 | 3.85E-108 | 1.00E+00 |

Comparison of trees using the Robinson & Foulds, 1981 method

| | ARS3001 | ARS3002 | ARS3003 | ARS3004 | ARS3005 | ARS3006 | ARS3007 | ARS3008 | ARS3009 | ARS3010 | Metagenoms |
|---|---|---|---|---|---|---|---|---|---|---|---|
| ARS3001 | 1.00E+00 | 7.92E-01 | 8.59E-01 | 3.47E-02 | 1.06E-01 | 1.62E-02 | 1.41E-04 | 1.01E-04 | 4.15E-13 | 2.33E-01 | 1.13E-129 |
| ARS3002 | 7.92E-01 | 1.00E+00 | 7.92E-01 | 6.73E-01 | 4.18E-02 | 4.46E-04 | 1.22E-06 | 5.04E-06 | 7.98E-13 | 2.11E-02 | 1.40E-126 |
| ARS3003 | 8.59E-01 | 7.92E-01 | 1.00E+00 | 3.47E-02 | 7.34E-01 | 4.18E-02 | 9.13E-04 | 1.01E-04 | 3.69E-18 | 4.59E-01 | 8.78E-135 |
| ARS3004 | 3.47E-02 | 6.73E-01 | 3.47E-02 | 1.00E+00 | 1.41E-04 | 1.93E-05 | 1.80E-08 | 2.90E-10 | 2.90E-10 | 1.75E-03 | 2.21E-138 |
| ARS3005 | 1.06E-01 | 4.18E-02 | 7.34E-01 | 1.41E-04 | 1.00E+00 | 2.33E-01 | 5.27E-02 | 1.75E-03 | 1.09E-23 | 5.98E-01 | 4.56E-125 |
| ARS3006 | 1.62E-02 | 4.46E-04 | 4.18E-02 | 1.93E-05 | 2.33E-01 | 1.00E+00 | 9.18E-01 | 3.32E-01 | 1.25E-21 | 7.34E-01 | 1.62E-104 |
| ARS3007 | 1.41E-04 | 1.22E-06 | 9.13E-04 | 1.80E-08 | 5.27E-02 | 9.18E-01 | 1.00E+00 | 5.98E-01 | 1.60E-24 | 1.99E-01 | 5.81E-101 |
| ARS3008 | 1.01E-04 | 5.04E-06 | 1.01E-04 | 2.90E-10 | 1.75E-03 | 3.32E-01 | 5.98E-01 | 1.00E+00 | 4.54E-32 | 4.18E-02 | 1.55E-97 |
| ARS3009 | 4.15E-13 | 7.98E-13 | 3.69E-18 | 2.90E-10 | 1.09E-23 | 1.25E-21 | 1.60E-24 | 4.54E-32 | 1.00E+00 | 8.09E-18 | 4.40E-140 |
| ARS3010 | 2.33E-01 | 2.11E-02 | 4.59E-01 | 1.75E-03 | 5.98E-01 | 7.34E-01 | 1.99E-01 | 4.18E-02 | 8.09E-18 | 1.00E+00 | 1.40E-126 |
| Metagenoms | 1.13E-129 | 1.40E-126 | 8.78E-135 | 2.21E-138 | 4.56E-125 | 1.62E-104 | 5.81E-101 | 1.55E-97 | 4.40E-140 | 1.40E-126 | 1.00E+00 |

Comparison of trees using the Kuhner & Felsenstein, 1994 method

| | ARS3001 | ARS3002 | ARS3003 | ARS3004 | ARS3005 | ARS3006 | ARS3007 | ARS3008 | ARS3009 | ARS3010 | Metagenoms |
|---|---|---|---|---|---|---|---|---|---|---|---|
| ARS3001 | 1.00E+00 | 9.98E-08 | 6.76E-12 | 6.76E-12 | 1.37E-02 | 3.92E-04 | 2.18E-01 | 1.24E-01 | 4.85E-01 | 2.68E-05 | 1.82E-50 |
| ARS3002 | 9.98E-08 | 1.00E+00 | 1.47E-01 | 2.99E-01 | 4.19E-03 | 6.29E-02 | 9.02E-11 | 1.36E-04 | 3.86E-12 | 5.52E-01 | 1.96E-29 |
| ARS3003 | 6.76E-12 | 1.47E-01 | 1.00E+00 | 1.00E+00 | 2.68E-05 | 1.91E-04 | 2.91E-15 | 1.12E-06 | 6.24E-17 | 3.88E-02 | 9.88E-19 |
| ARS3004 | 6.76E-12 | 2.99E-01 | 1.00E+00 | 1.00E+00 | 3.99E-05 | 8.05E-04 | 5.86E-15 | 4.24E-07 | 1.30E-16 | 3.04E-02 | 4.94E-21 |
| ARS3005 | 1.37E-02 | 4.19E-03 | 2.68E-05 | 3.99E-05 | 1.00E+00 | 1.47E-01 | 3.92E-04 | 6.22E-01 | 1.91E-04 | 2.53E-01 | 1.02E-37 |
| ARS3006 | 3.92E-04 | 6.29E-02 | 1.91E-04 | 8.05E-04 | 1.47E-01 | 1.00E+00 | 1.77E-06 | 3.09E-03 | 4.24E-07 | 2.53E-01 | 4.01E-41 |
| ARS3007 | 2.18E-01 | 9.02E-11 | 2.91E-15 | 5.86E-15 | 3.92E-04 | 1.77E-06 | 1.00E+00 | 1.05E-02 | 1.00E+00 | 2.76E-07 | 6.59E-61 |
| ARS3008 | 1.24E-01 | 1.36E-04 | 1.12E-06 | 4.24E-07 | 6.22E-01 | 3.09E-03 | 1.05E-02 | 1.00E+00 | 1.12E-03 | 2.37E-02 | 5.71E-46 |
| ARS3009 | 4.85E-01 | 3.86E-12 | 6.24E-17 | 1.30E-16 | 1.91E-04 | 4.24E-07 | 1.00E+00 | 1.12E-03 | 1.00E+00 | 1.13E-08 | 1.15E-59 |
| ARS3010 | 2.68E-05 | 5.52E-01 | 3.88E-02 | 3.04E-02 | 2.53E-01 | 2.53E-01 | 2.76E-07 | 2.37E-02 | 1.13E-08 | 1.00E+00 | 7.35E-31 |
| Metagenoms | 1.82E-50 | 1.96E-29 | 9.88E-19 | 4.94E-21 | 1.02E-37 | 4.01E-41 | 6.59E-61 | 5.71E-46 | 1.15E-59 | 7.35E-31 | 1.00E+00 |

**Supplementary Table S3C:** Pairwise FDR corrected p-values of the Kolmogorow-Smirnow test on the distances between phylogenetic trees for 16 non-ribosomal proteins in 10 samples of archaeal RefSeq genomes vs one sample containing archaeal MAGs (associated with Extended Data 1c)

Comparison of trees using the Nye et al., 2006 method

| | ARS3001 | ARS3002 | ARS3003 | ARS3004 | ARS3005 | ARS3006 | ARS3007 | ARS3008 | ARS3009 | ARS3010 | Metagenomes |
|---|---|---|---|---|---|---|---|---|---|---|---|
| ARS3001 | 1.00E+00 | 3.29E-03 | 3.71E-01 | 5.33E-02 | 5.32E-06 | 3.84E-02 | 7.95E-04 | 1.27E-03 | 2.42E-01 | 8.81E-01 | 3.19E-68 |
| ARS3002 | 3.29E-03 | 1.00E+00 | 1.09E-04 | 7.46E-02 | 5.33E-02 | 3.84E-02 | 3.71E-01 | 1.36E-01 | 7.95E-04 | 7.95E-04 | 3.19E-68 |
| ARS3003 | 3.71E-01 | 1.09E-04 | 1.00E+00 | 1.91E-04 | 5.66E-09 | 1.09E-04 | 1.33E-06 | 3.62E-05 | 3.71E-01 | 5.74E-01 | 3.19E-68 |
| ARS3004 | 5.33E-02 | 7.46E-02 | 1.91E-04 | 1.00E+00 | 1.27E-03 | 8.81E-01 | 1.36E-01 | 2.81E-02 | 7.95E-04 | 7.97E-03 | 3.19E-68 |
| ARS3005 | 5.32E-06 | 5.33E-02 | 5.66E-09 | 1.27E-03 | 1.00E+00 | 2.06E-03 | 3.71E-01 | 3.12E-01 | 3.10E-07 | 1.52E-07 | 3.19E-68 |
| ARS3006 | 3.84E-02 | 3.84E-02 | 1.09E-04 | 8.81E-01 | 2.06E-03 | 1.00E+00 | 1.03E-01 | 3.84E-02 | 5.34E-04 | 5.16E-03 | 3.19E-68 |
| ARS3007 | 7.95E-04 | 3.71E-01 | 1.33E-06 | 1.36E-01 | 3.71E-01 | 1.03E-01 | 1.00E+00 | 8.81E-01 | 6.50E-05 | 1.98E-05 | 3.19E-68 |
| ARS3008 | 1.27E-03 | 1.36E-01 | 3.62E-05 | 2.81E-02 | 3.12E-01 | 3.84E-02 | 8.81E-01 | 1.00E+00 | 5.34E-04 | 5.34E-04 | 3.19E-68 |
| ARS3009 | 2.42E-01 | 7.95E-04 | 3.71E-01 | 7.95E-04 | 3.10E-07 | 5.34E-04 | 6.50E-05 | 5.34E-04 | 1.00E+00 | 6.84E-01 | 3.19E-68 |
| ARS3010 | 8.81E-01 | 7.95E-04 | 5.74E-01 | 7.97E-03 | 1.52E-07 | 5.16E-03 | 1.98E-05 | 5.34E-04 | 6.84E-01 | 1.00E+00 | 3.19E-68 |
| Metagenomes | 3.19E-68 | 3.19E-68 | 3.19E-68 | 3.19E-68 | 3.19E-68 | 3.19E-68 | 3.19E-68 | 3.19E-68 | 3.19E-68 | 3.19E-68 | 1.00E+00 |

Comparison of trees using the Williams & Clifford, 1971 method

| | ARS3001 | ARS3002 | ARS3003 | ARS3004 | ARS3005 | ARS3006 | ARS3007 | ARS3008 | ARS3009 | ARS3010 | Metagenomes |
|---|---|---|---|---|---|---|---|---|---|---|---|
| ARS3001 | 1.00E+00 | 1.13E-01 | 5.63E-01 | 2.17E-02 | 1.55E-06 | 2.17E-02 | 2.98E-02 | 2.51E-04 | 2.99E-01 | 6.09E-02 | 1.91E-65 |
| ARS3002 | 1.13E-01 | 1.00E+00 | 1.13E-01 | 6.59E-01 | 3.36E-03 | 6.59E-01 | 4.71E-01 | 1.33E-01 | 1.57E-02 | 7.66E-01 | 3.79E-64 |
| ARS3003 | 5.63E-01 | 1.13E-01 | 1.00E+00 | 8.49E-02 | 4.39E-05 | 1.57E-02 | 2.98E-02 | 1.24E-03 | 1.33E-01 | 1.77E-01 | 3.79E-64 |
| ARS3004 | 2.17E-02 | 6.59E-01 | 8.49E-02 | 1.00E+00 | 2.44E-05 | 1.33E-01 | 1.33E-01 | 7.74E-03 | 4.25E-04 | 2.99E-01 | 1.91E-65 |
| ARS3005 | 1.55E-06 | 3.36E-03 | 4.39E-05 | 2.44E-05 | 1.00E+00 | 1.57E-02 | 1.18E-02 | 6.09E-02 | 3.66E-07 | 1.57E-02 | 3.23E-56 |
| ARS3006 | 2.17E-02 | 6.59E-01 | 1.57E-02 | 1.33E-01 | 1.57E-02 | 1.00E+00 | 1.33E-01 | 5.63E-01 | 7.74E-03 | 3.79E-01 | 3.79E-64 |
| ARS3007 | 2.98E-02 | 4.71E-01 | 2.98E-02 | 1.33E-01 | 1.18E-02 | 1.33E-01 | 1.00E+00 | 2.98E-02 | 2.17E-02 | 1.33E-01 | 3.79E-64 |
| ARS3008 | 2.51E-04 | 1.33E-01 | 1.24E-03 | 7.74E-03 | 6.09E-02 | 5.63E-01 | 2.98E-02 | 1.00E+00 | 1.46E-04 | 1.33E-01 | 1.99E-62 |
| ARS3009 | 2.99E-01 | 1.57E-02 | 1.33E-01 | 4.25E-04 | 3.66E-07 | 7.74E-03 | 2.17E-02 | 1.46E-04 | 1.00E+00 | 7.74E-03 | 3.79E-64 |
| ARS3010 | 6.09E-02 | 7.66E-01 | 1.77E-01 | 2.99E-01 | 1.57E-02 | 3.79E-01 | 1.33E-01 | 1.33E-01 | 7.74E-03 | 1.00E+00 | 3.79E-64 |
| Metagenomes | 1.91E-65 | 3.79E-64 | 3.79E-64 | 1.91E-65 | 3.23E-56 | 3.79E-64 | 3.79E-64 | 1.99E-62 | 3.79E-64 | 3.79E-64 | 1.00E+00 |

Comparison of trees using the Robinson & Foulds, 1981 method

| | ARS3001 | ARS3002 | ARS3003 | ARS3004 | ARS3005 | ARS3006 | ARS3007 | ARS3008 | ARS3009 | ARS3010 | Metagenomes |
|---|---|---|---|---|---|---|---|---|---|---|---|
| ARS3001 | 1.00E+00 | 3.51E-02 | 5.35E-01 | 4.87E-02 | 6.74E-04 | 3.42E-01 | 1.73E-03 | 1.33E-06 | 2.41E-02 | 2.79E-01 | 5.45E-66 |
| ARS3002 | 3.51E-02 | 1.00E+00 | 9.50E-01 | 2.26E-01 | 4.31E-01 | 6.36E-01 | 4.87E-02 | 7.10E-03 | 5.32E-06 | 1.00E+00 | 5.45E-66 |
| ARS3003 | 5.35E-01 | 9.50E-01 | 1.00E+00 | 2.79E-01 | 1.81E-01 | 7.41E-01 | 1.04E-02 | 1.73E-03 | 1.73E-03 | 1.00E+00 | 5.45E-66 |
| ARS3004 | 4.87E-02 | 2.26E-01 | 2.79E-01 | 1.00E+00 | 9.50E-01 | 1.00E+00 | 9.50E-01 | 1.35E-01 | 1.98E-05 | 2.26E-01 | 3.37E-64 |
| ARS3005 | 6.74E-04 | 4.31E-01 | 1.81E-01 | 9.50E-01 | 1.00E+00 | 2.79E-01 | 1.00E+00 | 6.36E-01 | 1.24E-08 | 2.26E-01 | 1.79E-62 |
| ARS3006 | 3.42E-01 | 6.36E-01 | 7.41E-01 | 1.00E+00 | 2.79E-01 | 1.00E+00 | 3.42E-01 | 1.04E-02 | 4.25E-04 | 3.42E-01 | 5.45E-66 |
| ARS3007 | 1.73E-03 | 4.87E-02 | 1.04E-02 | 9.50E-01 | 1.00E+00 | 3.42E-01 | 1.00E+00 | 7.41E-01 | 6.20E-08 | 9.86E-02 | 5.45E-66 |
| ARS3008 | 1.33E-06 | 7.10E-03 | 1.73E-03 | 1.35E-01 | 6.36E-01 | 1.04E-02 | 7.41E-01 | 1.00E+00 | 8.69E-13 | 7.10E-03 | 3.37E-64 |
| ARS3009 | 2.41E-02 | 5.32E-06 | 1.73E-03 | 1.98E-05 | 1.24E-08 | 4.25E-04 | 6.20E-08 | 8.69E-13 | 1.00E+00 | 6.74E-04 | 5.45E-66 |
| ARS3010 | 2.79E-01 | 1.00E+00 | 1.00E+00 | 2.26E-01 | 2.26E-01 | 3.42E-01 | 9.86E-02 | 7.10E-03 | 6.74E-04 | 1.00E+00 | 5.45E-66 |
| Metagenomes | 5.45E-66 | 5.45E-66 | 5.45E-66 | 3.37E-64 | 1.79E-62 | 5.45E-66 | 5.45E-66 | 3.37E-64 | 5.45E-66 | 5.45E-66 | 1.00E+00 |

Comparison of trees using the Kuhner & Felsenstein, 1994 method

| | ARS3001 | ARS3002 | ARS3003 | ARS3004 | ARS3005 | ARS3006 | ARS3007 | ARS3008 | ARS3009 | ARS3010 | Metagenomes |
|---|---|---|---|---|---|---|---|---|---|---|---|
| ARS3001 | 1.00E+00 | 3.99E-03 | 3.78E-02 | 2.02E-02 | 8.54E-04 | 3.19E-01 | 5.41E-02 | 3.19E-01 | 4.90E-01 | 7.94E-07 | 2.12E-60 |
| ARS3002 | 3.99E-03 | 1.00E+00 | 1.13E-01 | 3.19E-01 | 8.49E-02 | 8.96E-03 | 3.78E-02 | 1.13E-01 | 8.49E-02 | 1.36E-02 | 3.32E-59 |
| ARS3003 | 3.78E-02 | 1.13E-01 | 1.00E+00 | 6.84E-01 | 7.95E-01 | 8.49E-02 | 1.49E-01 | 8.49E-02 | 6.19E-03 | 7.10E-02 | 3.32E-59 |
| ARS3004 | 2.02E-02 | 3.19E-01 | 6.84E-01 | 1.00E+00 | 8.98E-01 | 3.19E-01 | 3.95E-01 | 7.10E-02 | 2.83E-02 | 7.10E-02 | 3.32E-59 |
| ARS3005 | 8.54E-04 | 8.49E-02 | 7.95E-01 | 8.98E-01 | 1.00E+00 | 8.49E-02 | 3.19E-01 | 3.78E-02 | 2.83E-02 | 1.49E-01 | 2.12E-60 |
| ARS3006 | 3.19E-01 | 8.96E-03 | 8.49E-02 | 3.19E-01 | 8.49E-02 | 1.00E+00 | 9.79E-01 | 5.74E-01 | 3.95E-01 | 8.54E-04 | 3.03E-63 |
| ARS3007 | 5.41E-02 | 3.78E-02 | 1.49E-01 | 3.95E-01 | 3.19E-01 | 9.79E-01 | 1.00E+00 | 5.74E-01 | 5.74E-01 | 8.96E-03 | 2.12E-60 |
| ARS3008 | 3.19E-01 | 1.13E-01 | 8.49E-02 | 7.10E-02 | 3.78E-02 | 5.74E-01 | 5.74E-01 | 1.00E+00 | 8.49E-02 | 3.09E-04 | 3.32E-59 |
| ARS3009 | 4.90E-01 | 8.49E-02 | 6.19E-03 | 2.83E-02 | 2.83E-02 | 3.95E-01 | 5.74E-01 | 8.49E-02 | 1.00E+00 | 5.12E-05 | 3.32E-59 |
| ARS3010 | 7.94E-07 | 1.36E-02 | 7.10E-02 | 7.10E-02 | 1.49E-01 | 8.54E-04 | 8.96E-03 | 3.09E-04 | 5.12E-05 | 1.00E+00 | 3.32E-59 |
| Metagenomes | 2.12E-60 | 3.32E-59 | 3.32E-59 | 3.32E-59 | 2.12E-60 | 3.03E-63 | 2.12E-60 | 3.32E-59 | 3.32E-59 | 3.32E-59 | 1.00E+00 |

**Supplementary Table S3D:** Pairwise FDR corrected p-values of the Kolmogorow-Smirnow test on the distances between phylogenetic trees for ribosomal proteins in 10 samples of bacterial RefSeq genomes vs one sample containing CPR MAGs (associated with Extended Data 1d)

Comparison of trees using the Nye et al., 2006 method

| | BRS3001 | BRS3002 | BRS3003 | BRS3004 | BRS3005 | BRS3006 | BRS3007 | BRS3008 | BRS3009 | BRS3010 | Metagenomes |
|---|---|---|---|---|---|---|---|---|---|---|---|
| BRS3001 | 1.00E+00 | 7.95E-01 | 5.92E-02 | 4.87E-11 | 3.71E-01 | 5.34E-04 | 1.03E-01 | 2.98E-02 | 2.32E-01 | 5.57E-05 | 2.01E-51 |
| BRS3002 | 7.95E-01 | 1.00E+00 | 1.03E-01 | 1.19E-10 | 3.71E-01 | 5.57E-05 | 1.36E-01 | 2.10E-02 | 7.84E-02 | 9.94E-05 | 1.76E-42 |
| BRS3003 | 5.92E-02 | 1.03E-01 | 1.00E+00 | 1.76E-05 | 3.89E-03 | 1.03E-01 | 6.98E-01 | 1.00E+00 | 2.32E-01 | 5.92E-02 | 1.62E-41 |
| BRS3004 | 4.87E-11 | 1.19E-10 | 1.76E-05 | 1.00E+00 | 9.40E-16 | 2.10E-02 | 1.76E-05 | 5.57E-05 | 2.50E-08 | 4.30E-02 | 1.07E-38 |
| BRS3005 | 3.71E-01 | 3.71E-01 | 3.89E-03 | 9.40E-16 | 1.00E+00 | 5.03E-08 | 1.52E-03 | 1.74E-04 | 7.84E-02 | 5.03E-08 | 1.11E-56 |
| BRS3006 | 5.34E-04 | 5.57E-05 | 1.03E-01 | 2.10E-02 | 5.03E-08 | 1.00E+00 | 2.98E-02 | 2.32E-01 | 9.68E-03 | 8.98E-01 | 1.07E-38 |
| BRS3007 | 1.03E-01 | 1.36E-01 | 6.98E-01 | 1.76E-05 | 1.52E-03 | 2.98E-02 | 1.00E+00 | 7.95E-01 | 1.36E-01 | 1.46E-02 | 1.46E-40 |
| BRS3008 | 2.98E-02 | 2.10E-02 | 1.00E+00 | 5.57E-05 | 1.74E-04 | 2.32E-01 | 7.95E-01 | 1.00E+00 | 2.99E-01 | 7.84E-02 | 1.76E-42 |
| BRS3009 | 2.32E-01 | 7.84E-02 | 2.32E-01 | 2.50E-08 | 7.84E-02 | 9.68E-03 | 1.36E-01 | 2.99E-01 | 1.00E+00 | 3.89E-03 | 4.49E-49 |
| BRS3010 | 5.57E-05 | 9.94E-05 | 5.92E-02 | 4.30E-02 | 5.03E-08 | 8.98E-01 | 1.46E-02 | 7.84E-02 | 3.89E-03 | 1.00E+00 | 6.22E-36 |
| Metagenomes | 2.01E-51 | 1.76E-42 | 1.62E-41 | 1.07E-38 | 1.11E-56 | 1.07E-38 | 1.46E-40 | 1.76E-42 | 4.49E-49 | 6.22E-36 | 1.00E+00 |

Comparison of trees using the Williams & Clifford, 1971 method

| | BRS3001 | BRS3002 | BRS3003 | BRS3004 | BRS3005 | BRS3006 | BRS3007 | BRS3008 | BRS3009 | BRS3010 | Metagenomes |
|---|---|---|---|---|---|---|---|---|---|---|---|
| BRS3001 | 1.00E+00 | 7.41E-01 | 2.09E-01 | 6.40E-04 | 6.87E-02 | 5.35E-01 | 8.43E-01 | 1.00E+00 | 1.66E-01 | 6.40E-04 | 3.89E-23 |
| BRS3002 | 7.41E-01 | 1.00E+00 | 3.59E-01 | 1.86E-05 | 4.31E-01 | 1.66E-01 | 7.41E-01 | 9.98E-01 | 6.50E-01 | 1.73E-03 | 1.03E-23 |
| BRS3003 | 2.09E-01 | 3.59E-01 | 1.00E+00 | 6.24E-07 | 9.98E-01 | 6.81E-03 | 2.09E-01 | 2.09E-01 | 9.98E-01 | 4.98E-06 | 1.37E-32 |
| BRS3004 | 6.40E-04 | 1.86E-05 | 6.24E-07 | 1.00E+00 | 5.66E-09 | 2.25E-02 | 1.06E-03 | 6.50E-05 | 2.92E-08 | 1.66E-01 | 1.06E-12 |
| BRS3005 | 6.87E-02 | 4.31E-01 | 9.98E-01 | 5.66E-09 | 1.00E+00 | 4.38E-03 | 4.87E-02 | 1.66E-01 | 8.43E-01 | 6.24E-07 | 4.93E-34 |
| BRS3006 | 5.35E-01 | 1.66E-01 | 6.81E-03 | 2.25E-02 | 4.38E-03 | 1.00E+00 | 7.41E-01 | 4.31E-01 | 2.25E-02 | 2.25E-02 | 8.56E-18 |
| BRS3007 | 8.43E-01 | 7.41E-01 | 2.09E-01 | 1.06E-03 | 4.87E-02 | 7.41E-01 | 1.00E+00 | 9.98E-01 | 1.66E-01 | 1.63E-02 | 2.49E-21 |
| BRS3008 | 1.00E+00 | 9.98E-01 | 2.09E-01 | 6.50E-05 | 1.66E-01 | 4.31E-01 | 9.98E-01 | 1.00E+00 | 4.31E-01 | 2.77E-03 | 6.68E-22 |
| BRS3009 | 1.66E-01 | 6.50E-01 | 9.98E-01 | 2.92E-08 | 8.43E-01 | 2.25E-02 | 1.66E-01 | 4.31E-01 | 1.00E+00 | 2.59E-06 | 4.36E-31 |
| BRS3010 | 6.40E-04 | 1.73E-03 | 4.98E-06 | 1.66E-01 | 6.24E-07 | 2.25E-02 | 1.63E-02 | 2.77E-03 | 2.59E-06 | 1.00E+00 | 4.02E-10 |
| Metagenomes | 3.89E-23 | 1.03E-23 | 1.37E-32 | 1.06E-12 | 4.93E-34 | 8.56E-18 | 2.49E-21 | 6.68E-22 | 4.36E-31 | 4.02E-10 | 1.00E+00 |

Comparison of trees using the Robinson & Foulds, 1981 method

| | BRS3001 | BRS3002 | BRS3003 | BRS3004 | BRS3005 | BRS3006 | BRS3007 | BRS3008 | BRS3009 | BRS3010 | Metagenomes |
|---|---|---|---|---|---|---|---|---|---|---|---|
| BRS3001 | 1.00E+00 | 3.12E-01 | 7.84E-02 | 1.53E-18 | 1.00E+00 | 9.30E-07 | 1.08E-01 | 5.32E-05 | 5.97E-01 | 9.57E-08 | 2.01E-51 |
| BRS3002 | 3.12E-01 | 1.00E+00 | 1.89E-01 | 1.03E-10 | 1.42E-01 | 1.31E-03 | 1.00E+00 | 2.81E-02 | 7.95E-01 | 1.54E-04 | 1.76E-42 |
| BRS3003 | 7.84E-02 | 1.89E-01 | 1.00E+00 | 8.77E-09 | 2.81E-02 | 5.76E-02 | 1.42E-01 | 2.48E-01 | 4.90E-01 | 5.68E-03 | 1.62E-41 |
| BRS3004 | 1.53E-18 | 1.03E-10 | 8.77E-09 | 1.00E+00 | 2.78E-20 | 8.47E-03 | 1.97E-12 | 9.11E-05 | 1.97E-12 | 8.47E-03 | 3.28E-29 |
| BRS3005 | 1.00E+00 | 1.42E-01 | 2.81E-02 | 2.78E-20 | 1.00E+00 | 9.57E-08 | 7.84E-02 | 2.93E-05 | 3.95E-01 | 8.77E-09 | 3.23E-55 |
| BRS3006 | 9.30E-07 | 1.31E-03 | 5.76E-02 | 8.47E-03 | 9.57E-08 | 1.00E+00 | 4.74E-04 | 7.95E-01 | 1.31E-03 | 7.95E-01 | 1.07E-36 |
| BRS3007 | 1.08E-01 | 1.00E+00 | 1.42E-01 | 1.97E-12 | 7.84E-02 | 4.74E-04 | 1.00E+00 | 5.76E-02 | 7.11E-01 | 1.54E-04 | 1.89E-43 |
| BRS3008 | 5.32E-05 | 2.81E-02 | 2.48E-01 | 9.11E-05 | 2.93E-05 | 7.95E-01 | 5.76E-02 | 1.00E+00 | 2.81E-02 | 3.12E-01 | 3.05E-33 |
| BRS3009 | 5.97E-01 | 7.95E-01 | 4.90E-01 | 1.97E-12 | 3.95E-01 | 1.31E-03 | 7.11E-01 | 2.81E-02 | 1.00E+00 | 9.11E-05 | 1.10E-46 |
| BRS3010 | 9.57E-08 | 1.54E-04 | 5.68E-03 | 8.47E-03 | 8.77E-09 | 7.95E-01 | 1.54E-04 | 3.12E-01 | 9.11E-05 | 1.00E+00 | 3.05E-33 |
| Metagenomes | 2.01E-51 | 1.76E-42 | 1.62E-41 | 3.28E-29 | 3.23E-55 | 1.07E-36 | 1.89E-43 | 3.05E-33 | 1.10E-46 | 3.05E-33 | 1.00E+00 |

Comparison of trees using the Kuhner & Felsenstein, 1994 method

| | BRS3001 | BRS3002 | BRS3003 | BRS3004 | BRS3005 | BRS3006 | BRS3007 | BRS3008 | BRS3009 | BRS3010 | Metagenomes |
|---|---|---|---|---|---|---|---|---|---|---|---|
| BRS3001 | 1.00E+00 | 3.40E-02 | 4.03E-05 | 2.49E-13 | 1.27E-09 | 1.27E-05 | 4.03E-05 | 3.40E-02 | 6.36E-04 | 3.79E-01 | 2.01E-51 |
| BRS3002 | 3.40E-02 | 1.00E+00 | 3.05E-01 | 4.16E-07 | 2.36E-05 | 4.84E-02 | 2.42E-01 | 8.46E-07 | 3.05E-01 | 2.52E-02 | 5.48E-47 |
| BRS3003 | 4.03E-05 | 3.05E-01 | 1.00E+00 | 1.34E-04 | 2.84E-03 | 5.85E-01 | 8.81E-01 | 6.37E-13 | 7.81E-01 | 4.03E-05 | 4.36E-48 |
| BRS3004 | 2.49E-13 | 4.16E-07 | 1.34E-04 | 1.00E+00 | 6.71E-01 | 6.36E-04 | 6.36E-04 | 1.59E-21 | 4.00E-04 | 6.37E-13 | 6.91E-36 |
| BRS3005 | 1.27E-09 | 2.36E-05 | 2.84E-03 | 6.71E-01 | 1.00E+00 | 3.40E-02 | 2.84E-03 | 2.08E-17 | 1.71E-02 | 9.11E-11 | 4.93E-35 |
| BRS3006 | 1.27E-05 | 4.84E-02 | 5.85E-01 | 6.36E-04 | 3.40E-02 | 1.00E+00 | 6.71E-01 | 1.72E-12 | 3.79E-01 | 1.69E-06 | 1.08E-43 |
| BRS3007 | 4.03E-05 | 2.42E-01 | 8.81E-01 | 6.36E-04 | 2.84E-03 | 6.71E-01 | 1.00E+00 | 2.23E-10 | 6.71E-01 | 6.69E-06 | 4.36E-48 |
| BRS3008 | 3.40E-02 | 8.46E-07 | 6.37E-13 | 1.59E-21 | 2.08E-17 | 1.72E-12 | 2.23E-10 | 1.00E+00 | 1.27E-09 | 3.40E-02 | 8.47E-60 |
| BRS3009 | 6.36E-04 | 3.05E-01 | 7.81E-01 | 4.00E-04 | 1.71E-02 | 3.79E-01 | 6.71E-01 | 1.27E-09 | 1.00E+00 | 2.33E-04 | 1.28E-40 |
| BRS3010 | 3.79E-01 | 2.52E-02 | 4.03E-05 | 6.37E-13 | 9.11E-11 | 1.69E-06 | 6.69E-06 | 3.40E-02 | 2.33E-04 | 1.00E+00 | 2.56E-50 |
| Metagenomes | 2.01E-51 | 5.48E-47 | 4.36E-48 | 6.91E-36 | 4.93E-35 | 1.08E-43 | 4.36E-48 | 8.47E-60 | 1.28E-40 | 2.56E-50 | 1.00E+00 |

**Supplementary Table S3E:** Pairwise FDR corrected p-values of the Kolmogorow-Smirnow test on the distances between phylogenetic trees for 20 ribosomal proteins in 10 samples of bacterial RefSeq genomes vs one sample containing bacterial MAGs (associated with Extended Data 1e)

Comparison of trees using the Nye et al., 2006 method

| | BRS3001 | BRS3002 | BRS3003 | BRS3004 | BRS3005 | BRS3006 | BRS3007 | BRS3008 | BRS3009 | BRS3010 | Metagenomes |
|---|---|---|---|---|---|---|---|---|---|---|---|
| BRS3001 | 1.00E+00 | 1.73E-03 | 3.20E-02 | 3.13E-15 | 3.58E-03 | 6.95E-03 | 1.27E-02 | 2.50E-03 | 9.46E-01 | 5.19E-04 | 3.13E-15 |
| BRS3002 | 1.73E-03 | 1.00E+00 | 9.18E-07 | 1.79E-24 | 8.92E-01 | 1.73E-07 | 9.18E-07 | 4.66E-01 | 6.95E-03 | 8.46E-09 | 3.13E-15 |
| BRS3003 | 3.20E-02 | 9.18E-07 | 1.00E+00 | 2.71E-08 | 9.18E-07 | 8.25E-01 | 9.46E-01 | 2.19E-04 | 1.27E-02 | 6.45E-01 | 3.13E-15 |
| BRS3004 | 3.13E-15 | 1.79E-24 | 2.71E-08 | 1.00E+00 | 1.92E-27 | 4.45E-06 | 4.55E-09 | 3.02E-19 | 3.13E-15 | 1.57E-06 | 3.13E-15 |
| BRS3005 | 3.58E-03 | 8.92E-01 | 9.18E-07 | 1.92E-27 | 1.00E+00 | 2.71E-08 | 2.66E-06 | 2.06E-01 | 1.27E-02 | 1.56E-08 | 3.13E-15 |
| BRS3006 | 6.95E-03 | 1.73E-07 | 8.25E-01 | 4.45E-06 | 2.71E-08 | 1.00E+00 | 8.92E-01 | 5.42E-05 | 5.08E-03 | 1.00E+00 | 3.13E-15 |
| BRS3007 | 1.27E-02 | 9.18E-07 | 9.46E-01 | 4.55E-09 | 2.66E-06 | 8.92E-01 | 1.00E+00 | 1.22E-05 | 3.20E-02 | 7.36E-01 | 3.13E-15 |
| BRS3008 | 2.50E-03 | 4.66E-01 | 2.19E-04 | 3.02E-19 | 2.06E-01 | 5.42E-05 | 1.22E-05 | 1.00E+00 | 9.67E-03 | 7.39E-06 | 3.13E-15 |
| BRS3009 | 9.46E-01 | 6.95E-03 | 1.27E-02 | 3.13E-15 | 1.27E-02 | 5.08E-03 | 3.20E-02 | 9.67E-03 | 1.00E+00 | 7.75E-04 | 3.13E-15 |
| BRS3010 | 5.19E-04 | 8.46E-09 | 6.45E-01 | 1.57E-06 | 1.56E-08 | 1.00E+00 | 7.36E-01 | 7.39E-06 | 7.75E-04 | 1.00E+00 | 3.13E-15 |
| Metagenomes | 3.13E-15 | 3.13E-15 | 3.13E-15 | 3.13E-15 | 3.13E-15 | 3.13E-15 | 3.13E-15 | 3.13E-15 | 3.13E-15 | 3.13E-15 | 1.00E+00 |

Comparison of trees using the Williams & Clifford, 1971 method

| | BRS3001 | BRS3002 | BRS3003 | BRS3004 | BRS3005 | BRS3006 | BRS3007 | BRS3008 | BRS3009 | BRS3010 | Metagenomes |
|---|---|---|---|---|---|---|---|---|---|---|---|
| BRS3001 | 1.00E+00 | 1.05E-01 | 5.96E-01 | 1.36E-11 | 1.05E-01 | 9.14E-04 | 7.79E-01 | 1.33E-01 | 1.00E+00 | 6.15E-04 | 4.27E-15 |
| BRS3002 | 1.05E-01 | 1.00E+00 | 8.22E-03 | 1.82E-07 | 6.77E-05 | 2.12E-02 | 5.14E-01 | 4.30E-03 | 8.61E-02 | 1.05E-01 | 4.27E-15 |
| BRS3003 | 5.96E-01 | 8.22E-03 | 1.00E+00 | 4.27E-15 | 2.53E-01 | 3.14E-06 | 1.66E-01 | 2.53E-01 | 3.77E-01 | 5.23E-06 | 4.27E-15 |
| BRS3004 | 1.36E-11 | 1.82E-07 | 4.27E-15 | 1.00E+00 | 1.07E-19 | 8.22E-03 | 1.12E-09 | 7.41E-17 | 3.12E-12 | 2.02E-03 | 3.12E-12 |
| BRS3005 | 1.05E-01 | 6.77E-05 | 2.53E-01 | 1.07E-19 | 1.00E+00 | 4.28E-09 | 1.56E-02 | 5.14E-01 | 3.79E-02 | 7.99E-09 | 4.27E-15 |
| BRS3006 | 9.14E-04 | 2.12E-02 | 3.14E-06 | 8.22E-03 | 4.28E-09 | 1.00E+00 | 8.22E-03 | 1.82E-07 | 1.70E-04 | 9.46E-01 | 5.04E-15 |
| BRS3007 | 7.79E-01 | 5.14E-01 | 1.66E-01 | 1.12E-09 | 1.56E-02 | 8.22E-03 | 1.00E+00 | 3.79E-02 | 4.47E-01 | 1.56E-02 | 4.27E-15 |
| BRS3008 | 1.33E-01 | 4.30E-03 | 2.53E-01 | 7.41E-17 | 5.14E-01 | 1.82E-07 | 3.79E-02 | 1.00E+00 | 8.61E-02 | 5.82E-07 | 4.27E-15 |
| BRS3009 | 1.00E+00 | 8.61E-02 | 3.77E-01 | 3.12E-12 | 3.79E-02 | 1.70E-04 | 4.47E-01 | 8.61E-02 | 1.00E+00 | 4.10E-04 | 4.27E-15 |
| BRS3010 | 6.15E-04 | 1.05E-01 | 5.23E-06 | 2.02E-03 | 7.99E-09 | 9.46E-01 | 1.56E-02 | 5.82E-07 | 4.10E-04 | 1.00E+00 | 4.27E-15 |
| Metagenomes | 4.27E-15 | 4.27E-15 | 4.27E-15 | 3.12E-12 | 4.27E-15 | 5.04E-15 | 4.27E-15 | 4.27E-15 | 4.27E-15 | 4.27E-15 | 1.00E+00 |

Comparison of trees using the Robinson & Foulds, 1981 method

| | BRS3001 | BRS3002 | BRS3003 | BRS3004 | BRS3005 | BRS3006 | BRS3007 | BRS3008 | BRS3009 | BRS3010 | Metagenomes |
|---|---|---|---|---|---|---|---|---|---|---|---|
| BRS3001 | 1.00E+00 | 9.64E-01 | 2.51E-04 | 1.20E-20 | 2.26E-01 | 1.06E-04 | 5.73E-04 | 7.21E-01 | 7.21E-01 | 9.86E-06 | 3.13E-15 |
| BRS3002 | 9.64E-01 | 1.00E+00 | 7.75E-04 | 1.04E-19 | 1.00E+00 | 1.64E-04 | 1.73E-03 | 7.21E-01 | 4.87E-01 | 4.11E-05 | 3.13E-15 |
| BRS3003 | 2.51E-04 | 7.75E-04 | 1.00E+00 | 2.18E-06 | 9.86E-06 | 1.00E+00 | 6.59E-01 | 3.68E-03 | 4.99E-02 | 2.26E-01 | 3.13E-15 |
| BRS3004 | 1.20E-20 | 1.04E-19 | 2.18E-06 | 1.00E+00 | 1.80E-20 | 6.33E-06 | 7.64E-07 | 9.87E-17 | 3.13E-15 | 1.53E-05 | 3.13E-15 |
| BRS3005 | 2.26E-01 | 1.00E+00 | 9.86E-06 | 1.80E-20 | 1.00E+00 | 1.30E-06 | 7.75E-04 | 7.94E-01 | 4.99E-02 | 1.53E-05 | 3.13E-15 |
| BRS3006 | 1.06E-04 | 1.64E-04 | 1.00E+00 | 6.33E-06 | 1.30E-06 | 1.00E+00 | 3.35E-01 | 7.75E-04 | 2.85E-02 | 3.35E-01 | 3.13E-15 |
| BRS3007 | 5.73E-04 | 1.73E-03 | 6.59E-01 | 7.64E-07 | 7.75E-04 | 3.35E-01 | 1.00E+00 | 2.12E-02 | 8.61E-02 | 5.70E-01 | 3.13E-15 |
| BRS3008 | 7.21E-01 | 7.21E-01 | 3.68E-03 | 9.87E-17 | 7.94E-01 | 7.75E-04 | 2.12E-02 | 1.00E+00 | 7.94E-01 | 7.75E-04 | 3.13E-15 |
| BRS3009 | 7.21E-01 | 4.87E-01 | 4.99E-02 | 3.13E-15 | 4.99E-02 | 2.85E-02 | 8.61E-02 | 7.94E-01 | 1.00E+00 | 5.73E-04 | 3.13E-15 |
| BRS3010 | 9.86E-06 | 4.11E-05 | 2.26E-01 | 1.53E-05 | 1.53E-05 | 3.35E-01 | 5.70E-01 | 7.75E-04 | 5.73E-04 | 1.00E+00 | 3.13E-15 |
| Metagenomes | 3.13E-15 | 3.13E-15 | 3.13E-15 | 3.13E-15 | 3.13E-15 | 3.13E-15 | 3.13E-15 | 3.13E-15 | 3.13E-15 | 3.13E-15 | 1.00E+00 |

Comparison of trees using the Kuhner & Felsenstein, 1994 method

| | BRS3001 | BRS3002 | BRS3003 | BRS3004 | BRS3005 | BRS3006 | BRS3007 | BRS3008 | BRS3009 | BRS3010 | Metagenomes |
|---|---|---|---|---|---|---|---|---|---|---|---|
| BRS3001 | 1.00E+00 | 5.35E-01 | 3.95E-05 | 1.56E-08 | 1.44E-10 | 5.73E-04 | 3.79E-03 | 1.30E-01 | 3.81E-04 | 2.58E-01 | 4.70E-15 |
| BRS3002 | 5.35E-01 | 1.00E+00 | 3.79E-03 | 1.63E-06 | 1.63E-06 | 3.79E-03 | 2.01E-02 | 2.63E-02 | 4.63E-02 | 4.63E-02 | 4.70E-15 |
| BRS3003 | 3.95E-05 | 3.79E-03 | 1.00E+00 | 6.05E-02 | 1.02E-01 | 4.56E-01 | 5.96E-01 | 8.46E-09 | 5.96E-01 | 1.63E-06 | 4.70E-15 |
| BRS3004 | 1.56E-08 | 1.63E-06 | 6.05E-02 | 1.00E+00 | 4.56E-01 | 7.53E-03 | 1.05E-02 | 1.41E-12 | 2.82E-03 | 2.43E-09 | 4.70E-15 |
| BRS3005 | 1.44E-10 | 1.63E-06 | 1.02E-01 | 4.56E-01 | 1.00E+00 | 7.53E-03 | 2.82E-03 | 9.74E-15 | 2.63E-02 | 6.70E-13 | 4.70E-15 |
| BRS3006 | 5.73E-04 | 3.79E-03 | 4.56E-01 | 7.53E-03 | 7.53E-03 | 1.00E+00 | 7.50E-01 | 8.46E-09 | 5.96E-01 | 1.05E-06 | 4.70E-15 |
| BRS3007 | 3.79E-03 | 2.01E-02 | 5.96E-01 | 1.05E-02 | 2.82E-03 | 7.50E-01 | 1.00E+00 | 1.92E-07 | 6.67E-01 | 6.02E-05 | 4.70E-15 |
| BRS3008 | 1.30E-01 | 2.63E-02 | 8.46E-09 | 1.41E-12 | 9.74E-15 | 8.46E-09 | 1.92E-07 | 1.00E+00 | 1.05E-06 | 6.67E-01 | 4.70E-15 |
| BRS3009 | 3.81E-04 | 4.63E-02 | 5.96E-01 | 2.82E-03 | 2.63E-02 | 5.96E-01 | 6.67E-01 | 1.05E-06 | 1.00E+00 | 6.02E-05 | 4.70E-15 |
| BRS3010 | 2.58E-01 | 4.63E-02 | 1.63E-06 | 2.43E-09 | 6.70E-13 | 1.05E-06 | 6.02E-05 | 6.67E-01 | 6.02E-05 | 1.00E+00 | 4.70E-15 |
| Metagenomes | 4.70E-15 | 4.70E-15 | 4.70E-15 | 4.70E-15 | 4.70E-15 | 4.70E-15 | 4.70E-15 | 4.70E-15 | 4.70E-15 | 4.70E-15 | 1.00E+00 |

**Supplementary Table S6A:** Pairwise FDR corrected p-values of the Kolmogorow-Smirnow test on the distances between phylogenetic trees for 39 universal proteins in 10 samples of archaeal RefSeq genomes vs one sample containing archaeal MAGs (associated with main Fig. 4a)

| | ARS3001 | ARS3002 | ARS3003 | ARS3004 | ARS3005 | ARS3006 | ARS3007 | ARS3008 | ARS3009 | ARS3010 | Metagenomes |
|---|---|---|---|---|---|---|---|---|---|---|---|
| ARS3001 | 1.00E+00 | 1.93E-03 | 3.85E-03 | 3.60E-02 | 3.74E-05 | 1.08E-04 | 3.85E-03 | 3.74E-05 | 1.78E-01 | 3.85E-03 | 9.42E-18 |
| ARS3002 | 1.93E-03 | 1.00E+00 | 1.78E-01 | 1.15E-01 | 1.78E-01 | 9.00E-01 | 9.00E-01 | 1.78E-01 | 1.31E-05 | 5.77E-01 | 9.42E-18 |
| ARS3003 | 3.85E-03 | 1.78E-01 | 1.00E+00 | 5.77E-01 | 8.66E-03 | 4.22E-01 | 2.85E-01 | 3.60E-02 | 2.83E-04 | 9.00E-01 | 9.42E-18 |
| ARS3004 | 3.60E-02 | 1.15E-01 | 5.77E-01 | 1.00E+00 | 1.85E-02 | 1.15E-01 | 1.15E-01 | 3.85E-03 | 3.85E-03 | 4.22E-01 | 9.42E-18 |
| ARS3005 | 3.74E-05 | 1.78E-01 | 8.66E-03 | 1.85E-02 | 1.00E+00 | 1.78E-01 | 2.85E-01 | 9.81E-01 | 2.01E-08 | 6.91E-02 | 9.42E-18 |
| ARS3006 | 1.08E-04 | 9.00E-01 | 4.22E-01 | 1.15E-01 | 1.78E-01 | 1.00E+00 | 9.00E-01 | 4.22E-01 | 1.22E-06 | 5.77E-01 | 9.42E-18 |
| ARS3007 | 3.85E-03 | 9.00E-01 | 2.85E-01 | 1.15E-01 | 2.85E-01 | 9.00E-01 | 1.00E+00 | 5.77E-01 | 1.31E-05 | 7.79E-01 | 9.42E-18 |
| ARS3008 | 3.74E-05 | 1.78E-01 | 3.60E-02 | 3.85E-03 | 9.81E-01 | 4.22E-01 | 5.77E-01 | 1.00E+00 | 4.87E-09 | 1.15E-01 | 9.42E-18 |
| ARS3009 | 1.78E-01 | 1.31E-05 | 2.83E-04 | 3.85E-03 | 2.01E-08 | 1.22E-06 | 1.31E-05 | 4.87E-09 | 1.00E+00 | 2.83E-04 | 9.42E-18 |
| ARS3010 | 3.85E-03 | 5.77E-01 | 9.00E-01 | 4.22E-01 | 6.91E-02 | 5.77E-01 | 7.79E-01 | 1.15E-01 | 2.83E-04 | 1.00E+00 | 9.42E-18 |
| Metagenomes | 9.42E-18 | 9.42E-18 | 9.42E-18 | 9.42E-18 | 9.42E-18 | 9.42E-18 | 9.42E-18 | 9.42E-18 | 9.42E-18 | 9.42E-18 | 1.00E+00 |

**Supplementary Table S6B:** Pairwise FDR corrected p-values of the Kolmogorow-Smirnow test on the distances between phylogenetic trees for 23 ribosomal proteins (including SecY) in 10 samples of archaeal RefSeq genomes vs one sample containing archaeal MAGs (associated with main Fig. 4b)

| | ARS3001 | ARS3002 | ARS3003 | ARS3004 | ARS3005 | ARS3006 | ARS3007 | ARS3008 | ARS3009 | ARS3010 | Metagenomes |
|---|---|---|---|---|---|---|---|---|---|---|---|
| ARS3001 | 1.00E+00 | 1.47E-02 | 1.65E-01 | 1.65E-01 | 6.25E-04 | 1.47E-02 | 2.00E-03 | 6.25E-04 | 1.65E-01 | 3.82E-02 | 1.36E-10 |
| ARS3002 | 1.47E-02 | 1.00E+00 | 8.40E-02 | 4.83E-01 | 9.83E-01 | 8.91E-01 | 8.91E-01 | 6.66E-01 | 6.25E-04 | 6.66E-01 | 1.36E-10 |
| ARS3003 | 1.65E-01 | 8.40E-02 | 1.00E+00 | 6.66E-01 | 1.65E-01 | 4.83E-01 | 8.40E-02 | 1.47E-02 | 5.63E-03 | 4.83E-01 | 1.36E-10 |
| ARS3004 | 1.65E-01 | 4.83E-01 | 6.66E-01 | 1.00E+00 | 2.98E-01 | 6.66E-01 | 2.98E-01 | 8.40E-02 | 5.63E-03 | 6.66E-01 | 1.36E-10 |
| ARS3005 | 6.25E-04 | 9.83E-01 | 1.65E-01 | 2.98E-01 | 1.00E+00 | 6.66E-01 | 9.83E-01 | 4.83E-01 | 6.25E-04 | 8.91E-01 | 1.36E-10 |
| ARS3006 | 1.47E-02 | 8.91E-01 | 4.83E-01 | 6.66E-01 | 6.66E-01 | 1.00E+00 | 6.66E-01 | 2.98E-01 | 5.63E-03 | 9.83E-01 | 1.36E-10 |
| ARS3007 | 2.00E-03 | 8.91E-01 | 8.40E-02 | 2.98E-01 | 9.83E-01 | 6.66E-01 | 1.00E+00 | 6.66E-01 | 2.10E-04 | 4.83E-01 | 1.36E-10 |
| ARS3008 | 6.25E-04 | 6.66E-01 | 1.47E-02 | 8.40E-02 | 4.83E-01 | 2.98E-01 | 6.66E-01 | 1.00E+00 | 1.10E-05 | 2.98E-01 | 1.36E-10 |
| ARS3009 | 1.65E-01 | 6.25E-04 | 5.63E-03 | 5.63E-03 | 6.25E-04 | 5.63E-03 | 2.10E-04 | 1.10E-05 | 1.00E+00 | 2.00E-03 | 1.36E-10 |
| ARS3010 | 3.82E-02 | 6.66E-01 | 4.83E-01 | 6.66E-01 | 8.91E-01 | 9.83E-01 | 4.83E-01 | 2.98E-01 | 2.00E-03 | 1.00E+00 | 1.36E-10 |
| Metagenomes | 1.36E-10 | 1.36E-10 | 1.36E-10 | 1.36E-10 | 1.36E-10 | 1.36E-10 | 1.36E-10 | 1.36E-10 | 1.36E-10 | 1.36E-10 | 1.00E+00 |

**Supplementary Table S6C:** Pairwise FDR corrected p-values of the Kolmogorow-Smirnow test on the distances between phylogenetic trees for 16 non-ribosomal proteins in 10 samples of archaeal RefSeq genomes vs one sample containing archaeal MAGs (associated with main Fig. 4c)

| | ARS3001 | ARS3002 | ARS3003 | ARS3004 | ARS3005 | ARS3006 | ARS3007 | ARS3008 | ARS3009 | ARS3010 | Metagenomes |
|---|---|---|---|---|---|---|---|---|---|---|---|
| ARS3001 | 1.00E+00 | 2.88E-01 | 2.88E-01 | 9.12E-01 | 2.16E-02 | 1.44E-01 | 4.79E-01 | 6.04E-02 | 4.79E-01 | 4.79E-01 | 1.95E-07 |
| ARS3002 | 2.88E-01 | 1.00E+00 | 7.25E-01 | 4.79E-01 | 4.79E-01 | 9.12E-01 | 9.12E-01 | 7.25E-01 | 6.88E-03 | 9.12E-01 | 1.95E-07 |
| ARS3003 | 2.88E-01 | 7.25E-01 | 1.00E+00 | 2.88E-01 | 2.88E-01 | 9.12E-01 | 7.25E-01 | 7.25E-01 | 6.88E-03 | 7.25E-01 | 1.95E-07 |
| ARS3004 | 9.12E-01 | 4.79E-01 | 2.88E-01 | 1.00E+00 | 6.04E-02 | 1.44E-01 | 4.79E-01 | 6.04E-02 | 1.44E-01 | 9.12E-01 | 1.95E-07 |
| ARS3005 | 2.16E-02 | 4.79E-01 | 2.88E-01 | 6.04E-02 | 1.00E+00 | 4.79E-01 | 2.88E-01 | 7.25E-01 | 7.64E-05 | 1.44E-01 | 1.95E-07 |
| ARS3006 | 1.44E-01 | 9.12E-01 | 9.12E-01 | 1.44E-01 | 4.79E-01 | 1.00E+00 | 7.25E-01 | 7.25E-01 | 1.84E-03 | 4.79E-01 | 1.95E-07 |
| ARS3007 | 4.79E-01 | 9.12E-01 | 7.25E-01 | 4.79E-01 | 2.88E-01 | 7.25E-01 | 1.00E+00 | 4.79E-01 | 2.16E-02 | 9.12E-01 | 1.95E-07 |
| ARS3008 | 6.04E-02 | 7.25E-01 | 7.25E-01 | 6.04E-02 | 7.25E-01 | 7.25E-01 | 4.79E-01 | 1.00E+00 | 4.00E-04 | 2.88E-01 | 1.95E-07 |
| ARS3009 | 4.79E-01 | 6.88E-03 | 6.88E-03 | 1.44E-01 | 7.64E-05 | 1.84E-03 | 2.16E-02 | 4.00E-04 | 1.00E+00 | 2.16E-02 | 1.95E-07 |
| ARS3010 | 4.79E-01 | 9.12E-01 | 7.25E-01 | 9.12E-01 | 1.44E-01 | 4.79E-01 | 9.12E-01 | 2.88E-01 | 2.16E-02 | 1.00E+00 | 1.95E-07 |
| Metagenomes | 1.95E-07 | 1.95E-07 | 1.95E-07 | 1.95E-07 | 1.95E-07 | 1.95E-07 | 1.95E-07 | 1.95E-07 | 1.95E-07 | 1.95E-07 | 1.00E+00 |

**Supplementary Table S6D:** Pairwise FDR corrected p-values of the Kolmogorow-Smirnow test on the distances between phylogenetic trees for ribosomal proteins in 10 samples of bacterial RefSeq genomes vs one sample containing CPR MAGs (associated with main Fig. 4d)

| | BRS3001 | BRS3002 | BRS3003 | BRS3004 | BRS3005 | BRS3006 | BRS3007 | BRS3008 | BRS3009 | BRS3010 | Meta_CPR3004 |
|---|---|---|---|---|---|---|---|---|---|---|---|
| BRS3001 | 1.00E+00 | 2.48E-01 | 6.69E-01 | 7.37E-03 | 6.69E-01 | 5.29E-02 | 9.12E-01 | 2.48E-01 | 6.69E-01 | 2.04E-02 | 1.95E-07 |
| BRS3002 | 2.48E-01 | 1.00E+00 | 4.56E-01 | 2.17E-03 | 1.29E-01 | 2.04E-02 | 6.69E-01 | 5.29E-02 | 1.29E-01 | 7.37E-03 | 1.95E-07 |
| BRS3003 | 6.69E-01 | 4.56E-01 | 1.00E+00 | 5.29E-02 | 4.56E-01 | 4.56E-01 | 9.12E-01 | 6.69E-01 | 6.69E-01 | 2.48E-01 | 1.95E-07 |
| BRS3004 | 7.37E-03 | 2.17E-03 | 5.29E-02 | 1.00E+00 | 2.04E-02 | 9.12E-01 | 2.04E-02 | 2.48E-01 | 5.29E-02 | 2.48E-01 | 1.95E-07 |
| BRS3005 | 6.69E-01 | 1.29E-01 | 4.56E-01 | 2.04E-02 | 1.00E+00 | 2.04E-02 | 4.56E-01 | 2.48E-01 | 6.69E-01 | 7.37E-03 | 1.95E-07 |
| BRS3006 | 5.29E-02 | 2.04E-02 | 4.56E-01 | 9.12E-01 | 2.04E-02 | 1.00E+00 | 1.29E-01 | 6.69E-01 | 5.29E-02 | 4.56E-01 | 1.95E-07 |
| BRS3007 | 9.12E-01 | 6.69E-01 | 9.12E-01 | 2.04E-02 | 4.56E-01 | 1.29E-01 | 1.00E+00 | 6.69E-01 | 4.56E-01 | 1.29E-01 | 1.95E-07 |
| BRS3008 | 2.48E-01 | 5.29E-02 | 6.69E-01 | 2.48E-01 | 2.48E-01 | 6.69E-01 | 6.69E-01 | 1.00E+00 | 2.48E-01 | 6.69E-01 | 1.95E-07 |
| BRS3009 | 6.69E-01 | 1.29E-01 | 6.69E-01 | 5.29E-02 | 6.69E-01 | 5.29E-02 | 4.56E-01 | 2.48E-01 | 1.00E+00 | 2.48E-01 | 1.95E-07 |
| BRS3010 | 2.04E-02 | 7.37E-03 | 2.48E-01 | 2.48E-01 | 7.37E-03 | 4.56E-01 | 1.29E-01 | 6.69E-01 | 2.48E-01 | 1.00E+00 | 1.95E-07 |
| Meta_CPR3004 | 1.95E-07 | 1.95E-07 | 1.95E-07 | 1.95E-07 | 1.95E-07 | 1.95E-07 | 1.95E-07 | 1.95E-07 | 1.95E-07 | 1.95E-07 | 1.00E+00 |

**Supplementary Table S6E:** Pairwise FDR corrected p-values of the Kolmogorow-Smirnow test on the distances between phylogenetic trees for 20 ribosomal proteins in 10 samples of bacterial RefSeq genomes vs one sample containing bacterial MAGs (associated with main Fig. 4e)

| | BRS3001 | BRS3002 | BRS3003 | BRS3004 | BRS3005 | BRS3006 | BRS3007 | BRS3008 | BRS3009 | BRS3010 | Meta_BMS3002 |
|---|---|---|---|---|---|---|---|---|---|---|---|
| BRS3001 | 1.00E+00 | 5.47E-01 | 3.78E-01 | 3.32E-03 | 5.47E-01 | 9.43E-03 | 5.47E-01 | 7.85E-01 | 2.32E-01 | 1.25E-01 | 3.05E-09 |
| BRS3002 | 5.47E-01 | 1.00E+00 | 3.78E-01 | 9.43E-03 | 2.32E-01 | 2.36E-02 | 3.78E-01 | 9.65E-01 | 2.32E-01 | 1.25E-01 | 3.05E-09 |
| BRS3003 | 3.78E-01 | 3.78E-01 | 1.00E+00 | 2.32E-01 | 3.78E-01 | 3.78E-01 | 5.47E-01 | 5.47E-01 | 5.47E-01 | 5.47E-01 | 3.05E-09 |
| BRS3004 | 3.32E-03 | 9.43E-03 | 2.32E-01 | 1.00E+00 | 3.32E-03 | 7.85E-01 | 5.55E-02 | 2.36E-02 | 5.55E-02 | 5.55E-02 | 3.05E-09 |
| BRS3005 | 5.47E-01 | 2.32E-01 | 3.78E-01 | 3.32E-03 | 1.00E+00 | 9.43E-03 | 5.47E-01 | 3.78E-01 | 5.47E-01 | 5.47E-01 | 3.05E-09 |
| BRS3006 | 9.43E-03 | 2.36E-02 | 3.78E-01 | 7.85E-01 | 9.43E-03 | 1.00E+00 | 1.25E-01 | 5.55E-02 | 2.36E-02 | 2.36E-02 | 3.05E-09 |
| BRS3007 | 5.47E-01 | 3.78E-01 | 5.47E-01 | 5.55E-02 | 5.47E-01 | 1.25E-01 | 1.00E+00 | 7.85E-01 | 3.78E-01 | 2.32E-01 | 3.05E-09 |
| BRS3008 | 7.85E-01 | 9.65E-01 | 5.47E-01 | 2.36E-02 | 3.78E-01 | 5.55E-02 | 7.85E-01 | 1.00E+00 | 3.78E-01 | 2.32E-01 | 3.05E-09 |
| BRS3009 | 2.32E-01 | 2.32E-01 | 5.47E-01 | 5.55E-02 | 5.47E-01 | 2.36E-02 | 3.78E-01 | 3.78E-01 | 1.00E+00 | 7.85E-01 | 3.05E-09 |
| BRS3010 | 1.25E-01 | 1.25E-01 | 5.47E-01 | 5.55E-02 | 5.47E-01 | 2.36E-02 | 2.32E-01 | 2.32E-01 | 7.85E-01 | 1.00E+00 | 3.05E-09 |
| Meta_BMS3002 | 3.05E-09 | 3.05E-09 | 3.05E-09 | 3.05E-09 | 3.05E-09 | 3.05E-09 | 3.05E-09 | 3.05E-09 | 3.05E-09 | 3.05E-09 | 1.00E+00 |

**Supplementary Table S7A:** Pairwise FDR corrected p-values of the Kolmogorow-Smirnow test on the distances between phylogenetic trees for 39 universal proteins in 10 samples of archaeal RefSeq genomes vs 10 samples of archaeal MAGs (associated with Supplementary Fig. S8a)

| | ARS3001 | ARS3002 | ARS3003 | ARS3004 | ARS3005 | ARS3006 | ARS3007 | ARS3008 | ARS3009 | ARS3010 | Meta_AMS3001 | Meta_AMS3002 | Meta_AMS3003 | Meta_AMS3004 | Meta_AMS3005 | Meta_AMS3006 | Meta_AMS3007 | Meta_AMS3008 | Meta_AMS3009 | Meta_AMS3010 |
|---|---|---|---|---|---|---|---|---|---|---|---|---|---|---|---|---|---|---|---|---|
| ARS3001 | 1.00E+00 | 1.13E-03 | 2.68E-03 | 2.67E-02 | 1.98E-05 | 5.91E-05 | 2.68E-03 | 1.98E-05 | 1.56E-01 | 2.68E-03 | 7.23E-18 | 7.23E-18 | 7.23E-18 | 7.23E-18 | 7.23E-18 | 7.23E-18 | 7.23E-18 | 7.23E-18 | 7.23E-18 | 7.23E-18 |
| ARS3002 | 1.13E-03 | 1.00E+00 | 1.56E-01 | 9.19E-02 | 1.56E-01 | 8.89E-01 | 8.89E-01 | 1.56E-01 | 6.21E-06 | 5.46E-01 | 7.23E-18 | 4.79E-13 | 2.72E-12 | 1.29E-14 | 7.23E-18 | 4.79E-13 | 7.23E-18 | 1.29E-14 | 8.10E-14 | 7.23E-18 |
| ARS3003 | 2.68E-03 | 1.56E-01 | 1.00E+00 | 5.46E-01 | 6.12E-03 | 3.82E-01 | 2.56E-01 | 2.67E-02 | 1.66E-04 | 8.89E-01 | 7.23E-18 | 2.16E-15 | 4.79E-13 | 4.86E-17 | 7.23E-18 | 7.23E-18 | 7.23E-18 | 7.23E-18 | 7.23E-18 | 7.23E-18 |
| ARS3004 | 2.67E-02 | 9.19E-02 | 5.46E-01 | 1.00E+00 | 1.31E-02 | 9.19E-02 | 9.19E-02 | 2.68E-03 | 2.68E-03 | 3.82E-01 | 7.23E-18 | 3.20E-16 | 8.10E-14 | 4.86E-17 | 7.23E-18 | 7.23E-18 | 7.23E-18 | 7.23E-18 | 7.23E-18 | 7.23E-18 |
| ARS3005 | 1.98E-05 | 1.56E-01 | 6.12E-03 | 1.31E-02 | 1.00E+00 | 1.56E-01 | 2.56E-01 | 9.81E-01 | 8.16E-09 | 5.14E-02 | 3.20E-16 | 2.72E-12 | 1.85E-09 | 2.16E-15 | 7.23E-18 | 2.72E-12 | 3.20E-16 | 3.20E-16 | 1.29E-14 | 3.20E-16 |
| ARS3006 | 5.91E-05 | 8.89E-01 | 3.82E-01 | 9.19E-02 | 1.56E-01 | 1.00E+00 | 8.89E-01 | 3.82E-01 | 5.21E-07 | 5.46E-01 | 4.86E-17 | 8.10E-14 | 1.51E-11 | 3.20E-16 | 7.23E-18 | 1.29E-14 | 4.86E-17 | 3.20E-16 | 2.16E-15 | 4.86E-17 |
| ARS3007 | 2.68E-03 | 8.89E-01 | 2.56E-01 | 9.19E-02 | 2.56E-01 | 8.89E-01 | 1.00E+00 | 5.46E-01 | 6.21E-06 | 7.40E-01 | 4.86E-17 | 4.79E-13 | 3.97E-10 | 1.29E-14 | 7.23E-18 | 1.29E-14 | 4.86E-17 | 2.16E-15 | 1.29E-14 | 4.86E-17 |
| ARS3008 | 1.98E-05 | 1.56E-01 | 2.67E-02 | 2.68E-03 | 9.81E-01 | 3.82E-01 | 5.46E-01 | 1.00E+00 | 1.85E-09 | 9.19E-02 | 3.20E-16 | 2.72E-12 | 8.16E-09 | 8.10E-14 | 7.23E-18 | 1.29E-14 | 1.29E-14 | 1.29E-14 | 1.29E-14 | 1.29E-14 |
| ARS3009 | 1.56E-01 | 6.21E-06 | 1.66E-04 | 2.68E-03 | 8.16E-09 | 5.21E-07 | 6.21E-06 | 1.85E-09 | 1.00E+00 | 1.66E-04 | 7.23E-18 | 7.23E-18 | 4.86E-17 | 7.23E-18 | 7.23E-18 | 7.23E-18 | 7.23E-18 | 7.23E-18 | 7.23E-18 | 7.23E-18 |
| ARS3010 | 2.68E-03 | 5.46E-01 | 8.89E-01 | 3.82E-01 | 5.14E-02 | 5.46E-01 | 7.40E-01 | 9.19E-02 | 1.66E-04 | 1.00E+00 | 7.23E-18 | 2.16E-15 | 4.79E-13 | 3.20E-16 | 7.23E-18 | 2.16E-15 | 7.23E-18 | 4.86E-17 | 3.20E-16 | 7.23E-18 |
| Meta_AMS3001 | 7.23E-18 | 7.23E-18 | 7.23E-18 | 7.23E-18 | 3.20E-16 | 4.86E-17 | 4.86E-17 | 3.20E-16 | 7.23E-18 | 7.23E-18 | 1.00E+00 | 1.13E-03 | 5.91E-05 | 6.12E-03 | 6.12E-03 | 1.13E-03 | 2.56E-01 | 3.82E-01 | 1.31E-02 | 9.19E-02 |
| Meta_AMS3002 | 7.23E-18 | 4.79E-13 | 2.16E-15 | 3.20E-16 | 2.72E-12 | 8.10E-14 | 4.79E-13 | 2.72E-12 | 7.23E-18 | 2.16E-15 | 1.13E-03 | 1.00E+00 | 3.82E-01 | 3.82E-01 | 1.38E-07 | 5.46E-01 | 2.67E-02 | 1.56E-01 | 7.40E-01 | 5.14E-02 |
| Meta_AMS3003 | 7.23E-18 | 2.72E-12 | 4.79E-13 | 8.10E-14 | 1.85E-09 | 1.51E-11 | 3.97E-10 | 8.16E-09 | 4.86E-17 | 4.79E-13 | 5.91E-05 | 3.82E-01 | 1.00E+00 | 1.31E-02 | 3.44E-08 | 5.14E-02 | 4.49E-04 | 1.31E-02 | 2.56E-01 | 4.49E-04 |
| Meta_AMS3004 | 7.23E-18 | 1.29E-14 | 4.86E-17 | 4.86E-17 | 2.16E-15 | 3.20E-16 | 1.29E-14 | 8.10E-14 | 7.23E-18 | 3.20E-16 | 6.12E-03 | 3.82E-01 | 1.31E-02 | 1.00E+00 | 1.85E-06 | 3.82E-01 | 3.82E-01 | 3.82E-01 | 7.40E-01 | 5.46E-01 |
| Meta_AMS3005 | 7.23E-18 | 7.23E-18 | 7.23E-18 | 7.23E-18 | 7.23E-18 | 7.23E-18 | 7.23E-18 | 7.23E-18 | 7.23E-18 | 7.23E-18 | 6.12E-03 | 1.38E-07 | 3.44E-08 | 1.85E-06 | 1.00E+00 | 1.85E-06 | 1.66E-04 | 1.13E-03 | 1.66E-04 | 5.91E-05 |
| Meta_AMS3006 | 7.23E-18 | 4.79E-13 | 7.23E-18 | 7.23E-18 | 2.72E-12 | 1.29E-14 | 1.29E-14 | 1.29E-14 | 7.23E-18 | 2.16E-15 | 1.13E-03 | 5.46E-01 | 5.14E-02 | 3.82E-01 | 1.85E-06 | 1.00E+00 | 2.67E-02 | 1.56E-01 | 8.89E-01 | 9.19E-02 |
| Meta_AMS3007 | 7.23E-18 | 7.23E-18 | 7.23E-18 | 7.23E-18 | 3.20E-16 | 4.86E-17 | 4.86E-17 | 1.29E-14 | 7.23E-18 | 7.23E-18 | 2.56E-01 | 2.67E-02 | 4.49E-04 | 3.82E-01 | 1.66E-04 | 2.67E-02 | 1.00E+00 | 8.89E-01 | 9.19E-02 | 5.46E-01 |
| Meta_AMS3008 | 7.23E-18 | 1.29E-14 | 7.23E-18 | 7.23E-18 | 3.20E-16 | 3.20E-16 | 2.16E-15 | 1.29E-14 | 7.23E-18 | 4.86E-17 | 3.82E-01 | 1.56E-01 | 1.31E-02 | 3.82E-01 | 1.13E-03 | 1.56E-01 | 8.89E-01 | 1.00E+00 | 3.82E-01 | 8.89E-01 |
| Meta_AMS3009 | 7.23E-18 | 8.10E-14 | 7.23E-18 | 7.23E-18 | 1.29E-14 | 2.16E-15 | 1.29E-14 | 1.29E-14 | 7.23E-18 | 3.20E-16 | 1.31E-02 | 7.40E-01 | 2.56E-01 | 7.40E-01 | 1.66E-04 | 8.89E-01 | 9.19E-02 | 3.82E-01 | 1.00E+00 | 5.14E-02 |
| Meta_AMS3010 | 7.23E-18 | 7.23E-18 | 7.23E-18 | 7.23E-18 | 3.20E-16 | 4.86E-17 | 4.86E-17 | 1.29E-14 | 7.23E-18 | 7.23E-18 | 9.19E-02 | 5.14E-02 | 4.49E-04 | 5.46E-01 | 5.91E-05 | 9.19E-02 | 5.46E-01 | 8.89E-01 | 5.14E-02 | 1.00E+00 |

**Supplementary Table S7B:** Pairwise FDR corrected p-values of the Kolmogorow-Smirnow test on the distances between phylogenetic trees for 23 ribosomal proteins (including SecY) in 10 samples of archaeal RefSeq genomes vs 10 samples of archaeal MAGs (associated with Supplementary Fig. S8b)

| | ARS3001 | ARS3002 | ARS3003 | ARS3004 | ARS3005 | ARS3006 | ARS3007 | ARS3008 | ARS3009 | ARS3010 | Meta_AMS3001 | Meta_AMS3002 | Meta_AMS3003 | Meta_AMS3004 | Meta_AMS3005 | Meta_AMS3006 | Meta_AMS3007 | Meta_AMS3008 | Meta_AMS3009 | Meta_AMS3010 |
|---|---|---|---|---|---|---|---|---|---|---|---|---|---|---|---|---|---|---|---|---|
| ARS3001 | 1.00E+00 | 1.01E-02 | 1.21E-01 | 1.21E-01 | 3.76E-04 | 1.01E-02 | 1.23E-03 | 3.76E-04 | 1.21E-01 | 2.55E-02 | 3.14E-10 | 3.76E-04 | 1.01E-02 | 1.90E-09 | 3.14E-10 | 2.82E-05 | 3.14E-10 | 2.74E-07 | 1.39E-06 | 3.14E-10 |
| ARS3002 | 1.01E-02 | 1.00E+00 | 5.76E-02 | 4.05E-01 | 9.83E-01 | 8.56E-01 | 8.56E-01 | 6.19E-01 | 3.76E-04 | 6.19E-01 | 5.51E-08 | 1.21E-01 | 4.05E-01 | 6.53E-06 | 3.14E-10 | 3.61E-03 | 5.51E-08 | 1.10E-04 | 3.61E-03 | 2.74E-07 |
| ARS3003 | 1.21E-01 | 5.76E-02 | 1.00E+00 | 6.19E-01 | 1.21E-01 | 4.05E-01 | 5.76E-02 | 1.01E-02 | 3.61E-03 | 4.05E-01 | 1.90E-09 | 1.23E-03 | 2.55E-02 | 9.90E-09 | 3.14E-10 | 2.82E-05 | 1.90E-09 | 2.74E-07 | 1.39E-06 | 9.90E-09 |
| ARS3004 | 1.21E-01 | 4.05E-01 | 6.19E-01 | 1.00E+00 | 2.35E-01 | 6.19E-01 | 2.35E-01 | 5.76E-02 | 3.61E-03 | 6.19E-01 | 9.90E-09 | 1.01E-02 | 1.21E-01 | 6.53E-06 | 3.14E-10 | 3.61E-03 | 9.90E-09 | 1.10E-04 | 3.76E-04 | 2.74E-07 |
| ARS3005 | 3.76E-04 | 9.83E-01 | 1.21E-01 | 2.35E-01 | 1.00E+00 | 6.19E-01 | 9.83E-01 | 4.05E-01 | 3.76E-04 | 8.56E-01 | 5.51E-08 | 1.21E-01 | 6.19E-01 | 2.82E-05 | 3.14E-10 | 1.01E-02 | 2.74E-07 | 3.76E-04 | 3.61E-03 | 2.74E-07 |
| ARS3006 | 1.01E-02 | 8.56E-01 | 4.05E-01 | 6.19E-01 | 6.19E-01 | 1.00E+00 | 6.19E-01 | 2.35E-01 | 3.61E-03 | 9.83E-01 | 5.51E-08 | 5.76E-02 | 4.05E-01 | 2.82E-05 | 1.90E-09 | 3.61E-03 | 2.74E-07 | 3.76E-04 | 3.76E-04 | 2.74E-07 |
| ARS3007 | 1.23E-03 | 8.56E-01 | 5.76E-02 | 2.35E-01 | 9.83E-01 | 6.19E-01 | 1.00E+00 | 6.19E-01 | 1.10E-04 | 4.05E-01 | 5.51E-08 | 4.05E-01 | 8.56E-01 | 1.10E-04 | 3.14E-10 | 2.55E-02 | 5.51E-08 | 3.76E-04 | 1.01E-02 | 2.82E-05 |
| ARS3008 | 3.76E-04 | 6.19E-01 | 1.01E-02 | 5.76E-02 | 4.05E-01 | 2.35E-01 | 6.19E-01 | 1.00E+00 | 6.53E-06 | 2.35E-01 | 2.74E-07 | 6.19E-01 | 6.19E-01 | 3.76E-04 | 9.90E-09 | 1.21E-01 | 6.53E-06 | 3.61E-03 | 5.76E-02 | 6.53E-06 |
| ARS3009 | 1.21E-01 | 3.76E-04 | 3.61E-03 | 3.61E-03 | 3.76E-04 | 3.61E-03 | 1.10E-04 | 6.53E-06 | 1.00E+00 | 1.23E-03 | 3.14E-10 | 2.74E-07 | 1.10E-04 | 3.14E-10 | 3.14E-10 | 9.90E-09 | 3.14E-10 | 9.90E-09 | 9.90E-09 | 3.14E-10 |
| ARS3010 | 2.55E-02 | 6.19E-01 | 4.05E-01 | 6.19E-01 | 8.56E-01 | 9.83E-01 | 4.05E-01 | 2.35E-01 | 1.23E-03 | 1.00E+00 | 9.90E-09 | 2.55E-02 | 4.05E-01 | 6.53E-06 | 3.14E-10 | 3.61E-03 | 9.90E-09 | 2.82E-05 | 1.23E-03 | 5.51E-08 |
| Meta_AMS3001 | 3.14E-10 | 5.51E-08 | 1.90E-09 | 9.90E-09 | 5.51E-08 | 5.51E-08 | 5.51E-08 | 2.74E-07 | 3.14E-10 | 9.90E-09 | 1.00E+00 | 1.39E-06 | 2.74E-07 | 3.76E-04 | 5.76E-02 | 1.10E-04 | 6.19E-01 | 2.55E-02 | 1.23E-03 | 3.61E-03 |
| Meta_AMS3002 | 3.76E-04 | 1.21E-01 | 1.23E-03 | 1.01E-02 | 1.21E-01 | 5.76E-02 | 4.05E-01 | 6.19E-01 | 2.74E-07 | 2.55E-02 | 1.39E-06 | 1.00E+00 | 6.19E-01 | 1.23E-03 | 1.90E-09 | 5.76E-02 | 1.39E-06 | 1.23E-03 | 5.76E-02 | 1.10E-04 |
| Meta_AMS3003 | 1.01E-02 | 4.05E-01 | 2.55E-02 | 1.21E-01 | 6.19E-01 | 4.05E-01 | 8.56E-01 | 6.19E-01 | 1.10E-04 | 4.05E-01 | 2.74E-07 | 6.19E-01 | 1.00E+00 | 1.10E-04 | 1.90E-09 | 2.55E-02 | 1.39E-06 | 1.23E-03 | 5.76E-02 | 1.39E-06 |
| Meta_AMS3004 | 1.90E-09 | 6.53E-06 | 9.90E-09 | 6.53E-06 | 2.82E-05 | 2.82E-05 | 1.10E-04 | 3.76E-04 | 3.14E-10 | 6.53E-06 | 3.76E-04 | 1.23E-03 | 1.10E-04 | 1.00E+00 | 1.39E-06 | 2.35E-01 | 2.55E-02 | 4.05E-01 | 2.35E-01 | 8.56E-01 |
| Meta_AMS3005 | 3.14E-10 | 3.14E-10 | 3.14E-10 | 3.14E-10 | 3.14E-10 | 1.90E-09 | 3.14E-10 | 9.90E-09 | 3.14E-10 | 3.14E-10 | 5.76E-02 | 1.90E-09 | 1.90E-09 | 1.39E-06 | 1.00E+00 | 5.51E-08 | 2.55E-02 | 3.76E-04 | 2.82E-05 | 6.53E-06 |
| Meta_AMS3006 | 2.82E-05 | 3.61E-03 | 2.82E-05 | 3.61E-03 | 1.01E-02 | 3.61E-03 | 2.55E-02 | 1.21E-01 | 9.90E-09 | 3.61E-03 | 1.10E-04 | 5.76E-02 | 2.55E-02 | 2.35E-01 | 5.51E-08 | 1.00E+00 | 1.23E-03 | 5.76E-02 | 2.35E-01 | 5.76E-02 |
| Meta_AMS3007 | 3.14E-10 | 5.51E-08 | 1.90E-09 | 9.90E-09 | 2.74E-07 | 2.74E-07 | 5.51E-08 | 6.53E-06 | 3.14E-10 | 9.90E-09 | 6.19E-01 | 1.39E-06 | 1.39E-06 | 2.55E-02 | 2.55E-02 | 1.23E-03 | 1.00E+00 | 2.35E-01 | 3.61E-03 | 2.55E-02 |
| Meta_AMS3008 | 2.74E-07 | 1.10E-04 | 2.74E-07 | 1.10E-04 | 3.76E-04 | 3.76E-04 | 3.76E-04 | 3.61E-03 | 9.90E-09 | 2.82E-05 | 2.55E-02 | 1.23E-03 | 1.23E-03 | 4.05E-01 | 3.76E-04 | 5.76E-02 | 2.35E-01 | 1.00E+00 | 1.21E-01 | 4.05E-01 |
| Meta_AMS3009 | 1.39E-06 | 3.61E-03 | 1.39E-06 | 3.76E-04 | 3.61E-03 | 3.76E-04 | 1.01E-02 | 5.76E-02 | 9.90E-09 | 1.23E-03 | 1.23E-03 | 5.76E-02 | 5.76E-02 | 2.35E-01 | 2.82E-05 | 2.35E-01 | 3.61E-03 | 1.21E-01 | 1.00E+00 | 5.76E-02 |
| Meta_AMS3010 | 3.14E-10 | 2.74E-07 | 9.90E-09 | 2.74E-07 | 2.74E-07 | 2.74E-07 | 2.82E-05 | 6.53E-06 | 3.14E-10 | 5.51E-08 | 3.61E-03 | 1.10E-04 | 1.39E-06 | 8.56E-01 | 6.53E-06 | 5.76E-02 | 2.55E-02 | 4.05E-01 | 5.76E-02 | 1.00E+00 |

**Supplementary Table S7C:** Pairwise FDR corrected p-values of the Kolmogorow-Smirnow test on the distances between phylogenetic trees for 16 non-ribosomal proteins in 10 samples of archaeal RefSeq genomes vs 10 samples of archaeal MAGs (associated with Supplementary Fig. S8c)

| | ARS3001 | ARS3002 | ARS3003 | ARS3004 | ARS3005 | ARS3006 | ARS3007 | ARS3008 | ARS3009 | ARS3010 | Meta_AMS3001 | Meta_AMS3002 | Meta_AMS3003 | Meta_AMS3004 | Meta_AMS3005 | Meta_AMS3006 | Meta_AMS3007 | Meta_AMS3008 | Meta_AMS3009 | Meta_AMS3010 |
|---|---|---|---|---|---|---|---|---|---|---|---|---|---|---|---|---|---|---|---|---|
| ARS3001 | 1.00E+00 | 2.25E-01 | 2.25E-01 | 9.12E-01 | 1.23E-02 | 1.01E-01 | 4.29E-01 | 3.85E-02 | 4.29E-01 | 4.29E-01 | 6.74E-08 | 6.74E-08 | 6.74E-08 | 6.74E-08 | 6.74E-08 | 6.74E-08 | 6.74E-08 | 6.74E-08 | 6.74E-08 | 6.74E-08 |
| ARS3002 | 2.25E-01 | 1.00E+00 | 6.95E-01 | 4.29E-01 | 4.29E-01 | 9.12E-01 | 9.12E-01 | 6.95E-01 | 3.39E-03 | 9.12E-01 | 6.74E-08 | 6.74E-08 | 6.74E-08 | 6.74E-08 | 6.74E-08 | 6.74E-08 | 6.74E-08 | 6.74E-08 | 6.74E-08 | 6.74E-08 |
| ARS3003 | 2.25E-01 | 6.95E-01 | 1.00E+00 | 2.25E-01 | 2.25E-01 | 9.12E-01 | 6.95E-01 | 6.95E-01 | 3.39E-03 | 6.95E-01 | 6.74E-08 | 6.74E-08 | 6.74E-08 | 6.74E-08 | 6.74E-08 | 6.74E-08 | 6.74E-08 | 6.74E-08 | 6.74E-08 | 6.74E-08 |
| ARS3004 | 9.12E-01 | 4.29E-01 | 2.25E-01 | 1.00E+00 | 3.85E-02 | 1.01E-01 | 4.29E-01 | 3.85E-02 | 1.01E-01 | 9.12E-01 | 6.74E-08 | 6.74E-08 | 6.74E-08 | 6.74E-08 | 6.74E-08 | 6.74E-08 | 6.74E-08 | 6.74E-08 | 6.74E-08 | 6.74E-08 |
| ARS3005 | 1.23E-02 | 4.29E-01 | 2.25E-01 | 3.85E-02 | 1.00E+00 | 4.29E-01 | 2.25E-01 | 6.95E-01 | 2.87E-05 | 1.01E-01 | 6.74E-08 | 6.74E-08 | 6.74E-08 | 6.74E-08 | 6.74E-08 | 6.74E-08 | 6.74E-08 | 6.74E-08 | 6.74E-08 | 6.74E-08 |
| ARS3006 | 1.01E-01 | 9.12E-01 | 9.12E-01 | 1.01E-01 | 4.29E-01 | 1.00E+00 | 6.95E-01 | 6.95E-01 | 8.00E-04 | 4.29E-01 | 6.74E-08 | 6.74E-08 | 6.74E-08 | 6.74E-08 | 6.74E-08 | 6.74E-08 | 6.74E-08 | 6.74E-08 | 6.74E-08 | 6.74E-08 |
| ARS3007 | 4.29E-01 | 9.12E-01 | 6.95E-01 | 4.29E-01 | 2.25E-01 | 6.95E-01 | 1.00E+00 | 4.29E-01 | 1.23E-02 | 9.12E-01 | 6.74E-08 | 6.74E-08 | 6.74E-08 | 6.74E-08 | 6.74E-08 | 6.74E-08 | 6.74E-08 | 6.74E-08 | 6.74E-08 | 6.74E-08 |
| ARS3008 | 3.85E-02 | 6.95E-01 | 6.95E-01 | 3.85E-02 | 6.95E-01 | 6.95E-01 | 4.29E-01 | 1.00E+00 | 1.63E-04 | 2.25E-01 | 6.74E-08 | 6.74E-08 | 6.74E-08 | 6.74E-08 | 6.74E-08 | 6.74E-08 | 6.74E-08 | 6.74E-08 | 6.74E-08 | 6.74E-08 |
| ARS3009 | 4.29E-01 | 3.39E-03 | 3.39E-03 | 1.01E-01 | 2.87E-05 | 8.00E-04 | 1.23E-02 | 1.63E-04 | 1.00E+00 | 1.23E-02 | 6.74E-08 | 6.74E-08 | 6.74E-08 | 6.74E-08 | 6.74E-08 | 6.74E-08 | 6.74E-08 | 6.74E-08 | 6.74E-08 | 6.74E-08 |
| ARS3010 | 4.29E-01 | 9.12E-01 | 6.95E-01 | 9.12E-01 | 1.01E-01 | 4.29E-01 | 9.12E-01 | 2.25E-01 | 1.23E-02 | 1.00E+00 | 6.74E-08 | 6.74E-08 | 6.74E-08 | 6.74E-08 | 6.74E-08 | 6.74E-08 | 6.74E-08 | 6.74E-08 | 6.74E-08 | 6.74E-08 |
| Meta_AMS3001 | 6.74E-08 | 6.74E-08 | 6.74E-08 | 6.74E-08 | 6.74E-08 | 6.74E-08 | 6.74E-08 | 6.74E-08 | 6.74E-08 | 6.74E-08 | 1.00E+00 | 6.95E-01 | 4.29E-01 | 9.12E-01 | 1.01E-01 | 2.25E-01 | 9.12E-01 | 9.12E-01 | 9.12E-01 | 9.12E-01 |
| Meta_AMS3002 | 6.74E-08 | 6.74E-08 | 6.74E-08 | 6.74E-08 | 6.74E-08 | 6.74E-08 | 6.74E-08 | 6.74E-08 | 6.74E-08 | 6.74E-08 | 6.95E-01 | 1.00E+00 | 2.25E-01 | 2.25E-01 | 4.29E-01 | 3.85E-02 | 6.95E-01 | 2.25E-01 | 4.29E-01 | 6.95E-01 |
| Meta_AMS3003 | 6.74E-08 | 6.74E-08 | 6.74E-08 | 6.74E-08 | 6.74E-08 | 6.74E-08 | 6.74E-08 | 6.74E-08 | 6.74E-08 | 6.74E-08 | 4.29E-01 | 2.25E-01 | 1.00E+00 | 9.12E-01 | 1.01E-01 | 4.29E-01 | 4.29E-01 | 6.95E-01 | 6.95E-01 | 2.25E-01 |
| Meta_AMS3004 | 6.74E-08 | 6.74E-08 | 6.74E-08 | 6.74E-08 | 6.74E-08 | 6.74E-08 | 6.74E-08 | 6.74E-08 | 6.74E-08 | 6.74E-08 | 9.12E-01 | 2.25E-01 | 9.12E-01 | 1.00E+00 | 1.01E-01 | 2.25E-01 | 6.95E-01 | 9.12E-01 | 6.95E-01 | 9.12E-01 |
| Meta_AMS3005 | 6.74E-08 | 6.74E-08 | 6.74E-08 | 6.74E-08 | 6.74E-08 | 6.74E-08 | 6.74E-08 | 6.74E-08 | 6.74E-08 | 6.74E-08 | 1.01E-01 | 4.29E-01 | 1.01E-01 | 1.01E-01 | 1.00E+00 | 1.23E-02 | 1.01E-01 | 1.01E-01 | 3.85E-02 | 2.25E-01 |
| Meta_AMS3006 | 6.74E-08 | 6.74E-08 | 6.74E-08 | 6.74E-08 | 6.74E-08 | 6.74E-08 | 6.74E-08 | 6.74E-08 | 6.74E-08 | 6.74E-08 | 2.25E-01 | 3.85E-02 | 4.29E-01 | 2.25E-01 | 1.23E-02 | 1.00E+00 | 6.95E-01 | 4.29E-01 | 6.95E-01 | 4.29E-01 |
| Meta_AMS3007 | 6.74E-08 | 6.74E-08 | 6.74E-08 | 6.74E-08 | 6.74E-08 | 6.74E-08 | 6.74E-08 | 6.74E-08 | 6.74E-08 | 6.74E-08 | 9.12E-01 | 6.95E-01 | 4.29E-01 | 6.95E-01 | 1.01E-01 | 6.95E-01 | 1.00E+00 | 9.12E-01 | 6.95E-01 | 4.29E-01 |
| Meta_AMS3008 | 6.74E-08 | 6.74E-08 | 6.74E-08 | 6.74E-08 | 6.74E-08 | 6.74E-08 | 6.74E-08 | 6.74E-08 | 6.74E-08 | 6.74E-08 | 9.12E-01 | 2.25E-01 | 6.95E-01 | 9.12E-01 | 1.01E-01 | 4.29E-01 | 9.12E-01 | 1.00E+00 | 9.12E-01 | 2.25E-01 |
| Meta_AMS3009 | 6.74E-08 | 6.74E-08 | 6.74E-08 | 6.74E-08 | 6.74E-08 | 6.74E-08 | 6.74E-08 | 6.74E-08 | 6.74E-08 | 6.74E-08 | 9.12E-01 | 4.29E-01 | 6.95E-01 | 6.95E-01 | 3.85E-02 | 6.95E-01 | 6.95E-01 | 9.12E-01 | 1.00E+00 | 6.95E-01 |
| Meta_AMS3010 | 6.74E-08 | 6.74E-08 | 6.74E-08 | 6.74E-08 | 6.74E-08 | 6.74E-08 | 6.74E-08 | 6.74E-08 | 6.74E-08 | 6.74E-08 | 9.12E-01 | 6.95E-01 | 2.25E-01 | 9.12E-01 | 2.25E-01 | 4.29E-01 | 4.29E-01 | 2.25E-01 | 6.95E-01 | 1.00E+00 |

**Supplementary Table S7D:** Pairwise FDR corrected p-values of the Kolmogorow-Smirnow test on the distances between phylogenetic trees for ribosomal proteins in 10 samples of bacterial RefSeq genomes vs 10 samples of CPR MAGs (associated with Supplementary Fig. S8d)

| | BRS3001 | BRS3002 | BRS3003 | BRS3004 | BRS3005 | BRS3006 | BRS3007 | BRS3008 | BRS3009 | BRS3010 | Meta_CPR3001 | Meta_CPR3002 | Meta_CPR3003 | Meta_CPR3004 | Meta_CPR3005 | Meta_CPR3006 | Meta_CPR3007 | Meta_CPR3008 | Meta_CPR3009 | Meta_CPR3010 |
|---|---|---|---|---|---|---|---|---|---|---|---|---|---|---|---|---|---|---|---|---|
| BRS3001 | 1.00E+00 | 1.99E-01 | 6.68E-01 | 3.18E-03 | 6.68E-01 | 3.32E-02 | 9.12E-01 | 1.99E-01 | 6.68E-01 | 1.13E-02 | 1.04E-07 | 1.04E-07 | 1.04E-07 | 1.04E-07 | 1.04E-07 | 1.04E-07 | 1.04E-07 | 1.04E-07 | 1.04E-07 | 1.04E-07 |
| BRS3002 | 1.99E-01 | 1.00E+00 | 4.01E-01 | 7.85E-04 | 8.79E-02 | 1.13E-02 | 6.68E-01 | 3.32E-02 | 8.79E-02 | 3.18E-03 | 1.04E-07 | 1.04E-07 | 6.57E-07 | 1.04E-07 | 1.04E-07 | 1.04E-07 | 1.04E-07 | 6.57E-07 | 6.57E-07 | 1.04E-07 |
| BRS3003 | 6.68E-01 | 4.01E-01 | 1.00E+00 | 3.32E-02 | 4.01E-01 | 4.01E-01 | 9.12E-01 | 6.68E-01 | 6.68E-01 | 1.99E-01 | 1.04E-07 | 1.04E-07 | 6.57E-07 | 1.04E-07 | 1.04E-07 | 1.04E-07 | 1.04E-07 | 4.55E-06 | 4.55E-06 | 6.57E-07 |
| BRS3004 | 3.18E-03 | 7.85E-04 | 3.32E-02 | 1.00E+00 | 1.13E-02 | 9.12E-01 | 1.13E-02 | 1.99E-01 | 3.32E-02 | 1.99E-01 | 1.04E-07 | 1.04E-07 | 6.57E-07 | 1.04E-07 | 1.04E-07 | 1.04E-07 | 1.04E-07 | 4.55E-06 | 1.64E-04 | 6.57E-07 |
| BRS3005 | 6.68E-01 | 8.79E-02 | 4.01E-01 | 1.13E-02 | 1.00E+00 | 1.13E-02 | 4.01E-01 | 1.99E-01 | 6.68E-01 | 3.18E-03 | 1.04E-07 | 1.04E-07 | 1.04E-07 | 1.04E-07 | 1.04E-07 | 1.04E-07 | 1.04E-07 | 1.04E-07 | 1.04E-07 | 1.04E-07 |
| BRS3006 | 3.32E-02 | 1.13E-02 | 4.01E-01 | 9.12E-01 | 1.13E-02 | 1.00E+00 | 8.79E-02 | 6.68E-01 | 3.32E-02 | 4.01E-01 | 1.04E-07 | 6.57E-07 | 4.55E-06 | 1.04E-07 | 6.57E-07 | 6.57E-07 | 1.04E-07 | 4.55E-06 | 2.90E-05 | 6.57E-07 |
| BRS3007 | 9.12E-01 | 6.68E-01 | 9.12E-01 | 1.13E-02 | 4.01E-01 | 8.79E-02 | 1.00E+00 | 6.68E-01 | 4.01E-01 | 8.79E-02 | 1.04E-07 | 1.04E-07 | 6.57E-07 | 1.04E-07 | 6.57E-07 | 1.04E-07 | 1.04E-07 | 6.57E-07 | 2.90E-05 | 6.57E-07 |
| BRS3008 | 1.99E-01 | 3.32E-02 | 6.68E-01 | 1.99E-01 | 1.99E-01 | 6.68E-01 | 6.68E-01 | 1.00E+00 | 1.99E-01 | 6.68E-01 | 1.04E-07 | 6.57E-07 | 6.57E-07 | 1.04E-07 | 6.57E-07 | 6.57E-07 | 1.04E-07 | 6.57E-07 | 4.55E-06 | 6.57E-07 |
| BRS3009 | 6.68E-01 | 8.79E-02 | 6.68E-01 | 3.32E-02 | 6.68E-01 | 3.32E-02 | 4.01E-01 | 1.99E-01 | 1.00E+00 | 1.99E-01 | 1.04E-07 | 1.04E-07 | 1.04E-07 | 1.04E-07 | 1.04E-07 | 1.04E-07 | 1.04E-07 | 1.04E-07 | 6.57E-07 | 1.04E-07 |
| BRS3010 | 1.13E-02 | 3.18E-03 | 1.99E-01 | 1.99E-01 | 3.18E-03 | 4.01E-01 | 8.79E-02 | 6.68E-01 | 1.99E-01 | 1.00E+00 | 1.04E-07 | 1.04E-07 | 6.57E-07 | 1.04E-07 | 1.04E-07 | 1.04E-07 | 1.04E-07 | 4.55E-06 | 4.55E-06 | 6.57E-07 |
| Meta_CPR3001 | 1.04E-07 | 1.04E-07 | 1.04E-07 | 1.04E-07 | 1.04E-07 | 1.04E-07 | 1.04E-07 | 1.04E-07 | 1.04E-07 | 1.04E-07 | 1.00E+00 | 8.79E-02 | 8.79E-02 | 9.12E-01 | 1.99E-01 | 4.01E-01 | 1.99E-01 | 3.32E-02 | 3.18E-03 | 3.32E-02 |
| Meta_CPR3002 | 1.04E-07 | 1.04E-07 | 1.04E-07 | 1.04E-07 | 1.04E-07 | 6.57E-07 | 1.04E-07 | 6.57E-07 | 1.04E-07 | 1.04E-07 | 8.79E-02 | 1.00E+00 | 6.68E-01 | 3.32E-02 | 9.12E-01 | 8.79E-02 | 3.18E-03 | 6.68E-01 | 3.32E-02 | 6.68E-01 |
| Meta_CPR3003 | 1.04E-07 | 6.57E-07 | 6.57E-07 | 6.57E-07 | 1.04E-07 | 4.55E-06 | 6.57E-07 | 6.57E-07 | 1.04E-07 | 6.57E-07 | 8.79E-02 | 6.68E-01 | 1.00E+00 | 3.32E-02 | 6.68E-01 | 8.79E-02 | 7.85E-04 | 9.12E-01 | 4.01E-01 | 9.12E-01 |
| Meta_CPR3004 | 1.04E-07 | 1.04E-07 | 1.04E-07 | 1.04E-07 | 1.04E-07 | 1.04E-07 | 1.04E-07 | 1.04E-07 | 1.04E-07 | 1.04E-07 | 9.12E-01 | 3.32E-02 | 3.32E-02 | 1.00E+00 | 8.79E-02 | 1.99E-01 | 9.12E-01 | 3.18E-03 | 7.85E-04 | 1.13E-02 |
| Meta_CPR3005 | 1.04E-07 | 1.04E-07 | 1.04E-07 | 1.04E-07 | 1.04E-07 | 6.57E-07 | 6.57E-07 | 6.57E-07 | 1.04E-07 | 1.04E-07 | 1.99E-01 | 9.12E-01 | 6.68E-01 | 8.79E-02 | 1.00E+00 | 4.01E-01 | 3.32E-02 | 6.68E-01 | 3.32E-02 | 9.12E-01 |
| Meta_CPR3006 | 1.04E-07 | 1.04E-07 | 1.04E-07 | 1.04E-07 | 1.04E-07 | 6.57E-07 | 1.04E-07 | 6.57E-07 | 1.04E-07 | 1.04E-07 | 4.01E-01 | 8.79E-02 | 8.79E-02 | 1.99E-01 | 4.01E-01 | 1.00E+00 | 1.99E-01 | 3.32E-02 | 1.13E-02 | 1.99E-01 |
| Meta_CPR3007 | 1.04E-07 | 1.04E-07 | 1.04E-07 | 1.04E-07 | 1.04E-07 | 1.04E-07 | 1.04E-07 | 1.04E-07 | 1.04E-07 | 1.04E-07 | 1.99E-01 | 3.18E-03 | 7.85E-04 | 9.12E-01 | 3.32E-02 | 1.99E-01 | 1.00E+00 | 7.85E-04 | 2.90E-05 | 3.18E-03 |
| Meta_CPR3008 | 1.04E-07 | 6.57E-07 | 4.55E-06 | 4.55E-06 | 1.04E-07 | 4.55E-06 | 6.57E-07 | 6.57E-07 | 1.04E-07 | 4.55E-06 | 3.32E-02 | 6.68E-01 | 9.12E-01 | 3.18E-03 | 6.68E-01 | 3.32E-02 | 7.85E-04 | 1.00E+00 | 4.01E-01 | 9.12E-01 |
| Meta_CPR3009 | 1.04E-07 | 6.57E-07 | 4.55E-06 | 1.64E-04 | 1.04E-07 | 2.90E-05 | 2.90E-05 | 4.55E-06 | 6.57E-07 | 4.55E-06 | 3.18E-03 | 3.32E-02 | 4.01E-01 | 7.85E-04 | 3.32E-02 | 1.13E-02 | 2.90E-05 | 4.01E-01 | 1.00E+00 | 8.79E-02 |
| Meta_CPR3010 | 1.04E-07 | 1.04E-07 | 6.57E-07 | 6.57E-07 | 1.04E-07 | 6.57E-07 | 6.57E-07 | 6.57E-07 | 1.04E-07 | 6.57E-07 | 3.32E-02 | 6.68E-01 | 9.12E-01 | 1.13E-02 | 9.12E-01 | 1.99E-01 | 3.18E-03 | 9.12E-01 | 8.79E-02 | 1.00E+00 |

**Supplementary Table S7E:** Pairwise FDR corrected p-values of the Kolmogorow-Smirnow test on the distances between phylogenetic trees for 20 ribosomal proteins in 10 samples of bacterial RefSeq genomes vs 10 samples of bacterial MAGs(associated with Supplementary Fig. S8e)

| | BRS3001 | BRS3002 | BRS3003 | BRS3004 | BRS3005 | BRS3006 | BRS3007 | BRS3008 | BRS3009 | BRS3010 | Meta_BMS3001 | Meta_BMS3002 | Meta_BMS3003 | Meta_BMS3004 | Meta_BMS3005 | Meta_BMS3006 | Meta_BMS3007 | Meta_BMS3008 | Meta_BMS3009 | Meta_BMS3010 |
|---|---|---|---|---|---|---|---|---|---|---|---|---|---|---|---|---|---|---|---|---|
| BRS3001 | 1.00E+00 | 5.19E-01 | 3.11E-01 | 1.09E-03 | 5.19E-01 | 3.64E-03 | 5.19E-01 | 7.83E-01 | 1.62E-01 | 7.49E-02 | 1.85E-09 | 1.85E-09 | 1.85E-09 | 1.85E-09 | 1.85E-09 | 1.85E-09 | 1.85E-09 | 1.15E-08 | 1.85E-09 | 1.85E-09 |
| BRS3002 | 5.19E-01 | 1.00E+00 | 3.11E-01 | 3.64E-03 | 1.62E-01 | 1.12E-02 | 3.11E-01 | 9.65E-01 | 1.62E-01 | 7.49E-02 | 1.85E-09 | 1.85E-09 | 7.18E-08 | 1.85E-09 | 1.85E-09 | 1.85E-09 | 7.18E-08 | 7.18E-08 | 1.85E-09 | 1.15E-08 |
| BRS3003 | 3.11E-01 | 3.11E-01 | 1.00E+00 | 1.62E-01 | 3.11E-01 | 3.11E-01 | 5.19E-01 | 5.19E-01 | 5.19E-01 | 5.19E-01 | 1.85E-09 | 1.85E-09 | 2.64E-06 | 1.15E-08 | 1.85E-09 | 1.85E-09 | 2.64E-06 | 2.64E-06 | 1.15E-08 | 4.59E-07 |
| BRS3004 | 1.09E-03 | 3.64E-03 | 1.62E-01 | 1.00E+00 | 1.09E-03 | 7.83E-01 | 3.06E-02 | 1.12E-02 | 3.06E-02 | 3.06E-02 | 1.85E-09 | 1.85E-09 | 2.64E-06 | 1.15E-08 | 1.85E-09 | 1.85E-09 | 1.39E-05 | 2.64E-06 | 1.15E-08 | 4.59E-07 |
| BRS3005 | 5.19E-01 | 1.62E-01 | 3.11E-01 | 1.09E-03 | 1.00E+00 | 3.64E-03 | 5.19E-01 | 3.11E-01 | 5.19E-01 | 5.19E-01 | 1.85E-09 | 1.85E-09 | 1.15E-08 | 1.85E-09 | 1.85E-09 | 1.85E-09 | 1.15E-08 | 1.15E-08 | 1.15E-08 | 1.15E-08 |
| BRS3006 | 3.64E-03 | 1.12E-02 | 3.11E-01 | 7.83E-01 | 3.64E-03 | 1.00E+00 | 7.49E-02 | 3.06E-02 | 1.12E-02 | 1.12E-02 | 1.85E-09 | 1.85E-09 | 7.18E-08 | 1.15E-08 | 1.85E-09 | 1.85E-09 | 7.18E-08 | 7.18E-08 | 1.15E-08 | 1.15E-08 |
| BRS3007 | 5.19E-01 | 3.11E-01 | 5.19E-01 | 3.06E-02 | 5.19E-01 | 7.49E-02 | 1.00E+00 | 7.83E-01 | 3.11E-01 | 1.62E-01 | 1.85E-09 | 1.85E-09 | 7.18E-08 | 1.85E-09 | 1.85E-09 | 1.85E-09 | 4.59E-07 | 4.59E-07 | 1.85E-09 | 7.18E-08 |
| BRS3008 | 7.83E-01 | 9.65E-01 | 5.19E-01 | 1.12E-02 | 3.11E-01 | 3.06E-02 | 7.83E-01 | 1.00E+00 | 3.11E-01 | 1.62E-01 | 1.85E-09 | 1.85E-09 | 1.15E-08 | 1.85E-09 | 1.85E-09 | 1.85E-09 | 1.15E-08 | 1.15E-08 | 1.85E-09 | 1.85E-09 |
| BRS3009 | 1.62E-01 | 1.62E-01 | 5.19E-01 | 3.06E-02 | 5.19E-01 | 1.12E-02 | 3.11E-01 | 3.11E-01 | 1.00E+00 | 7.83E-01 | 1.85E-09 | 1.85E-09 | 7.18E-08 | 1.85E-09 | 1.85E-09 | 1.85E-09 | 7.18E-08 | 7.18E-08 | 1.85E-09 | 7.18E-08 |
| BRS3010 | 7.49E-02 | 7.49E-02 | 5.19E-01 | 3.06E-02 | 5.19E-01 | 1.12E-02 | 1.62E-01 | 1.62E-01 | 7.83E-01 | 1.00E+00 | 1.85E-09 | 1.85E-09 | 7.18E-08 | 1.85E-09 | 1.85E-09 | 1.85E-09 | 7.18E-08 | 4.59E-07 | 1.85E-09 | 7.18E-08 |
| Meta_BMS3001 | 1.85E-09 | 1.85E-09 | 1.85E-09 | 1.85E-09 | 1.85E-09 | 1.85E-09 | 1.85E-09 | 1.85E-09 | 1.85E-09 | 1.85E-09 | 1.00E+00 | 1.09E-03 | 1.09E-03 | 3.64E-03 | 9.65E-01 | 3.64E-03 | 1.39E-05 | 4.59E-07 | 7.49E-02 | 1.39E-05 |
| Meta_BMS3002 | 1.85E-09 | 1.85E-09 | 1.85E-09 | 1.85E-09 | 1.85E-09 | 1.85E-09 | 1.85E-09 | 1.85E-09 | 1.85E-09 | 1.85E-09 | 1.09E-03 | 1.00E+00 | 7.18E-08 | 2.64E-06 | 1.09E-03 | 7.83E-01 | 7.18E-08 | 1.15E-08 | 6.68E-05 | 1.15E-08 |
| Meta_BMS3003 | 1.85E-09 | 7.18E-08 | 2.64E-06 | 2.64E-06 | 1.15E-08 | 7.18E-08 | 7.18E-08 | 1.15E-08 | 7.18E-08 | 7.18E-08 | 1.09E-03 | 7.18E-08 | 1.00E+00 | 5.19E-01 | 3.64E-03 | 4.59E-07 | 5.19E-01 | 3.06E-02 | 3.11E-01 | 5.19E-01 |
| Meta_BMS3004 | 1.85E-09 | 1.85E-09 | 1.15E-08 | 1.15E-08 | 1.85E-09 | 1.15E-08 | 1.85E-09 | 1.85E-09 | 1.85E-09 | 1.85E-09 | 3.64E-03 | 2.64E-06 | 5.19E-01 | 1.00E+00 | 3.64E-03 | 6.68E-05 | 7.49E-02 | 1.09E-03 | 5.19E-01 | 7.49E-02 |
| Meta_BMS3005 | 1.85E-09 | 1.85E-09 | 1.85E-09 | 1.85E-09 | 1.85E-09 | 1.85E-09 | 1.85E-09 | 1.85E-09 | 1.85E-09 | 1.85E-09 | 9.65E-01 | 1.09E-03 | 3.64E-03 | 3.64E-03 | 1.00E+00 | 1.09E-03 | 1.39E-05 | 2.64E-06 | 1.62E-01 | 1.39E-05 |
| Meta_BMS3006 | 1.85E-09 | 1.85E-09 | 1.85E-09 | 1.85E-09 | 1.85E-09 | 1.85E-09 | 1.85E-09 | 1.85E-09 | 1.85E-09 | 1.85E-09 | 3.64E-03 | 7.83E-01 | 4.59E-07 | 6.68E-05 | 1.09E-03 | 1.00E+00 | 7.18E-08 | 1.15E-08 | 2.93E-04 | 1.15E-08 |
| Meta_BMS3007 | 1.85E-09 | 7.18E-08 | 2.64E-06 | 1.39E-05 | 1.15E-08 | 7.18E-08 | 4.59E-07 | 1.15E-08 | 7.18E-08 | 7.18E-08 | 1.39E-05 | 7.18E-08 | 5.19E-01 | 7.49E-02 | 1.39E-05 | 7.18E-08 | 1.00E+00 | 1.62E-01 | 3.64E-03 | 9.65E-01 |
| Meta_BMS3008 | 1.15E-08 | 7.18E-08 | 2.64E-06 | 2.64E-06 | 1.15E-08 | 7.18E-08 | 4.59E-07 | 1.15E-08 | 7.18E-08 | 4.59E-07 | 4.59E-07 | 1.15E-08 | 3.06E-02 | 1.09E-03 | 2.64E-06 | 1.15E-08 | 1.62E-01 | 1.00E+00 | 6.68E-05 | 3.11E-01 |
| Meta_BMS3009 | 1.85E-09 | 1.85E-09 | 1.15E-08 | 1.15E-08 | 1.15E-08 | 1.15E-08 | 1.85E-09 | 1.85E-09 | 1.85E-09 | 1.85E-09 | 7.49E-02 | 6.68E-05 | 3.11E-01 | 5.19E-01 | 1.62E-01 | 2.93E-04 | 3.64E-03 | 6.68E-05 | 1.00E+00 | 1.12E-02 |
| Meta_BMS3010 | 1.85E-09 | 1.15E-08 | 4.59E-07 | 4.59E-07 | 1.15E-08 | 1.15E-08 | 7.18E-08 | 1.85E-09 | 7.18E-08 | 7.18E-08 | 1.39E-05 | 1.15E-08 | 5.19E-01 | 7.49E-02 | 1.39E-05 | 1.15E-08 | 9.65E-01 | 3.11E-01 | 1.12E-02 | 1.00E+00 |
